# scHiCTools: a computational toolbox for analyzing single-cell Hi-C data

**DOI:** 10.1101/769513

**Authors:** Xinjun Li, Fan Feng, Wai Yan Leung, Jie Liu

**Affiliations:** Department of Statistics, University of Michigan, Ann Arbor, Michigan, United States of America; Department of Computational Medicine and Bioinformatics, University of Michigan, Ann Arbor, Michigan, United States of America

## Abstract

Single-cell Hi-C (scHi-C) sequencing technologies allow us to investigate three-dimensional chromatin organization at the single-cell level. However, we still need computational tools to deal with the sparsity of the contact maps from single cells and embed single cells in a lower-dimensional Euclidean space. This embedding helps us understand relationships between the cells in different dimensions such as cell-cycle dynamics and cell differentiation. Here, we present an open-source computational toolbox, **scHiCTools**, for analyzing single cell Hi-C data. The toolbox takes singlecell Hi-C data files as input, and projects single cells in a lower-dimensional Euclidean space. The toolbox includes three commonly used methods for smoothing scHi-C data (linear convolution, random walk, and network enhancing), three projection methods for embedding single cells (fastHiCRep, Selfish, and InnerProduct), three clustering methods for clustering cells (*k*-means, spectral clustering, and HiCluster) and a build-in function to visualize the cells embedding in a two-dimensional or three-dimensional plot. We benchmark the embedding performance and run time of these methods on a number of scHi-C datasets, and provide some suggestions for practice use. **scHiCTools**, based on Python3, can run on different platforms, including Linux, macOS, and Windows. Our software package is available at https://github.com/liu-bioinfo-lab/scHiCTools.

## Introduction

Single-cell Hi-C sequencing (scHi-C) technologies [10,11] allow us to understand chromatin organization dynamics and cell-to-cell heterogeneity, and connect many important genome research areas, including transcription regulation and epigenomics. However, interpretation of scHi-C data exposes several inherent data analysis challenges. Unlike RNA-seq data and ATAC-seq data, which are vectors of *m*-dimensional measures, Hi-C data are essentially symmetric matrices of *m×m*-dimensional pairwise measures, where the number of genomic loci *m* is usually more than tens of thousands, depending on the resolution of the contact maps. Due to the cost of scHi-C experiments, up to thousands of single cells are usually profiled, and the number of contacts in each cell ranges between a few thousands and hundreds of thousands. Therefore, scHi-C analysis also suffers from common high-dimensional challenges, including sparsity of the contact maps, batch effects, and sequencing noise. Thus, reasonable low-dimensional representation of scHi-C data is vital in scHi-C data analysis.

Previously, most similarity measures for comparing Hi-C contact matrices focus on bulk Hi-C data [19]. These methods [12,14,17,18] evaluate how likely two bulk Hi-C experiments are generated from the same biological sample. In a previous study [7], similarity methods have been applied to single-cell Hi-C data to evaluate similarity among *n* single cells. Those methods also couple with multidimensional scaling (MDS) to project these n single cells into a lower-dimensional Euclidean space. Among these methods, HiCRep [18] yields reasonable embedding of the single cells, but its *O*(*n*^2^) computational complexity makes it impractical when the number of cells is large.

In our **scHiCTools**, we implement a new “InnerProduct” approach to measure pairwise similarity among single cells. Our tool also implements a faster version of HiCRep, together with another Hi-C similarity measure named Selfish [1]. Among the three methods implemented, InnerProduct provides the most efficient way of embedding scHi-C data into a lower-dimensional space, and yields better clustering of the data, compared with existing scHi-C clustering method HiCluster [20]. All of the three approaches have *O*(*n*) computational complexity. Benchmarking experiments demonstrate that the new InnerProduct approach runs faster than the original HiCRep, and produces comparably accurate projection. To deal with the sparsity in scHi-C data, three smoothing approaches are implemented, including linear convolution, random walk, and network enhancing [16]. Among the three approaches, linear convolution appears to be most effective for smoothing sparse Hi-C contact maps from our experiments. In addition to the computational components, our toolbox supports different input file formats, summary diagnostic plots, and flexible projection plots. Our open-source toolbox, **scHiCTools**, as the first toolbox of such kind, can be useful for analyzing scHi-C data.

## Design and implementation

### Overview

Our **scHiCTools** implements commonly used approaches to analyze single cell Hi-C data. The core of the toolbox is a number of dimension reduction approaches which takes a number of single cells’ contact maps as input, and embeds the cells in a low-dimensional Euclidean space, so that the users can easily explore the heterogeneity among the cells. The toolbox also implements a number of built-in auxiliary functions for flexible and interactive visualization. The entire workflow of scHiCTools, illustrated in Figure 1, includes the following five steps: (1) reading single-cell data in .txt, .hic, or .cool format, generating summary diagnostic plots, and screening cells by their contact number and contact distance profile, (2) smoothing scHi-C contact maps using linear convolution, random walk, or network enhancing, (3) calculating pairwise similarity between the cells using fastHiCRep, InnerProduct, or Selfish, (4) embedding or clustering the cells in a lower dimensional space using dimensionality reduction methods, and (5) visualizing the two-dimensional or three-dimensional embedding in a scatter plot. We next describe the five steps in more details.

**Figure 1:**
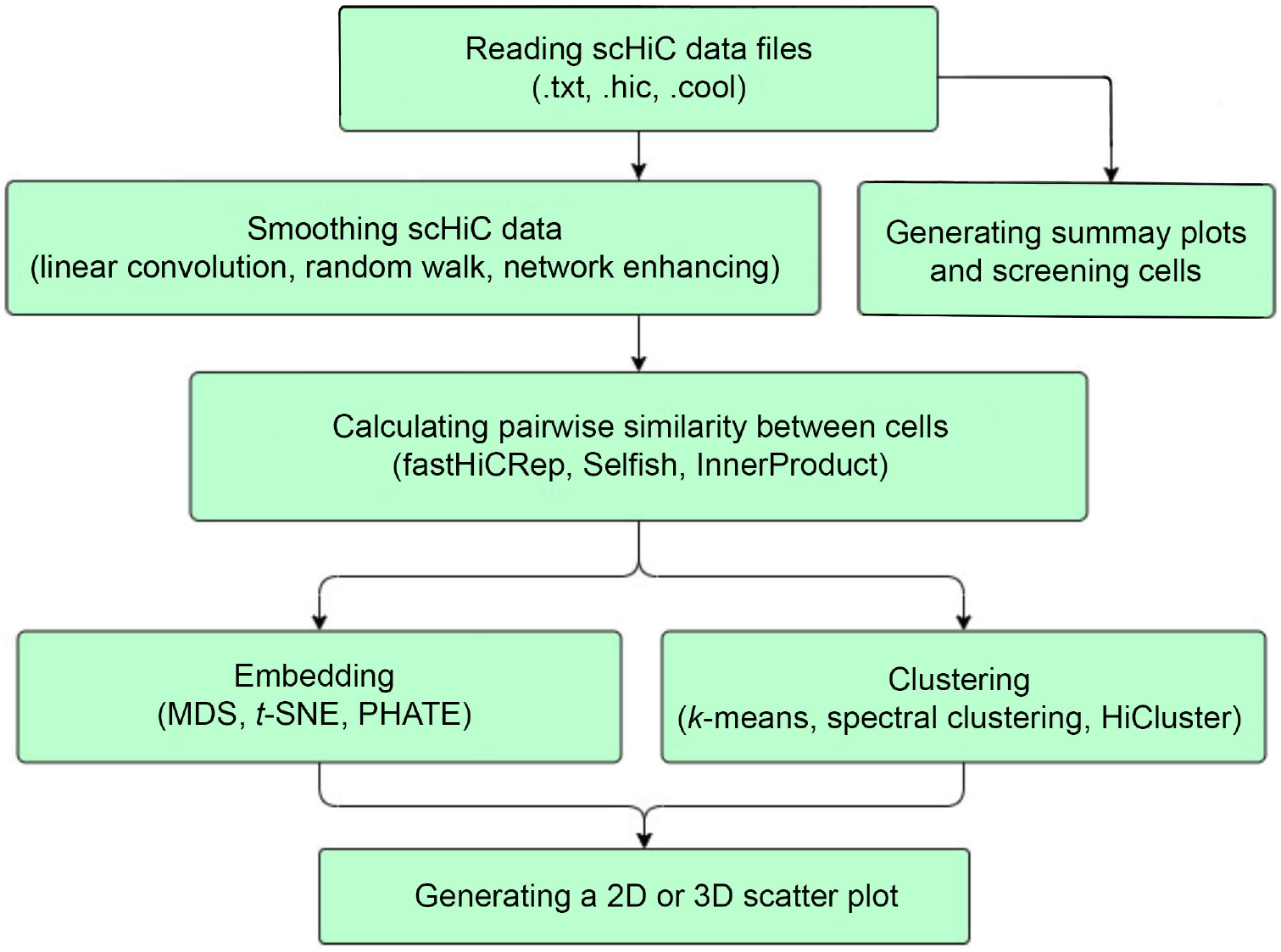
The workflow of scHiCTools. The workflow of scHiCTools includes the following five steps: (1) reading multiple files containing the single-cell data in .txt, .hic, or .cool format, generating the summary plots of the cells, and screening cells based on their contact number and contact distance profile, (2) smoothing the scHi-C contact maps using linear convolution, random walk, or network enhancing, (3) calculating the pairwise similarity between cells using fastHiCRep, InnerProduct, or Selfish, (4) embedding or clustering the cells in a lower dimensional space using dimensionality reduction methods, and (5) visualizing the two-dimensional or three-dimensional embedding in a scatter plot.

### Loading data and screening cells

Users can load scHi-C data in different file formats, including .hic files, .cool files, and sparse contact matrices in text files. When users choose to load sparse matrices in text files, they can customize the format of each column in the text files, and specify additional information including reference genome and the resolution of the contact maps. After loading the data files, **scHiCTools** allows users to plot two summary plots to examine the quality of the loaded single cell Hi-C data, namely a histogram of contact numbers in the cells, and a scatter plot of cells with the proportion of shortrange contacts (< 2 Mb) versus the proportion of the contacts at the mitotic band (2 – 12 Mb) (Fig. 2). A low-quality contact map usually shows a low number of contacts and a relatively high proportion of short-range contacts. Users can further screen the cells by thresholding the number of contacts and the proportion of short-range contacts in the cells.

**Figure 2:**
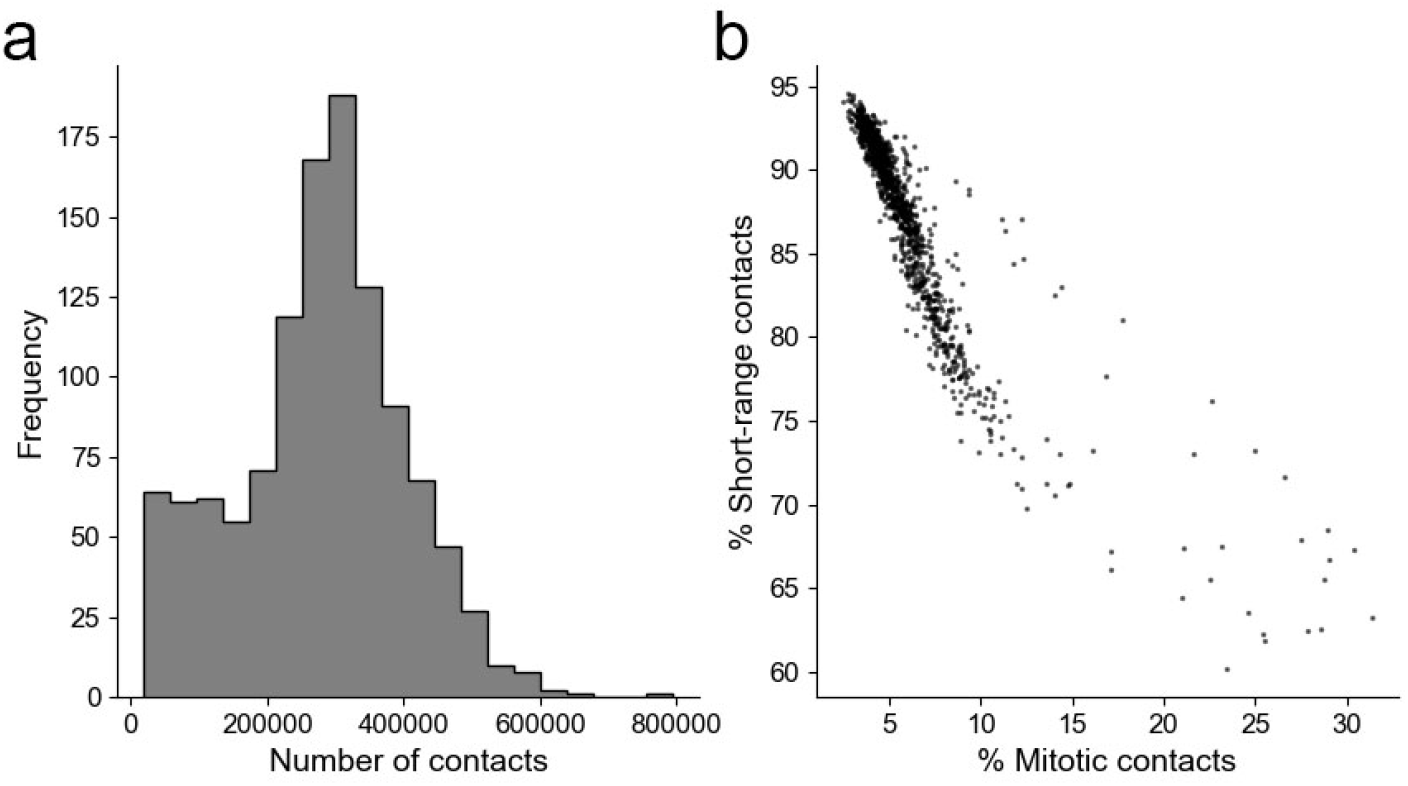
Summary plots for examining the quality of input scHi-C data. (a) A histogram of contact numbers in the individual cells. (b) A scatter plot showing the percentage of short-range contacts (< 2 Mb) versus the percentage of contacts at the mitotic band (2 – 12 Mb) in individual cells.

### Smoothing contact maps

Smoothing chromatin contact maps is necessary when they are sparse, especially in a comparative analysis of single cell contact maps. Our toolbox **scHiCTools** includes three smoothing approaches.

**Linear convolution** is essentially a two-dimensional convolution filter with equal weights in every position, which can be viewed as smoothing over neighboring bins in Hi-C contact maps. For example, original HiCRep [18] uses a parameter *h* to describe a (2*h* + 1)×(2*h* + 1) convolution filter, i.e., *h* = 1 indicating a 3×3 kernel with each element 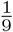. Because this approach is similar to reducing the resolution of the contact map, it is effective when contact maps are sparse.

**Random walk** [4] can also be used to smooth chromatin contact maps. Unlike linear convolution which takes information from neighbors on the contact map, random walk captures the signals from a global setting as follows. Let *W* be the input *m × m* contact matrix. Random walk updates *W* by *W*(*t*) = *W*^(*t*−1)^ · *B* in the *t*-th iteration, where 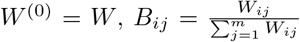. The matrix *B* is the input matrix divided by its row sum (i.e., every row of B sum up to 1). We can write B as *B* = *D*^-1^ · *W*, where 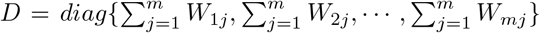. After *t* steps of update, *W*^(*t*)^ = *W · B^t^* is the output matrix after smoothing.

**Network enhancing** [16] is a special type of random walk which increases gaps between leading eigenvalues of a doubly stochastic contact matrix to enhance network signals. Because the random walk procedure needs to be applied on a doubly stochastic matrix (DSM, the sum of each row and the sum of each column are both 1), Knight-Ruiz (KR) normalization [6] is applied to the original contact matrix *W*, i.e., finding a vector *a*, such that *W*′ = *aWa^T^* is a DSM. Network enhancing makes the partition of contact maps more prominent, enhancing boundaries of topologically associated domains (TADs) in chromatin contact maps [16].

### Calculating pairwise similarity

Hi-C contact maps are *m × m*-dimensional measures, where *m* is the number of genomic loci. By calculating pairwise similarity among the cells, we avoid dealing with the high-dimensional measure in the following embedding and clustering steps. Our toolbox **scHiCTools** includes three approaches for calculating pairwise similarity among the single cells, as follows.

**InnerProduct** calculates the pairwise similarity matrix among cells in two steps. The first step is “scaling”, which directly sets the concatenated *z*-normalized first *s* strata as feature vectors. The second step is “multiplication”, which calculates an inner product of the feature vectors to obtain the similarity matrix of the single cells. InnerProduct starts with the calculation of the vector of strata. Denote *υ_i_* = [*υ*_*i*,1_, *υ*_*i*,2_,⋯, *υ*_*i,m*−1_]^*T*^ to be the *i*-th stratum of the chromosome, *i* from 1 to *s*. *υ_i,k_ = W_k,k+i_* is the *k*-th element of *υ_i_*, where *W* is the contact matrix of a chromosome. Then, *z*-normalization is applied to each *υ_i_* to get a zero-mean and unit-variance vector *υ_i_*. By concatenating all strata, the feature vector for each contact map is 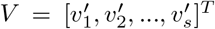. If we directly calculate the inner product of the two feature vectors *V_x_* and *V_y_* of map *x* and map *y*, then the inner product defines a kernel

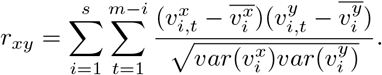

We can obtain a kernel matrix from each chromosome. Taking the average of *r_xy_* over all chromosomes keeps the matrix positive definite, and thus gives us an overall kernel matrix of all the cells in the input. However in practice, although taking the median may make the matrix no longer positive definite, it makes the similarity more stable and robust to noisy measurements.

**fastHiCRep** is a faster implementation of the original HiCRep approach [18]. Original HiCRep [18] calculates s stratum-adjusted correlation coefficients (SCCs) of the s strata near the diagonal of two contact maps, namely

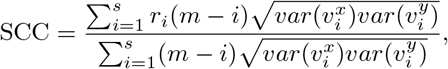

where *υ_i_* is the *i*-th stratum of the chromosome, *m − i* is the length of *υ_i_*, and *r_i_* is the Pearson’s correlation coefficient of 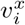 and 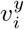. The length of *υ_i_* is *m − i*.

We have 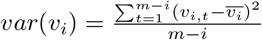, then

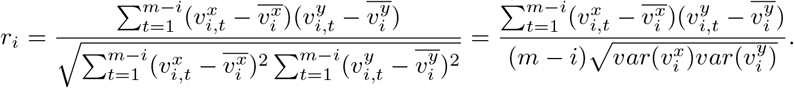

We can write SCC between contact maps *x* and *y* as a simple inner product of concatenated strata vectors divided by a constant related to variances and lengths of strata, namely

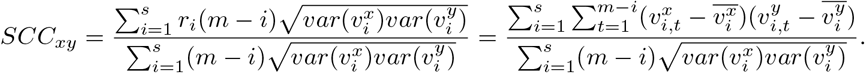

From above, the numerator is the inner product of all *s* strata (subtracted by its mean), and the denominator is a normalization factor, and therefore SCC can also be represented as an inner product. As a result, we are able to implement fastHiCRep, a fast version of HiCRep, by first presenting the contact maps as vectors and then calculating SCC as an inner product. Our fastHiCRep calculation reduces HiCRep’s computation complexity from *O*(*n*^2^) to *O*(*n*).

The third embedding approach **Selfish** [1] was recently proposed for bulk Hi-C comparative analysis. It first uses a sliding window to obtain a number of rectangular regions along the diagonal of the contact map, and then counts overall contact numbers in each region. Then, it generates a one-hot “fingerprint matrix” for each contact map. Gaussian kernels over the fingerprint matrices are finally calculated as similarities among the cells.

### Embedding and clustering

By embedding the cells in a lower-dimensional Euclidean space, we can easily explore and analyze the heterogeneity among the single cells. **scHiCTools** includes three different dimension reduction methods that convert the pairwise similarity matrices into embedding in a Euclidean space. The three dimension reduction methods are as follows.

**MDS** (Multidimensional scaling) takes in a pairwise distance matrix evaluated in the original space, and embeds the data points in a lower-dimensional space which preserves the pairwise distance matrix. In our package, we use the classical MDS which finds the *p* dimensional embedding 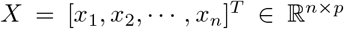 that minimizes the loss function: *loss* = ‖*XX^T^ − G*‖_*F*_, where 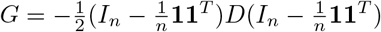, *D_ij_* is the distance between i-th and *j*-th cells, and **1** denotes a column vector of all ones.

*t***-SNE** [8] embeds high-dimensional data in a lower-dimensional space with an emphasis of preserving local neighborhood. *t*-SNE assumes data points *x*_1_,⋯,*x_n_* in the higher-dimensional space follow a Gaussian distribution, and the embedded points *y*_1_,⋯,*y_n_* follow a Student’s *t*-distribution. In the higher-dimensional space, the conditional probability of *x_i_* picking *x_j_* as its neighbor is 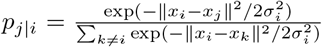. In our implementation, instead of computing the norm between two points, we directly use distances between two cells to calculate the similarity. The similarity between *x_i_* and *x_j_* is defined as the probability of picking *x_i_* and *x_j_* as neighbors, which is 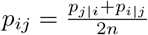. In the embedding space, since *y*_1_,⋯,*y_n_ ~ t*_1_, the probability of picking *y_i_* and *y_j_* as neighbors is 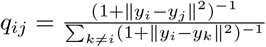. The Kullback–Leibler divergence of the distribution of data points from the distribution of embedded points is 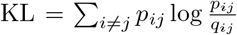. We choose embedding points which minimize KL-divergence.

**PHATE** (Potential of Heat-diffusion for Affinity-based Trajectory Embedding) [9] is a dimension reduction approach which emphasizes both local and global similarity. PHATE first calculates a local affinity matrix based on *k*-nearest neighbor distance *K_k_*, which captures the local structure of the data points. PHATE normalizes K_k_ using a Gaussian kernel to obtain a new matrix *P*, then diffuses *P* by *t* steps and gets a new matrix *P^t^* which preserves the global structure of data points. Based on matrix *P^t^*, we can find a potential representation of data *U_t_ = −log*(*P_t_*). Finally, PHATE applies non-metric MDS to *U_t_* to generate the lower-dimensional embedding.

If the user assumes that the single cells come from several clusters rather than a continuous manifold, then they can use a clustering method during the generation of the lower dimensional embedding. The following optional clustering methods are implemented in our toolbox.

*k***-means** assigns an observation to the cluster with the nearest cluster centroid, which is the mean of all observations belonging to the cluster. We use *k*-means++ [2] to initialize the centroids of clusters. Iterations of *k*-means algorithm renew the clusters by finding the points closest to the centroid in the last iteration, and update the centroid of each cluster by taking the mean of points in each cluster. Since *k*-means needs coordinates of cells in a Euclidean space to find the centroid of each cluster, we use MDS to embed the cells into a *l*-dimensional space first, and then perform *k*-means in the *l*-dimensional space.

**Spectral clustering** [15] takes in a distance matrix of data points and divides *n* observations into *k* clusters. Spectral clustering directly takes in the distance matrix and constructs a similarity graph with Gaussian similarity function. Based on the similarity graph, spectral clustering calculates the graph Laplacian of the similarity graph, and projects the points into a *k*-dimensional space based on the first *k* eigenvectors of the graph Laplacian matrix. Finally, it uses *k*-means to divide the data points into *k* group.

**HiCluster** [20] is a clustering method designed explicitly for single-cell Hi-C data. It uses convolution and random walk for smoothing and imputation, then converts the contact matrices to binary matrices. HiCluster conducts principal component analysis for embedding, and then *k*-means for clustering.

### Visualization of embedding

Users can plot the two-dimensional or three-dimensional embedding of cells using **scHiCTools**. Fig. 3 shows the scatter plots of the two-dimensional embedding of the cells in a cell cycle study [10]. Users can also visualize three-dimensional embedding of the cells if they assume the cells exists on a three-dimensional manifold (see Supplementary File 2). Users can also generate an interactive scatter plot of the cell embedding, in which the cell label is displayed when the user’s mouse hovers (see Supplementary 3). In order to use this feature, users need to install *ploty* on their device.

**Figure 3:**
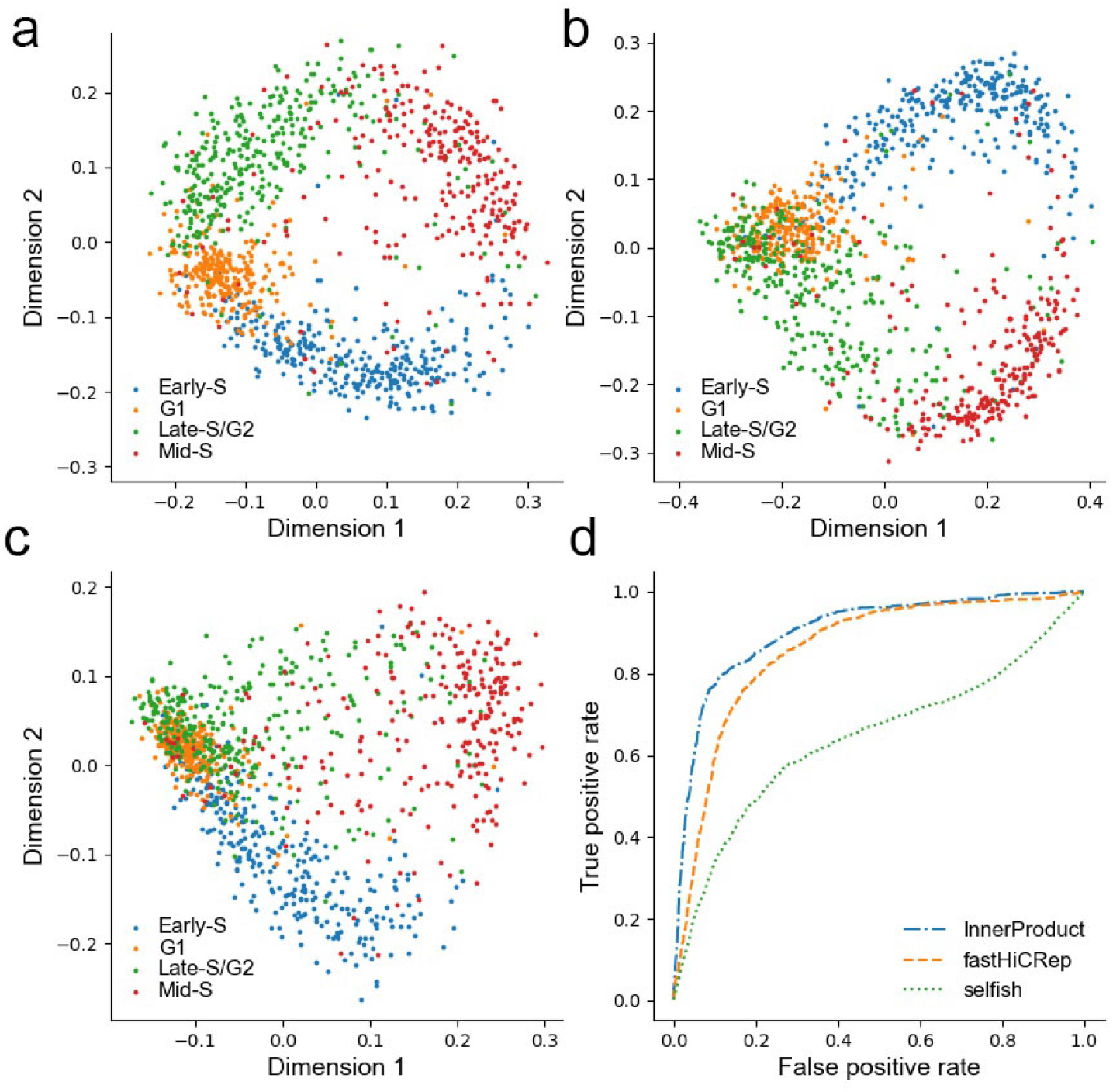
Two-dimensional scatter plots of the embedding from the three methods that calculate the similarity between contact matrices, including InnerProduct, fastHiCRep and Selfish. (Dataset: Nagano et al., 2017). (a) Two-dimensional projection from InnerProduct and MDS shows a clear circular pattern along the four stages of cell cycle. (b) Two-dimensional projection using fastHiCRep and MDS does not show clear separation between the four stages of cell cycle. (c) Two-dimensional projection using Selfish and MDS does not show clear separation between the four stages of cell cycle. (d) Evaluating the three embedding methods in a cell-cycle phasing task by ACROC. The ACROC values from InnerProduct, fastHiCRep and Selfish are 0.904, 0.858 and 0.642, respectively.

## Results

In this section, we apply the toolbox on a number of scHi-C datasets [3, 10], and benchmark the performance when different smoothing, embedding, and clustering methods are used. In addition to plotting the two-dimensional embedding of cells and examining whether the embedding is sensible, we benchmark the projection performance on a scHi-C dataset [10] with the average area under the curve of a circular ROC calculation (ACROC) proposed in a recent work [7]. We record the run time of these methods on the scHi-C dataset [10] to compare the efficiency of different methods. We evaluate the clustering performance with two evaluation criteria: normalized mutual information (NMI) [13] and adjusted rand index (ARI) [5] on a recent scHi-C dataset [3]. We have the following observations.

### InnerProduct is effective for calculating the pairwise similarity among single-cell Hi-C contact maps

When InnerProduct is coupled with MDS, it produces an accurate projection of single cells along the four stages of cell cycle (Fig. 3a), achieving an average area under the curve of a circular ROC calculation (ACROC) of 0.904, which is as good as original HiCRep reported in the recent work [7]. However, two-dimensional scatter plots from fastHiCRep and Selfish show a circular pattern along the four stages of cell cycle, but the separation between the stages is not as clear as that from InnerProduct (Fig 3b and Fig 3c). The ACROC measures from fastHiCRep is 0.858 and the ACROC measures from Selfish is 0.642, which are both lower than that from InnerProduct (Fig. 3d).

### PHATE and *t*-SNE produce satisfactory projection

Since MDS recovers global pairwise distance in its projection, it is unclear whether methods that preserve local pairwise distance (i.e., PHATE and *t*-SNE) can produce a better projection. Fig. 4a and Fig. 4b show that PHATE and *t*-SNE are suitable embedding methods that project the smoothed contact maps into a lower-dimensional space while preserving a cell-cycle pattern. The ACROC measure of PHATE is 0.920, which is better than *t*-SNE (0.901) and MDS (0.904). Therefore, PHATE and *t*-SNE can be used alternatively, especially when faraway neighbors’ similarity cannot be properly evaluated.

**Figure 4:**
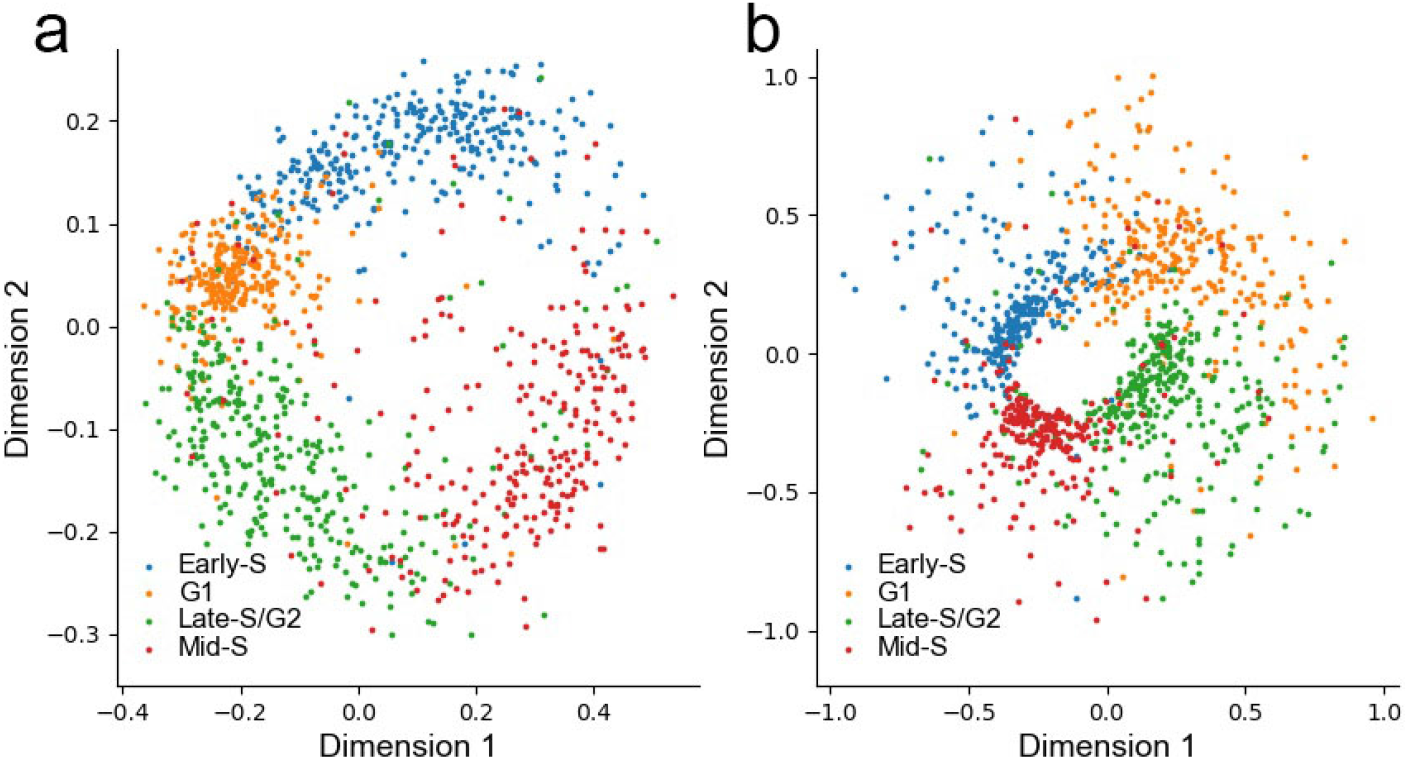
Two-dimensional embedding using different dimension reduction methods. (Dataset: Nagano et al., 2017). (a) Two-dimensional projection from InnerProduct/PHATE shows a circular pattern similiar to MDS projection (ACROC: 0.920). (b) Two-dimensional projection from InnerProduct/*t*-SNE shows a circular pattern (ACROC: 0.901).

### All three embedding methods are computationally efficient

The run time of the three methods is compared in Table 1. For embedding 500 cells randomly selected from data from Nagano et al. [10], three methods in our package finish within minutes. Given the fact that all of the three embedding approaches have an *O*(*n*) computation complexity, they scale up very well for a large number of cells. FastHiCRep is slightly faster than InnerProduct, which is faster than Selfish under the default parameters. In Table 1, we also report the run time from the two steps: “scaling” and “multiplication” from InnerProduct separately. It is observed that the “scaling” step in which the feature vector is normalized contributes to most of the run time. In contrast, the “multiplication” step, including calculating the inner product of two feature vectors, does not contribute much to the run time.

**Table 1:** Run time (in seconds) of different methods as the number of cells varies. (Dataset: Nagano et al., 2017)

| # cells | HiCRep | fastHiCRep | InnerProduct |  |  | Selfish |
| --- | --- | --- | --- | --- | --- | --- |
|  |  |  | Scaling | Multiplication | Total |  |
| 100 | 323.26 | 5.30 | 4.71 | 0.59 | 5.30 | 11.47 |
| 200 | 1303.59 | 10.25 | 8.89 | 1.82 | 10.71 | 24.26 |
| 300 | 2938.93 | 17.67 | 14.65 | 3.72 | 18.37 | 40.30 |
| 400 | 5224.05 | 21.67 | 21.10 | 5.46 | 26.56 | 59.56 |
| 500 | 8156.38 | 28.44 | 25.98 | 7.99 | 33.97 | 84.95 |

### Linear convolution smoothing and random walk improve projection at high dropout rates

We examine the three smoothing approaches when they are coupled with InnerProduct on a scHi-C dataset [10] which is sparsified with two different methods. The first sparsification method is to remove 40% ~ 99.9% of the contacts randomly from all genomic positions (i.e., contact number ranging from ~200,000 to ~500 in each cell). The second sparsification method is to simulate dropout events in sequencing data which discard contacts from 5% ~ 60% of the genomic loci. Fig. 5a shows that none of the three smoothing methods, including linear convolution, random walk, and network enhancing, improve the projection performance when the down-sampling was used. The ACROC decreases from around 0.9 to around 0.6 as down-sampling rate ranges from 1 to 0.05. Fig. 5b shows that under the second sparsification method, linear convolution and random walk showed some consistent improvement. At a high dropout rate (0.7), linear convolution keeps the average ACROC above 0.9, whereas other methods’ average ACROC’s drop to below 0.9. Therefore, we recommend users use linear convolution to smooth scHi-C contact maps if they suspect dropout events exist moderately in their scHi-C data.

**Figure 5:**
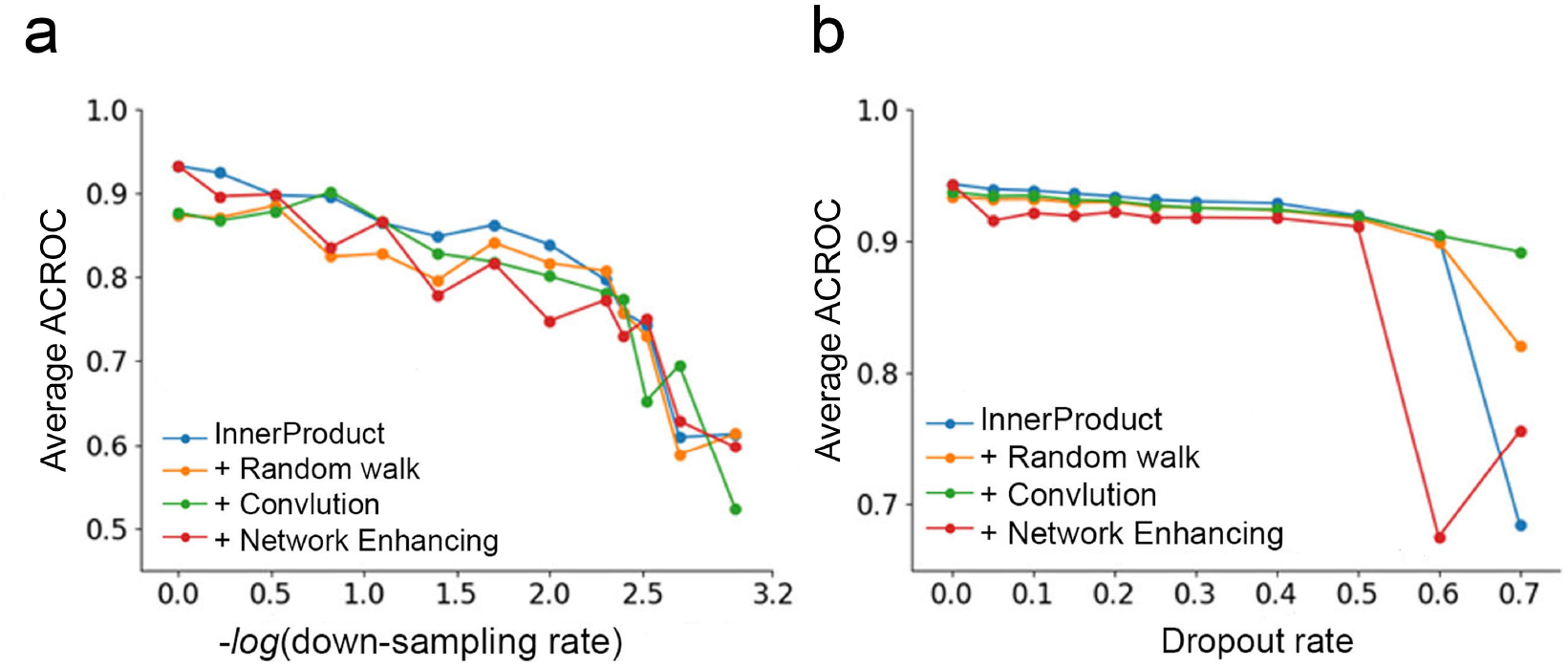
Average ACROC measures from InnerProduct without any smoothing, InnerProduct with random walk smoothing, InnerProduct with linear convolution, and InnerProduct with network enhancing. (a) When the dataset is sparsified with the first sparsification method, ACROC values from the four approaches decrease when down-sampling rate increases. (b) When the dataset is sparsified with the second sparsification method, ACROC values from the four approaches decrease when down-sampling rate increases, but at a high dropout rate (0.7), linear convolution’s ACROC remains high, whereas other methods’ ACROC drops to below 0.9.

### HiCluster produces better clustering results than InnerProduct coupled with spectral clustering or with *k*-means, but less efficient

Table 2 shows the performance of different clustering methods implemented in **scHiCTools** on the dataset of Collombet et al., 2020 [3]. We use all 750 mouse embryo cells at five differentiation stages in the study of Collombet et al., including the 1-cell, 2-cell, 4-cell, 8-cell, and 64-cell stages. We separate the cells into five clusters and evaluate the clusters using two evaluation measures, including normalized mutual information (NMI) [13], and adjusted rand index (ARI) [5]. HiCluster shows better NMI and ARI values than InnerProduct with *k*-means and InnerProduct with spectral clustering. InnerProduct with *k*-means and InnerProduct with spectral clustering are more efficient than HiCluster. The run time of HiCluster is 98.635 seconds, which is much longer than InnerProduct with *k*-means (1.414 seconds) and InnerProduct with spectral clustering (1.435 seconds).

**Table 2:** The normalized mutual information (NMI), adjusted rand index (ARI), and run time of three clustering approaches. (Data: 750 embryo cells at five differentiation stages, including 1-cell, 2-cell, 4-cell, 8-cell and 64-cell stages, Collombet et al., 2020)

|  | NMI | ARI | Run time (in seconds) |
| --- | --- | --- | --- |
| HiCluster | 0.266 | 0.259 | 98.635 |
| InnerProduct+MDS+$k$-means | 0.222 | 0.208 | 1.414 |
| InnerProduct+spectral clustering | 0.241 | 0.197 | 1.435 |

## Availability and future directions

Our scHiCTools is implemented in Python. The source code is available and maintained at Github: https://github.com/liu-bioinfo-lab/scHiCTools. This package is also available on PyPI python package manager. The current code runs under Python 3.7 or newer versions. Other dependency includes numpy, scipy, matplotlib, pandas, simplejson, six, and h5py. For the interactive scatter plot function, you need to have plotly installed. In the future, we will keep updating the toolbox with new scHi-C analysis algorithms, including new embedding methods such as UMAP and new clustering methods such as hierarchical clustering.

## Supporting information

**S1 File. Plots for different similarity measures and embedding methods applied to Nagano single-cell dataset.** This zip file contains the plots of other combination of similarity measures (InnerProduct, fastHiCRep and Selfish) and embedding methods (MDS, *t*-SNE and PHATE).

(ZIP)

**S2 File. Three-dimensional scatter plots.** This zip file contains the 3D scatter plots of different embedding methods (MDS, *t*-SNE and PHATE) applied to Nagano single-cell dataset. (ZIP)

**S3 File. Interactive plots.** This file contiains an example of interactive 2D scatter plot and an example of interactive 3D scatter plot of cells from Nagano et al. (ZIP)

**S4 File. scHiCTools source code, documentation and test dataset.** This zip file is a clone of scHiCTools public Git repository. To install from this file rather than from PyPI, see the readme file and follow installation instructions. (ZIP)

