## Supplementary figures and images for "scHiCTools: a computational toolbox for analyzing single-cell Hi-C data"

### file.png

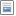

### HiCRep+MDS 3D.png

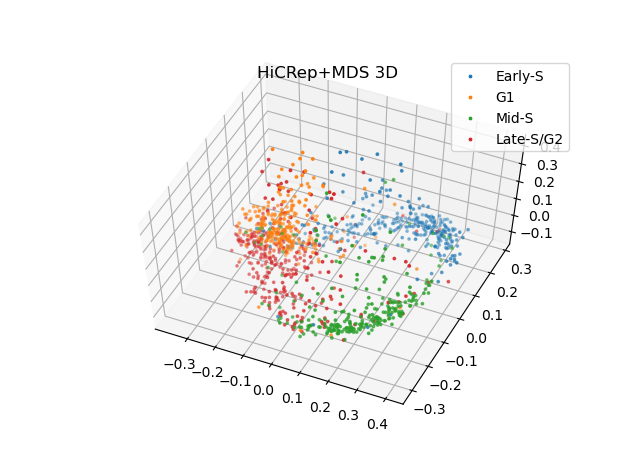

### HiCRep+MDS.png

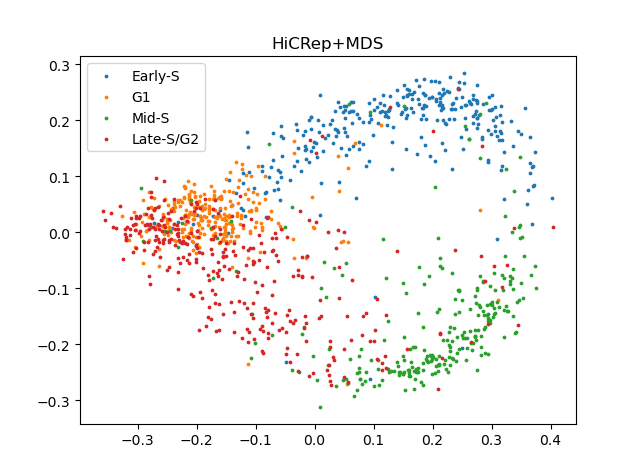

### HiCRep+PHATE 3D.png

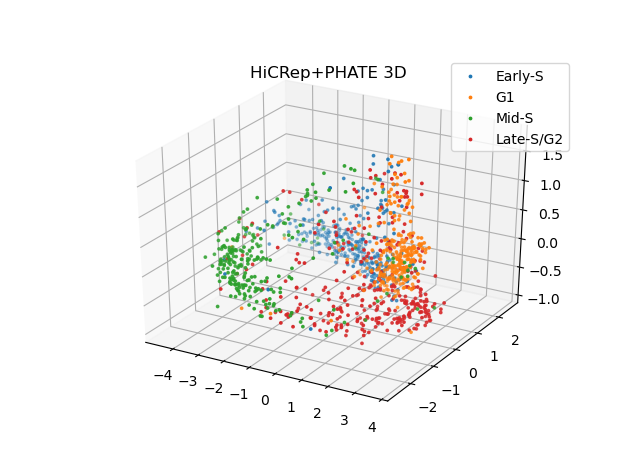

### HiCRep+PHATE.png

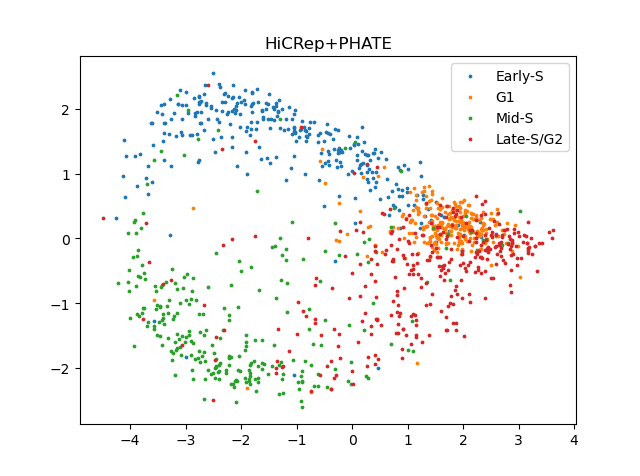

### HiCRep+tSNE 3D.png

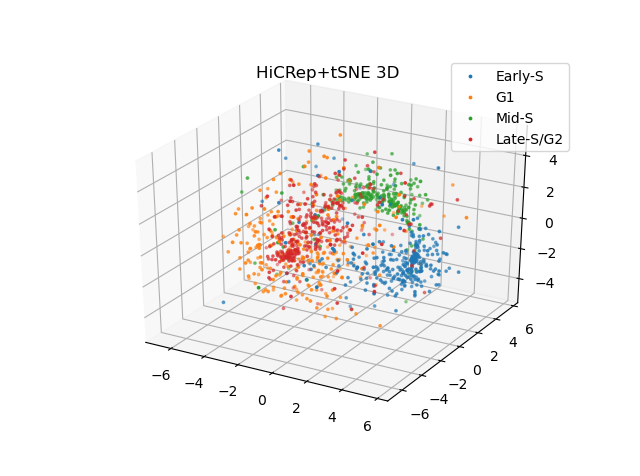

### HiCRep+tSNE.png

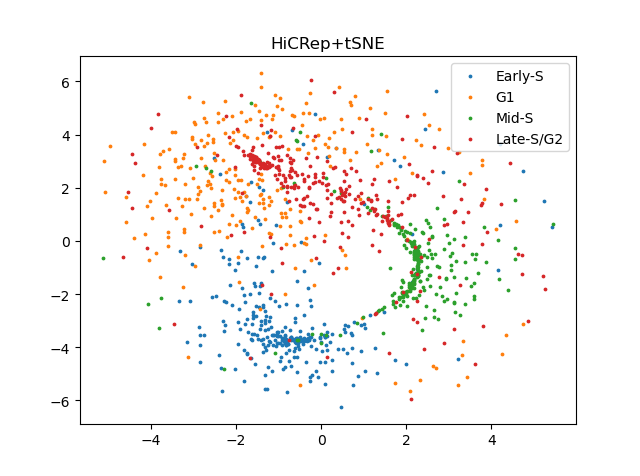

### InnerProduct+MDS.png

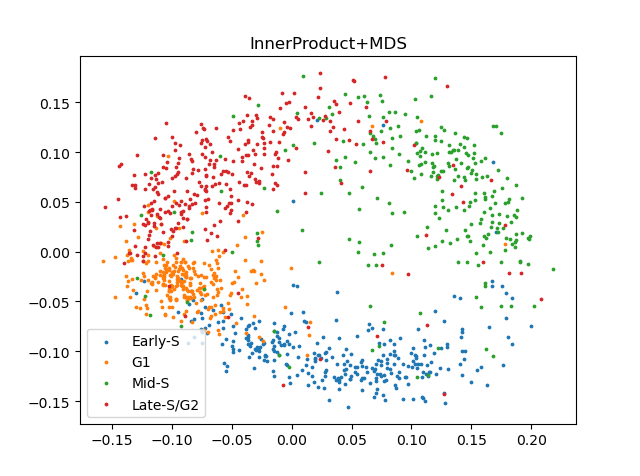

### InnerProduct+MDS_3D.png

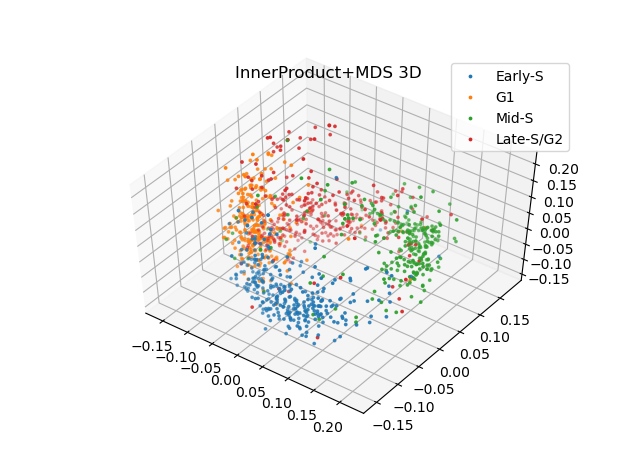

### InnerProduct+PHATE 3D.png

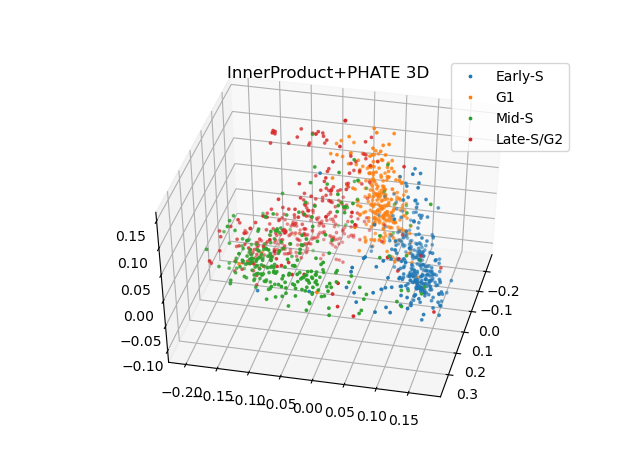

### InnerProduct+PHATE.png

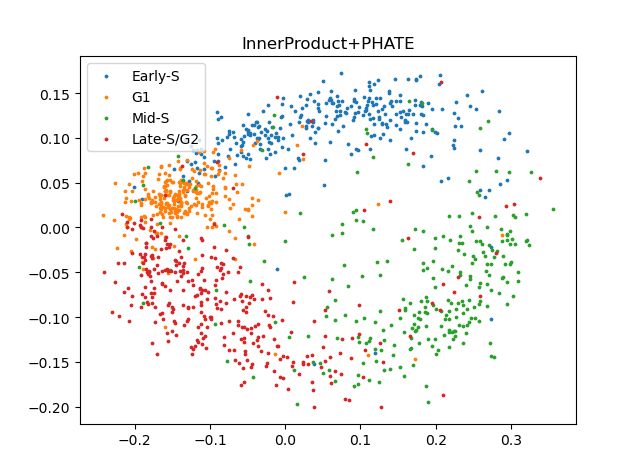

### InnerProduct+tSNE 3D.png

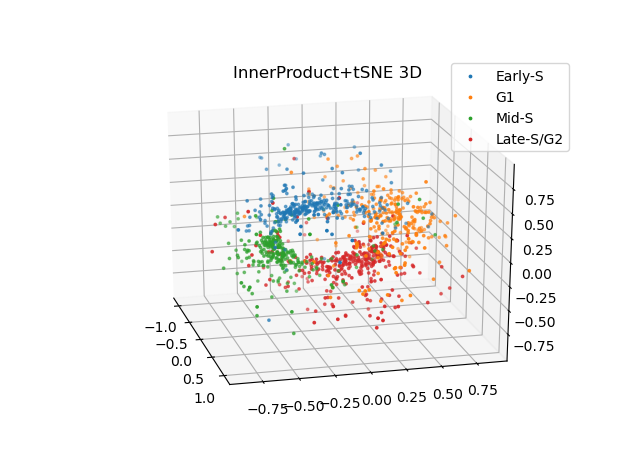

### InnerProduct+tSNE.png

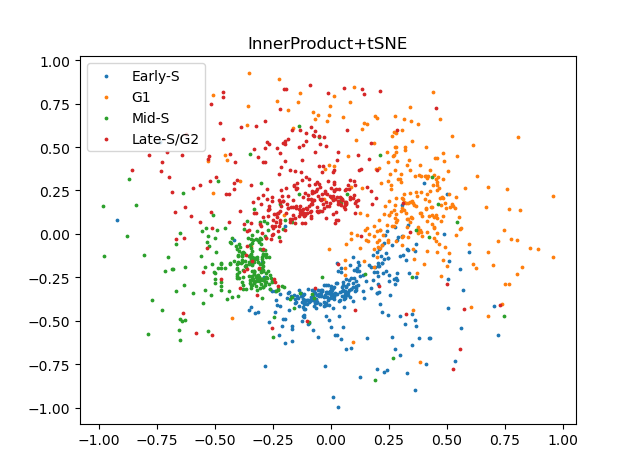

### minus.png

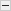

### plus.png

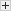

### Selfish+MDS 3D.png

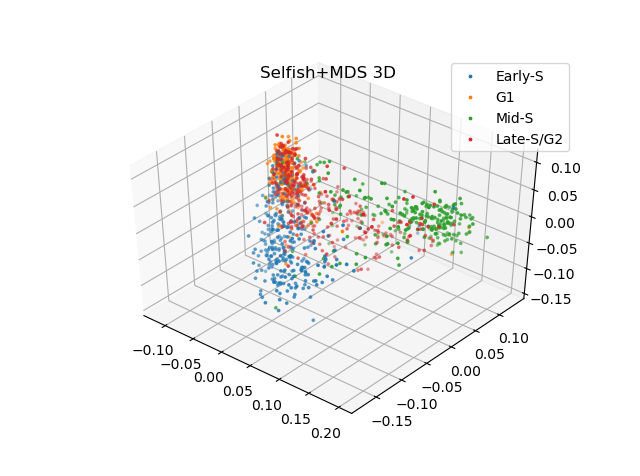

### Selfish+MDS.png

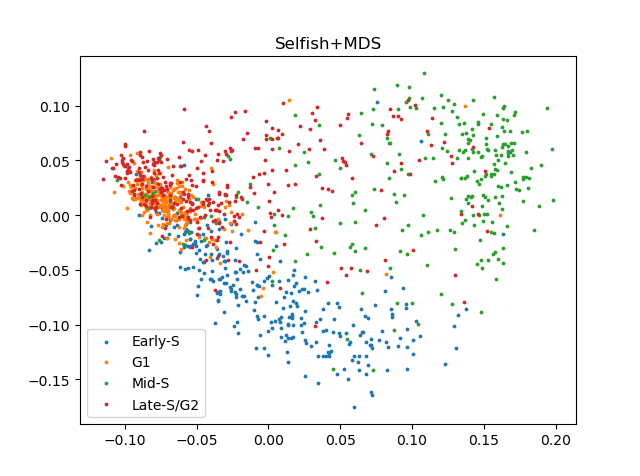

### Selfish+PHATE 3D.png

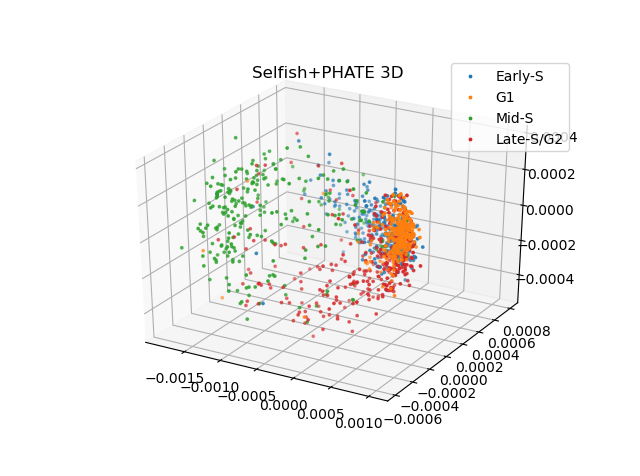

### Selfish+PHATE.png

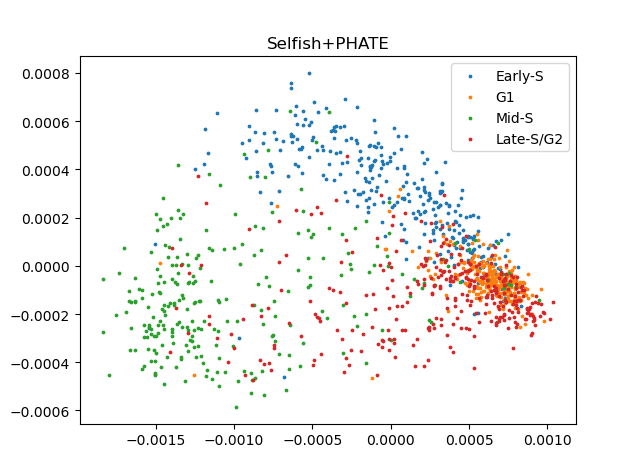

### Selfish+tSNE 3D.png

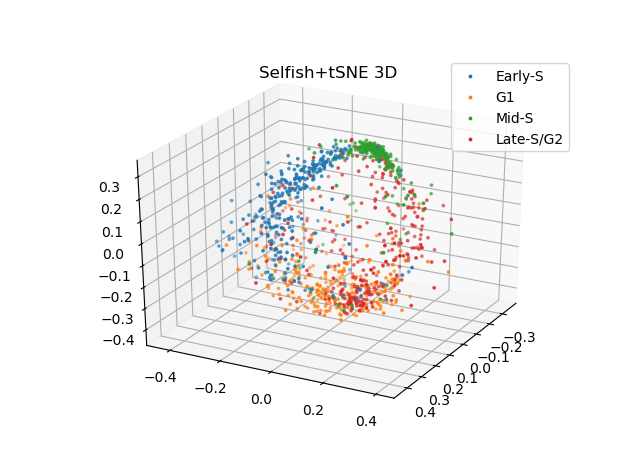

### Selfish+tSNE.png

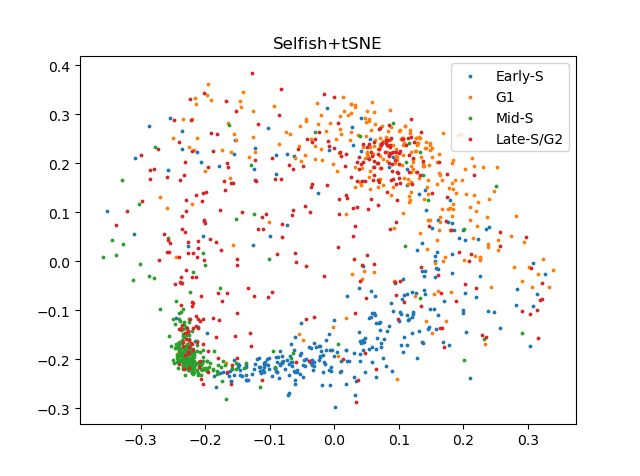
