## Supplementary material for "scHiCTools: a computational toolbox for analyzing single-cell Hi-C data": S4: index.rst.txt

.. scHiCTools documentation master file, created by
   sphinx-quickstart on Sun May 31 19:56:09 2020.
   You can adapt this file completely to your liking, but it should at least
   contain the root `toctree` directive.

Welcome to scHiCTools's documentation!
======================================

.. toctree::
   :maxdepth: 2
   :caption: Contents:


Indices and tables
==================

* :ref:`genindex`
* :ref:`modindex`
* :ref:`search`

Introduction
============

scHiCTools is a computational toolbox for analyzing single cell Hi-C (high-throughput sequencing for 3C) data which includes functions for:

1. Load single-cell HiC datasets
2. Smoothing the contact maps with linear convolution, random walk or network enhancing
3. Calculating embeddings for single cell HiC datasets efficiently with reproducibility measures include InnerProduct, HiCRep and Selfish


Setup procedure
===============
