## Supplementary material for "scHiCTools: a computational toolbox for analyzing single-cell Hi-C data": S4: doctools.js

```
/*
 * doctools.js
 * ~~~~~~~~~~~
 *
 * Sphinx JavaScript utilities for all documentation.
 *
 * :copyright: Copyright 2007-2020 by the Sphinx team, see AUTHORS.
 * :license: BSD, see LICENSE for details.
 *
 */

/**
 * select a different prefix for underscore
 */
$u = _.noConflict();

/**
 * make the code below compatible with browsers without
 * an installed firebug like debugger
if (!window.console || !console.firebug) {
  var names = ["log", "debug", "info", "warn", "error", "assert", "dir",
    "dirxml", "group", "groupEnd", "time", "timeEnd", "count", "trace",
    "profile", "profileEnd"];
  window.console = {};
  for (var i = 0; i < names.length; ++i)
    window.console[names[i]] = function() {};
}
 */

/**
 * small helper function to urldecode strings
 */
jQuery.urldecode = function(x) {
  return decodeURIComponent(x).replace(/\+/g, ' ');
};

/**
 * small helper function to urlencode strings
 */
jQuery.urlencode = encodeURIComponent;

/**
 * This function returns the parsed url parameters of the
 * current request. Multiple values per key are supported,
 * it will always return arrays of strings for the value parts.
 */
jQuery.getQueryParameters = function(s) {
  if (typeof s === 'undefined')
    s = document.location.search;
  var parts = s.substr(s.indexOf('?') + 1).split('&');
  var result = {};
  for (var i = 0; i < parts.length; i++) {
    var tmp = parts[i].split('=', 2);
    var key = jQuery.urldecode(tmp[0]);
    var value = jQuery.urldecode(tmp[1]);
    if (key in result)
      result[key].push(value);
    else
      result[key] = [value];
  }
  return result;
};

/**
 * highlight a given string on a jquery object by wrapping it in
 * span elements with the given class name.
 */
jQuery.fn.highlightText = function(text, className) {
  function highlight(node, addItems) {
    if (node.nodeType === 3) {
      var val = node.nodeValue;
      var pos = val.toLowerCase().indexOf(text);
      if (pos >= 0 &&
          !jQuery(node.parentNode).hasClass(className) &&
          !jQuery(node.parentNode).hasClass("nohighlight")) {
        var span;
        var isInSVG = jQuery(node).closest("body, svg, foreignObject").is("svg");
        if (isInSVG) {
          span = document.createElementNS("http://www.w3.org/2000/svg", "tspan");
        } else {
          span = document.createElement("span");
          span.className = className;
        }
        span.appendChild(document.createTextNode(val.substr(pos, text.length)));
        node.parentNode.insertBefore(span, node.parentNode.insertBefore(
          document.createTextNode(val.substr(pos + text.length)),
          node.nextSibling));
        node.nodeValue = val.substr(0, pos);
        if (isInSVG) {
          var rect = document.createElementNS("http://www.w3.org/2000/svg", "rect");
          var bbox = node.parentElement.getBBox();
          rect.x.baseVal.value = bbox.x;
          rect.y.baseVal.value = bbox.y;
          rect.width.baseVal.value = bbox.width;
          rect.height.baseVal.value = bbox.height;
          rect.setAttribute('class', className);
          addItems.push({
              "parent": node.parentNode,
              "target": rect});
        }
      }
    }
    else if (!jQuery(node).is("button, select, textarea")) {
      jQuery.each(node.childNodes, function() {
        highlight(this, addItems);
      });
    }
  }
  var addItems = [];
  var result = this.each(function() {
    highlight(this, addItems);
  });
  for (var i = 0; i < addItems.length; ++i) {
    jQuery(addItems[i].parent).before(addItems[i].target);
  }
  return result;
};

/*
 * backward compatibility for jQuery.browser
 * This will be supported until firefox bug is fixed.
 */
if (!jQuery.browser) {
  jQuery.uaMatch = function(ua) {
    ua = ua.toLowerCase();

    var match = /(chrome)[ \/]([\w.]+)/.exec(ua) ||
      /(webkit)[ \/]([\w.]+)/.exec(ua) ||
      /(opera)(?:.*version|)[ \/]([\w.]+)/.exec(ua) ||
      /(msie) ([\w.]+)/.exec(ua) ||
      ua.indexOf("compatible") < 0 && /(mozilla)(?:.*? rv:([\w.]+)|)/.exec(ua) ||
      [];

    return {
      browser: match[ 1 ] || "",
      version: match[ 2 ] || "0"
    };
  };
  jQuery.browser = {};
  jQuery.browser[jQuery.uaMatch(navigator.userAgent).browser] = true;
}

/**
 * Small JavaScript module for the documentation.
 */
var Documentation = {

  init : function() {
    this.fixFirefoxAnchorBug();
    this.highlightSearchWords();
    this.initIndexTable();
    if (DOCUMENTATION_OPTIONS.NAVIGATION_WITH_KEYS) {
      this.initOnKeyListeners();
    }
  },

  /**
   * i18n support
   */
  TRANSLATIONS : {},
  PLURAL_EXPR : function(n) { return n === 1 ? 0 : 1; },
  LOCALE : 'unknown',

  // gettext and ngettext don't access this so that the functions
  // can safely bound to a different name (_ = Documentation.gettext)
  gettext : function(string) {
    var translated = Documentation.TRANSLATIONS[string];
    if (typeof translated === 'undefined')
      return string;
    return (typeof translated === 'string') ? translated : translated[0];
  },

  ngettext : function(singular, plural, n) {
    var translated = Documentation.TRANSLATIONS[singular];
    if (typeof translated === 'undefined')
      return (n == 1) ? singular : plural;
    return translated[Documentation.PLURALEXPR(n)];
  },

  addTranslations : function(catalog) {
    for (var key in catalog.messages)
      this.TRANSLATIONS[key] = catalog.messages[key];
    this.PLURAL_EXPR = new Function('n', 'return +(' + catalog.plural_expr + ')');
    this.LOCALE = catalog.locale;
  },

  /**
   * add context elements like header anchor links
   */
  addContextElements : function() {
    $('div[id] > :header:first').each(function() {
      $('<a class="headerlink">\u00B6</a>').
      attr('href', '#' + this.id).
      attr('title', _('Permalink to this headline')).
      appendTo(this);
    });
    $('dt[id]').each(function() {
      $('<a class="headerlink">\u00B6</a>').
      attr('href', '#' + this.id).
      attr('title', _('Permalink to this definition')).
      appendTo(this);
    });
  },

  /**
   * workaround a firefox stupidity
   * see: https://bugzilla.mozilla.org/show_bug.cgi?id=645075
   */
  fixFirefoxAnchorBug : function() {
    if (document.location.hash && $.browser.mozilla)
      window.setTimeout(function() {
        document.location.href += '';
      }, 10);
  },

  /**
   * highlight the search words provided in the url in the text
   */
  highlightSearchWords : function() {
    var params = $.getQueryParameters();
    var terms = (params.highlight) ? params.highlight[0].split(/\s+/) : [];
    if (terms.length) {
      var body = $('div.body');
      if (!body.length) {
        body = $('body');
      }
      window.setTimeout(function() {
        $.each(terms, function() {
          body.highlightText(this.toLowerCase(), 'highlighted');
        });
      }, 10);
      $('<p class="highlight-link"><a href="javascript:Documentation.' +
        'hideSearchWords()">' + _('Hide Search Matches') + '</a></p>')
          .appendTo($('#searchbox'));
    }
  },

  /**
   * init the domain index toggle buttons
   */
  initIndexTable : function() {
    var togglers = $('img.toggler').click(function() {
      var src = $(this).attr('src');
      var idnum = $(this).attr('id').substr(7);
      $('tr.cg-' + idnum).toggle();
      if (src.substr(-9) === 'minus.png')
        $(this).attr('src', src.substr(0, src.length-9) + 'plus.png');
      else
        $(this).attr('src', src.substr(0, src.length-8) + 'minus.png');
    }).css('display', '');
    if (DOCUMENTATION_OPTIONS.COLLAPSE_INDEX) {
        togglers.click();
    }
  },

  /**
   * helper function to hide the search marks again
   */
  hideSearchWords : function() {
    $('#searchbox .highlight-link').fadeOut(300);
    $('span.highlighted').removeClass('highlighted');
  },

  /**
   * make the url absolute
   */
  makeURL : function(relativeURL) {
    return DOCUMENTATION_OPTIONS.URL_ROOT + '/' + relativeURL;
  },

  /**
   * get the current relative url
   */
  getCurrentURL : function() {
    var path = document.location.pathname;
    var parts = path.split(/\//);
    $.each(DOCUMENTATION_OPTIONS.URL_ROOT.split(/\//), function() {
      if (this === '..')
        parts.pop();
    });
    var url = parts.join('/');
    return path.substring(url.lastIndexOf('/') + 1, path.length - 1);
  },

  initOnKeyListeners: function() {
    $(document).keydown(function(event) {
      var activeElementType = document.activeElement.tagName;
      // don't navigate when in search box or textarea
      if (activeElementType !== 'TEXTAREA' && activeElementType !== 'INPUT' && activeElementType !== 'SELECT'
          && !event.altKey && !event.ctrlKey && !event.metaKey && !event.shiftKey) {
        switch (event.keyCode) {
          case 37: // left
            var prevHref = $('link[rel="prev"]').prop('href');
            if (prevHref) {
              window.location.href = prevHref;
              return false;
            }
          case 39: // right
            var nextHref = $('link[rel="next"]').prop('href');
            if (nextHref) {
              window.location.href = nextHref;
              return false;
            }
        }
      }
    });
  }
};

// quick alias for translations
_ = Documentation.gettext;

$(document).ready(function() {
  Documentation.init();
});
```
