## Supplementary material for "scHiCTools: a computational toolbox for analyzing single-cell Hi-C data": S4: jquery.js

```
/*! jQuery v3.5.1 | (c) JS Foundation and other contributors | jquery.org/license */
!function(e,t){"use strict";"object"==typeof module&&"object"==typeof module.exports?module.exports=e.document?t(e,!0):function(e){if(!e.document)throw new Error("jQuery requires a window with a document");return t(e)}:t(e)}("undefined"!=typeof window?window:this,function(C,e){"use strict";var t=[],r=Object.getPrototypeOf,s=t.slice,g=t.flat?function(e){return t.flat.call(e)}:function(e){return t.concat.apply([],e)},u=t.push,i=t.indexOf,n={},o=n.toString,v=n.hasOwnProperty,a=v.toString,l=a.call(Object),y={},m=function(e){return"function"==typeof e&&"number"!=typeof e.nodeType},x=function(e){return null!=e&&e===e.window},E=C.document,c={type:!0,src:!0,nonce:!0,noModule:!0};function b(e,t,n){var r,i,o=(n=n||E).createElement("script");if(o.text=e,t)for(r in c)(i=t[r]||t.getAttribute&&t.getAttribute(r))&&o.setAttribute(r,i);n.head.appendChild(o).parentNode.removeChild(o)}function w(e){return null==e?e+"":"object"==typeof e||"function"==typeof e?n[o.call(e)]||"object":typeof e}var f="3.5.1",S=function(e,t){return new S.fn.init(e,t)};function p(e){var t=!!e&&"length"in e&&e.length,n=w(e);return!m(e)&&!x(e)&&("array"===n||0===t||"number"==typeof t&&0<t&&t-1 in e)}S.fn=S.prototype={jquery:f,constructor:S,length:0,toArray:function(){return s.call(this)},get:function(e){return null==e?s.call(this):e<0?this[e+this.length]:this[e]},pushStack:function(e){var t=S.merge(this.constructor(),e);return t.prevObject=this,t},each:function(e){return S.each(this,e)},map:function(n){return this.pushStack(S.map(this,function(e,t){return n.call(e,t,e)}))},slice:function(){return this.pushStack(s.apply(this,arguments))},first:function(){return this.eq(0)},last:function(){return this.eq(-1)},even:function(){return this.pushStack(S.grep(this,function(e,t){return(t+1)%2}))},odd:function(){return this.pushStack(S.grep(this,function(e,t){return t%2}))},eq:function(e){var t=this.length,n=+e+(e<0?t:0);return this.pushStack(0<=n&&n<t?[this[n]]:[])},end:function(){return this.prevObject||this.constructor()},push:u,sort:t.sort,splice:t.splice},S.extend=S.fn.extend=function(){var e,t,n,r,i,o,a=arguments[0]||{},s=1,u=arguments.length,l=!1;for("boolean"==typeof a&&(l=a,a=arguments[s]||{},s++),"object"==typeof a||m(a)||(a={}),s===u&&(a=this,s--);s<u;s++)if(null!=(e=arguments[s]))for(t in e)r=e[t],"__proto__"!==t&&a!==r&&(l&&r&&(S.isPlainObject(r)||(i=Array.isArray(r)))?(n=a[t],o=i&&!Array.isArray(n)?[]:i||S.isPlainObject(n)?n:{},i=!1,a[t]=S.extend(l,o,r)):void 0!==r&&(a[t]=r));return a},S.extend({expando:"jQuery"+(f+Math.random()).replace(/\D/g,""),isReady:!0,error:function(e){throw new Error(e)},noop:function(){},isPlainObject:function(e){var t,n;return!(!e||"[object Object]"!==o.call(e))&&(!(t=r(e))||"function"==typeof(n=v.call(t,"constructor")&&t.constructor)&&a.call(n)===l)},isEmptyObject:function(e){var t;for(t in e)return!1;return!0},globalEval:function(e,t,n){b(e,{nonce:t&&t.nonce},n)},each:function(e,t){var n,r=0;if(p(e)){for(n=e.length;r<n;r++)if(!1===t.call(e[r],r,e[r]))break}else for(r in e)if(!1===t.call(e[r],r,e[r]))break;return e},makeArray:function(e,t){var n=t||[];return null!=e&&(p(Object(e))?S.merge(n,"string"==typeof e?[e]:e):u.call(n,e)),n},inArray:function(e,t,n){return null==t?-1:i.call(t,e,n)},merge:function(e,t){for(var n=+t.length,r=0,i=e.length;r<n;r++)e[i++]=t[r];return e.length=i,e},grep:function(e,t,n){for(var r=[],i=0,o=e.length,a=!n;i<o;i++)!t(e[i],i)!==a&&r.push(e[i]);return r},map:function(e,t,n){var r,i,o=0,a=[];if(p(e))for(r=e.length;o<r;o++)null!=(i=t(e[o],o,n))&&a.push(i);else for(o in e)null!=(i=t(e[o],o,n))&&a.push(i);return g(a)},guid:1,support:y}),"function"==typeof Symbol&&(S.fn[Symbol.iterator]=t[Symbol.iterator]),S.each("Boolean Number String Function Array Date RegExp Object Error Symbol".split(" "),function(e,t){n["[object "+t+"]"]=t.toLowerCase()});var d=function(n){var e,d,b,o,i,h,f,g,w,u,l,T,C,a,E,v,s,c,y,S="sizzle"+1*new Date,p=n.document,k=0,r=0,m=ue(),x=ue(),A=ue(),N=ue(),D=function(e,t){return e===t&&(l=!0),0},j={}.hasOwnProperty,t=[],q=t.pop,L=t.push,H=t.push,O=t.slice,P=function(e,t){for(var n=0,r=e.length;n<r;n++)if(e[n]===t)return n;return-1},R="checked|selected|async|autofocus|autoplay|controls|defer|disabled|hidden|ismap|loop|multiple|open|readonly|required|scoped",M="[\\x20\\t\\r\\n\\f]",I="(?:\\\\[\\da-fA-F]{1,6}"+M+"?|\\\\[^\\r\\n\\f]|[\\w-]|[^\0-\\x7f])+",W="\\["+M+"*("+I+")(?:"+M+"*([*^$|!~]?=)"+M+"*(?:'((?:\\\\.|[^\\\\'])*)'|\"((?:\\\\.|[^\\\\\"])*)\"|("+I+"))|)"+M+"*\\]",F=":("+I+")(?:\\((('((?:\\\\.|[^\\\\'])*)'|\"((?:\\\\.|[^\\\\\"])*)\")|((?:\\\\.|[^\\\\()[\\]]|"+W+")*)|.*)\\)|)",B=new RegExp(M+"+","g"),$=new RegExp("^"+M+"+|((?:^|[^\\\\])(?:\\\\.)*)"+M+"+$","g"),_=new RegExp("^"+M+"*,"+M+"*"),z=new RegExp("^"+M+"*([>+~]|"+M+")"+M+"*"),U=new RegExp(M+"|>"),X=new RegExp(F),V=new RegExp("^"+I+"$"),G={ID:new RegExp("^#("+I+")"),CLASS:new RegExp("^\\.("+I+")"),TAG:new RegExp("^("+I+"|[*])"),ATTR:new RegExp("^"+W),PSEUDO:new RegExp("^"+F),CHILD:new RegExp("^:(only|first|last|nth|nth-last)-(child|of-type)(?:\\("+M+"*(even|odd|(([+-]|)(\\d*)n|)"+M+"*(?:([+-]|)"+M+"*(\\d+)|))"+M+"*\\)|)","i"),bool:new RegExp("^(?:"+R+")$","i"),needsContext:new RegExp("^"+M+"*[>+~]|:(even|odd|eq|gt|lt|nth|first|last)(?:\\("+M+"*((?:-\\d)?\\d*)"+M+"*\\)|)(?=[^-]|$)","i")},Y=/HTML$/i,Q=/^(?:input|select|textarea|button)$/i,J=/^h\d$/i,K=/^[^{]+\{\s*\[native \w/,Z=/^(?:#([\w-]+)|(\w+)|\.([\w-]+))$/,ee=/[+~]/,te=new RegExp("\\\\[\\da-fA-F]{1,6}"+M+"?|\\\\([^\\r\\n\\f])","g"),ne=function(e,t){var n="0x"+e.slice(1)-65536;return t||(n<0?String.fromCharCode(n+65536):String.fromCharCode(n>>10|55296,1023&n|56320))},re=/([\0-\x1f\x7f]|^-?\d)|^-$|[^\0-\x1f\x7f-\uFFFF\w-]/g,ie=function(e,t){return t?"\0"===e?"\ufffd":e.slice(0,-1)+"\\"+e.charCodeAt(e.length-1).toString(16)+" ":"\\"+e},oe=function(){T()},ae=be(function(e){return!0===e.disabled&&"fieldset"===e.nodeName.toLowerCase()},{dir:"parentNode",next:"legend"});try{H.apply(t=O.call(p.childNodes),p.childNodes),t[p.childNodes.length].nodeType}catch(e){H={apply:t.length?function(e,t){L.apply(e,O.call(t))}:function(e,t){var n=e.length,r=0;while(e[n++]=t[r++]);e.length=n-1}}}function se(t,e,n,r){var i,o,a,s,u,l,c,f=e&&e.ownerDocument,p=e?e.nodeType:9;if(n=n||[],"string"!=typeof t||!t||1!==p&&9!==p&&11!==p)return n;if(!r&&(T(e),e=e||C,E)){if(11!==p&&(u=Z.exec(t)))if(i=u[1]){if(9===p){if(!(a=e.getElementById(i)))return n;if(a.id===i)return n.push(a),n}else if(f&&(a=f.getElementById(i))&&y(e,a)&&a.id===i)return n.push(a),n}else{if(u[2])return H.apply(n,e.getElementsByTagName(t)),n;if((i=u[3])&&d.getElementsByClassName&&e.getElementsByClassName)return H.apply(n,e.getElementsByClassName(i)),n}if(d.qsa&&!N[t+" "]&&(!v||!v.test(t))&&(1!==p||"object"!==e.nodeName.toLowerCase())){if(c=t,f=e,1===p&&(U.test(t)||z.test(t))){(f=ee.test(t)&&ye(e.parentNode)||e)===e&&d.scope||((s=e.getAttribute("id"))?s=s.replace(re,ie):e.setAttribute("id",s=S)),o=(l=h(t)).length;while(o--)l[o]=(s?"#"+s:":scope")+" "+xe(l[o]);c=l.join(",")}try{return H.apply(n,f.querySelectorAll(c)),n}catch(e){N(t,!0)}finally{s===S&&e.removeAttribute("id")}}}return g(t.replace($,"$1"),e,n,r)}function ue(){var r=[];return function e(t,n){return r.push(t+" ")>b.cacheLength&&delete e[r.shift()],e[t+" "]=n}}function le(e){return e[S]=!0,e}function ce(e){var t=C.createElement("fieldset");try{return!!e(t)}catch(e){return!1}finally{t.parentNode&&t.parentNode.removeChild(t),t=null}}function fe(e,t){var n=e.split("|"),r=n.length;while(r--)b.attrHandle[n[r]]=t}function pe(e,t){var n=t&&e,r=n&&1===e.nodeType&&1===t.nodeType&&e.sourceIndex-t.sourceIndex;if(r)return r;if(n)while(n=n.nextSibling)if(n===t)return-1;return e?1:-1}function de(t){return function(e){return"input"===e.nodeName.toLowerCase()&&e.type===t}}function he(n){return function(e){var t=e.nodeName.toLowerCase();return("input"===t||"button"===t)&&e.type===n}}function ge(t){return function(e){return"form"in e?e.parentNode&&!1===e.disabled?"label"in e?"label"in e.parentNode?e.parentNode.disabled===t:e.disabled===t:e.isDisabled===t||e.isDisabled!==!t&&ae(e)===t:e.disabled===t:"label"in e&&e.disabled===t}}function ve(a){return le(function(o){return o=+o,le(function(e,t){var n,r=a([],e.length,o),i=r.length;while(i--)e[n=r[i]]&&(e[n]=!(t[n]=e[n]))})})}function ye(e){return e&&"undefined"!=typeof e.getElementsByTagName&&e}for(e in d=se.support={},i=se.isXML=function(e){var t=e.namespaceURI,n=(e.ownerDocument||e).documentElement;return!Y.test(t||n&&n.nodeName||"HTML")},T=se.setDocument=function(e){var t,n,r=e?e.ownerDocument||e:p;return r!=C&&9===r.nodeType&&r.documentElement&&(a=(C=r).documentElement,E=!i(C),p!=C&&(n=C.defaultView)&&n.top!==n&&(n.addEventListener?n.addEventListener("unload",oe,!1):n.attachEvent&&n.attachEvent("onunload",oe)),d.scope=ce(function(e){return a.appendChild(e).appendChild(C.createElement("div")),"undefined"!=typeof e.querySelectorAll&&!e.querySelectorAll(":scope fieldset div").length}),d.attributes=ce(function(e){return e.className="i",!e.getAttribute("className")}),d.getElementsByTagName=ce(function(e){return e.appendChild(C.createComment("")),!e.getElementsByTagName("*").length}),d.getElementsByClassName=K.test(C.getElementsByClassName),d.getById=ce(function(e){return a.appendChild(e).id=S,!C.getElementsByName||!C.getElementsByName(S).length}),d.getById?(b.filter.ID=function(e){var t=e.replace(te,ne);return function(e){return e.getAttribute("id")===t}},b.find.ID=function(e,t){if("undefined"!=typeof t.getElementById&&E){var n=t.getElementById(e);return n?[n]:[]}}):(b.filter.ID=function(e){var n=e.replace(te,ne);return function(e){var t="undefined"!=typeof e.getAttributeNode&&e.getAttributeNode("id");return t&&t.value===n}},b.find.ID=function(e,t){if("undefined"!=typeof t.getElementById&&E){var n,r,i,o=t.getElementById(e);if(o){if((n=o.getAttributeNode("id"))&&n.value===e)return[o];i=t.getElementsByName(e),r=0;while(o=i[r++])if((n=o.getAttributeNode("id"))&&n.value===e)return[o]}return[]}}),b.find.TAG=d.getElementsByTagName?function(e,t){return"undefined"!=typeof t.getElementsByTagName?t.getElementsByTagName(e):d.qsa?t.querySelectorAll(e):void 0}:function(e,t){var n,r=[],i=0,o=t.getElementsByTagName(e);if("*"===e){while(n=o[i++])1===n.nodeType&&r.push(n);return r}return o},b.find.CLASS=d.getElementsByClassName&&function(e,t){if("undefined"!=typeof t.getElementsByClassName&&E)return t.getElementsByClassName(e)},s=[],v=[],(d.qsa=K.test(C.querySelectorAll))&&(ce(function(e){var t;a.appendChild(e).innerHTML="<a id='"+S+"'></a><select id='"+S+"-\r\\' msallowcapture=''><option selected=''></option></select>",e.querySelectorAll("[msallowcapture^='']").length&&v.push("[*^$]="+M+"*(?:''|\"\")"),e.querySelectorAll("[selected]").length||v.push("\\["+M+"*(?:value|"+R+")"),e.querySelectorAll("[id~="+S+"-]").length||v.push("~="),(t=C.createElement("input")).setAttribute("name",""),e.appendChild(t),e.querySelectorAll("[name='']").length||v.push("\\["+M+"*name"+M+"*="+M+"*(?:''|\"\")"),e.querySelectorAll(":checked").length||v.push(":checked"),e.querySelectorAll("a#"+S+"+*").length||v.push(".#.+[+~]"),e.querySelectorAll("\\\f"),v.push("[\\r\\n\\f]")}),ce(function(e){e.innerHTML="<a href='' disabled='disabled'></a><select disabled='disabled'><option/></select>";var t=C.createElement("input");t.setAttribute("type","hidden"),e.appendChild(t).setAttribute("name","D"),e.querySelectorAll("[name=d]").length&&v.push("name"+M+"*[*^$|!~]?="),2!==e.querySelectorAll(":enabled").length&&v.push(":enabled",":disabled"),a.appendChild(e).disabled=!0,2!==e.querySelectorAll(":disabled").length&&v.push(":enabled",":disabled"),e.querySelectorAll("*,:x"),v.push(",.*:")})),(d.matchesSelector=K.test(c=a.matches||a.webkitMatchesSelector||a.mozMatchesSelector||a.oMatchesSelector||a.msMatchesSelector))&&ce(function(e){d.disconnectedMatch=c.call(e,"*"),c.call(e,"[s!='']:x"),s.push("!=",F)}),v=v.length&&new RegExp(v.join("|")),s=s.length&&new RegExp(s.join("|")),t=K.test(a.compareDocumentPosition),y=t||K.test(a.contains)?function(e,t){var n=9===e.nodeType?e.documentElement:e,r=t&&t.parentNode;return e===r||!(!r||1!==r.nodeType||!(n.contains?n.contains(r):e.compareDocumentPosition&&16&e.compareDocumentPosition(r)))}:function(e,t){if(t)while(t=t.parentNode)if(t===e)return!0;return!1},D=t?function(e,t){if(e===t)return l=!0,0;var n=!e.compareDocumentPosition-!t.compareDocumentPosition;return n||(1&(n=(e.ownerDocument||e)==(t.ownerDocument||t)?e.compareDocumentPosition(t):1)||!d.sortDetached&&t.compareDocumentPosition(e)===n?e==C||e.ownerDocument==p&&y(p,e)?-1:t==C||t.ownerDocument==p&&y(p,t)?1:u?P(u,e)-P(u,t):0:4&n?-1:1)}:function(e,t){if(e===t)return l=!0,0;var n,r=0,i=e.parentNode,o=t.parentNode,a=[e],s=[t];if(!i||!o)return e==C?-1:t==C?1:i?-1:o?1:u?P(u,e)-P(u,t):0;if(i===o)return pe(e,t);n=e;while(n=n.parentNode)a.unshift(n);n=t;while(n=n.parentNode)s.unshift(n);while(a[r]===s[r])r++;return r?pe(a[r],s[r]):a[r]==p?-1:s[r]==p?1:0}),C},se.matches=function(e,t){return se(e,null,null,t)},se.matchesSelector=function(e,t){if(T(e),d.matchesSelector&&E&&!N[t+" "]&&(!s||!s.test(t))&&(!v||!v.test(t)))try{var n=c.call(e,t);if(n||d.disconnectedMatch||e.document&&11!==e.document.nodeType)return n}catch(e){N(t,!0)}return 0<se(t,C,null,[e]).length},se.contains=function(e,t){return(e.ownerDocument||e)!=C&&T(e),y(e,t)},se.attr=function(e,t){(e.ownerDocument||e)!=C&&T(e);var n=b.attrHandle[t.toLowerCase()],r=n&&j.call(b.attrHandle,t.toLowerCase())?n(e,t,!E):void 0;return void 0!==r?r:d.attributes||!E?e.getAttribute(t):(r=e.getAttributeNode(t))&&r.specified?r.value:null},se.escape=function(e){return(e+"").replace(re,ie)},se.error=function(e){throw new Error("Syntax error, unrecognized expression: "+e)},se.uniqueSort=function(e){var t,n=[],r=0,i=0;if(l=!d.detectDuplicates,u=!d.sortStable&&e.slice(0),e.sort(D),l){while(t=e[i++])t===e[i]&&(r=n.push(i));while(r--)e.splice(n[r],1)}return u=null,e},o=se.getText=function(e){var t,n="",r=0,i=e.nodeType;if(i){if(1===i||9===i||11===i){if("string"==typeof e.textContent)return e.textContent;for(e=e.firstChild;e;e=e.nextSibling)n+=o(e)}else if(3===i||4===i)return e.nodeValue}else while(t=e[r++])n+=o(t);return n},(b=se.selectors={cacheLength:50,createPseudo:le,match:G,attrHandle:{},find:{},relative:{">":{dir:"parentNode",first:!0}," ":{dir:"parentNode"},"+":{dir:"previousSibling",first:!0},"~":{dir:"previousSibling"}},preFilter:{ATTR:function(e){return e[1]=e[1].replace(te,ne),e[3]=(e[3]||e[4]||e[5]||"").replace(te,ne),"~="===e[2]&&(e[3]=" "+e[3]+" "),e.slice(0,4)},CHILD:function(e){return e[1]=e[1].toLowerCase(),"nth"===e[1].slice(0,3)?(e[3]||se.error(e[0]),e[4]=+(e[4]?e[5]+(e[6]||1):2*("even"===e[3]||"odd"===e[3])),e[5]=+(e[7]+e[8]||"odd"===e[3])):e[3]&&se.error(e[0]),e},PSEUDO:function(e){var t,n=!e[6]&&e[2];return G.CHILD.test(e[0])?null:(e[3]?e[2]=e[4]||e[5]||"":n&&X.test(n)&&(t=h(n,!0))&&(t=n.indexOf(")",n.length-t)-n.length)&&(e[0]=e[0].slice(0,t),e[2]=n.slice(0,t)),e.slice(0,3))}},filter:{TAG:function(e){var t=e.replace(te,ne).toLowerCase();return"*"===e?function(){return!0}:function(e){return e.nodeName&&e.nodeName.toLowerCase()===t}},CLASS:function(e){var t=m[e+" "];return t||(t=new RegExp("(^|"+M+")"+e+"("+M+"|$)"))&&m(e,function(e){return t.test("string"==typeof e.className&&e.className||"undefined"!=typeof e.getAttribute&&e.getAttribute("class")||"")})},ATTR:function(n,r,i){return function(e){var t=se.attr(e,n);return null==t?"!="===r:!r||(t+="","="===r?t===i:"!="===r?t!==i:"^="===r?i&&0===t.indexOf(i):"*="===r?i&&-1<t.indexOf(i):"$="===r?i&&t.slice(-i.length)===i:"~="===r?-1<(" "+t.replace(B," ")+" ").indexOf(i):"|="===r&&(t===i||t.slice(0,i.length+1)===i+"-"))}},CHILD:function(h,e,t,g,v){var y="nth"!==h.slice(0,3),m="last"!==h.slice(-4),x="of-type"===e;return 1===g&&0===v?function(e){return!!e.parentNode}:function(e,t,n){var r,i,o,a,s,u,l=y!==m?"nextSibling":"previousSibling",c=e.parentNode,f=x&&e.nodeName.toLowerCase(),p=!n&&!x,d=!1;if(c){if(y){while(l){a=e;while(a=a[l])if(x?a.nodeName.toLowerCase()===f:1===a.nodeType)return!1;u=l="only"===h&&!u&&"nextSibling"}return!0}if(u=[m?c.firstChild:c.lastChild],m&&p){d=(s=(r=(i=(o=(a=c)[S]||(a[S]={}))[a.uniqueID]||(o[a.uniqueID]={}))[h]||[])[0]===k&&r[1])&&r[2],a=s&&c.childNodes[s];while(a=++s&&a&&a[l]||(d=s=0)||u.pop())if(1===a.nodeType&&++d&&a===e){i[h]=[k,s,d];break}}else if(p&&(d=s=(r=(i=(o=(a=e)[S]||(a[S]={}))[a.uniqueID]||(o[a.uniqueID]={}))[h]||[])[0]===k&&r[1]),!1===d)while(a=++s&&a&&a[l]||(d=s=0)||u.pop())if((x?a.nodeName.toLowerCase()===f:1===a.nodeType)&&++d&&(p&&((i=(o=a[S]||(a[S]={}))[a.uniqueID]||(o[a.uniqueID]={}))[h]=[k,d]),a===e))break;return(d-=v)===g||d%g==0&&0<=d/g}}},PSEUDO:function(e,o){var t,a=b.pseudos[e]||b.setFilters[e.toLowerCase()]||se.error("unsupported pseudo: "+e);return a[S]?a(o):1<a.length?(t=[e,e,"",o],b.setFilters.hasOwnProperty(e.toLowerCase())?le(function(e,t){var n,r=a(e,o),i=r.length;while(i--)e[n=P(e,r[i])]=!(t[n]=r[i])}):function(e){return a(e,0,t)}):a}},pseudos:{not:le(function(e){var r=[],i=[],s=f(e.replace($,"$1"));return s[S]?le(function(e,t,n,r){var i,o=s(e,null,r,[]),a=e.length;while(a--)(i=o[a])&&(e[a]=!(t[a]=i))}):function(e,t,n){return r[0]=e,s(r,null,n,i),r[0]=null,!i.pop()}}),has:le(function(t){return function(e){return 0<se(t,e).length}}),contains:le(function(t){return t=t.replace(te,ne),function(e){return-1<(e.textContent||o(e)).indexOf(t)}}),lang:le(function(n){return V.test(n||"")||se.error("unsupported lang: "+n),n=n.replace(te,ne).toLowerCase(),function(e){var t;do{if(t=E?e.lang:e.getAttribute("xml:lang")||e.getAttribute("lang"))return(t=t.toLowerCase())===n||0===t.indexOf(n+"-")}while((e=e.parentNode)&&1===e.nodeType);return!1}}),target:function(e){var t=n.location&&n.location.hash;return t&&t.slice(1)===e.id},root:function(e){return e===a},focus:function(e){return e===C.activeElement&&(!C.hasFocus||C.hasFocus())&&!!(e.type||e.href||~e.tabIndex)},enabled:ge(!1),disabled:ge(!0),checked:function(e){var t=e.nodeName.toLowerCase();return"input"===t&&!!e.checked||"option"===t&&!!e.selected},selected:function(e){return e.parentNode&&e.parentNode.selectedIndex,!0===e.selected},empty:function(e){for(e=e.firstChild;e;e=e.nextSibling)if(e.nodeType<6)return!1;return!0},parent:function(e){return!b.pseudos.empty(e)},header:function(e){return J.test(e.nodeName)},input:function(e){return Q.test(e.nodeName)},button:function(e){var t=e.nodeName.toLowerCase();return"input"===t&&"button"===e.type||"button"===t},text:function(e){var t;return"input"===e.nodeName.toLowerCase()&&"text"===e.type&&(null==(t=e.getAttribute("type"))||"text"===t.toLowerCase())},first:ve(function(){return[0]}),last:ve(function(e,t){return[t-1]}),eq:ve(function(e,t,n){return[n<0?n+t:n]}),even:ve(function(e,t){for(var n=0;n<t;n+=2)e.push(n);return e}),odd:ve(function(e,t){for(var n=1;n<t;n+=2)e.push(n);return e}),lt:ve(function(e,t,n){for(var r=n<0?n+t:t<n?t:n;0<=--r;)e.push(r);return e}),gt:ve(function(e,t,n){for(var r=n<0?n+t:n;++r<t;)e.push(r);return e})}}).pseudos.nth=b.pseudos.eq,{radio:!0,checkbox:!0,file:!0,password:!0,image:!0})b.pseudos[e]=de(e);for(e in{submit:!0,reset:!0})b.pseudos[e]=he(e);function me(){}function xe(e){for(var t=0,n=e.length,r="";t<n;t++)r+=e[t].value;return r}function be(s,e,t){var u=e.dir,l=e.next,c=l||u,f=t&&"parentNode"===c,p=r++;return e.first?function(e,t,n){while(e=e[u])if(1===e.nodeType||f)return s(e,t,n);return!1}:function(e,t,n){var r,i,o,a=[k,p];if(n){while(e=e[u])if((1===e.nodeType||f)&&s(e,t,n))return!0}else while(e=e[u])if(1===e.nodeType||f)if(i=(o=e[S]||(e[S]={}))[e.uniqueID]||(o[e.uniqueID]={}),l&&l===e.nodeName.toLowerCase())e=e[u]||e;else{if((r=i[c])&&r[0]===k&&r[1]===p)return a[2]=r[2];if((i[c]=a)[2]=s(e,t,n))return!0}return!1}}function we(i){return 1<i.length?function(e,t,n){var r=i.length;while(r--)if(!i[r](e,t,n))return!1;return!0}:i[0]}function Te(e,t,n,r,i){for(var o,a=[],s=0,u=e.length,l=null!=t;s<u;s++)(o=e[s])&&(n&&!n(o,r,i)||(a.push(o),l&&t.push(s)));return a}function Ce(d,h,g,v,y,e){return v&&!v[S]&&(v=Ce(v)),y&&!y[S]&&(y=Ce(y,e)),le(function(e,t,n,r){var i,o,a,s=[],u=[],l=t.length,c=e||function(e,t,n){for(var r=0,i=t.length;r<i;r++)se(e,t[r],n);return n}(h||"*",n.nodeType?[n]:n,[]),f=!d||!e&&h?c:Te(c,s,d,n,r),p=g?y||(e?d:l||v)?[]:t:f;if(g&&g(f,p,n,r),v){i=Te(p,u),v(i,[],n,r),o=i.length;while(o--)(a=i[o])&&(p[u[o]]=!(f[u[o]]=a))}if(e){if(y||d){if(y){i=[],o=p.length;while(o--)(a=p[o])&&i.push(f[o]=a);y(null,p=[],i,r)}o=p.length;while(o--)(a=p[o])&&-1<(i=y?P(e,a):s[o])&&(e[i]=!(t[i]=a))}}else p=Te(p===t?p.splice(l,p.length):p),y?y(null,t,p,r):H.apply(t,p)})}function Ee(e){for(var i,t,n,r=e.length,o=b.relative[e[0].type],a=o||b.relative[" "],s=o?1:0,u=be(function(e){return e===i},a,!0),l=be(function(e){return-1<P(i,e)},a,!0),c=[function(e,t,n){var r=!o&&(n||t!==w)||((i=t).nodeType?u(e,t,n):l(e,t,n));return i=null,r}];s<r;s++)if(t=b.relative[e[s].type])c=[be(we(c),t)];else{if((t=b.filter[e[s].type].apply(null,e[s].matches))[S]){for(n=++s;n<r;n++)if(b.relative[e[n].type])break;return Ce(1<s&&we(c),1<s&&xe(e.slice(0,s-1).concat({value:" "===e[s-2].type?"*":""})).replace($,"$1"),t,s<n&&Ee(e.slice(s,n)),n<r&&Ee(e=e.slice(n)),n<r&&xe(e))}c.push(t)}return we(c)}return me.prototype=b.filters=b.pseudos,b.setFilters=new me,h=se.tokenize=function(e,t){var n,r,i,o,a,s,u,l=x[e+" "];if(l)return t?0:l.slice(0);a=e,s=[],u=b.preFilter;while(a){for(o in n&&!(r=_.exec(a))||(r&&(a=a.slice(r[0].length)||a),s.push(i=[])),n=!1,(r=z.exec(a))&&(n=r.shift(),i.push({value:n,type:r[0].replace($," ")}),a=a.slice(n.length)),b.filter)!(r=G[o].exec(a))||u[o]&&!(r=u[o](r))||(n=r.shift(),i.push({value:n,type:o,matches:r}),a=a.slice(n.length));if(!n)break}return t?a.length:a?se.error(e):x(e,s).slice(0)},f=se.compile=function(e,t){var n,v,y,m,x,r,i=[],o=[],a=A[e+" "];if(!a){t||(t=h(e)),n=t.length;while(n--)(a=Ee(t[n]))[S]?i.push(a):o.push(a);(a=A(e,(v=o,m=0<(y=i).length,x=0<v.length,r=function(e,t,n,r,i){var o,a,s,u=0,l="0",c=e&&[],f=[],p=w,d=e||x&&b.find.TAG("*",i),h=k+=null==p?1:Math.random()||.1,g=d.length;for(i&&(w=t==C||t||i);l!==g&&null!=(o=d[l]);l++){if(x&&o){a=0,t||o.ownerDocument==C||(T(o),n=!E);while(s=v[a++])if(s(o,t||C,n)){r.push(o);break}i&&(k=h)}m&&((o=!s&&o)&&u--,e&&c.push(o))}if(u+=l,m&&l!==u){a=0;while(s=y[a++])s(c,f,t,n);if(e){if(0<u)while(l--)c[l]||f[l]||(f[l]=q.call(r));f=Te(f)}H.apply(r,f),i&&!e&&0<f.length&&1<u+y.length&&se.uniqueSort(r)}return i&&(k=h,w=p),c},m?le(r):r))).selector=e}return a},g=se.select=function(e,t,n,r){var i,o,a,s,u,l="function"==typeof e&&e,c=!r&&h(e=l.selector||e);if(n=n||[],1===c.length){if(2<(o=c[0]=c[0].slice(0)).length&&"ID"===(a=o[0]).type&&9===t.nodeType&&E&&b.relative[o[1].type]){if(!(t=(b.find.ID(a.matches[0].replace(te,ne),t)||[])[0]))return n;l&&(t=t.parentNode),e=e.slice(o.shift().value.length)}i=G.needsContext.test(e)?0:o.length;while(i--){if(a=o[i],b.relative[s=a.type])break;if((u=b.find[s])&&(r=u(a.matches[0].replace(te,ne),ee.test(o[0].type)&&ye(t.parentNode)||t))){if(o.splice(i,1),!(e=r.length&&xe(o)))return H.apply(n,r),n;break}}}return(l||f(e,c))(r,t,!E,n,!t||ee.test(e)&&ye(t.parentNode)||t),n},d.sortStable=S.split("").sort(D).join("")===S,d.detectDuplicates=!!l,T(),d.sortDetached=ce(function(e){return 1&e.compareDocumentPosition(C.createElement("fieldset"))}),ce(function(e){return e.innerHTML="<a href='#'></a>","#"===e.firstChild.getAttribute("href")})||fe("type|href|height|width",function(e,t,n){if(!n)return e.getAttribute(t,"type"===t.toLowerCase()?1:2)}),d.attributes&&ce(function(e){return e.innerHTML="<input/>",e.firstChild.setAttribute("value",""),""===e.firstChild.getAttribute("value")})||fe("value",function(e,t,n){if(!n&&"input"===e.nodeName.toLowerCase())return e.defaultValue}),ce(function(e){return null==e.getAttribute("disabled")})||fe(R,function(e,t,n){var r;if(!n)return!0===e[t]?t.toLowerCase():(r=e.getAttributeNode(t))&&r.specified?r.value:null}),se}(C);S.find=d,S.expr=d.selectors,S.expr[":"]=S.expr.pseudos,S.uniqueSort=S.unique=d.uniqueSort,S.text=d.getText,S.isXMLDoc=d.isXML,S.contains=d.contains,S.escapeSelector=d.escape;var h=function(e,t,n){var r=[],i=void 0!==n;while((e=e[t])&&9!==e.nodeType)if(1===e.nodeType){if(i&&S(e).is(n))break;r.push(e)}return r},T=function(e,t){for(var n=[];e;e=e.nextSibling)1===e.nodeType&&e!==t&&n.push(e);return n},k=S.expr.match.needsContext;function A(e,t){return e.nodeName&&e.nodeName.toLowerCase()===t.toLowerCase()}var N=/^<([a-z][^\/\0>:\x20\t\r\n\f]*)[\x20\t\r\n\f]*\/?>(?:<\/\1>|)$/i;function D(e,n,r){return m(n)?S.grep(e,function(e,t){return!!n.call(e,t,e)!==r}):n.nodeType?S.grep(e,function(e){return e===n!==r}):"string"!=typeof n?S.grep(e,function(e){return-1<i.call(n,e)!==r}):S.filter(n,e,r)}S.filter=function(e,t,n){var r=t[0];return n&&(e=":not("+e+")"),1===t.length&&1===r.nodeType?S.find.matchesSelector(r,e)?[r]:[]:S.find.matches(e,S.grep(t,function(e){return 1===e.nodeType}))},S.fn.extend({find:function(e){var t,n,r=this.length,i=this;if("string"!=typeof e)return this.pushStack(S(e).filter(function(){for(t=0;t<r;t++)if(S.contains(i[t],this))return!0}));for(n=this.pushStack([]),t=0;t<r;t++)S.find(e,i[t],n);return 1<r?S.uniqueSort(n):n},filter:function(e){return this.pushStack(D(this,e||[],!1))},not:function(e){return this.pushStack(D(this,e||[],!0))},is:function(e){return!!D(this,"string"==typeof e&&k.test(e)?S(e):e||[],!1).length}});var j,q=/^(?:\s*(<[\w\W]+>)[^>]*|#([\w-]+))$/;(S.fn.init=function(e,t,n){var r,i;if(!e)return this;if(n=n||j,"string"==typeof e){if(!(r="<"===e[0]&&">"===e[e.length-1]&&3<=e.length?[null,e,null]:q.exec(e))||!r[1]&&t)return!t||t.jquery?(t||n).find(e):this.constructor(t).find(e);if(r[1]){if(t=t instanceof S?t[0]:t,S.merge(this,S.parseHTML(r[1],t&&t.nodeType?t.ownerDocument||t:E,!0)),N.test(r[1])&&S.isPlainObject(t))for(r in t)m(this[r])?this[r](t[r]):this.attr(r,t[r]);return this}return(i=E.getElementById(r[2]))&&(this[0]=i,this.length=1),this}return e.nodeType?(this[0]=e,this.length=1,this):m(e)?void 0!==n.ready?n.ready(e):e(S):S.makeArray(e,this)}).prototype=S.fn,j=S(E);var L=/^(?:parents|prev(?:Until|All))/,H={children:!0,contents:!0,next:!0,prev:!0};function O(e,t){while((e=e[t])&&1!==e.nodeType);return e}S.fn.extend({has:function(e){var t=S(e,this),n=t.length;return this.filter(function(){for(var e=0;e<n;e++)if(S.contains(this,t[e]))return!0})},closest:function(e,t){var n,r=0,i=this.length,o=[],a="string"!=typeof e&&S(e);if(!k.test(e))for(;r<i;r++)for(n=this[r];n&&n!==t;n=n.parentNode)if(n.nodeType<11&&(a?-1<a.index(n):1===n.nodeType&&S.find.matchesSelector(n,e))){o.push(n);break}return this.pushStack(1<o.length?S.uniqueSort(o):o)},index:function(e){return e?"string"==typeof e?i.call(S(e),this[0]):i.call(this,e.jquery?e[0]:e):this[0]&&this[0].parentNode?this.first().prevAll().length:-1},add:function(e,t){return this.pushStack(S.uniqueSort(S.merge(this.get(),S(e,t))))},addBack:function(e){return this.add(null==e?this.prevObject:this.prevObject.filter(e))}}),S.each({parent:function(e){var t=e.parentNode;return t&&11!==t.nodeType?t:null},parents:function(e){return h(e,"parentNode")},parentsUntil:function(e,t,n){return h(e,"parentNode",n)},next:function(e){return O(e,"nextSibling")},prev:function(e){return O(e,"previousSibling")},nextAll:function(e){return h(e,"nextSibling")},prevAll:function(e){return h(e,"previousSibling")},nextUntil:function(e,t,n){return h(e,"nextSibling",n)},prevUntil:function(e,t,n){return h(e,"previousSibling",n)},siblings:function(e){return T((e.parentNode||{}).firstChild,e)},children:function(e){return T(e.firstChild)},contents:function(e){return null!=e.contentDocument&&r(e.contentDocument)?e.contentDocument:(A(e,"template")&&(e=e.content||e),S.merge([],e.childNodes))}},function(r,i){S.fn[r]=function(e,t){var n=S.map(this,i,e);return"Until"!==r.slice(-5)&&(t=e),t&&"string"==typeof t&&(n=S.filter(t,n)),1<this.length&&(H[r]||S.uniqueSort(n),L.test(r)&&n.reverse()),this.pushStack(n)}});var P=/[^\x20\t\r\n\f]+/g;function R(e){return e}function M(e){throw e}function I(e,t,n,r){var i;try{e&&m(i=e.promise)?i.call(e).done(t).fail(n):e&&m(i=e.then)?i.call(e,t,n):t.apply(void 0,[e].slice(r))}catch(e){n.apply(void 0,[e])}}S.Callbacks=function(r){var e,n;r="string"==typeof r?(e=r,n={},S.each(e.match(P)||[],function(e,t){n[t]=!0}),n):S.extend({},r);var i,t,o,a,s=[],u=[],l=-1,c=function(){for(a=a||r.once,o=i=!0;u.length;l=-1){t=u.shift();while(++l<s.length)!1===s[l].apply(t[0],t[1])&&r.stopOnFalse&&(l=s.length,t=!1)}r.memory||(t=!1),i=!1,a&&(s=t?[]:"")},f={add:function(){return s&&(t&&!i&&(l=s.length-1,u.push(t)),function n(e){S.each(e,function(e,t){m(t)?r.unique&&f.has(t)||s.push(t):t&&t.length&&"string"!==w(t)&&n(t)})}(arguments),t&&!i&&c()),this},remove:function(){return S.each(arguments,function(e,t){var n;while(-1<(n=S.inArray(t,s,n)))s.splice(n,1),n<=l&&l--}),this},has:function(e){return e?-1<S.inArray(e,s):0<s.length},empty:function(){return s&&(s=[]),this},disable:function(){return a=u=[],s=t="",this},disabled:function(){return!s},lock:function(){return a=u=[],t||i||(s=t=""),this},locked:function(){return!!a},fireWith:function(e,t){return a||(t=[e,(t=t||[]).slice?t.slice():t],u.push(t),i||c()),this},fire:function(){return f.fireWith(this,arguments),this},fired:function(){return!!o}};return f},S.extend({Deferred:function(e){var o=[["notify","progress",S.Callbacks("memory"),S.Callbacks("memory"),2],["resolve","done",S.Callbacks("once memory"),S.Callbacks("once memory"),0,"resolved"],["reject","fail",S.Callbacks("once memory"),S.Callbacks("once memory"),1,"rejected"]],i="pending",a={state:function(){return i},always:function(){return s.done(arguments).fail(arguments),this},"catch":function(e){return a.then(null,e)},pipe:function(){var i=arguments;return S.Deferred(function(r){S.each(o,function(e,t){var n=m(i[t[4]])&&i[t[4]];s[t[1]](function(){var e=n&&n.apply(this,arguments);e&&m(e.promise)?e.promise().progress(r.notify).done(r.resolve).fail(r.reject):r[t[0]+"With"](this,n?[e]:arguments)})}),i=null}).promise()},then:function(t,n,r){var u=0;function l(i,o,a,s){return function(){var n=this,r=arguments,e=function(){var e,t;if(!(i<u)){if((e=a.apply(n,r))===o.promise())throw new TypeError("Thenable self-resolution");t=e&&("object"==typeof e||"function"==typeof e)&&e.then,m(t)?s?t.call(e,l(u,o,R,s),l(u,o,M,s)):(u++,t.call(e,l(u,o,R,s),l(u,o,M,s),l(u,o,R,o.notifyWith))):(a!==R&&(n=void 0,r=[e]),(s||o.resolveWith)(n,r))}},t=s?e:function(){try{e()}catch(e){S.Deferred.exceptionHook&&S.Deferred.exceptionHook(e,t.stackTrace),u<=i+1&&(a!==M&&(n=void 0,r=[e]),o.rejectWith(n,r))}};i?t():(S.Deferred.getStackHook&&(t.stackTrace=S.Deferred.getStackHook()),C.setTimeout(t))}}return S.Deferred(function(e){o[0][3].add(l(0,e,m(r)?r:R,e.notifyWith)),o[1][3].add(l(0,e,m(t)?t:R)),o[2][3].add(l(0,e,m(n)?n:M))}).promise()},promise:function(e){return null!=e?S.extend(e,a):a}},s={};return S.each(o,function(e,t){var n=t[2],r=t[5];a[t[1]]=n.add,r&&n.add(function(){i=r},o[3-e][2].disable,o[3-e][3].disable,o[0][2].lock,o[0][3].lock),n.add(t[3].fire),s[t[0]]=function(){return s[t[0]+"With"](this===s?void 0:this,arguments),this},s[t[0]+"With"]=n.fireWith}),a.promise(s),e&&e.call(s,s),s},when:function(e){var n=arguments.length,t=n,r=Array(t),i=s.call(arguments),o=S.Deferred(),a=function(t){return function(e){r[t]=this,i[t]=1<arguments.length?s.call(arguments):e,--n||o.resolveWith(r,i)}};if(n<=1&&(I(e,o.done(a(t)).resolve,o.reject,!n),"pending"===o.state()||m(i[t]&&i[t].then)))return o.then();while(t--)I(i[t],a(t),o.reject);return o.promise()}});var W=/^(Eval|Internal|Range|Reference|Syntax|Type|URI)Error$/;S.Deferred.exceptionHook=function(e,t){C.console&&C.console.warn&&e&&W.test(e.name)&&C.console.warn("jQuery.Deferred exception: "+e.message,e.stack,t)},S.readyException=function(e){C.setTimeout(function(){throw e})};var F=S.Deferred();function B(){E.removeEventListener("DOMContentLoaded",B),C.removeEventListener("load",B),S.ready()}S.fn.ready=function(e){return F.then(e)["catch"](function(e){S.readyException(e)}),this},S.extend({isReady:!1,readyWait:1,ready:function(e){(!0===e?--S.readyWait:S.isReady)||(S.isReady=!0)!==e&&0<--S.readyWait||F.resolveWith(E,[S])}}),S.ready.then=F.then,"complete"===E.readyState||"loading"!==E.readyState&&!E.documentElement.doScroll?C.setTimeout(S.ready):(E.addEventListener("DOMContentLoaded",B),C.addEventListener("load",B));var $=function(e,t,n,r,i,o,a){var s=0,u=e.length,l=null==n;if("object"===w(n))for(s in i=!0,n)$(e,t,s,n[s],!0,o,a);else if(void 0!==r&&(i=!0,m(r)||(a=!0),l&&(a?(t.call(e,r),t=null):(l=t,t=function(e,t,n){return l.call(S(e),n)})),t))for(;s<u;s++)t(e[s],n,a?r:r.call(e[s],s,t(e[s],n)));return i?e:l?t.call(e):u?t(e[0],n):o},_=/^-ms-/,z=/-([a-z])/g;function U(e,t){return t.toUpperCase()}function X(e){return e.replace(_,"ms-").replace(z,U)}var V=function(e){return 1===e.nodeType||9===e.nodeType||!+e.nodeType};function G(){this.expando=S.expando+G.uid++}G.uid=1,G.prototype={cache:function(e){var t=e[this.expando];return t||(t={},V(e)&&(e.nodeType?e[this.expando]=t:Object.defineProperty(e,this.expando,{value:t,configurable:!0}))),t},set:function(e,t,n){var r,i=this.cache(e);if("string"==typeof t)i[X(t)]=n;else for(r in t)i[X(r)]=t[r];return i},get:function(e,t){return void 0===t?this.cache(e):e[this.expando]&&e[this.expando][X(t)]},access:function(e,t,n){return void 0===t||t&&"string"==typeof t&&void 0===n?this.get(e,t):(this.set(e,t,n),void 0!==n?n:t)},remove:function(e,t){var n,r=e[this.expando];if(void 0!==r){if(void 0!==t){n=(t=Array.isArray(t)?t.map(X):(t=X(t))in r?[t]:t.match(P)||[]).length;while(n--)delete r[t[n]]}(void 0===t||S.isEmptyObject(r))&&(e.nodeType?e[this.expando]=void 0:delete e[this.expando])}},hasData:function(e){var t=e[this.expando];return void 0!==t&&!S.isEmptyObject(t)}};var Y=new G,Q=new G,J=/^(?:\{[\w\W]*\}|\[[\w\W]*\])$/,K=/[A-Z]/g;function Z(e,t,n){var r,i;if(void 0===n&&1===e.nodeType)if(r="data-"+t.replace(K,"-$&").toLowerCase(),"string"==typeof(n=e.getAttribute(r))){try{n="true"===(i=n)||"false"!==i&&("null"===i?null:i===+i+""?+i:J.test(i)?JSON.parse(i):i)}catch(e){}Q.set(e,t,n)}else n=void 0;return n}S.extend({hasData:function(e){return Q.hasData(e)||Y.hasData(e)},data:function(e,t,n){return Q.access(e,t,n)},removeData:function(e,t){Q.remove(e,t)},_data:function(e,t,n){return Y.access(e,t,n)},_removeData:function(e,t){Y.remove(e,t)}}),S.fn.extend({data:function(n,e){var t,r,i,o=this[0],a=o&&o.attributes;if(void 0===n){if(this.length&&(i=Q.get(o),1===o.nodeType&&!Y.get(o,"hasDataAttrs"))){t=a.length;while(t--)a[t]&&0===(r=a[t].name).indexOf("data-")&&(r=X(r.slice(5)),Z(o,r,i[r]));Y.set(o,"hasDataAttrs",!0)}return i}return"object"==typeof n?this.each(function(){Q.set(this,n)}):$(this,function(e){var t;if(o&&void 0===e)return void 0!==(t=Q.get(o,n))?t:void 0!==(t=Z(o,n))?t:void 0;this.each(function(){Q.set(this,n,e)})},null,e,1<arguments.length,null,!0)},removeData:function(e){return this.each(function(){Q.remove(this,e)})}}),S.extend({queue:function(e,t,n){var r;if(e)return t=(t||"fx")+"queue",r=Y.get(e,t),n&&(!r||Array.isArray(n)?r=Y.access(e,t,S.makeArray(n)):r.push(n)),r||[]},dequeue:function(e,t){t=t||"fx";var n=S.queue(e,t),r=n.length,i=n.shift(),o=S._queueHooks(e,t);"inprogress"===i&&(i=n.shift(),r--),i&&("fx"===t&&n.unshift("inprogress"),delete o.stop,i.call(e,function(){S.dequeue(e,t)},o)),!r&&o&&o.empty.fire()},_queueHooks:function(e,t){var n=t+"queueHooks";return Y.get(e,n)||Y.access(e,n,{empty:S.Callbacks("once memory").add(function(){Y.remove(e,[t+"queue",n])})})}}),S.fn.extend({queue:function(t,n){var e=2;return"string"!=typeof t&&(n=t,t="fx",e--),arguments.length<e?S.queue(this[0],t):void 0===n?this:this.each(function(){var e=S.queue(this,t,n);S._queueHooks(this,t),"fx"===t&&"inprogress"!==e[0]&&S.dequeue(this,t)})},dequeue:function(e){return this.each(function(){S.dequeue(this,e)})},clearQueue:function(e){return this.queue(e||"fx",[])},promise:function(e,t){var n,r=1,i=S.Deferred(),o=this,a=this.length,s=function(){--r||i.resolveWith(o,[o])};"string"!=typeof e&&(t=e,e=void 0),e=e||"fx";while(a--)(n=Y.get(o[a],e+"queueHooks"))&&n.empty&&(r++,n.empty.add(s));return s(),i.promise(t)}});var ee=/[+-]?(?:\d*\.|)\d+(?:[eE][+-]?\d+|)/.source,te=new RegExp("^(?:([+-])=|)("+ee+")([a-z%]*)$","i"),ne=["Top","Right","Bottom","Left"],re=E.documentElement,ie=function(e){return S.contains(e.ownerDocument,e)},oe={composed:!0};re.getRootNode&&(ie=function(e){return S.contains(e.ownerDocument,e)||e.getRootNode(oe)===e.ownerDocument});var ae=function(e,t){return"none"===(e=t||e).style.display||""===e.style.display&&ie(e)&&"none"===S.css(e,"display")};function se(e,t,n,r){var i,o,a=20,s=r?function(){return r.cur()}:function(){return S.css(e,t,"")},u=s(),l=n&&n[3]||(S.cssNumber[t]?"":"px"),c=e.nodeType&&(S.cssNumber[t]||"px"!==l&&+u)&&te.exec(S.css(e,t));if(c&&c[3]!==l){u/=2,l=l||c[3],c=+u||1;while(a--)S.style(e,t,c+l),(1-o)*(1-(o=s()/u||.5))<=0&&(a=0),c/=o;c*=2,S.style(e,t,c+l),n=n||[]}return n&&(c=+c||+u||0,i=n[1]?c+(n[1]+1)*n[2]:+n[2],r&&(r.unit=l,r.start=c,r.end=i)),i}var ue={};function le(e,t){for(var n,r,i,o,a,s,u,l=[],c=0,f=e.length;c<f;c++)(r=e[c]).style&&(n=r.style.display,t?("none"===n&&(l[c]=Y.get(r,"display")||null,l[c]||(r.style.display="")),""===r.style.display&&ae(r)&&(l[c]=(u=a=o=void 0,a=(i=r).ownerDocument,s=i.nodeName,(u=ue[s])||(o=a.body.appendChild(a.createElement(s)),u=S.css(o,"display"),o.parentNode.removeChild(o),"none"===u&&(u="block"),ue[s]=u)))):"none"!==n&&(l[c]="none",Y.set(r,"display",n)));for(c=0;c<f;c++)null!=l[c]&&(e[c].style.display=l[c]);return e}S.fn.extend({show:function(){return le(this,!0)},hide:function(){return le(this)},toggle:function(e){return"boolean"==typeof e?e?this.show():this.hide():this.each(function(){ae(this)?S(this).show():S(this).hide()})}});var ce,fe,pe=/^(?:checkbox|radio)$/i,de=/<([a-z][^\/\0>\x20\t\r\n\f]*)/i,he=/^$|^module$|\/(?:java|ecma)script/i;ce=E.createDocumentFragment().appendChild(E.createElement("div")),(fe=E.createElement("input")).setAttribute("type","radio"),fe.setAttribute("checked","checked"),fe.setAttribute("name","t"),ce.appendChild(fe),y.checkClone=ce.cloneNode(!0).cloneNode(!0).lastChild.checked,ce.innerHTML="<textarea>x</textarea>",y.noCloneChecked=!!ce.cloneNode(!0).lastChild.defaultValue,ce.innerHTML="<option></option>",y.option=!!ce.lastChild;var ge={thead:[1,"<table>","</table>"],col:[2,"<table><colgroup>","</colgroup></table>"],tr:[2,"<table><tbody>","</tbody></table>"],td:[3,"<table><tbody><tr>","</tr></tbody></table>"],_default:[0,"",""]};function ve(e,t){var n;return n="undefined"!=typeof e.getElementsByTagName?e.getElementsByTagName(t||"*"):"undefined"!=typeof e.querySelectorAll?e.querySelectorAll(t||"*"):[],void 0===t||t&&A(e,t)?S.merge([e],n):n}function ye(e,t){for(var n=0,r=e.length;n<r;n++)Y.set(e[n],"globalEval",!t||Y.get(t[n],"globalEval"))}ge.tbody=ge.tfoot=ge.colgroup=ge.caption=ge.thead,ge.th=ge.td,y.option||(ge.optgroup=ge.option=[1,"<select multiple='multiple'>","</select>"]);var me=/<|&#?\w+;/;function xe(e,t,n,r,i){for(var o,a,s,u,l,c,f=t.createDocumentFragment(),p=[],d=0,h=e.length;d<h;d++)if((o=e[d])||0===o)if("object"===w(o))S.merge(p,o.nodeType?[o]:o);else if(me.test(o)){a=a||f.appendChild(t.createElement("div")),s=(de.exec(o)||["",""])[1].toLowerCase(),u=ge[s]||ge._default,a.innerHTML=u[1]+S.htmlPrefilter(o)+u[2],c=u[0];while(c--)a=a.lastChild;S.merge(p,a.childNodes),(a=f.firstChild).textContent=""}else p.push(t.createTextNode(o));f.textContent="",d=0;while(o=p[d++])if(r&&-1<S.inArray(o,r))i&&i.push(o);else if(l=ie(o),a=ve(f.appendChild(o),"script"),l&&ye(a),n){c=0;while(o=a[c++])he.test(o.type||"")&&n.push(o)}return f}var be=/^key/,we=/^(?:mouse|pointer|contextmenu|drag|drop)|click/,Te=/^([^.]*)(?:\.(.+)|)/;function Ce(){return!0}function Ee(){return!1}function Se(e,t){return e===function(){try{return E.activeElement}catch(e){}}()==("focus"===t)}function ke(e,t,n,r,i,o){var a,s;if("object"==typeof t){for(s in"string"!=typeof n&&(r=r||n,n=void 0),t)ke(e,s,n,r,t[s],o);return e}if(null==r&&null==i?(i=n,r=n=void 0):null==i&&("string"==typeof n?(i=r,r=void 0):(i=r,r=n,n=void 0)),!1===i)i=Ee;else if(!i)return e;return 1===o&&(a=i,(i=function(e){return S().off(e),a.apply(this,arguments)}).guid=a.guid||(a.guid=S.guid++)),e.each(function(){S.event.add(this,t,i,r,n)})}function Ae(e,i,o){o?(Y.set(e,i,!1),S.event.add(e,i,{namespace:!1,handler:function(e){var t,n,r=Y.get(this,i);if(1&e.isTrigger&&this[i]){if(r.length)(S.event.special[i]||{}).delegateType&&e.stopPropagation();else if(r=s.call(arguments),Y.set(this,i,r),t=o(this,i),this[i](),r!==(n=Y.get(this,i))||t?Y.set(this,i,!1):n={},r!==n)return e.stopImmediatePropagation(),e.preventDefault(),n.value}else r.length&&(Y.set(this,i,{value:S.event.trigger(S.extend(r[0],S.Event.prototype),r.slice(1),this)}),e.stopImmediatePropagation())}})):void 0===Y.get(e,i)&&S.event.add(e,i,Ce)}S.event={global:{},add:function(t,e,n,r,i){var o,a,s,u,l,c,f,p,d,h,g,v=Y.get(t);if(V(t)){n.handler&&(n=(o=n).handler,i=o.selector),i&&S.find.matchesSelector(re,i),n.guid||(n.guid=S.guid++),(u=v.events)||(u=v.events=Object.create(null)),(a=v.handle)||(a=v.handle=function(e){return"undefined"!=typeof S&&S.event.triggered!==e.type?S.event.dispatch.apply(t,arguments):void 0}),l=(e=(e||"").match(P)||[""]).length;while(l--)d=g=(s=Te.exec(e[l])||[])[1],h=(s[2]||"").split(".").sort(),d&&(f=S.event.special[d]||{},d=(i?f.delegateType:f.bindType)||d,f=S.event.special[d]||{},c=S.extend({type:d,origType:g,data:r,handler:n,guid:n.guid,selector:i,needsContext:i&&S.expr.match.needsContext.test(i),namespace:h.join(".")},o),(p=u[d])||((p=u[d]=[]).delegateCount=0,f.setup&&!1!==f.setup.call(t,r,h,a)||t.addEventListener&&t.addEventListener(d,a)),f.add&&(f.add.call(t,c),c.handler.guid||(c.handler.guid=n.guid)),i?p.splice(p.delegateCount++,0,c):p.push(c),S.event.global[d]=!0)}},remove:function(e,t,n,r,i){var o,a,s,u,l,c,f,p,d,h,g,v=Y.hasData(e)&&Y.get(e);if(v&&(u=v.events)){l=(t=(t||"").match(P)||[""]).length;while(l--)if(d=g=(s=Te.exec(t[l])||[])[1],h=(s[2]||"").split(".").sort(),d){f=S.event.special[d]||{},p=u[d=(r?f.delegateType:f.bindType)||d]||[],s=s[2]&&new RegExp("(^|\\.)"+h.join("\\.(?:.*\\.|)")+"(\\.|$)"),a=o=p.length;while(o--)c=p[o],!i&&g!==c.origType||n&&n.guid!==c.guid||s&&!s.test(c.namespace)||r&&r!==c.selector&&("**"!==r||!c.selector)||(p.splice(o,1),c.selector&&p.delegateCount--,f.remove&&f.remove.call(e,c));a&&!p.length&&(f.teardown&&!1!==f.teardown.call(e,h,v.handle)||S.removeEvent(e,d,v.handle),delete u[d])}else for(d in u)S.event.remove(e,d+t[l],n,r,!0);S.isEmptyObject(u)&&Y.remove(e,"handle events")}},dispatch:function(e){var t,n,r,i,o,a,s=new Array(arguments.length),u=S.event.fix(e),l=(Y.get(this,"events")||Object.create(null))[u.type]||[],c=S.event.special[u.type]||{};for(s[0]=u,t=1;t<arguments.length;t++)s[t]=arguments[t];if(u.delegateTarget=this,!c.preDispatch||!1!==c.preDispatch.call(this,u)){a=S.event.handlers.call(this,u,l),t=0;while((i=a[t++])&&!u.isPropagationStopped()){u.currentTarget=i.elem,n=0;while((o=i.handlers[n++])&&!u.isImmediatePropagationStopped())u.rnamespace&&!1!==o.namespace&&!u.rnamespace.test(o.namespace)||(u.handleObj=o,u.data=o.data,void 0!==(r=((S.event.special[o.origType]||{}).handle||o.handler).apply(i.elem,s))&&!1===(u.result=r)&&(u.preventDefault(),u.stopPropagation()))}return c.postDispatch&&c.postDispatch.call(this,u),u.result}},handlers:function(e,t){var n,r,i,o,a,s=[],u=t.delegateCount,l=e.target;if(u&&l.nodeType&&!("click"===e.type&&1<=e.button))for(;l!==this;l=l.parentNode||this)if(1===l.nodeType&&("click"!==e.type||!0!==l.disabled)){for(o=[],a={},n=0;n<u;n++)void 0===a[i=(r=t[n]).selector+" "]&&(a[i]=r.needsContext?-1<S(i,this).index(l):S.find(i,this,null,[l]).length),a[i]&&o.push(r);o.length&&s.push({elem:l,handlers:o})}return l=this,u<t.length&&s.push({elem:l,handlers:t.slice(u)}),s},addProp:function(t,e){Object.defineProperty(S.Event.prototype,t,{enumerable:!0,configurable:!0,get:m(e)?function(){if(this.originalEvent)return e(this.originalEvent)}:function(){if(this.originalEvent)return this.originalEvent[t]},set:function(e){Object.defineProperty(this,t,{enumerable:!0,configurable:!0,writable:!0,value:e})}})},fix:function(e){return e[S.expando]?e:new S.Event(e)},special:{load:{noBubble:!0},click:{setup:function(e){var t=this||e;return pe.test(t.type)&&t.click&&A(t,"input")&&Ae(t,"click",Ce),!1},trigger:function(e){var t=this||e;return pe.test(t.type)&&t.click&&A(t,"input")&&Ae(t,"click"),!0},_default:function(e){var t=e.target;return pe.test(t.type)&&t.click&&A(t,"input")&&Y.get(t,"click")||A(t,"a")}},beforeunload:{postDispatch:function(e){void 0!==e.result&&e.originalEvent&&(e.originalEvent.returnValue=e.result)}}}},S.removeEvent=function(e,t,n){e.removeEventListener&&e.removeEventListener(t,n)},S.Event=function(e,t){if(!(this instanceof S.Event))return new S.Event(e,t);e&&e.type?(this.originalEvent=e,this.type=e.type,this.isDefaultPrevented=e.defaultPrevented||void 0===e.defaultPrevented&&!1===e.returnValue?Ce:Ee,this.target=e.target&&3===e.target.nodeType?e.target.parentNode:e.target,this.currentTarget=e.currentTarget,this.relatedTarget=e.relatedTarget):this.type=e,t&&S.extend(this,t),this.timeStamp=e&&e.timeStamp||Date.now(),this[S.expando]=!0},S.Event.prototype={constructor:S.Event,isDefaultPrevented:Ee,isPropagationStopped:Ee,isImmediatePropagationStopped:Ee,isSimulated:!1,preventDefault:function(){var e=this.originalEvent;this.isDefaultPrevented=Ce,e&&!this.isSimulated&&e.preventDefault()},stopPropagation:function(){var e=this.originalEvent;this.isPropagationStopped=Ce,e&&!this.isSimulated&&e.stopPropagation()},stopImmediatePropagation:function(){var e=this.originalEvent;this.isImmediatePropagationStopped=Ce,e&&!this.isSimulated&&e.stopImmediatePropagation(),this.stopPropagation()}},S.each({altKey:!0,bubbles:!0,cancelable:!0,changedTouches:!0,ctrlKey:!0,detail:!0,eventPhase:!0,metaKey:!0,pageX:!0,pageY:!0,shiftKey:!0,view:!0,"char":!0,code:!0,charCode:!0,key:!0,keyCode:!0,button:!0,buttons:!0,clientX:!0,clientY:!0,offsetX:!0,offsetY:!0,pointerId:!0,pointerType:!0,screenX:!0,screenY:!0,targetTouches:!0,toElement:!0,touches:!0,which:function(e){var t=e.button;return null==e.which&&be.test(e.type)?null!=e.charCode?e.charCode:e.keyCode:!e.which&&void 0!==t&&we.test(e.type)?1&t?1:2&t?3:4&t?2:0:e.which}},S.event.addProp),S.each({focus:"focusin",blur:"focusout"},function(e,t){S.event.special[e]={setup:function(){return Ae(this,e,Se),!1},trigger:function(){return Ae(this,e),!0},delegateType:t}}),S.each({mouseenter:"mouseover",mouseleave:"mouseout",pointerenter:"pointerover",pointerleave:"pointerout"},function(e,i){S.event.special[e]={delegateType:i,bindType:i,handle:function(e){var t,n=e.relatedTarget,r=e.handleObj;return n&&(n===this||S.contains(this,n))||(e.type=r.origType,t=r.handler.apply(this,arguments),e.type=i),t}}}),S.fn.extend({on:function(e,t,n,r){return ke(this,e,t,n,r)},one:function(e,t,n,r){return ke(this,e,t,n,r,1)},off:function(e,t,n){var r,i;if(e&&e.preventDefault&&e.handleObj)return r=e.handleObj,S(e.delegateTarget).off(r.namespace?r.origType+"."+r.namespace:r.origType,r.selector,r.handler),this;if("object"==typeof e){for(i in e)this.off(i,t,e[i]);return this}return!1!==t&&"function"!=typeof t||(n=t,t=void 0),!1===n&&(n=Ee),this.each(function(){S.event.remove(this,e,n,t)})}});var Ne=/<script|<style|<link/i,De=/checked\s*(?:[^=]|=\s*.checked.)/i,je=/^\s*<!(?:\[CDATA\[|--)|(?:\]\]|--)>\s*$/g;function qe(e,t){return A(e,"table")&&A(11!==t.nodeType?t:t.firstChild,"tr")&&S(e).children("tbody")[0]||e}function Le(e){return e.type=(null!==e.getAttribute("type"))+"/"+e.type,e}function He(e){return"true/"===(e.type||"").slice(0,5)?e.type=e.type.slice(5):e.removeAttribute("type"),e}function Oe(e,t){var n,r,i,o,a,s;if(1===t.nodeType){if(Y.hasData(e)&&(s=Y.get(e).events))for(i in Y.remove(t,"handle events"),s)for(n=0,r=s[i].length;n<r;n++)S.event.add(t,i,s[i][n]);Q.hasData(e)&&(o=Q.access(e),a=S.extend({},o),Q.set(t,a))}}function Pe(n,r,i,o){r=g(r);var e,t,a,s,u,l,c=0,f=n.length,p=f-1,d=r[0],h=m(d);if(h||1<f&&"string"==typeof d&&!y.checkClone&&De.test(d))return n.each(function(e){var t=n.eq(e);h&&(r[0]=d.call(this,e,t.html())),Pe(t,r,i,o)});if(f&&(t=(e=xe(r,n[0].ownerDocument,!1,n,o)).firstChild,1===e.childNodes.length&&(e=t),t||o)){for(s=(a=S.map(ve(e,"script"),Le)).length;c<f;c++)u=e,c!==p&&(u=S.clone(u,!0,!0),s&&S.merge(a,ve(u,"script"))),i.call(n[c],u,c);if(s)for(l=a[a.length-1].ownerDocument,S.map(a,He),c=0;c<s;c++)u=a[c],he.test(u.type||"")&&!Y.access(u,"globalEval")&&S.contains(l,u)&&(u.src&&"module"!==(u.type||"").toLowerCase()?S._evalUrl&&!u.noModule&&S._evalUrl(u.src,{nonce:u.nonce||u.getAttribute("nonce")},l):b(u.textContent.replace(je,""),u,l))}return n}function Re(e,t,n){for(var r,i=t?S.filter(t,e):e,o=0;null!=(r=i[o]);o++)n||1!==r.nodeType||S.cleanData(ve(r)),r.parentNode&&(n&&ie(r)&&ye(ve(r,"script")),r.parentNode.removeChild(r));return e}S.extend({htmlPrefilter:function(e){return e},clone:function(e,t,n){var r,i,o,a,s,u,l,c=e.cloneNode(!0),f=ie(e);if(!(y.noCloneChecked||1!==e.nodeType&&11!==e.nodeType||S.isXMLDoc(e)))for(a=ve(c),r=0,i=(o=ve(e)).length;r<i;r++)s=o[r],u=a[r],void 0,"input"===(l=u.nodeName.toLowerCase())&&pe.test(s.type)?u.checked=s.checked:"input"!==l&&"textarea"!==l||(u.defaultValue=s.defaultValue);if(t)if(n)for(o=o||ve(e),a=a||ve(c),r=0,i=o.length;r<i;r++)Oe(o[r],a[r]);else Oe(e,c);return 0<(a=ve(c,"script")).length&&ye(a,!f&&ve(e,"script")),c},cleanData:function(e){for(var t,n,r,i=S.event.special,o=0;void 0!==(n=e[o]);o++)if(V(n)){if(t=n[Y.expando]){if(t.events)for(r in t.events)i[r]?S.event.remove(n,r):S.removeEvent(n,r,t.handle);n[Y.expando]=void 0}n[Q.expando]&&(n[Q.expando]=void 0)}}}),S.fn.extend({detach:function(e){return Re(this,e,!0)},remove:function(e){return Re(this,e)},text:function(e){return $(this,function(e){return void 0===e?S.text(this):this.empty().each(function(){1!==this.nodeType&&11!==this.nodeType&&9!==this.nodeType||(this.textContent=e)})},null,e,arguments.length)},append:function(){return Pe(this,arguments,function(e){1!==this.nodeType&&11!==this.nodeType&&9!==this.nodeType||qe(this,e).appendChild(e)})},prepend:function(){return Pe(this,arguments,function(e){if(1===this.nodeType||11===this.nodeType||9===this.nodeType){var t=qe(this,e);t.insertBefore(e,t.firstChild)}})},before:function(){return Pe(this,arguments,function(e){this.parentNode&&this.parentNode.insertBefore(e,this)})},after:function(){return Pe(this,arguments,function(e){this.parentNode&&this.parentNode.insertBefore(e,this.nextSibling)})},empty:function(){for(var e,t=0;null!=(e=this[t]);t++)1===e.nodeType&&(S.cleanData(ve(e,!1)),e.textContent="");return this},clone:function(e,t){return e=null!=e&&e,t=null==t?e:t,this.map(function(){return S.clone(this,e,t)})},html:function(e){return $(this,function(e){var t=this[0]||{},n=0,r=this.length;if(void 0===e&&1===t.nodeType)return t.innerHTML;if("string"==typeof e&&!Ne.test(e)&&!ge[(de.exec(e)||["",""])[1].toLowerCase()]){e=S.htmlPrefilter(e);try{for(;n<r;n++)1===(t=this[n]||{}).nodeType&&(S.cleanData(ve(t,!1)),t.innerHTML=e);t=0}catch(e){}}t&&this.empty().append(e)},null,e,arguments.length)},replaceWith:function(){var n=[];return Pe(this,arguments,function(e){var t=this.parentNode;S.inArray(this,n)<0&&(S.cleanData(ve(this)),t&&t.replaceChild(e,this))},n)}}),S.each({appendTo:"append",prependTo:"prepend",insertBefore:"before",insertAfter:"after",replaceAll:"replaceWith"},function(e,a){S.fn[e]=function(e){for(var t,n=[],r=S(e),i=r.length-1,o=0;o<=i;o++)t=o===i?this:this.clone(!0),S(r[o])[a](t),u.apply(n,t.get());return this.pushStack(n)}});var Me=new RegExp("^("+ee+")(?!px)[a-z%]+$","i"),Ie=function(e){var t=e.ownerDocument.defaultView;return t&&t.opener||(t=C),t.getComputedStyle(e)},We=function(e,t,n){var r,i,o={};for(i in t)o[i]=e.style[i],e.style[i]=t[i];for(i in r=n.call(e),t)e.style[i]=o[i];return r},Fe=new RegExp(ne.join("|"),"i");function Be(e,t,n){var r,i,o,a,s=e.style;return(n=n||Ie(e))&&(""!==(a=n.getPropertyValue(t)||n[t])||ie(e)||(a=S.style(e,t)),!y.pixelBoxStyles()&&Me.test(a)&&Fe.test(t)&&(r=s.width,i=s.minWidth,o=s.maxWidth,s.minWidth=s.maxWidth=s.width=a,a=n.width,s.width=r,s.minWidth=i,s.maxWidth=o)),void 0!==a?a+"":a}function $e(e,t){return{get:function(){if(!e())return(this.get=t).apply(this,arguments);delete this.get}}}!function(){function e(){if(l){u.style.cssText="position:absolute;left:-11111px;width:60px;margin-top:1px;padding:0;border:0",l.style.cssText="position:relative;display:block;box-sizing:border-box;overflow:scroll;margin:auto;border:1px;padding:1px;width:60%;top:1%",re.appendChild(u).appendChild(l);var e=C.getComputedStyle(l);n="1%"!==e.top,s=12===t(e.marginLeft),l.style.right="60%",o=36===t(e.right),r=36===t(e.width),l.style.position="absolute",i=12===t(l.offsetWidth/3),re.removeChild(u),l=null}}function t(e){return Math.round(parseFloat(e))}var n,r,i,o,a,s,u=E.createElement("div"),l=E.createElement("div");l.style&&(l.style.backgroundClip="content-box",l.cloneNode(!0).style.backgroundClip="",y.clearCloneStyle="content-box"===l.style.backgroundClip,S.extend(y,{boxSizingReliable:function(){return e(),r},pixelBoxStyles:function(){return e(),o},pixelPosition:function(){return e(),n},reliableMarginLeft:function(){return e(),s},scrollboxSize:function(){return e(),i},reliableTrDimensions:function(){var e,t,n,r;return null==a&&(e=E.createElement("table"),t=E.createElement("tr"),n=E.createElement("div"),e.style.cssText="position:absolute;left:-11111px",t.style.height="1px",n.style.height="9px",re.appendChild(e).appendChild(t).appendChild(n),r=C.getComputedStyle(t),a=3<parseInt(r.height),re.removeChild(e)),a}}))}();var _e=["Webkit","Moz","ms"],ze=E.createElement("div").style,Ue={};function Xe(e){var t=S.cssProps[e]||Ue[e];return t||(e in ze?e:Ue[e]=function(e){var t=e[0].toUpperCase()+e.slice(1),n=_e.length;while(n--)if((e=_e[n]+t)in ze)return e}(e)||e)}var Ve=/^(none|table(?!-c[ea]).+)/,Ge=/^--/,Ye={position:"absolute",visibility:"hidden",display:"block"},Qe={letterSpacing:"0",fontWeight:"400"};function Je(e,t,n){var r=te.exec(t);return r?Math.max(0,r[2]-(n||0))+(r[3]||"px"):t}function Ke(e,t,n,r,i,o){var a="width"===t?1:0,s=0,u=0;if(n===(r?"border":"content"))return 0;for(;a<4;a+=2)"margin"===n&&(u+=S.css(e,n+ne[a],!0,i)),r?("content"===n&&(u-=S.css(e,"padding"+ne[a],!0,i)),"margin"!==n&&(u-=S.css(e,"border"+ne[a]+"Width",!0,i))):(u+=S.css(e,"padding"+ne[a],!0,i),"padding"!==n?u+=S.css(e,"border"+ne[a]+"Width",!0,i):s+=S.css(e,"border"+ne[a]+"Width",!0,i));return!r&&0<=o&&(u+=Math.max(0,Math.ceil(e["offset"+t[0].toUpperCase()+t.slice(1)]-o-u-s-.5))||0),u}function Ze(e,t,n){var r=Ie(e),i=(!y.boxSizingReliable()||n)&&"border-box"===S.css(e,"boxSizing",!1,r),o=i,a=Be(e,t,r),s="offset"+t[0].toUpperCase()+t.slice(1);if(Me.test(a)){if(!n)return a;a="auto"}return(!y.boxSizingReliable()&&i||!y.reliableTrDimensions()&&A(e,"tr")||"auto"===a||!parseFloat(a)&&"inline"===S.css(e,"display",!1,r))&&e.getClientRects().length&&(i="border-box"===S.css(e,"boxSizing",!1,r),(o=s in e)&&(a=e[s])),(a=parseFloat(a)||0)+Ke(e,t,n||(i?"border":"content"),o,r,a)+"px"}function et(e,t,n,r,i){return new et.prototype.init(e,t,n,r,i)}S.extend({cssHooks:{opacity:{get:function(e,t){if(t){var n=Be(e,"opacity");return""===n?"1":n}}}},cssNumber:{animationIterationCount:!0,columnCount:!0,fillOpacity:!0,flexGrow:!0,flexShrink:!0,fontWeight:!0,gridArea:!0,gridColumn:!0,gridColumnEnd:!0,gridColumnStart:!0,gridRow:!0,gridRowEnd:!0,gridRowStart:!0,lineHeight:!0,opacity:!0,order:!0,orphans:!0,widows:!0,zIndex:!0,zoom:!0},cssProps:{},style:function(e,t,n,r){if(e&&3!==e.nodeType&&8!==e.nodeType&&e.style){var i,o,a,s=X(t),u=Ge.test(t),l=e.style;if(u||(t=Xe(s)),a=S.cssHooks[t]||S.cssHooks[s],void 0===n)return a&&"get"in a&&void 0!==(i=a.get(e,!1,r))?i:l[t];"string"===(o=typeof n)&&(i=te.exec(n))&&i[1]&&(n=se(e,t,i),o="number"),null!=n&&n==n&&("number"!==o||u||(n+=i&&i[3]||(S.cssNumber[s]?"":"px")),y.clearCloneStyle||""!==n||0!==t.indexOf("background")||(l[t]="inherit"),a&&"set"in a&&void 0===(n=a.set(e,n,r))||(u?l.setProperty(t,n):l[t]=n))}},css:function(e,t,n,r){var i,o,a,s=X(t);return Ge.test(t)||(t=Xe(s)),(a=S.cssHooks[t]||S.cssHooks[s])&&"get"in a&&(i=a.get(e,!0,n)),void 0===i&&(i=Be(e,t,r)),"normal"===i&&t in Qe&&(i=Qe[t]),""===n||n?(o=parseFloat(i),!0===n||isFinite(o)?o||0:i):i}}),S.each(["height","width"],function(e,u){S.cssHooks[u]={get:function(e,t,n){if(t)return!Ve.test(S.css(e,"display"))||e.getClientRects().length&&e.getBoundingClientRect().width?Ze(e,u,n):We(e,Ye,function(){return Ze(e,u,n)})},set:function(e,t,n){var r,i=Ie(e),o=!y.scrollboxSize()&&"absolute"===i.position,a=(o||n)&&"border-box"===S.css(e,"boxSizing",!1,i),s=n?Ke(e,u,n,a,i):0;return a&&o&&(s-=Math.ceil(e["offset"+u[0].toUpperCase()+u.slice(1)]-parseFloat(i[u])-Ke(e,u,"border",!1,i)-.5)),s&&(r=te.exec(t))&&"px"!==(r[3]||"px")&&(e.style[u]=t,t=S.css(e,u)),Je(0,t,s)}}}),S.cssHooks.marginLeft=$e(y.reliableMarginLeft,function(e,t){if(t)return(parseFloat(Be(e,"marginLeft"))||e.getBoundingClientRect().left-We(e,{marginLeft:0},function(){return e.getBoundingClientRect().left}))+"px"}),S.each({margin:"",padding:"",border:"Width"},function(i,o){S.cssHooks[i+o]={expand:function(e){for(var t=0,n={},r="string"==typeof e?e.split(" "):[e];t<4;t++)n[i+ne[t]+o]=r[t]||r[t-2]||r[0];return n}},"margin"!==i&&(S.cssHooks[i+o].set=Je)}),S.fn.extend({css:function(e,t){return $(this,function(e,t,n){var r,i,o={},a=0;if(Array.isArray(t)){for(r=Ie(e),i=t.length;a<i;a++)o[t[a]]=S.css(e,t[a],!1,r);return o}return void 0!==n?S.style(e,t,n):S.css(e,t)},e,t,1<arguments.length)}}),((S.Tween=et).prototype={constructor:et,init:function(e,t,n,r,i,o){this.elem=e,this.prop=n,this.easing=i||S.easing._default,this.options=t,this.start=this.now=this.cur(),this.end=r,this.unit=o||(S.cssNumber[n]?"":"px")},cur:function(){var e=et.propHooks[this.prop];return e&&e.get?e.get(this):et.propHooks._default.get(this)},run:function(e){var t,n=et.propHooks[this.prop];return this.options.duration?this.pos=t=S.easing[this.easing](e,this.options.duration*e,0,1,this.options.duration):this.pos=t=e,this.now=(this.end-this.start)*t+this.start,this.options.step&&this.options.step.call(this.elem,this.now,this),n&&n.set?n.set(this):et.propHooks._default.set(this),this}}).init.prototype=et.prototype,(et.propHooks={_default:{get:function(e){var t;return 1!==e.elem.nodeType||null!=e.elem[e.prop]&&null==e.elem.style[e.prop]?e.elem[e.prop]:(t=S.css(e.elem,e.prop,""))&&"auto"!==t?t:0},set:function(e){S.fx.step[e.prop]?S.fx.step[e.prop](e):1!==e.elem.nodeType||!S.cssHooks[e.prop]&&null==e.elem.style[Xe(e.prop)]?e.elem[e.prop]=e.now:S.style(e.elem,e.prop,e.now+e.unit)}}}).scrollTop=et.propHooks.scrollLeft={set:function(e){e.elem.nodeType&&e.elem.parentNode&&(e.elem[e.prop]=e.now)}},S.easing={linear:function(e){return e},swing:function(e){return.5-Math.cos(e*Math.PI)/2},_default:"swing"},S.fx=et.prototype.init,S.fx.step={};var tt,nt,rt,it,ot=/^(?:toggle|show|hide)$/,at=/queueHooks$/;function st(){nt&&(!1===E.hidden&&C.requestAnimationFrame?C.requestAnimationFrame(st):C.setTimeout(st,S.fx.interval),S.fx.tick())}function ut(){return C.setTimeout(function(){tt=void 0}),tt=Date.now()}function lt(e,t){var n,r=0,i={height:e};for(t=t?1:0;r<4;r+=2-t)i["margin"+(n=ne[r])]=i["padding"+n]=e;return t&&(i.opacity=i.width=e),i}function ct(e,t,n){for(var r,i=(ft.tweeners[t]||[]).concat(ft.tweeners["*"]),o=0,a=i.length;o<a;o++)if(r=i[o].call(n,t,e))return r}function ft(o,e,t){var n,a,r=0,i=ft.prefilters.length,s=S.Deferred().always(function(){delete u.elem}),u=function(){if(a)return!1;for(var e=tt||ut(),t=Math.max(0,l.startTime+l.duration-e),n=1-(t/l.duration||0),r=0,i=l.tweens.length;r<i;r++)l.tweens[r].run(n);return s.notifyWith(o,[l,n,t]),n<1&&i?t:(i||s.notifyWith(o,[l,1,0]),s.resolveWith(o,[l]),!1)},l=s.promise({elem:o,props:S.extend({},e),opts:S.extend(!0,{specialEasing:{},easing:S.easing._default},t),originalProperties:e,originalOptions:t,startTime:tt||ut(),duration:t.duration,tweens:[],createTween:function(e,t){var n=S.Tween(o,l.opts,e,t,l.opts.specialEasing[e]||l.opts.easing);return l.tweens.push(n),n},stop:function(e){var t=0,n=e?l.tweens.length:0;if(a)return this;for(a=!0;t<n;t++)l.tweens[t].run(1);return e?(s.notifyWith(o,[l,1,0]),s.resolveWith(o,[l,e])):s.rejectWith(o,[l,e]),this}}),c=l.props;for(!function(e,t){var n,r,i,o,a;for(n in e)if(i=t[r=X(n)],o=e[n],Array.isArray(o)&&(i=o[1],o=e[n]=o[0]),n!==r&&(e[r]=o,delete e[n]),(a=S.cssHooks[r])&&"expand"in a)for(n in o=a.expand(o),delete e[r],o)n in e||(e[n]=o[n],t[n]=i);else t[r]=i}(c,l.opts.specialEasing);r<i;r++)if(n=ft.prefilters[r].call(l,o,c,l.opts))return m(n.stop)&&(S._queueHooks(l.elem,l.opts.queue).stop=n.stop.bind(n)),n;return S.map(c,ct,l),m(l.opts.start)&&l.opts.start.call(o,l),l.progress(l.opts.progress).done(l.opts.done,l.opts.complete).fail(l.opts.fail).always(l.opts.always),S.fx.timer(S.extend(u,{elem:o,anim:l,queue:l.opts.queue})),l}S.Animation=S.extend(ft,{tweeners:{"*":[function(e,t){var n=this.createTween(e,t);return se(n.elem,e,te.exec(t),n),n}]},tweener:function(e,t){m(e)?(t=e,e=["*"]):e=e.match(P);for(var n,r=0,i=e.length;r<i;r++)n=e[r],ft.tweeners[n]=ft.tweeners[n]||[],ft.tweeners[n].unshift(t)},prefilters:[function(e,t,n){var r,i,o,a,s,u,l,c,f="width"in t||"height"in t,p=this,d={},h=e.style,g=e.nodeType&&ae(e),v=Y.get(e,"fxshow");for(r in n.queue||(null==(a=S._queueHooks(e,"fx")).unqueued&&(a.unqueued=0,s=a.empty.fire,a.empty.fire=function(){a.unqueued||s()}),a.unqueued++,p.always(function(){p.always(function(){a.unqueued--,S.queue(e,"fx").length||a.empty.fire()})})),t)if(i=t[r],ot.test(i)){if(delete t[r],o=o||"toggle"===i,i===(g?"hide":"show")){if("show"!==i||!v||void 0===v[r])continue;g=!0}d[r]=v&&v[r]||S.style(e,r)}if((u=!S.isEmptyObject(t))||!S.isEmptyObject(d))for(r in f&&1===e.nodeType&&(n.overflow=[h.overflow,h.overflowX,h.overflowY],null==(l=v&&v.display)&&(l=Y.get(e,"display")),"none"===(c=S.css(e,"display"))&&(l?c=l:(le([e],!0),l=e.style.display||l,c=S.css(e,"display"),le([e]))),("inline"===c||"inline-block"===c&&null!=l)&&"none"===S.css(e,"float")&&(u||(p.done(function(){h.display=l}),null==l&&(c=h.display,l="none"===c?"":c)),h.display="inline-block")),n.overflow&&(h.overflow="hidden",p.always(function(){h.overflow=n.overflow[0],h.overflowX=n.overflow[1],h.overflowY=n.overflow[2]})),u=!1,d)u||(v?"hidden"in v&&(g=v.hidden):v=Y.access(e,"fxshow",{display:l}),o&&(v.hidden=!g),g&&le([e],!0),p.done(function(){for(r in g||le([e]),Y.remove(e,"fxshow"),d)S.style(e,r,d[r])})),u=ct(g?v[r]:0,r,p),r in v||(v[r]=u.start,g&&(u.end=u.start,u.start=0))}],prefilter:function(e,t){t?ft.prefilters.unshift(e):ft.prefilters.push(e)}}),S.speed=function(e,t,n){var r=e&&"object"==typeof e?S.extend({},e):{complete:n||!n&&t||m(e)&&e,duration:e,easing:n&&t||t&&!m(t)&&t};return S.fx.off?r.duration=0:"number"!=typeof r.duration&&(r.duration in S.fx.speeds?r.duration=S.fx.speeds[r.duration]:r.duration=S.fx.speeds._default),null!=r.queue&&!0!==r.queue||(r.queue="fx"),r.old=r.complete,r.complete=function(){m(r.old)&&r.old.call(this),r.queue&&S.dequeue(this,r.queue)},r},S.fn.extend({fadeTo:function(e,t,n,r){return this.filter(ae).css("opacity",0).show().end().animate({opacity:t},e,n,r)},animate:function(t,e,n,r){var i=S.isEmptyObject(t),o=S.speed(e,n,r),a=function(){var e=ft(this,S.extend({},t),o);(i||Y.get(this,"finish"))&&e.stop(!0)};return a.finish=a,i||!1===o.queue?this.each(a):this.queue(o.queue,a)},stop:function(i,e,o){var a=function(e){var t=e.stop;delete e.stop,t(o)};return"string"!=typeof i&&(o=e,e=i,i=void 0),e&&this.queue(i||"fx",[]),this.each(function(){var e=!0,t=null!=i&&i+"queueHooks",n=S.timers,r=Y.get(this);if(t)r[t]&&r[t].stop&&a(r[t]);else for(t in r)r[t]&&r[t].stop&&at.test(t)&&a(r[t]);for(t=n.length;t--;)n[t].elem!==this||null!=i&&n[t].queue!==i||(n[t].anim.stop(o),e=!1,n.splice(t,1));!e&&o||S.dequeue(this,i)})},finish:function(a){return!1!==a&&(a=a||"fx"),this.each(function(){var e,t=Y.get(this),n=t[a+"queue"],r=t[a+"queueHooks"],i=S.timers,o=n?n.length:0;for(t.finish=!0,S.queue(this,a,[]),r&&r.stop&&r.stop.call(this,!0),e=i.length;e--;)i[e].elem===this&&i[e].queue===a&&(i[e].anim.stop(!0),i.splice(e,1));for(e=0;e<o;e++)n[e]&&n[e].finish&&n[e].finish.call(this);delete t.finish})}}),S.each(["toggle","show","hide"],function(e,r){var i=S.fn[r];S.fn[r]=function(e,t,n){return null==e||"boolean"==typeof e?i.apply(this,arguments):this.animate(lt(r,!0),e,t,n)}}),S.each({slideDown:lt("show"),slideUp:lt("hide"),slideToggle:lt("toggle"),fadeIn:{opacity:"show"},fadeOut:{opacity:"hide"},fadeToggle:{opacity:"toggle"}},function(e,r){S.fn[e]=function(e,t,n){return this.animate(r,e,t,n)}}),S.timers=[],S.fx.tick=function(){var e,t=0,n=S.timers;for(tt=Date.now();t<n.length;t++)(e=n[t])()||n[t]!==e||n.splice(t--,1);n.length||S.fx.stop(),tt=void 0},S.fx.timer=function(e){S.timers.push(e),S.fx.start()},S.fx.interval=13,S.fx.start=function(){nt||(nt=!0,st())},S.fx.stop=function(){nt=null},S.fx.speeds={slow:600,fast:200,_default:400},S.fn.delay=function(r,e){return r=S.fx&&S.fx.speeds[r]||r,e=e||"fx",this.queue(e,function(e,t){var n=C.setTimeout(e,r);t.stop=function(){C.clearTimeout(n)}})},rt=E.createElement("input"),it=E.createElement("select").appendChild(E.createElement("option")),rt.type="checkbox",y.checkOn=""!==rt.value,y.optSelected=it.selected,(rt=E.createElement("input")).value="t",rt.type="radio",y.radioValue="t"===rt.value;var pt,dt=S.expr.attrHandle;S.fn.extend({attr:function(e,t){return $(this,S.attr,e,t,1<arguments.length)},removeAttr:function(e){return this.each(function(){S.removeAttr(this,e)})}}),S.extend({attr:function(e,t,n){var r,i,o=e.nodeType;if(3!==o&&8!==o&&2!==o)return"undefined"==typeof e.getAttribute?S.prop(e,t,n):(1===o&&S.isXMLDoc(e)||(i=S.attrHooks[t.toLowerCase()]||(S.expr.match.bool.test(t)?pt:void 0)),void 0!==n?null===n?void S.removeAttr(e,t):i&&"set"in i&&void 0!==(r=i.set(e,n,t))?r:(e.setAttribute(t,n+""),n):i&&"get"in i&&null!==(r=i.get(e,t))?r:null==(r=S.find.attr(e,t))?void 0:r)},attrHooks:{type:{set:function(e,t){if(!y.radioValue&&"radio"===t&&A(e,"input")){var n=e.value;return e.setAttribute("type",t),n&&(e.value=n),t}}}},removeAttr:function(e,t){var n,r=0,i=t&&t.match(P);if(i&&1===e.nodeType)while(n=i[r++])e.removeAttribute(n)}}),pt={set:function(e,t,n){return!1===t?S.removeAttr(e,n):e.setAttribute(n,n),n}},S.each(S.expr.match.bool.source.match(/\w+/g),function(e,t){var a=dt[t]||S.find.attr;dt[t]=function(e,t,n){var r,i,o=t.toLowerCase();return n||(i=dt[o],dt[o]=r,r=null!=a(e,t,n)?o:null,dt[o]=i),r}});var ht=/^(?:input|select|textarea|button)$/i,gt=/^(?:a|area)$/i;function vt(e){return(e.match(P)||[]).join(" ")}function yt(e){return e.getAttribute&&e.getAttribute("class")||""}function mt(e){return Array.isArray(e)?e:"string"==typeof e&&e.match(P)||[]}S.fn.extend({prop:function(e,t){return $(this,S.prop,e,t,1<arguments.length)},removeProp:function(e){return this.each(function(){delete this[S.propFix[e]||e]})}}),S.extend({prop:function(e,t,n){var r,i,o=e.nodeType;if(3!==o&&8!==o&&2!==o)return 1===o&&S.isXMLDoc(e)||(t=S.propFix[t]||t,i=S.propHooks[t]),void 0!==n?i&&"set"in i&&void 0!==(r=i.set(e,n,t))?r:e[t]=n:i&&"get"in i&&null!==(r=i.get(e,t))?r:e[t]},propHooks:{tabIndex:{get:function(e){var t=S.find.attr(e,"tabindex");return t?parseInt(t,10):ht.test(e.nodeName)||gt.test(e.nodeName)&&e.href?0:-1}}},propFix:{"for":"htmlFor","class":"className"}}),y.optSelected||(S.propHooks.selected={get:function(e){var t=e.parentNode;return t&&t.parentNode&&t.parentNode.selectedIndex,null},set:function(e){var t=e.parentNode;t&&(t.selectedIndex,t.parentNode&&t.parentNode.selectedIndex)}}),S.each(["tabIndex","readOnly","maxLength","cellSpacing","cellPadding","rowSpan","colSpan","useMap","frameBorder","contentEditable"],function(){S.propFix[this.toLowerCase()]=this}),S.fn.extend({addClass:function(t){var e,n,r,i,o,a,s,u=0;if(m(t))return this.each(function(e){S(this).addClass(t.call(this,e,yt(this)))});if((e=mt(t)).length)while(n=this[u++])if(i=yt(n),r=1===n.nodeType&&" "+vt(i)+" "){a=0;while(o=e[a++])r.indexOf(" "+o+" ")<0&&(r+=o+" ");i!==(s=vt(r))&&n.setAttribute("class",s)}return this},removeClass:function(t){var e,n,r,i,o,a,s,u=0;if(m(t))return this.each(function(e){S(this).removeClass(t.call(this,e,yt(this)))});if(!arguments.length)return this.attr("class","");if((e=mt(t)).length)while(n=this[u++])if(i=yt(n),r=1===n.nodeType&&" "+vt(i)+" "){a=0;while(o=e[a++])while(-1<r.indexOf(" "+o+" "))r=r.replace(" "+o+" "," ");i!==(s=vt(r))&&n.setAttribute("class",s)}return this},toggleClass:function(i,t){var o=typeof i,a="string"===o||Array.isArray(i);return"boolean"==typeof t&&a?t?this.addClass(i):this.removeClass(i):m(i)?this.each(function(e){S(this).toggleClass(i.call(this,e,yt(this),t),t)}):this.each(function(){var e,t,n,r;if(a){t=0,n=S(this),r=mt(i);while(e=r[t++])n.hasClass(e)?n.removeClass(e):n.addClass(e)}else void 0!==i&&"boolean"!==o||((e=yt(this))&&Y.set(this,"__className__",e),this.setAttribute&&this.setAttribute("class",e||!1===i?"":Y.get(this,"__className__")||""))})},hasClass:function(e){var t,n,r=0;t=" "+e+" ";while(n=this[r++])if(1===n.nodeType&&-1<(" "+vt(yt(n))+" ").indexOf(t))return!0;return!1}});var xt=/\r/g;S.fn.extend({val:function(n){var r,e,i,t=this[0];return arguments.length?(i=m(n),this.each(function(e){var t;1===this.nodeType&&(null==(t=i?n.call(this,e,S(this).val()):n)?t="":"number"==typeof t?t+="":Array.isArray(t)&&(t=S.map(t,function(e){return null==e?"":e+""})),(r=S.valHooks[this.type]||S.valHooks[this.nodeName.toLowerCase()])&&"set"in r&&void 0!==r.set(this,t,"value")||(this.value=t))})):t?(r=S.valHooks[t.type]||S.valHooks[t.nodeName.toLowerCase()])&&"get"in r&&void 0!==(e=r.get(t,"value"))?e:"string"==typeof(e=t.value)?e.replace(xt,""):null==e?"":e:void 0}}),S.extend({valHooks:{option:{get:function(e){var t=S.find.attr(e,"value");return null!=t?t:vt(S.text(e))}},select:{get:function(e){var t,n,r,i=e.options,o=e.selectedIndex,a="select-one"===e.type,s=a?null:[],u=a?o+1:i.length;for(r=o<0?u:a?o:0;r<u;r++)if(((n=i[r]).selected||r===o)&&!n.disabled&&(!n.parentNode.disabled||!A(n.parentNode,"optgroup"))){if(t=S(n).val(),a)return t;s.push(t)}return s},set:function(e,t){var n,r,i=e.options,o=S.makeArray(t),a=i.length;while(a--)((r=i[a]).selected=-1<S.inArray(S.valHooks.option.get(r),o))&&(n=!0);return n||(e.selectedIndex=-1),o}}}}),S.each(["radio","checkbox"],function(){S.valHooks[this]={set:function(e,t){if(Array.isArray(t))return e.checked=-1<S.inArray(S(e).val(),t)}},y.checkOn||(S.valHooks[this].get=function(e){return null===e.getAttribute("value")?"on":e.value})}),y.focusin="onfocusin"in C;var bt=/^(?:focusinfocus|focusoutblur)$/,wt=function(e){e.stopPropagation()};S.extend(S.event,{trigger:function(e,t,n,r){var i,o,a,s,u,l,c,f,p=[n||E],d=v.call(e,"type")?e.type:e,h=v.call(e,"namespace")?e.namespace.split("."):[];if(o=f=a=n=n||E,3!==n.nodeType&&8!==n.nodeType&&!bt.test(d+S.event.triggered)&&(-1<d.indexOf(".")&&(d=(h=d.split(".")).shift(),h.sort()),u=d.indexOf(":")<0&&"on"+d,(e=e[S.expando]?e:new S.Event(d,"object"==typeof e&&e)).isTrigger=r?2:3,e.namespace=h.join("."),e.rnamespace=e.namespace?new RegExp("(^|\\.)"+h.join("\\.(?:.*\\.|)")+"(\\.|$)"):null,e.result=void 0,e.target||(e.target=n),t=null==t?[e]:S.makeArray(t,[e]),c=S.event.special[d]||{},r||!c.trigger||!1!==c.trigger.apply(n,t))){if(!r&&!c.noBubble&&!x(n)){for(s=c.delegateType||d,bt.test(s+d)||(o=o.parentNode);o;o=o.parentNode)p.push(o),a=o;a===(n.ownerDocument||E)&&p.push(a.defaultView||a.parentWindow||C)}i=0;while((o=p[i++])&&!e.isPropagationStopped())f=o,e.type=1<i?s:c.bindType||d,(l=(Y.get(o,"events")||Object.create(null))[e.type]&&Y.get(o,"handle"))&&l.apply(o,t),(l=u&&o[u])&&l.apply&&V(o)&&(e.result=l.apply(o,t),!1===e.result&&e.preventDefault());return e.type=d,r||e.isDefaultPrevented()||c._default&&!1!==c._default.apply(p.pop(),t)||!V(n)||u&&m(n[d])&&!x(n)&&((a=n[u])&&(n[u]=null),S.event.triggered=d,e.isPropagationStopped()&&f.addEventListener(d,wt),n[d](),e.isPropagationStopped()&&f.removeEventListener(d,wt),S.event.triggered=void 0,a&&(n[u]=a)),e.result}},simulate:function(e,t,n){var r=S.extend(new S.Event,n,{type:e,isSimulated:!0});S.event.trigger(r,null,t)}}),S.fn.extend({trigger:function(e,t){return this.each(function(){S.event.trigger(e,t,this)})},triggerHandler:function(e,t){var n=this[0];if(n)return S.event.trigger(e,t,n,!0)}}),y.focusin||S.each({focus:"focusin",blur:"focusout"},function(n,r){var i=function(e){S.event.simulate(r,e.target,S.event.fix(e))};S.event.special[r]={setup:function(){var e=this.ownerDocument||this.document||this,t=Y.access(e,r);t||e.addEventListener(n,i,!0),Y.access(e,r,(t||0)+1)},teardown:function(){var e=this.ownerDocument||this.document||this,t=Y.access(e,r)-1;t?Y.access(e,r,t):(e.removeEventListener(n,i,!0),Y.remove(e,r))}}});var Tt=C.location,Ct={guid:Date.now()},Et=/\?/;S.parseXML=function(e){var t;if(!e||"string"!=typeof e)return null;try{t=(new C.DOMParser).parseFromString(e,"text/xml")}catch(e){t=void 0}return t&&!t.getElementsByTagName("parsererror").length||S.error("Invalid XML: "+e),t};var St=/\[\]$/,kt=/\r?\n/g,At=/^(?:submit|button|image|reset|file)$/i,Nt=/^(?:input|select|textarea|keygen)/i;function Dt(n,e,r,i){var t;if(Array.isArray(e))S.each(e,function(e,t){r||St.test(n)?i(n,t):Dt(n+"["+("object"==typeof t&&null!=t?e:"")+"]",t,r,i)});else if(r||"object"!==w(e))i(n,e);else for(t in e)Dt(n+"["+t+"]",e[t],r,i)}S.param=function(e,t){var n,r=[],i=function(e,t){var n=m(t)?t():t;r[r.length]=encodeURIComponent(e)+"="+encodeURIComponent(null==n?"":n)};if(null==e)return"";if(Array.isArray(e)||e.jquery&&!S.isPlainObject(e))S.each(e,function(){i(this.name,this.value)});else for(n in e)Dt(n,e[n],t,i);return r.join("&")},S.fn.extend({serialize:function(){return S.param(this.serializeArray())},serializeArray:function(){return this.map(function(){var e=S.prop(this,"elements");return e?S.makeArray(e):this}).filter(function(){var e=this.type;return this.name&&!S(this).is(":disabled")&&Nt.test(this.nodeName)&&!At.test(e)&&(this.checked||!pe.test(e))}).map(function(e,t){var n=S(this).val();return null==n?null:Array.isArray(n)?S.map(n,function(e){return{name:t.name,value:e.replace(kt,"\r\n")}}):{name:t.name,value:n.replace(kt,"\r\n")}}).get()}});var jt=/%20/g,qt=/#.*$/,Lt=/([?&])_=[^&]*/,Ht=/^(.*?):[ \t]*([^\r\n]*)$/gm,Ot=/^(?:GET|HEAD)$/,Pt=/^\/\//,Rt={},Mt={},It="*/".concat("*"),Wt=E.createElement("a");function Ft(o){return function(e,t){"string"!=typeof e&&(t=e,e="*");var n,r=0,i=e.toLowerCase().match(P)||[];if(m(t))while(n=i[r++])"+"===n[0]?(n=n.slice(1)||"*",(o[n]=o[n]||[]).unshift(t)):(o[n]=o[n]||[]).push(t)}}function Bt(t,i,o,a){var s={},u=t===Mt;function l(e){var r;return s[e]=!0,S.each(t[e]||[],function(e,t){var n=t(i,o,a);return"string"!=typeof n||u||s[n]?u?!(r=n):void 0:(i.dataTypes.unshift(n),l(n),!1)}),r}return l(i.dataTypes[0])||!s["*"]&&l("*")}function $t(e,t){var n,r,i=S.ajaxSettings.flatOptions||{};for(n in t)void 0!==t[n]&&((i[n]?e:r||(r={}))[n]=t[n]);return r&&S.extend(!0,e,r),e}Wt.href=Tt.href,S.extend({active:0,lastModified:{},etag:{},ajaxSettings:{url:Tt.href,type:"GET",isLocal:/^(?:about|app|app-storage|.+-extension|file|res|widget):$/.test(Tt.protocol),global:!0,processData:!0,async:!0,contentType:"application/x-www-form-urlencoded; charset=UTF-8",accepts:{"*":It,text:"text/plain",html:"text/html",xml:"application/xml, text/xml",json:"application/json, text/javascript"},contents:{xml:/\bxml\b/,html:/\bhtml/,json:/\bjson\b/},responseFields:{xml:"responseXML",text:"responseText",json:"responseJSON"},converters:{"* text":String,"text html":!0,"text json":JSON.parse,"text xml":S.parseXML},flatOptions:{url:!0,context:!0}},ajaxSetup:function(e,t){return t?$t($t(e,S.ajaxSettings),t):$t(S.ajaxSettings,e)},ajaxPrefilter:Ft(Rt),ajaxTransport:Ft(Mt),ajax:function(e,t){"object"==typeof e&&(t=e,e=void 0),t=t||{};var c,f,p,n,d,r,h,g,i,o,v=S.ajaxSetup({},t),y=v.context||v,m=v.context&&(y.nodeType||y.jquery)?S(y):S.event,x=S.Deferred(),b=S.Callbacks("once memory"),w=v.statusCode||{},a={},s={},u="canceled",T={readyState:0,getResponseHeader:function(e){var t;if(h){if(!n){n={};while(t=Ht.exec(p))n[t[1].toLowerCase()+" "]=(n[t[1].toLowerCase()+" "]||[]).concat(t[2])}t=n[e.toLowerCase()+" "]}return null==t?null:t.join(", ")},getAllResponseHeaders:function(){return h?p:null},setRequestHeader:function(e,t){return null==h&&(e=s[e.toLowerCase()]=s[e.toLowerCase()]||e,a[e]=t),this},overrideMimeType:function(e){return null==h&&(v.mimeType=e),this},statusCode:function(e){var t;if(e)if(h)T.always(e[T.status]);else for(t in e)w[t]=[w[t],e[t]];return this},abort:function(e){var t=e||u;return c&&c.abort(t),l(0,t),this}};if(x.promise(T),v.url=((e||v.url||Tt.href)+"").replace(Pt,Tt.protocol+"//"),v.type=t.method||t.type||v.method||v.type,v.dataTypes=(v.dataType||"*").toLowerCase().match(P)||[""],null==v.crossDomain){r=E.createElement("a");try{r.href=v.url,r.href=r.href,v.crossDomain=Wt.protocol+"//"+Wt.host!=r.protocol+"//"+r.host}catch(e){v.crossDomain=!0}}if(v.data&&v.processData&&"string"!=typeof v.data&&(v.data=S.param(v.data,v.traditional)),Bt(Rt,v,t,T),h)return T;for(i in(g=S.event&&v.global)&&0==S.active++&&S.event.trigger("ajaxStart"),v.type=v.type.toUpperCase(),v.hasContent=!Ot.test(v.type),f=v.url.replace(qt,""),v.hasContent?v.data&&v.processData&&0===(v.contentType||"").indexOf("application/x-www-form-urlencoded")&&(v.data=v.data.replace(jt,"+")):(o=v.url.slice(f.length),v.data&&(v.processData||"string"==typeof v.data)&&(f+=(Et.test(f)?"&":"?")+v.data,delete v.data),!1===v.cache&&(f=f.replace(Lt,"$1"),o=(Et.test(f)?"&":"?")+"_="+Ct.guid+++o),v.url=f+o),v.ifModified&&(S.lastModified[f]&&T.setRequestHeader("If-Modified-Since",S.lastModified[f]),S.etag[f]&&T.setRequestHeader("If-None-Match",S.etag[f])),(v.data&&v.hasContent&&!1!==v.contentType||t.contentType)&&T.setRequestHeader("Content-Type",v.contentType),T.setRequestHeader("Accept",v.dataTypes[0]&&v.accepts[v.dataTypes[0]]?v.accepts[v.dataTypes[0]]+("*"!==v.dataTypes[0]?", "+It+"; q=0.01":""):v.accepts["*"]),v.headers)T.setRequestHeader(i,v.headers[i]);if(v.beforeSend&&(!1===v.beforeSend.call(y,T,v)||h))return T.abort();if(u="abort",b.add(v.complete),T.done(v.success),T.fail(v.error),c=Bt(Mt,v,t,T)){if(T.readyState=1,g&&m.trigger("ajaxSend",[T,v]),h)return T;v.async&&0<v.timeout&&(d=C.setTimeout(function(){T.abort("timeout")},v.timeout));try{h=!1,c.send(a,l)}catch(e){if(h)throw e;l(-1,e)}}else l(-1,"No Transport");function l(e,t,n,r){var i,o,a,s,u,l=t;h||(h=!0,d&&C.clearTimeout(d),c=void 0,p=r||"",T.readyState=0<e?4:0,i=200<=e&&e<300||304===e,n&&(s=function(e,t,n){var r,i,o,a,s=e.contents,u=e.dataTypes;while("*"===u[0])u.shift(),void 0===r&&(r=e.mimeType||t.getResponseHeader("Content-Type"));if(r)for(i in s)if(s[i]&&s[i].test(r)){u.unshift(i);break}if(u[0]in n)o=u[0];else{for(i in n){if(!u[0]||e.converters[i+" "+u[0]]){o=i;break}a||(a=i)}o=o||a}if(o)return o!==u[0]&&u.unshift(o),n[o]}(v,T,n)),!i&&-1<S.inArray("script",v.dataTypes)&&(v.converters["text script"]=function(){}),s=function(e,t,n,r){var i,o,a,s,u,l={},c=e.dataTypes.slice();if(c[1])for(a in e.converters)l[a.toLowerCase()]=e.converters[a];o=c.shift();while(o)if(e.responseFields[o]&&(n[e.responseFields[o]]=t),!u&&r&&e.dataFilter&&(t=e.dataFilter(t,e.dataType)),u=o,o=c.shift())if("*"===o)o=u;else if("*"!==u&&u!==o){if(!(a=l[u+" "+o]||l["* "+o]))for(i in l)if((s=i.split(" "))[1]===o&&(a=l[u+" "+s[0]]||l["* "+s[0]])){!0===a?a=l[i]:!0!==l[i]&&(o=s[0],c.unshift(s[1]));break}if(!0!==a)if(a&&e["throws"])t=a(t);else try{t=a(t)}catch(e){return{state:"parsererror",error:a?e:"No conversion from "+u+" to "+o}}}return{state:"success",data:t}}(v,s,T,i),i?(v.ifModified&&((u=T.getResponseHeader("Last-Modified"))&&(S.lastModified[f]=u),(u=T.getResponseHeader("etag"))&&(S.etag[f]=u)),204===e||"HEAD"===v.type?l="nocontent":304===e?l="notmodified":(l=s.state,o=s.data,i=!(a=s.error))):(a=l,!e&&l||(l="error",e<0&&(e=0))),T.status=e,T.statusText=(t||l)+"",i?x.resolveWith(y,[o,l,T]):x.rejectWith(y,[T,l,a]),T.statusCode(w),w=void 0,g&&m.trigger(i?"ajaxSuccess":"ajaxError",[T,v,i?o:a]),b.fireWith(y,[T,l]),g&&(m.trigger("ajaxComplete",[T,v]),--S.active||S.event.trigger("ajaxStop")))}return T},getJSON:function(e,t,n){return S.get(e,t,n,"json")},getScript:function(e,t){return S.get(e,void 0,t,"script")}}),S.each(["get","post"],function(e,i){S[i]=function(e,t,n,r){return m(t)&&(r=r||n,n=t,t=void 0),S.ajax(S.extend({url:e,type:i,dataType:r,data:t,success:n},S.isPlainObject(e)&&e))}}),S.ajaxPrefilter(function(e){var t;for(t in e.headers)"content-type"===t.toLowerCase()&&(e.contentType=e.headers[t]||"")}),S._evalUrl=function(e,t,n){return S.ajax({url:e,type:"GET",dataType:"script",cache:!0,async:!1,global:!1,converters:{"text script":function(){}},dataFilter:function(e){S.globalEval(e,t,n)}})},S.fn.extend({wrapAll:function(e){var t;return this[0]&&(m(e)&&(e=e.call(this[0])),t=S(e,this[0].ownerDocument).eq(0).clone(!0),this[0].parentNode&&t.insertBefore(this[0]),t.map(function(){var e=this;while(e.firstElementChild)e=e.firstElementChild;return e}).append(this)),this},wrapInner:function(n){return m(n)?this.each(function(e){S(this).wrapInner(n.call(this,e))}):this.each(function(){var e=S(this),t=e.contents();t.length?t.wrapAll(n):e.append(n)})},wrap:function(t){var n=m(t);return this.each(function(e){S(this).wrapAll(n?t.call(this,e):t)})},unwrap:function(e){return this.parent(e).not("body").each(function(){S(this).replaceWith(this.childNodes)}),this}}),S.expr.pseudos.hidden=function(e){return!S.expr.pseudos.visible(e)},S.expr.pseudos.visible=function(e){return!!(e.offsetWidth||e.offsetHeight||e.getClientRects().length)},S.ajaxSettings.xhr=function(){try{return new C.XMLHttpRequest}catch(e){}};var _t={0:200,1223:204},zt=S.ajaxSettings.xhr();y.cors=!!zt&&"withCredentials"in zt,y.ajax=zt=!!zt,S.ajaxTransport(function(i){var o,a;if(y.cors||zt&&!i.crossDomain)return{send:function(e,t){var n,r=i.xhr();if(r.open(i.type,i.url,i.async,i.username,i.password),i.xhrFields)for(n in i.xhrFields)r[n]=i.xhrFields[n];for(n in i.mimeType&&r.overrideMimeType&&r.overrideMimeType(i.mimeType),i.crossDomain||e["X-Requested-With"]||(e["X-Requested-With"]="XMLHttpRequest"),e)r.setRequestHeader(n,e[n]);o=function(e){return function(){o&&(o=a=r.onload=r.onerror=r.onabort=r.ontimeout=r.onreadystatechange=null,"abort"===e?r.abort():"error"===e?"number"!=typeof r.status?t(0,"error"):t(r.status,r.statusText):t(_t[r.status]||r.status,r.statusText,"text"!==(r.responseType||"text")||"string"!=typeof r.responseText?{binary:r.response}:{text:r.responseText},r.getAllResponseHeaders()))}},r.onload=o(),a=r.onerror=r.ontimeout=o("error"),void 0!==r.onabort?r.onabort=a:r.onreadystatechange=function(){4===r.readyState&&C.setTimeout(function(){o&&a()})},o=o("abort");try{r.send(i.hasContent&&i.data||null)}catch(e){if(o)throw e}},abort:function(){o&&o()}}}),S.ajaxPrefilter(function(e){e.crossDomain&&(e.contents.script=!1)}),S.ajaxSetup({accepts:{script:"text/javascript, application/javascript, application/ecmascript, application/x-ecmascript"},contents:{script:/\b(?:java|ecma)script\b/},converters:{"text script":function(e){return S.globalEval(e),e}}}),S.ajaxPrefilter("script",function(e){void 0===e.cache&&(e.cache=!1),e.crossDomain&&(e.type="GET")}),S.ajaxTransport("script",function(n){var r,i;if(n.crossDomain||n.scriptAttrs)return{send:function(e,t){r=S("<script>").attr(n.scriptAttrs||{}).prop({charset:n.scriptCharset,src:n.url}).on("load error",i=function(e){r.remove(),i=null,e&&t("error"===e.type?404:200,e.type)}),E.head.appendChild(r[0])},abort:function(){i&&i()}}});var Ut,Xt=[],Vt=/(=)\?(?=&|$)|\?\?/;S.ajaxSetup({jsonp:"callback",jsonpCallback:function(){var e=Xt.pop()||S.expando+"_"+Ct.guid++;return this[e]=!0,e}}),S.ajaxPrefilter("json jsonp",function(e,t,n){var r,i,o,a=!1!==e.jsonp&&(Vt.test(e.url)?"url":"string"==typeof e.data&&0===(e.contentType||"").indexOf("application/x-www-form-urlencoded")&&Vt.test(e.data)&&"data");if(a||"jsonp"===e.dataTypes[0])return r=e.jsonpCallback=m(e.jsonpCallback)?e.jsonpCallback():e.jsonpCallback,a?e[a]=e[a].replace(Vt,"$1"+r):!1!==e.jsonp&&(e.url+=(Et.test(e.url)?"&":"?")+e.jsonp+"="+r),e.converters["script json"]=function(){return o||S.error(r+" was not called"),o[0]},e.dataTypes[0]="json",i=C[r],C[r]=function(){o=arguments},n.always(function(){void 0===i?S(C).removeProp(r):C[r]=i,e[r]&&(e.jsonpCallback=t.jsonpCallback,Xt.push(r)),o&&m(i)&&i(o[0]),o=i=void 0}),"script"}),y.createHTMLDocument=((Ut=E.implementation.createHTMLDocument("").body).innerHTML="<form></form><form></form>",2===Ut.childNodes.length),S.parseHTML=function(e,t,n){return"string"!=typeof e?[]:("boolean"==typeof t&&(n=t,t=!1),t||(y.createHTMLDocument?((r=(t=E.implementation.createHTMLDocument("")).createElement("base")).href=E.location.href,t.head.appendChild(r)):t=E),o=!n&&[],(i=N.exec(e))?[t.createElement(i[1])]:(i=xe([e],t,o),o&&o.length&&S(o).remove(),S.merge([],i.childNodes)));var r,i,o},S.fn.load=function(e,t,n){var r,i,o,a=this,s=e.indexOf(" ");return-1<s&&(r=vt(e.slice(s)),e=e.slice(0,s)),m(t)?(n=t,t=void 0):t&&"object"==typeof t&&(i="POST"),0<a.length&&S.ajax({url:e,type:i||"GET",dataType:"html",data:t}).done(function(e){o=arguments,a.html(r?S("<div>").append(S.parseHTML(e)).find(r):e)}).always(n&&function(e,t){a.each(function(){n.apply(this,o||[e.responseText,t,e])})}),this},S.expr.pseudos.animated=function(t){return S.grep(S.timers,function(e){return t===e.elem}).length},S.offset={setOffset:function(e,t,n){var r,i,o,a,s,u,l=S.css(e,"position"),c=S(e),f={};"static"===l&&(e.style.position="relative"),s=c.offset(),o=S.css(e,"top"),u=S.css(e,"left"),("absolute"===l||"fixed"===l)&&-1<(o+u).indexOf("auto")?(a=(r=c.position()).top,i=r.left):(a=parseFloat(o)||0,i=parseFloat(u)||0),m(t)&&(t=t.call(e,n,S.extend({},s))),null!=t.top&&(f.top=t.top-s.top+a),null!=t.left&&(f.left=t.left-s.left+i),"using"in t?t.using.call(e,f):("number"==typeof f.top&&(f.top+="px"),"number"==typeof f.left&&(f.left+="px"),c.css(f))}},S.fn.extend({offset:function(t){if(arguments.length)return void 0===t?this:this.each(function(e){S.offset.setOffset(this,t,e)});var e,n,r=this[0];return r?r.getClientRects().length?(e=r.getBoundingClientRect(),n=r.ownerDocument.defaultView,{top:e.top+n.pageYOffset,left:e.left+n.pageXOffset}):{top:0,left:0}:void 0},position:function(){if(this[0]){var e,t,n,r=this[0],i={top:0,left:0};if("fixed"===S.css(r,"position"))t=r.getBoundingClientRect();else{t=this.offset(),n=r.ownerDocument,e=r.offsetParent||n.documentElement;while(e&&(e===n.body||e===n.documentElement)&&"static"===S.css(e,"position"))e=e.parentNode;e&&e!==r&&1===e.nodeType&&((i=S(e).offset()).top+=S.css(e,"borderTopWidth",!0),i.left+=S.css(e,"borderLeftWidth",!0))}return{top:t.top-i.top-S.css(r,"marginTop",!0),left:t.left-i.left-S.css(r,"marginLeft",!0)}}},offsetParent:function(){return this.map(function(){var e=this.offsetParent;while(e&&"static"===S.css(e,"position"))e=e.offsetParent;return e||re})}}),S.each({scrollLeft:"pageXOffset",scrollTop:"pageYOffset"},function(t,i){var o="pageYOffset"===i;S.fn[t]=function(e){return $(this,function(e,t,n){var r;if(x(e)?r=e:9===e.nodeType&&(r=e.defaultView),void 0===n)return r?r[i]:e[t];r?r.scrollTo(o?r.pageXOffset:n,o?n:r.pageYOffset):e[t]=n},t,e,arguments.length)}}),S.each(["top","left"],function(e,n){S.cssHooks[n]=$e(y.pixelPosition,function(e,t){if(t)return t=Be(e,n),Me.test(t)?S(e).position()[n]+"px":t})}),S.each({Height:"height",Width:"width"},function(a,s){S.each({padding:"inner"+a,content:s,"":"outer"+a},function(r,o){S.fn[o]=function(e,t){var n=arguments.length&&(r||"boolean"!=typeof e),i=r||(!0===e||!0===t?"margin":"border");return $(this,function(e,t,n){var r;return x(e)?0===o.indexOf("outer")?e["inner"+a]:e.document.documentElement["client"+a]:9===e.nodeType?(r=e.documentElement,Math.max(e.body["scroll"+a],r["scroll"+a],e.body["offset"+a],r["offset"+a],r["client"+a])):void 0===n?S.css(e,t,i):S.style(e,t,n,i)},s,n?e:void 0,n)}})}),S.each(["ajaxStart","ajaxStop","ajaxComplete","ajaxError","ajaxSuccess","ajaxSend"],function(e,t){S.fn[t]=function(e){return this.on(t,e)}}),S.fn.extend({bind:function(e,t,n){return this.on(e,null,t,n)},unbind:function(e,t){return this.off(e,null,t)},delegate:function(e,t,n,r){return this.on(t,e,n,r)},undelegate:function(e,t,n){return 1===arguments.length?this.off(e,"**"):this.off(t,e||"**",n)},hover:function(e,t){return this.mouseenter(e).mouseleave(t||e)}}),S.each("blur focus focusin focusout resize scroll click dblclick mousedown mouseup mousemove mouseover mouseout mouseenter mouseleave change select submit keydown keypress keyup contextmenu".split(" "),function(e,n){S.fn[n]=function(e,t){return 0<arguments.length?this.on(n,null,e,t):this.trigger(n)}});var Gt=/^[\s\uFEFF\xA0]+|[\s\uFEFF\xA0]+$/g;S.proxy=function(e,t){var n,r,i;if("string"==typeof t&&(n=e[t],t=e,e=n),m(e))return r=s.call(arguments,2),(i=function(){return e.apply(t||this,r.concat(s.call(arguments)))}).guid=e.guid=e.guid||S.guid++,i},S.holdReady=function(e){e?S.readyWait++:S.ready(!0)},S.isArray=Array.isArray,S.parseJSON=JSON.parse,S.nodeName=A,S.isFunction=m,S.isWindow=x,S.camelCase=X,S.type=w,S.now=Date.now,S.isNumeric=function(e){var t=S.type(e);return("number"===t||"string"===t)&&!isNaN(e-parseFloat(e))},S.trim=function(e){return null==e?"":(e+"").replace(Gt,"")},"function"==typeof define&&define.amd&&define("jquery",[],function(){return S});var Yt=C.jQuery,Qt=C.$;return S.noConflict=function(e){return C.$===S&&(C.$=Qt),e&&C.jQuery===S&&(C.jQuery=Yt),S},"undefined"==typeof e&&(C.jQuery=C.$=S),S});
```
