## Supplementary material for "scHiCTools: a computational toolbox for analyzing single-cell Hi-C data": S4: theme.js

```
/* sphinx_rtd_theme version 0.4.3 | MIT license */
/* Built 20190212 16:02 */
require=function r(s,a,l){function c(e,n){if(!a[e]){if(!s[e]){var i="function"==typeof require&&require;if(!n&&i)return i(e,!0);if(u)return u(e,!0);var t=new Error("Cannot find module '"+e+"'");throw t.code="MODULE_NOT_FOUND",t}var o=a[e]={exports:{}};s[e][0].call(o.exports,function(n){return c(s[e][1][n]||n)},o,o.exports,r,s,a,l)}return a[e].exports}for(var u="function"==typeof require&&require,n=0;n<l.length;n++)c(l[n]);return c}({"sphinx-rtd-theme":[function(n,e,i){var jQuery="undefined"!=typeof window?window.jQuery:n("jquery");e.exports.ThemeNav={navBar:null,win:null,winScroll:!1,winResize:!1,linkScroll:!1,winPosition:0,winHeight:null,docHeight:null,isRunning:!1,enable:function(e){var i=this;void 0===e&&(e=!0),i.isRunning||(i.isRunning=!0,jQuery(function(n){i.init(n),i.reset(),i.win.on("hashchange",i.reset),e&&i.win.on("scroll",function(){i.linkScroll||i.winScroll||(i.winScroll=!0,requestAnimationFrame(function(){i.onScroll()}))}),i.win.on("resize",function(){i.winResize||(i.winResize=!0,requestAnimationFrame(function(){i.onResize()}))}),i.onResize()}))},enableSticky:function(){this.enable(!0)},init:function(i){i(document);var t=this;this.navBar=i("div.wy-side-scroll:first"),this.win=i(window),i(document).on("click","[data-toggle='wy-nav-top']",function(){i("[data-toggle='wy-nav-shift']").toggleClass("shift"),i("[data-toggle='rst-versions']").toggleClass("shift")}).on("click",".wy-menu-vertical .current ul li a",function(){var n=i(this);i("[data-toggle='wy-nav-shift']").removeClass("shift"),i("[data-toggle='rst-versions']").toggleClass("shift"),t.toggleCurrent(n),t.hashChange()}).on("click","[data-toggle='rst-current-version']",function(){i("[data-toggle='rst-versions']").toggleClass("shift-up")}),i("table.docutils:not(.field-list,.footnote,.citation)").wrap("<div class='wy-table-responsive'></div>"),i("table.docutils.footnote").wrap("<div class='wy-table-responsive footnote'></div>"),i("table.docutils.citation").wrap("<div class='wy-table-responsive citation'></div>"),i(".wy-menu-vertical ul").not(".simple").siblings("a").each(function(){var e=i(this);expand=i('<span class="toctree-expand"></span>'),expand.on("click",function(n){return t.toggleCurrent(e),n.stopPropagation(),!1}),e.prepend(expand)})},reset:function(){var n=encodeURI(window.location.hash)||"#";try{var e=$(".wy-menu-vertical"),i=e.find('[href="'+n+'"]');if(0===i.length){var t=$('.document [id="'+n.substring(1)+'"]').closest("div.section");0===(i=e.find('[href="#'+t.attr("id")+'"]')).length&&(i=e.find('[href="#"]'))}0<i.length&&($(".wy-menu-vertical .current").removeClass("current"),i.addClass("current"),i.closest("li.toctree-l1").addClass("current"),i.closest("li.toctree-l1").parent().addClass("current"),i.closest("li.toctree-l1").addClass("current"),i.closest("li.toctree-l2").addClass("current"),i.closest("li.toctree-l3").addClass("current"),i.closest("li.toctree-l4").addClass("current"),i[0].scrollIntoView())}catch(o){console.log("Error expanding nav for anchor",o)}},onScroll:function(){this.winScroll=!1;var n=this.win.scrollTop(),e=n+this.winHeight,i=this.navBar.scrollTop()+(n-this.winPosition);n<0||e>this.docHeight||(this.navBar.scrollTop(i),this.winPosition=n)},onResize:function(){this.winResize=!1,this.winHeight=this.win.height(),this.docHeight=$(document).height()},hashChange:function(){this.linkScroll=!0,this.win.one("hashchange",function(){this.linkScroll=!1})},toggleCurrent:function(n){var e=n.closest("li");e.siblings("li.current").removeClass("current"),e.siblings().find("li.current").removeClass("current"),e.find("> ul li.current").removeClass("current"),e.toggleClass("current")}},"undefined"!=typeof window&&(window.SphinxRtdTheme={Navigation:e.exports.ThemeNav,StickyNav:e.exports.ThemeNav}),function(){for(var r=0,n=["ms","moz","webkit","o"],e=0;e<n.length&&!window.requestAnimationFrame;++e)window.requestAnimationFrame=window[n[e]+"RequestAnimationFrame"],window.cancelAnimationFrame=window[n[e]+"CancelAnimationFrame"]||window[n[e]+"CancelRequestAnimationFrame"];window.requestAnimationFrame||(window.requestAnimationFrame=function(n,e){var i=(new Date).getTime(),t=Math.max(0,16-(i-r)),o=window.setTimeout(function(){n(i+t)},t);return r=i+t,o}),window.cancelAnimationFrame||(window.cancelAnimationFrame=function(n){clearTimeout(n)})}()},{jquery:"jquery"}]},{},["sphinx-rtd-theme"]);
```
