## Supplementary material for "scHiCTools: a computational toolbox for analyzing single-cell Hi-C data": S4: language_data.js

```
/*
 * language_data.js
 * ~~~~~~~~~~~~~~~~
 *
 * This script contains the language-specific data used by searchtools.js,
 * namely the list of stopwords, stemmer, scorer and splitter.
 *
 * :copyright: Copyright 2007-2020 by the Sphinx team, see AUTHORS.
 * :license: BSD, see LICENSE for details.
 *
 */

var stopwords = ["a","and","are","as","at","be","but","by","for","if","in","into","is","it","near","no","not","of","on","or","such","that","the","their","then","there","these","they","this","to","was","will","with"];


/* Non-minified version JS is _stemmer.js if file is provided */ 
/**
 * Porter Stemmer
 */
var Stemmer = function() {

  var step2list = {
    ational: 'ate',
    tional: 'tion',
    enci: 'ence',
    anci: 'ance',
    izer: 'ize',
    bli: 'ble',
    alli: 'al',
    entli: 'ent',
    eli: 'e',
    ousli: 'ous',
    ization: 'ize',
    ation: 'ate',
    ator: 'ate',
    alism: 'al',
    iveness: 'ive',
    fulness: 'ful',
    ousness: 'ous',
    aliti: 'al',
    iviti: 'ive',
    biliti: 'ble',
    logi: 'log'
  };

  var step3list = {
    icate: 'ic',
    ative: '',
    alize: 'al',
    iciti: 'ic',
    ical: 'ic',
    ful: '',
    ness: ''
  };

  var c = "[^aeiou]";          // consonant
  var v = "[aeiouy]";          // vowel
  var C = c + "[^aeiouy]*";    // consonant sequence
  var V = v + "[aeiou]*";      // vowel sequence

  var mgr0 = "^(" + C + ")?" + V + C;                      // [C]VC... is m>0
  var meq1 = "^(" + C + ")?" + V + C + "(" + V + ")?$";    // [C]VC[V] is m=1
  var mgr1 = "^(" + C + ")?" + V + C + V + C;              // [C]VCVC... is m>1
  var s_v   = "^(" + C + ")?" + v;                         // vowel in stem

  this.stemWord = function (w) {
    var stem;
    var suffix;
    var firstch;
    var origword = w;

    if (w.length < 3)
      return w;

    var re;
    var re2;
    var re3;
    var re4;

    firstch = w.substr(0,1);
    if (firstch == "y")
      w = firstch.toUpperCase() + w.substr(1);

    // Step 1a
    re = /^(.+?)(ss|i)es$/;
    re2 = /^(.+?)([^s])s$/;

    if (re.test(w))
      w = w.replace(re,"$1$2");
    else if (re2.test(w))
      w = w.replace(re2,"$1$2");

    // Step 1b
    re = /^(.+?)eed$/;
    re2 = /^(.+?)(ed|ing)$/;
    if (re.test(w)) {
      var fp = re.exec(w);
      re = new RegExp(mgr0);
      if (re.test(fp[1])) {
        re = /.$/;
        w = w.replace(re,"");
      }
    }
    else if (re2.test(w)) {
      var fp = re2.exec(w);
      stem = fp[1];
      re2 = new RegExp(s_v);
      if (re2.test(stem)) {
        w = stem;
        re2 = /(at|bl|iz)$/;
        re3 = new RegExp("([^aeiouylsz])\\1$");
        re4 = new RegExp("^" + C + v + "[^aeiouwxy]$");
        if (re2.test(w))
          w = w + "e";
        else if (re3.test(w)) {
          re = /.$/;
          w = w.replace(re,"");
        }
        else if (re4.test(w))
          w = w + "e";
      }
    }

    // Step 1c
    re = /^(.+?)y$/;
    if (re.test(w)) {
      var fp = re.exec(w);
      stem = fp[1];
      re = new RegExp(s_v);
      if (re.test(stem))
        w = stem + "i";
    }

    // Step 2
    re = /^(.+?)(ational|tional|enci|anci|izer|bli|alli|entli|eli|ousli|ization|ation|ator|alism|iveness|fulness|ousness|aliti|iviti|biliti|logi)$/;
    if (re.test(w)) {
      var fp = re.exec(w);
      stem = fp[1];
      suffix = fp[2];
      re = new RegExp(mgr0);
      if (re.test(stem))
        w = stem + step2list[suffix];
    }

    // Step 3
    re = /^(.+?)(icate|ative|alize|iciti|ical|ful|ness)$/;
    if (re.test(w)) {
      var fp = re.exec(w);
      stem = fp[1];
      suffix = fp[2];
      re = new RegExp(mgr0);
      if (re.test(stem))
        w = stem + step3list[suffix];
    }

    // Step 4
    re = /^(.+?)(al|ance|ence|er|ic|able|ible|ant|ement|ment|ent|ou|ism|ate|iti|ous|ive|ize)$/;
    re2 = /^(.+?)(s|t)(ion)$/;
    if (re.test(w)) {
      var fp = re.exec(w);
      stem = fp[1];
      re = new RegExp(mgr1);
      if (re.test(stem))
        w = stem;
    }
    else if (re2.test(w)) {
      var fp = re2.exec(w);
      stem = fp[1] + fp[2];
      re2 = new RegExp(mgr1);
      if (re2.test(stem))
        w = stem;
    }

    // Step 5
    re = /^(.+?)e$/;
    if (re.test(w)) {
      var fp = re.exec(w);
      stem = fp[1];
      re = new RegExp(mgr1);
      re2 = new RegExp(meq1);
      re3 = new RegExp("^" + C + v + "[^aeiouwxy]$");
      if (re.test(stem) || (re2.test(stem) && !(re3.test(stem))))
        w = stem;
    }
    re = /ll$/;
    re2 = new RegExp(mgr1);
    if (re.test(w) && re2.test(w)) {
      re = /.$/;
      w = w.replace(re,"");
    }

    // and turn initial Y back to y
    if (firstch == "y")
      w = firstch.toLowerCase() + w.substr(1);
    return w;
  }
}


var splitChars = (function() {
    var result = {};
    var singles = [96, 180, 187, 191, 215, 247, 749, 885, 903, 907, 909, 930, 1014, 1648,
         1748, 1809, 2416, 2473, 2481, 2526, 2601, 2609, 2612, 2615, 2653, 2702,
         2706, 2729, 2737, 2740, 2857, 2865, 2868, 2910, 2928, 2948, 2961, 2971,
         2973, 3085, 3089, 3113, 3124, 3213, 3217, 3241, 3252, 3295, 3341, 3345,
         3369, 3506, 3516, 3633, 3715, 3721, 3736, 3744, 3748, 3750, 3756, 3761,
         3781, 3912, 4239, 4347, 4681, 4695, 4697, 4745, 4785, 4799, 4801, 4823,
         4881, 5760, 5901, 5997, 6313, 7405, 8024, 8026, 8028, 8030, 8117, 8125,
         8133, 8181, 8468, 8485, 8487, 8489, 8494, 8527, 11311, 11359, 11687, 11695,
         11703, 11711, 11719, 11727, 11735, 12448, 12539, 43010, 43014, 43019, 43587,
         43696, 43713, 64286, 64297, 64311, 64317, 64319, 64322, 64325, 65141];
    var i, j, start, end;
    for (i = 0; i < singles.length; i++) {
        result[singles[i]] = true;
    }
    var ranges = [[0, 47], [58, 64], [91, 94], [123, 169], [171, 177], [182, 184], [706, 709],
         [722, 735], [741, 747], [751, 879], [888, 889], [894, 901], [1154, 1161],
         [1318, 1328], [1367, 1368], [1370, 1376], [1416, 1487], [1515, 1519], [1523, 1568],
         [1611, 1631], [1642, 1645], [1750, 1764], [1767, 1773], [1789, 1790], [1792, 1807],
         [1840, 1868], [1958, 1968], [1970, 1983], [2027, 2035], [2038, 2041], [2043, 2047],
         [2070, 2073], [2075, 2083], [2085, 2087], [2089, 2307], [2362, 2364], [2366, 2383],
         [2385, 2391], [2402, 2405], [2419, 2424], [2432, 2436], [2445, 2446], [2449, 2450],
         [2483, 2485], [2490, 2492], [2494, 2509], [2511, 2523], [2530, 2533], [2546, 2547],
         [2554, 2564], [2571, 2574], [2577, 2578], [2618, 2648], [2655, 2661], [2672, 2673],
         [2677, 2692], [2746, 2748], [2750, 2767], [2769, 2783], [2786, 2789], [2800, 2820],
         [2829, 2830], [2833, 2834], [2874, 2876], [2878, 2907], [2914, 2917], [2930, 2946],
         [2955, 2957], [2966, 2968], [2976, 2978], [2981, 2983], [2987, 2989], [3002, 3023],
         [3025, 3045], [3059, 3076], [3130, 3132], [3134, 3159], [3162, 3167], [3170, 3173],
         [3184, 3191], [3199, 3204], [3258, 3260], [3262, 3293], [3298, 3301], [3312, 3332],
         [3386, 3388], [3390, 3423], [3426, 3429], [3446, 3449], [3456, 3460], [3479, 3481],
         [3518, 3519], [3527, 3584], [3636, 3647], [3655, 3663], [3674, 3712], [3717, 3718],
         [3723, 3724], [3726, 3731], [3752, 3753], [3764, 3772], [3774, 3775], [3783, 3791],
         [3802, 3803], [3806, 3839], [3841, 3871], [3892, 3903], [3949, 3975], [3980, 4095],
         [4139, 4158], [4170, 4175], [4182, 4185], [4190, 4192], [4194, 4196], [4199, 4205],
         [4209, 4212], [4226, 4237], [4250, 4255], [4294, 4303], [4349, 4351], [4686, 4687],
         [4702, 4703], [4750, 4751], [4790, 4791], [4806, 4807], [4886, 4887], [4955, 4968],
         [4989, 4991], [5008, 5023], [5109, 5120], [5741, 5742], [5787, 5791], [5867, 5869],
         [5873, 5887], [5906, 5919], [5938, 5951], [5970, 5983], [6001, 6015], [6068, 6102],
         [6104, 6107], [6109, 6111], [6122, 6127], [6138, 6159], [6170, 6175], [6264, 6271],
         [6315, 6319], [6390, 6399], [6429, 6469], [6510, 6511], [6517, 6527], [6572, 6592],
         [6600, 6607], [6619, 6655], [6679, 6687], [6741, 6783], [6794, 6799], [6810, 6822],
         [6824, 6916], [6964, 6980], [6988, 6991], [7002, 7042], [7073, 7085], [7098, 7167],
         [7204, 7231], [7242, 7244], [7294, 7400], [7410, 7423], [7616, 7679], [7958, 7959],
         [7966, 7967], [8006, 8007], [8014, 8015], [8062, 8063], [8127, 8129], [8141, 8143],
         [8148, 8149], [8156, 8159], [8173, 8177], [8189, 8303], [8306, 8307], [8314, 8318],
         [8330, 8335], [8341, 8449], [8451, 8454], [8456, 8457], [8470, 8472], [8478, 8483],
         [8506, 8507], [8512, 8516], [8522, 8525], [8586, 9311], [9372, 9449], [9472, 10101],
         [10132, 11263], [11493, 11498], [11503, 11516], [11518, 11519], [11558, 11567],
         [11622, 11630], [11632, 11647], [11671, 11679], [11743, 11822], [11824, 12292],
         [12296, 12320], [12330, 12336], [12342, 12343], [12349, 12352], [12439, 12444],
         [12544, 12548], [12590, 12592], [12687, 12689], [12694, 12703], [12728, 12783],
         [12800, 12831], [12842, 12880], [12896, 12927], [12938, 12976], [12992, 13311],
         [19894, 19967], [40908, 40959], [42125, 42191], [42238, 42239], [42509, 42511],
         [42540, 42559], [42592, 42593], [42607, 42622], [42648, 42655], [42736, 42774],
         [42784, 42785], [42889, 42890], [42893, 43002], [43043, 43055], [43062, 43071],
         [43124, 43137], [43188, 43215], [43226, 43249], [43256, 43258], [43260, 43263],
         [43302, 43311], [43335, 43359], [43389, 43395], [43443, 43470], [43482, 43519],
         [43561, 43583], [43596, 43599], [43610, 43615], [43639, 43641], [43643, 43647],
         [43698, 43700], [43703, 43704], [43710, 43711], [43715, 43738], [43742, 43967],
         [44003, 44015], [44026, 44031], [55204, 55215], [55239, 55242], [55292, 55295],
         [57344, 63743], [64046, 64047], [64110, 64111], [64218, 64255], [64263, 64274],
         [64280, 64284], [64434, 64466], [64830, 64847], [64912, 64913], [64968, 65007],
         [65020, 65135], [65277, 65295], [65306, 65312], [65339, 65344], [65371, 65381],
         [65471, 65473], [65480, 65481], [65488, 65489], [65496, 65497]];
    for (i = 0; i < ranges.length; i++) {
        start = ranges[i][0];
        end = ranges[i][1];
        for (j = start; j <= end; j++) {
            result[j] = true;
        }
    }
    return result;
})();

function splitQuery(query) {
    var result = [];
    var start = -1;
    for (var i = 0; i < query.length; i++) {
        if (splitChars[query.charCodeAt(i)]) {
            if (start !== -1) {
                result.push(query.slice(start, i));
                start = -1;
            }
        } else if (start === -1) {
            start = i;
        }
    }
    if (start !== -1) {
        result.push(query.slice(start));
    }
    return result;
}
```
