## Supplementary material for "scHiCTools: a computational toolbox for analyzing single-cell Hi-C data": S4: searchtools.js

```
/*
 * searchtools.js
 * ~~~~~~~~~~~~~~~~
 *
 * Sphinx JavaScript utilities for the full-text search.
 *
 * :copyright: Copyright 2007-2020 by the Sphinx team, see AUTHORS.
 * :license: BSD, see LICENSE for details.
 *
 */

if (!Scorer) {
  /**
   * Simple result scoring code.
   */
  var Scorer = {
    // Implement the following function to further tweak the score for each result
    // The function takes a result array [filename, title, anchor, descr, score]
    // and returns the new score.
    /*
    score: function(result) {
      return result[4];
    },
    */

    // query matches the full name of an object
    objNameMatch: 11,
    // or matches in the last dotted part of the object name
    objPartialMatch: 6,
    // Additive scores depending on the priority of the object
    objPrio: {0:  15,   // used to be importantResults
              1:  5,   // used to be objectResults
              2: -5},  // used to be unimportantResults
    //  Used when the priority is not in the mapping.
    objPrioDefault: 0,

    // query found in title
    title: 15,
    partialTitle: 7,
    // query found in terms
    term: 5,
    partialTerm: 2
  };
}

if (!splitQuery) {
  function splitQuery(query) {
    return query.split(/\s+/);
  }
}

/**
 * Search Module
 */
var Search = {

  _index : null,
  _queued_query : null,
  _pulse_status : -1,

  htmlToText : function(htmlString) {
      var htmlElement = document.createElement('span');
      htmlElement.innerHTML = htmlString;
      $(htmlElement).find('.headerlink').remove();
      docContent = $(htmlElement).find('[role=main]')[0];
      if(docContent === undefined) {
          console.warn("Content block not found. Sphinx search tries to obtain it " +
                       "via '[role=main]'. Could you check your theme or template.");
          return "";
      }
      return docContent.textContent || docContent.innerText;
  },

  init : function() {
      var params = $.getQueryParameters();
      if (params.q) {
          var query = params.q[0];
          $('input[name="q"]')[0].value = query;
          this.performSearch(query);
      }
  },

  loadIndex : function(url) {
    $.ajax({type: "GET", url: url, data: null,
            dataType: "script", cache: true,
            complete: function(jqxhr, textstatus) {
              if (textstatus != "success") {
                document.getElementById("searchindexloader").src = url;
              }
            }});
  },

  setIndex : function(index) {
    var q;
    this._index = index;
    if ((q = this._queued_query) !== null) {
      this._queued_query = null;
      Search.query(q);
    }
  },

  hasIndex : function() {
      return this._index !== null;
  },

  deferQuery : function(query) {
      this._queued_query = query;
  },

  stopPulse : function() {
      this._pulse_status = 0;
  },

  startPulse : function() {
    if (this._pulse_status >= 0)
        return;
    function pulse() {
      var i;
      Search._pulse_status = (Search._pulse_status + 1) % 4;
      var dotString = '';
      for (i = 0; i < Search._pulse_status; i++)
        dotString += '.';
      Search.dots.text(dotString);
      if (Search._pulse_status > -1)
        window.setTimeout(pulse, 500);
    }
    pulse();
  },

  /**
   * perform a search for something (or wait until index is loaded)
   */
  performSearch : function(query) {
    // create the required interface elements
    this.out = $('#search-results');
    this.title = $('<h2>' + _('Searching') + '</h2>').appendTo(this.out);
    this.dots = $('<span></span>').appendTo(this.title);
    this.status = $('<p class="search-summary">&nbsp;</p>').appendTo(this.out);
    this.output = $('<ul class="search"/>').appendTo(this.out);

    $('#search-progress').text(_('Preparing search...'));
    this.startPulse();

    // index already loaded, the browser was quick!
    if (this.hasIndex())
      this.query(query);
    else
      this.deferQuery(query);
  },

  /**
   * execute search (requires search index to be loaded)
   */
  query : function(query) {
    var i;

    // stem the searchterms and add them to the correct list
    var stemmer = new Stemmer();
    var searchterms = [];
    var excluded = [];
    var hlterms = [];
    var tmp = splitQuery(query);
    var objectterms = [];
    for (i = 0; i < tmp.length; i++) {
      if (tmp[i] !== "") {
          objectterms.push(tmp[i].toLowerCase());
      }

      if ($u.indexOf(stopwords, tmp[i].toLowerCase()) != -1 || tmp[i].match(/^\d+$/) ||
          tmp[i] === "") {
        // skip this "word"
        continue;
      }
      // stem the word
      var word = stemmer.stemWord(tmp[i].toLowerCase());
      // prevent stemmer from cutting word smaller than two chars
      if(word.length < 3 && tmp[i].length >= 3) {
        word = tmp[i];
      }
      var toAppend;
      // select the correct list
      if (word[0] == '-') {
        toAppend = excluded;
        word = word.substr(1);
      }
      else {
        toAppend = searchterms;
        hlterms.push(tmp[i].toLowerCase());
      }
      // only add if not already in the list
      if (!$u.contains(toAppend, word))
        toAppend.push(word);
    }
    var highlightstring = '?highlight=' + $.urlencode(hlterms.join(" "));

    // console.debug('SEARCH: searching for:');
    // console.info('required: ', searchterms);
    // console.info('excluded: ', excluded);

    // prepare search
    var terms = this._index.terms;
    var titleterms = this._index.titleterms;

    // array of [filename, title, anchor, descr, score]
    var results = [];
    $('#search-progress').empty();

    // lookup as object
    for (i = 0; i < objectterms.length; i++) {
      var others = [].concat(objectterms.slice(0, i),
                             objectterms.slice(i+1, objectterms.length));
      results = results.concat(this.performObjectSearch(objectterms[i], others));
    }

    // lookup as search terms in fulltext
    results = results.concat(this.performTermsSearch(searchterms, excluded, terms, titleterms));

    // let the scorer override scores with a custom scoring function
    if (Scorer.score) {
      for (i = 0; i < results.length; i++)
        results[i][4] = Scorer.score(results[i]);
    }

    // now sort the results by score (in opposite order of appearance, since the
    // display function below uses pop() to retrieve items) and then
    // alphabetically
    results.sort(function(a, b) {
      var left = a[4];
      var right = b[4];
      if (left > right) {
        return 1;
      } else if (left < right) {
        return -1;
      } else {
        // same score: sort alphabetically
        left = a[1].toLowerCase();
        right = b[1].toLowerCase();
        return (left > right) ? -1 : ((left < right) ? 1 : 0);
      }
    });

    // for debugging
    //Search.lastresults = results.slice();  // a copy
    //console.info('search results:', Search.lastresults);

    // print the results
    var resultCount = results.length;
    function displayNextItem() {
      // results left, load the summary and display it
      if (results.length) {
        var item = results.pop();
        var listItem = $('<li style="display:none"></li>');
        var requestUrl = "";
        var linkUrl = "";
        if (DOCUMENTATION_OPTIONS.BUILDER === 'dirhtml') {
          // dirhtml builder
          var dirname = item[0] + '/';
          if (dirname.match(/\/index\/$/)) {
            dirname = dirname.substring(0, dirname.length-6);
          } else if (dirname == 'index/') {
            dirname = '';
          }
          requestUrl = DOCUMENTATION_OPTIONS.URL_ROOT + dirname;
          linkUrl = requestUrl;

        } else {
          // normal html builders
          requestUrl = DOCUMENTATION_OPTIONS.URL_ROOT + item[0] + DOCUMENTATION_OPTIONS.FILE_SUFFIX;
          linkUrl = item[0] + DOCUMENTATION_OPTIONS.LINK_SUFFIX;
        }
        listItem.append($('<a/>').attr('href',
            linkUrl +
            highlightstring + item[2]).html(item[1]));
        if (item[3]) {
          listItem.append($('<span> (' + item[3] + ')</span>'));
          Search.output.append(listItem);
          listItem.slideDown(5, function() {
            displayNextItem();
          });
        } else if (DOCUMENTATION_OPTIONS.HAS_SOURCE) {
          $.ajax({url: requestUrl,
                  dataType: "text",
                  complete: function(jqxhr, textstatus) {
                    var data = jqxhr.responseText;
                    if (data !== '' && data !== undefined) {
                      listItem.append(Search.makeSearchSummary(data, searchterms, hlterms));
                    }
                    Search.output.append(listItem);
                    listItem.slideDown(5, function() {
                      displayNextItem();
                    });
                  }});
        } else {
          // no source available, just display title
          Search.output.append(listItem);
          listItem.slideDown(5, function() {
            displayNextItem();
          });
        }
      }
      // search finished, update title and status message
      else {
        Search.stopPulse();
        Search.title.text(_('Search Results'));
        if (!resultCount)
          Search.status.text(_('Your search did not match any documents. Please make sure that all words are spelled correctly and that you\'ve selected enough categories.'));
        else
            Search.status.text(_('Search finished, found %s page(s) matching the search query.').replace('%s', resultCount));
        Search.status.fadeIn(500);
      }
    }
    displayNextItem();
  },

  /**
   * search for object names
   */
  performObjectSearch : function(object, otherterms) {
    var filenames = this._index.filenames;
    var docnames = this._index.docnames;
    var objects = this._index.objects;
    var objnames = this._index.objnames;
    var titles = this._index.titles;

    var i;
    var results = [];

    for (var prefix in objects) {
      for (var name in objects[prefix]) {
        var fullname = (prefix ? prefix + '.' : '') + name;
        var fullnameLower = fullname.toLowerCase()
        if (fullnameLower.indexOf(object) > -1) {
          var score = 0;
          var parts = fullnameLower.split('.');
          // check for different match types: exact matches of full name or
          // "last name" (i.e. last dotted part)
          if (fullnameLower == object || parts[parts.length - 1] == object) {
            score += Scorer.objNameMatch;
          // matches in last name
          } else if (parts[parts.length - 1].indexOf(object) > -1) {
            score += Scorer.objPartialMatch;
          }
          var match = objects[prefix][name];
          var objname = objnames[match[1]][2];
          var title = titles[match[0]];
          // If more than one term searched for, we require other words to be
          // found in the name/title/description
          if (otherterms.length > 0) {
            var haystack = (prefix + ' ' + name + ' ' +
                            objname + ' ' + title).toLowerCase();
            var allfound = true;
            for (i = 0; i < otherterms.length; i++) {
              if (haystack.indexOf(otherterms[i]) == -1) {
                allfound = false;
                break;
              }
            }
            if (!allfound) {
              continue;
            }
          }
          var descr = objname + _(', in ') + title;

          var anchor = match[3];
          if (anchor === '')
            anchor = fullname;
          else if (anchor == '-')
            anchor = objnames[match[1]][1] + '-' + fullname;
          // add custom score for some objects according to scorer
          if (Scorer.objPrio.hasOwnProperty(match[2])) {
            score += Scorer.objPrio[match[2]];
          } else {
            score += Scorer.objPrioDefault;
          }
          results.push([docnames[match[0]], fullname, '#'+anchor, descr, score, filenames[match[0]]]);
        }
      }
    }

    return results;
  },

  /**
   * search for full-text terms in the index
   */
  performTermsSearch : function(searchterms, excluded, terms, titleterms) {
    var docnames = this._index.docnames;
    var filenames = this._index.filenames;
    var titles = this._index.titles;

    var i, j, file;
    var fileMap = {};
    var scoreMap = {};
    var results = [];

    // perform the search on the required terms
    for (i = 0; i < searchterms.length; i++) {
      var word = searchterms[i];
      var files = [];
      var _o = [
        {files: terms[word], score: Scorer.term},
        {files: titleterms[word], score: Scorer.title}
      ];
      // add support for partial matches
      if (word.length > 2) {
        for (var w in terms) {
          if (w.match(word) && !terms[word]) {
            _o.push({files: terms[w], score: Scorer.partialTerm})
          }
        }
        for (var w in titleterms) {
          if (w.match(word) && !titleterms[word]) {
              _o.push({files: titleterms[w], score: Scorer.partialTitle})
          }
        }
      }

      // no match but word was a required one
      if ($u.every(_o, function(o){return o.files === undefined;})) {
        break;
      }
      // found search word in contents
      $u.each(_o, function(o) {
        var _files = o.files;
        if (_files === undefined)
          return

        if (_files.length === undefined)
          _files = [_files];
        files = files.concat(_files);

        // set score for the word in each file to Scorer.term
        for (j = 0; j < _files.length; j++) {
          file = _files[j];
          if (!(file in scoreMap))
            scoreMap[file] = {};
          scoreMap[file][word] = o.score;
        }
      });

      // create the mapping
      for (j = 0; j < files.length; j++) {
        file = files[j];
        if (file in fileMap && fileMap[file].indexOf(word) === -1)
          fileMap[file].push(word);
        else
          fileMap[file] = [word];
      }
    }

    // now check if the files don't contain excluded terms
    for (file in fileMap) {
      var valid = true;

      // check if all requirements are matched
      var filteredTermCount = // as search terms with length < 3 are discarded: ignore
        searchterms.filter(function(term){return term.length > 2}).length
      if (
        fileMap[file].length != searchterms.length &&
        fileMap[file].length != filteredTermCount
      ) continue;

      // ensure that none of the excluded terms is in the search result
      for (i = 0; i < excluded.length; i++) {
        if (terms[excluded[i]] == file ||
            titleterms[excluded[i]] == file ||
            $u.contains(terms[excluded[i]] || [], file) ||
            $u.contains(titleterms[excluded[i]] || [], file)) {
          valid = false;
          break;
        }
      }

      // if we have still a valid result we can add it to the result list
      if (valid) {
        // select one (max) score for the file.
        // for better ranking, we should calculate ranking by using words statistics like basic tf-idf...
        var score = $u.max($u.map(fileMap[file], function(w){return scoreMap[file][w]}));
        results.push([docnames[file], titles[file], '', null, score, filenames[file]]);
      }
    }
    return results;
  },

  /**
   * helper function to return a node containing the
   * search summary for a given text. keywords is a list
   * of stemmed words, hlwords is the list of normal, unstemmed
   * words. the first one is used to find the occurrence, the
   * latter for highlighting it.
   */
  makeSearchSummary : function(htmlText, keywords, hlwords) {
    var text = Search.htmlToText(htmlText);
    var textLower = text.toLowerCase();
    var start = 0;
    $.each(keywords, function() {
      var i = textLower.indexOf(this.toLowerCase());
      if (i > -1)
        start = i;
    });
    start = Math.max(start - 120, 0);
    var excerpt = ((start > 0) ? '...' : '') +
      $.trim(text.substr(start, 240)) +
      ((start + 240 - text.length) ? '...' : '');
    var rv = $('<div class="context"></div>').text(excerpt);
    $.each(hlwords, function() {
      rv = rv.highlightText(this, 'highlighted');
    });
    return rv;
  }
};

$(document).ready(function() {
  Search.init();
});
```
