## Supplementary material for "scHiCTools: a computational toolbox for analyzing single-cell Hi-C data": S4: genindex.html

  


Index — scHiCTools 0.0.3 documentation


scHiCTools

scHiCTools

- Docs »
- Index

---

### Index

**\_**
| **C**
| **F**
| **G**
| **H**
| **I**
| **K**
| **L**
| **M**
| **N**
| **P**
| **S**
| **T**
| **V**

## \_

- \_\_init\_\_() (scHiCTools.load.ContactMaps.scHiCs method)

## C

|  |  |
| --- | --- |
| - cal\_strata() (scHiCTools.load.ContactMaps.scHiCs method) | - clustering() (scHiCTools.load.ContactMaps.scHiCs method) |

## F

- file\_line\_generator() (in module scHiCTools.load.load\_hic\_file)

## G

- get\_chromosome\_lengths() (in module scHiCTools.load.load\_hic\_file)

## H

- HAC() (in module scHiCTools.analysis.clustering)

## I

- interactive\_scatter() (in module scHiCTools.analysis.visualization)

## K

|  |  |
| --- | --- |
| - kl\_divergence() (in module scHiCTools.embedding.embedding) | - kmeans() (in module scHiCTools.analysis.clustering) |

## L

|  |  |
| --- | --- |
| - learn\_embedding() (scHiCTools.load.ContactMaps.scHiCs method) | - load\_HiC() (in module scHiCTools.load.load\_hic\_file) |

## M

- matrix\_operation() (in module scHiCTools.load.processing\_utils)
- MDS() (in module scHiCTools.embedding.embedding)
- module
  - scHiCTools
  - scHiCTools.analysis
  - scHiCTools.analysis.clustering
  - scHiCTools.analysis.visualization
  - scHiCTools.embedding
  - scHiCTools.embedding.embedding
  - scHiCTools.embedding.reproducibility
  - scHiCTools.load
  - scHiCTools.load.ContactMaps
  - scHiCTools.load.load\_hic\_file
  - scHiCTools.load.processing\_utils

## N

- nMDS() (in module scHiCTools.embedding.embedding)

## P

|  |  |
| --- | --- |
| - pairwise\_distances() (in module scHiCTools.embedding.reproducibility) - PCA() (in module scHiCTools.embedding.embedding) | - PHATE() (in module scHiCTools.embedding.embedding) - plot\_contacts() (scHiCTools.load.ContactMaps.scHiCs method) - processing() (scHiCTools.load.ContactMaps.scHiCs method) |

## S

|  |  |
| --- | --- |
| - scatter() (in module scHiCTools.analysis.visualization) - scHiCluster() (scHiCTools.load.ContactMaps.scHiCs method) - scHiCs (class in scHiCTools.load.ContactMaps) - scHiCTools   - module - scHiCTools.analysis   - module - scHiCTools.analysis.clustering   - module - scHiCTools.analysis.visualization   - module - scHiCTools.embedding   - module - scHiCTools.embedding.embedding   - module | - scHiCTools.embedding.reproducibility   - module - scHiCTools.load   - module - scHiCTools.load.ContactMaps   - module - scHiCTools.load.load\_hic\_file   - module - scHiCTools.load.processing\_utils   - module - select\_cells() (scHiCTools.load.ContactMaps.scHiCs method) - spectral\_clustering() (in module scHiCTools.analysis.clustering) - SpectralEmbedding() (in module scHiCTools.embedding.embedding) |

## T

- tSNE() (in module scHiCTools.embedding.embedding)

## V

- VNE() (in module scHiCTools.embedding.embedding)

---

© Copyright 2020, Xinjun Li

Built with Sphinx using a theme provided by Read the Docs.
