## Supplementary material for "scHiCTools: a computational toolbox for analyzing single-cell Hi-C data": S4: index.html

  


Welcome to scHiCTools’s documentation! — scHiCTools 0.0.3 documentation


scHiCTools

- Welcome to scHiCTools’s documentation!
- Indices and tables
- Introduction
- Setup procedure

scHiCTools

- Docs »
- Welcome to scHiCTools’s documentation!
- View page source

---

### Welcome to scHiCTools’s documentation!¶

### Indices and tables¶

- Index
- Module Index
- Search Page

### Introduction¶

scHiCTools is a computational toolbox for analyzing single cell Hi-C (high-throughput sequencing for 3C) data which includes functions for:

1. Load single-cell HiC datasets
2. Smoothing the contact maps with linear convolution, random walk or network enhancing
3. Calculating embeddings for single cell HiC datasets efficiently with reproducibility measures include InnerProduct, HiCRep and Selfish

### Setup procedure¶

---

© Copyright 2020, Xinjun Li

Built with Sphinx using a theme provided by Read the Docs.
