## Supplementary material for "scHiCTools: a computational toolbox for analyzing single-cell Hi-C data": S4: modules.html

  


scHiCTools — scHiCTools 0.0.3 documentation


scHiCTools

- scHiCTools

scHiCTools

- Docs »
- scHiCTools
- View page source

---

### scHiCTools¶

- scHiCTools package
  - Subpackages
    - scHiCTools.analysis package
      - Submodules
      - scHiCTools.analysis.clustering module
      - scHiCTools.analysis.visualization module
      - Module contents
    - scHiCTools.embedding package
      - Submodules
      - scHiCTools.embedding.embedding module
      - scHiCTools.embedding.reproducibility module
      - Module contents
    - scHiCTools.load package
      - Submodules
      - scHiCTools.load.ContactMaps module
      - scHiCTools.load.load\_hic\_file module
      - scHiCTools.load.processing\_utils module
      - Module contents
  - Module contents

---

© Copyright 2020, Xinjun Li

Built with Sphinx using a theme provided by Read the Docs.
