## Supplementary material for "scHiCTools: a computational toolbox for analyzing single-cell Hi-C data": S4: py-modindex.html

  

Python Module Index — scHiCTools 0.0.3 documentation

scHiCTools

scHiCTools

- Docs »
- Python Module Index

---

### Python Module Index

**s**

|  |  |
| --- | --- |
|  | **s** |
|  | `scHiCTools` |
|  | `scHiCTools.analysis` |
|  | `scHiCTools.analysis.clustering` |
|  | `scHiCTools.analysis.visualization` |
|  | `scHiCTools.embedding` |
|  | `scHiCTools.embedding.embedding` |
|  | `scHiCTools.embedding.reproducibility` |
|  | `scHiCTools.load` |
|  | `scHiCTools.load.ContactMaps` |
|  | `scHiCTools.load.load_hic_file` |
|  | `scHiCTools.load.processing_utils` |

---

© Copyright 2020, Xinjun Li

Built with Sphinx using a theme provided by Read the Docs.
