## Supplementary material for "scHiCTools: a computational toolbox for analyzing single-cell Hi-C data": S4: scHiCTools.analysis.html

  


scHiCTools.analysis package — scHiCTools 0.0.3 documentation


scHiCTools

- scHiCTools.analysis package
  - Submodules
  - scHiCTools.analysis.clustering module
  - scHiCTools.analysis.visualization module
  - Module contents

scHiCTools

- Docs »
- scHiCTools.analysis package
- View page source

---

### scHiCTools.analysis package¶

#### Submodules¶

#### scHiCTools.analysis.clustering module¶

Clustering component of scHiCTools

Author: Xinjun Li

This module define functions to clustering single cell data.

`scHiCTools.analysis.clustering.``HAC`(*data*, *data\_type='points'*, *n\_clusters=4*, *method='centroid'*)¶
:   This function is a impliment of hierarchical afflomerative clustering.
    Reference:

    > Christopher D. Manning, Prabhakar Raghavan and Hinrich Schütze,
    > “Introduction to Information Retrieval” chapter 17

    Parameters
    :   - **data** (*numpy.ndarray*) – Distance matrix or coordinate of the points.
        - **data\_type** (*str**,* *optional*) – Format of argument data, specify the type of data input.
          Must be in either ‘points’ or ‘distance\_matrix’.
          The default is ‘points’.
        - **n\_clusters** (*int**,* *optional*) – Number of clusters.
          The default is 4.
        - **method** (*str**,* *optional*) – Specify the hierarchical clustering method used.
          Now support ‘single-link’, ‘complete-link’, ‘centroid’,’group-average’.
          The default is ‘centroid’.

    Returns
    :   **label** (*numpy.ndarray*) – A list of the label of every point.

`scHiCTools.analysis.clustering.``kmeans`(*data*, *k=4*, *weights=None*, *iteration=1000*, *\*\*kwargs*)¶
:   k-means algorithm, with k-means++ to initialize the start points.

    Parameters
    :   - **data** (*numpy.ndarray*) – Coordiante of points.
        - **k** (*int**,* *optional*) – Number of clusters.
          The default is 4.
        - **weights** (*list**,* *optional*) – List of the weight of every point,
          should either be ‘None’ or have the length same as the number of rows in data.
          The default is None.
        - **iteration** (*int**,* *optional*) – Number of iterations in k-means algorithm.
          The default is 1000.

    Returns
    :   **label** (*numpy.ndarray*) – A list of the label of every point.

`scHiCTools.analysis.clustering.``spectral_clustering`(*data*, *data\_type='points'*, *n\_clusters=4*, *normalize=True*, *\*\*kwargs*)¶
:   This function is a impliment of unnormalized spectral clustering.
    Reference: “A Tutorial on Spectral Clustering” by Ulrike von Luxburg.

    Parameters
    :   - **data** (*numpy.ndarray*) – Distance matrix or coordinate of the points.
        - **data\_type** (*str**,* *optional*) – Format of argument data, specify the type of data input.
          Must be in either ‘points’ or ‘distance\_matrix’.
          The default is ‘points’.
        - **n\_clusters** (*int**,* *optional*) – Number of clusters.
          The default is 4.
        - **normalize** (*bool**,* *optional*) – Whether to use unnormalized or normalized spectral clustering.
          The default is True.
        - **\*\*kwargs** – Arguments pass to kmeans.

    Returns
    :   **label** (*numpy.ndarray*) – A list of the label of every point.

#### scHiCTools.analysis.visualization module¶

Visualization component of scHiCTools

Author: Xinjun Li

This module define a function to plot scatter plot of embedding points of single cell data.

`scHiCTools.analysis.visualization.``interactive_scatter`(*schic*, *data*, *out\_file*, *dimension='2D'*, *point\_size=3*, *label=None*, *title=None*, *alpha=1*, *aes\_label=None*)¶
:   This function is to generate an interactive scatter plot of embedded single cell data.

    Parameters
    :   - **schic** (*scHiCs*) – A scHiCs object.
        - **data** (*numpy.array*) – A numpy array which has 2 or 3 columns, every row represent a point.
        - **out\_file** (*str*) – Output file path.
        - **dimension** (*str**,* *optional*) – Specifiy the dimension of the plot, either “2D” or “3D”.
          The default is “2D”.
        - **point\_size** (*float**,* *optional*) – Set the size of the points in scatter plot.
          The default is 3.
        - **label** (*list* *or* *None**,* *optional*) – Specifiy the label of each point. The default is None.
        - **title** (*str**,* *optional*) – Title of the plot. The default is None.
        - **alpha** (*float**,* *optional*) – The alpha blending value. The default is 1.
        - **aes\_label** (*list**,* *optional*) – Set the label of every axis. The default is None.

`scHiCTools.analysis.visualization.``scatter`(*data*, *dimension='2D'*, *point\_size=3*, *sty='default'*, *label=None*, *title=None*, *alpha=None*, *aes\_label=None*, *\*\*kwargs*)¶
:   This function is to plot scatter plot of embedding points
    :   of single cell data.

    Parameters
    :   - **data** (*numpy.array*) – A numpy array which has 2 or 3 columns, every row represent a point.
        - **dimension** (*str**,* *optional*) – Specifiy the dimension of the plot, either “2D” or “3D”.
          The default is “2D”.
        - **point\_size** (*float**,* *optional*) – Set the size of the points in scatter plot.
          The default is 3.
        - **sty** (*str**,* *optional*) – Styles of Matplotlib. The default is ‘default’.
        - **label** (*list* *or* *None**,* *optional*) – Specifiy the label of each point. The default is None.
        - **title** (*str**,* *optional*) – Title of the plot. The default is None.
        - **alpha** (*float**,* *optional*) – The alpha blending value. The default is None.
        - **aes\_label** (*list**,* *optional*) – Set the label of every axis. The default is None.
        - **\*\*kwargs** – Other arguments passed to the matplotlib.pyplot.legend,
          controlling the plot’s legend.

    Returns
    :   *Scatter plot of either 2D or 3D.*

#### Module contents¶

---

© Copyright 2020, Xinjun Li

Built with Sphinx using a theme provided by Read the Docs.
