## Supplementary material for "scHiCTools: a computational toolbox for analyzing single-cell Hi-C data": S4: scHiCTools.embedding.html

  


scHiCTools.embedding package — scHiCTools 0.0.3 documentation


scHiCTools

- scHiCTools.embedding package
  - Submodules
  - scHiCTools.embedding.embedding module
  - scHiCTools.embedding.reproducibility module
  - Module contents

scHiCTools

- Docs »
- scHiCTools.embedding package
- View page source

---

### scHiCTools.embedding package¶

#### Submodules¶

#### scHiCTools.embedding.embedding module¶

Embedding component of scHiCTools

Author: Xinjun Li

This script define functions to embedding single cell data to a lower-dimensional space.

`scHiCTools.embedding.embedding.``MDS`(*mat*, *n=2*)¶
:   Multidimensional scaling, MDS.

    Parameters
    :   - **mat** (*numpy.ndarray*) – Distance matrix of the data points.
        - **n** (*int**,* *optional*) – The dimension of the projected points.
          The default is 2.

    Returns
    :   **co** (*numpy.ndarray*) – Coordinates of the projected points.

`scHiCTools.embedding.embedding.``PCA`(*X*, *dim=2*)¶
:   Principal components analysis，PCA.

    Parameters
    :   - **X** (*numpy.ndarray*) – Coordinates of input data points.
        - **dim** (*int**,* *optional*) – The dimension of the projected points.
          The default is 2.

    Returns
    :   **Y** (*numpy.ndarray*) – Coordinates of the projected points.

`scHiCTools.embedding.embedding.``PHATE`(*mat*, *dim=2*, *k=5*, *a=1*, *gamma=1*, *t\_max=100*, *momentum=0.1*, *iteration=1000*)¶
:   Potential of Heat-diffusion for Affinity-based Trajectory Embedding,
    PHATE.

    Parameters
    :   - **mat** (*numpy.ndarray*) – Distance matrix.
        - **dim** (*int**,* *optional*) – Desired embedding dimension.
          The default is 2.
        - **k** (*int**,* *optional*) – Neighborhood size.
          The default is 5.
        - **a** (*float**,* *optional*) – Locality scale.
          The default is 1.
        - **gamma** (*float**,* *optional**,* *must in* *[**-1**,* *1**]*) – Informational distance constant between -1 and 1.
          The default is 1.
        - **t\_max** (*int**,* *optional*) – Maximum time scale for diffusion.
          The default is 100.
        - **momentum** (*float**,* *optional*) – Momentum of gradient descent algorithm.
          The default is .1.
        - **iteration** (*TYPE**,* *optional*) – Number of iteration in gradient descent.
          The default is 1000.

    Returns
    :   **Y** (*numpy.ndarray*) – The PHATE embedding matrix.

`scHiCTools.embedding.embedding.``SpectralEmbedding`(*graph\_matrix*, *dim*)¶
:   Spectral embedding.

    Parameters
    :   - **graph\_matrix** (*numpy.ndarray*) – Adjecent matrix of the graph of points.
        - **dim** (*int*) – Dimension of embedding space.

    Returns
    :   **Y** (*numpy.ndarray*) – Embedding points.

`scHiCTools.embedding.embedding.``VNE`(*P*, *t*)¶
:   Von Neumann Entropy.

    Parameters
    :   - **P** (*numpy.ndarray*) – The density matrix.
        - **t** (*int*) – Number of time stages to calculate VNE.

    Returns
    :   **Ht** (*float*) – Von Neumann Entropy.

`scHiCTools.embedding.embedding.``kl_divergence`(*params*, *P*, *degrees\_of\_freedom*, *n\_samples*, *n\_components*)¶
:   t-SNE objective function: gradient of the KL divergence
    of p\_ijs and q\_ijs and the absolute error.

    Parameters
    :   - **params** (*numpy.ndarray**,* *shape* *(**n\_params**,**)*) – Unraveled embedding.
        - **P** (*numpy.ndarray**,* *shape* *(**n\_samples \** *(**n\_samples-1**)* */ 2**,**)*) – Condensed joint probability matrix.
        - **degrees\_of\_freedom** (*int*) – Degrees of freedom of the Student’s-t distribution.
        - **n\_samples** (*int*) – Number of samples.
        - **n\_components** (*int*) – Dimension of the embedded space.

    Returns
    :   - **kl\_divergence** (*float*) – Kullback-Leibler divergence of p\_ij and q\_ij.
        - **grad** (*numpy.ndarray, shape (n\_params,)*) – Unraveled gradient of the Kullback-Leibler divergence with respect to the embedding.

`scHiCTools.embedding.embedding.``nMDS`(*dist\_mat*, *init*, *momentum=0.1*, *iteration=1000*)¶
:   Non-metric multi-dimensional scaling, nMDS.

    Parameters
    :   - **dist\_mat** (*array*) – Distance matrix of the points.
        - **init** (*array*) – Initial embedding.
        - **momentum** (*float*) – Dimension of the embedding space.
          The default is 2.
        - **iteration** (*int*) – Number of iteration of gradient descent.
          The default is 1000.

    Returns
    :   **Y** (*array*) – Embedding points.

`scHiCTools.embedding.embedding.``tSNE`(*mat*, *n\_dim=2*, *perp=30.0*, *n\_iter=1000*, *momentum=0.5*, *rate=200.0*, *tol=1e-05*)¶
:   t-Distributed Stochastic Neighbor Embedding, t-SNE.

    Parameters
    :   - **mat** (*numpy.ndarray*) – Distance matrix of the data points.
        - **n\_dim** (*int**,* *optional*) – The dimension of the projected points.
          The default is 2.
        - **perp** (*float**,* *optional*) – Perplexity.
          The default is 30.0.
        - **n\_iter** (*int**,* *optional*) – Max number of iteration.
          The default is 1000.
        - **momentum** (*float**,* *optional*) – Momentum of gredient decendent.
          The default is 0.5.
        - **rate** (*float**,* *optional*) – Gredient decendent rate.
          The default is 200.
        - **tol** (*float**,* *optional*) – The threshold of gradient norm to stop the iteration of grendient decendent.
          The default is 1e-5.

    Returns
    :   **Y** (*numpy.ndarray*) – Coordinates of the projected points.

#### scHiCTools.embedding.reproducibility module¶

`scHiCTools.embedding.reproducibility.``pairwise_distances`(*all\_strata*, *similarity\_method*, *print\_time=False*, *sigma=0.5*, *window\_size=10*)¶
:   Find the pairwise distace using different similarity method,
    and return a distance matrix.

    Parameters
    :   - **all\_strata** (*numpy.ndarray*) – An array contained the strata of a specific chromesome for cells.
        - **similarity\_method** (*str*) – The similarity method used to calculate distance matrix.
          Now support ‘innerproduct’, ‘hicrep’, ‘selfish’.
        - **print\_time** (*bool**,* *optional*) – Whether to return the run time of similarity.
          The default is False.
        - **sigma** (*float**,* *optional*) – Parameter for method ‘Selfish’.
          Sigma for the Gaussian kernel used in Selfish.
          The default is .5.
        - **window\_size** (*int**,* *optional*) – Parameter for method ‘Selfish’.
          Length of every window used in Selfish.
          The default is 10.

    Returns
    :   *numpy.ndarray* – Distance matrix.

#### Module contents¶

---

© Copyright 2020, Xinjun Li

Built with Sphinx using a theme provided by Read the Docs.
