## Supplementary material for "scHiCTools: a computational toolbox for analyzing single-cell Hi-C data": S4: scHiCTools.load.html

  


scHiCTools.load package — scHiCTools 0.0.3 documentation


scHiCTools

- scHiCTools.load package
  - Submodules
  - scHiCTools.load.ContactMaps module
  - scHiCTools.load.load\_hic\_file module
  - scHiCTools.load.processing\_utils module
  - Module contents

scHiCTools

- Docs »
- scHiCTools.load package
- View page source

---

### scHiCTools.load package¶

#### Submodules¶

#### scHiCTools.load.ContactMaps module¶

*class* `scHiCTools.load.ContactMaps.``scHiCs`(*list\_of\_files*, *reference\_genome*, *resolution*, *adjust\_resolution=True*, *sparse=False*, *chromosomes='all'*, *format='customized'*, *keep\_n\_strata=10*, *store\_full\_map=False*, *operations=None*, *header=0*, *customized\_format=None*, *map\_filter=0.0*, *gzip=False*, *\*\*kwargs*)¶
:   Bases: `object`

    `__init__`(*list\_of\_files*, *reference\_genome*, *resolution*, *adjust\_resolution=True*, *sparse=False*, *chromosomes='all'*, *format='customized'*, *keep\_n\_strata=10*, *store\_full\_map=False*, *operations=None*, *header=0*, *customized\_format=None*, *map\_filter=0.0*, *gzip=False*, *\*\*kwargs*)¶
    :   Parameters
        :   - **list\_of\_files** (*list*) – List of HiC file paths.
            - **reference\_genome** (*str* *or* *dict*) – Now supporting ‘mm9’, ‘mm10’, ‘hg19’, ‘hg38’,
              if using other references,you can simply provide the chromosome name
              and corresponding size (bp) with a dictionary in Python.
              e.g. {‘chr1’: 150000000, ‘chr2’: 130000000, ‘chr3’: 200000000}
            - **resolution** (*int*) – The resolution to separate genome into bins. If using .hic file format,
              the given resolution must match with the resolutions in .hic file.
            - **adjust\_resolution** (*bool**,* *optional*) – Whether to adjust resolution for input file.
              Sometimes the input file is already in the proper resolution
              (e.g. position 3000000 has already been changed to 6 in 500kb resolution),
              then you can set adjust\_resolution=False. The default is True.
            - **sparse** (*bool**,* *optional*) – Whether to use sparse matrix to store (only effective when max\_distance=None). The default is False.
            - **chromosomes** (*list* *or* *str**,* *optional*) – Chromosomes to use,
              eg. [‘chr1’, ‘chr2’], or just ‘except Y’, ‘except XY’,’all’,
              which means chr 1-19 + XY for mouse and chr 1-22 + XY for human.
              The default is ‘all’.
            - **format** (*str**,* *optional*) – HiC files’ format.
              e.g., ‘.hic’, ‘customized’, ‘.cool’. The default is ‘customized’.
            - **keep\_n\_strata** (*int**,* *optional*) – Only consider contacts within this genomic distance.
              If None, it will store full matrices in numpy matrix or
              scipy sparse format, which will use too much memory sometimes.
              The default is 10.
            - **store\_full\_map** (*bool**,* *optional*) – Whether store contact maps. The default is False.
            - **operations** (*list**,* *optional*) – The methods use for pre-processing or smoothing the maps given in a list.
              The operations will happen in the given order.
              Operations: ‘convolution’, ‘random\_walk’, ‘network\_enhancing’.
              For pre-processing and smoothing operations, sometimes you need additional arguments.
              You can check docstrings for pre-processing and smoothing for more information.
              The default is None.
            - **header** (*int**,* *optional*) – The number of header line(s).
              If header=0, HiC files do not have header.
              The default is 0.
            - **customized\_format** (*int* *or* *list**,* *optional*) – Format for each line. The default is None.
            - **map\_filter** (*float**,* *optional*) – The threshold to filter some reads by map quality.
              The default is 0..
            - **gzip** (*bool**,* *optional*) – If the HiC files are zip files.
              If True, the HiC files are zip files.
              The default is False.
            - **\*\*kwargs** – Other arguments specify smoothing methods passed to function.
              See scHiCTools.load.processing\_utils.matrix\_operation function.

        Returns
        :   *None.*

    `cal_strata`(*n\_strata*)¶
    :   Alter the number of strata kept in a scHiCs object.

        Parameters
        :   **n\_strata** (*int*) – Number of strata to keep.

        Returns
        :   *dict* – Strata of cells.

    `clustering`(*n\_clusters*, *clustering\_method*, *similarity\_method*, *aggregation='median'*, *n\_strata=None*, *print\_time=False*, *\*\*kwargs*)¶
    :   Parameters
        :   - **n\_clusters** (*int*) – Number of clusters.
            - **clustering\_method** (*str*) – Clustering method in ‘kmeans’, ‘spectral\_clustering’ or ‘HAC’(hierarchical agglomerative clustering).
            - **similarity\_method** (*str*) – Reproducibility measure.
              Value in ‘InnerProduct’, ‘HiCRep’ or ‘Selfish’.
            - **aggregation** (*str**,* *optional*) – Method to aggregate different chromosomes.
              Value is either ‘mean’ or ‘median’.
              The default is ‘median’.
            - **n\_strata** (*int* *or* *None**,* *optional*) – Only consider contacts within this genomic distance.
              If it is None, it will use the all strata kept from previous loading process.
              The default is None.
            - **print\_time** (*bool**,* *optional*) – Whether to print the processing time. The default is False.
            - **\*\*kwargs** – Other arguments pass to function scHiCTools.embedding.reproducibility.pairwise\_distances `,
              and the clustering function in `scHiCTools.analysis.clustering.

        Returns
        :   **label** (*numpy.ndarray*) – An array of cell labels clustered.

    `learn_embedding`(*similarity\_method*, *embedding\_method*, *dim=2*, *aggregation='median'*, *n\_strata=None*, *return\_distance=False*, *print\_time=False*, *\*\*kwargs*)¶
    :   Function to find a low-dimensional embedding for cells.

        Parameters
        :   - **similarity\_method** (*str*) – The method used to calculate similarity matrix.
              Now support ‘inner\_product’, ‘HiCRep’ and ‘Selfish’.
            - **embedding\_method** (*str*) – The method used to project cells into lower-dimensional space.
              Now support ‘MDS’, ‘tSNE’, ‘phate’, ‘spectral\_embedding’.
            - **dim** (*int**,* *optional*) – Dimension of the embedding space.
              The default is 2.
            - **aggregation** (*str**,* *optional*) – Method to find the distance matrix based on distance matrices of chromesomes.
              Must be ‘mean’ or ‘median’.
              The default is ‘median’.
            - **n\_strata** (*int**,* *optional*) – Number of strata used in calculation.
              The default is None.
            - **return\_distance** (*bool**,* *optional*) – Whether to return the distance matrix of cells.
              If True, return (embeddings, distance\_matrix);
              if False, only return embeddings.
              The default is False.
            - **print\_time** (*bool**,* *optional*) – Whether to print process time. The default is False.
            - **\*\*kwargs** – Including two arguments for Selfish
              (see funciton pairwise\_distances): n\_windows: number of Selfish windows sigma: sigma in the Gaussian-like kernel and some arguments specify different embedding method
              (see functions in scHiCTools.embedding.embedding).

        Returns
        :   - **embeddings** (*numpy.ndarray*) – The embedding of cells in lower-dimensional space.
            - **final\_distance** (*numpy.ndarray, optional*) – The pairwise distance calculated.

    `plot_contacts`(*hist=True*, *percent=True*, *size=1.0*, *bins=10*, *color='#1f77b4'*)¶
    :   Generate two plots:
        Histogram of contacts and
        scatter plot of short-range contacts v.s. contacts at the mitotic band.

        Parameters
        :   - **hist** (*bool**,* *optional*) – Whether to plot Histogram of contacts.
              If True, plot Histogram of contacts.
              The default is True.
            - **percent** (*int**,* *optional*) – Whether to plot scatter plot of short-range contacts v.s. contacts at the mitotic band.
              If True, plot scatter plot of short-range contacts v.s. contacts at the mitotic band.
              The default is True.
            - **size** (*float**,* *optional*) – The point size of scatter plot.
              The default is 1.0.
            - **bins** (*int**,* *optional*) – Number of bins in histogram.
              The default is 10.
            - **color** (*str**,* *optional*) – The color of the plot.
              The default is ‘#1f77b4’.

        Returns
        :   *None.*

    `processing`(*operations*, *\*\*kwargs*)¶
    :   Apply a smoothing method to contact maps.
        Requre the scHiCs object to store the full map of contacts maps.

        Parameters
        :   - **operations** (*str*) – The methods use for smoothing the maps.
              Avaliable operations: ‘convolution’, ‘random\_walk’, ‘network\_enhancing’.
            - **\*\*kwargs** – Other arguments specify smoothing methods passed to function.
              See function scHiCTools.load.processing\_utils.matrix\_operation.

        Returns
        :   *None.*

    `scHiCluster`(*dim=2*, *n\_clusters=4*, *cutoff=0.8*, *n\_PCs=10*, *\*\*kwargs*)¶
    :   Embedding and clustering single cells using HiCluster.
        Reference:

        > Zhou J, Ma J, Chen Y, Cheng C, Bao B, Peng J, et al.
        > Robust single-cell Hi-C clustering by convolution- and random-walk–based imputation.
        > PNAS. 2019 Jul 9;116(28):14011–8.

        Parameters
        :   - **dim** (*int**,* *optional*) – Number of dimension of embedding. The default is 2.
            - **n\_clusters** (*int**,* *optional*) – Number of clusters. The default is 4.
            - **cutoff** (*float**,* *optional*) – The cutoff proportion to convert the real contact
              matrix into binary matrix. The default is 0.8.
            - **n\_PCs** (*int**,* *optional*) – Number of principal components. The default is 10.
            - **\*\*kwargs** – Other arguments passed to kmeans.
              See scHiCTools.analysis.clustering.kmeans function.

        Returns
        :   - **embeddings** (*numpy.ndarray*) – The embedding of cells using HiCluster.
            - **label** (*numpy.ndarray*) – An array of cell labels clustered by HiCluster.

    `select_cells`(*min\_n\_contacts=0*, *max\_short\_range\_contact=1*)¶
    :   Select qualify cells based on minimum number of contacts and
        maxium percent of short range contact.

        Parameters
        :   - **min\_n\_contacts** (*int**,* *optional*) – The threshold of minimum number of contacts in each cell.
              The default is 0.
            - **max\_short\_range\_contact** (*float**,* *optional*) – The threshold of maximum proportion of short range contact in every cell.
              The default is 1.

        Returns
        :   *list* – Selected files.

#### scHiCTools.load.load\_hic\_file module¶

This file is for loading a single HiC contact map from different type of file.

`scHiCTools.load.load_hic_file.``file_line_generator`(*file*, *format=None*, *chrom=None*, *header=0*, *resolution=1*, *resolution\_adjust=True*, *mapping\_filter=0.0*, *gzip=False*)¶
:   For formats other than .hic and .mcool

    Parameters
    :   - **file** (*str*) – File path.
        - **format** (*int* *or* *list*) – Format
          1) [chr1 pos1 chr2 pos2 mapq1 mapq2];
          2) [chr1 pos1 chr2 pos2 score];
          3) [chr1 pos1 chr2 pos2].
        - **chrom** (*str*) – Chromosome to extract.
        - **header** (*None* *or* *int*) – Whether to skip header (1 line).
        - **mapping\_filter** (*float*) – The threshold to filter some reads by map quality.

    Returns
    :   No return value.
        Yield a line each time in the format of (position\_1, position\_2, contact\_reads).

`scHiCTools.load.load_hic_file.``get_chromosome_lengths`(*ref*, *chromosomes*, *res=1*)¶
:   Get lengths for all chromosomes in a given resolution according to the reference genome.

    Parameters
    :   - **ref** (*str* *or* *dict*) – name of reference genome, eg.’mm10’, ‘hg38’
        - **res** (*int*) – resolution
        - **chromosomes** (*str*) – ‘All’, ‘except X’, ‘except Y’ or a list of chromosomes like [‘chr1’, ‘chr2’]

    Returns
    :   *chromosomes (set)* – lengths (dict): eg. {‘chr1’: 395, ‘chr2’: 390, …}

`scHiCTools.load.load_hic_file.``load_HiC`(*file*, *genome\_length*, *format=None*, *custom\_format=None*, *header=0*, *chromosome=None*, *resolution=10000*, *resolution\_adjust=True*, *map\_filter=0.0*, *sparse=False*, *gzip=False*, *keep\_n\_strata=False*, *operations=None*, *\*\*kwargs*)¶
:   Load HiC contact map into a matrix
    :param file: File path.
    :type file: str
    :param genome\_length: The length of each genome.
    :type genome\_length: dict
    :param format: Now support .txt, .hic, and .mcool file.
    :type format: str
    :param custom\_format: If the format is not in our provided list.
    :type custom\_format: int or list
    :param header: How many header lines to skip.
    :type header: int
    :param chromosome: Specify the chromosome.
    :type chromosome: str
    :param resolution: Resolution.
    :type resolution: int
    :param resolution\_adjust: In some situations, the input file is already pre-processed, and we don’t need to adjust resolution again.
    :type resolution\_adjust: bool
    :param map\_filter: The threshold to filter some reads by map quality
    :type map\_filter: float
    :param sparse: Whether store in sparse matrices
    :type sparse: bool
    :param gzip: Whether the file is zipped.
    :type gzip: bool
    :param keep\_n\_strata: Number of strata to keep.
    :type keep\_n\_strata: int or None

    Returns
    :   *Numpy.array* – loaded contact map

#### scHiCTools.load.processing\_utils module¶

`scHiCTools.load.processing_utils.``matrix_operation`(*mat*, *operations*, *\*\*kwargs*)¶
:   Parameters
    :   - **mat** (*numpy.ndarray*) – Matrix to apply smoothing operators.
        - **operations** (*list**[**str**]*) – A list of smoothing operators to apply.
          Now support perators: ‘oe\_norm’, ‘vc\_norm’, ‘vc\_sqrt\_norm’,
          ‘kr\_norm’, ‘convolution’, ‘random\_walk’, ‘network\_enhancing’,
          ‘logarithm’, ‘power’.
        - **\*\*kwargs** –

          Arguments for specific smoothing operators.

          ’kr\_norm’:
          :   ’maximum\_error\_rate’: error rate to stop iteration, default=1e-4.

          ’convolution’:
          :   ’kernel\_shape’: shape of kernel (‘kernel\_shape’ ,’kernel\_shape’), default=3.

          ’random\_walk’:
          :   ’random\_walk\_ratio’: propotion of random walk smooth matrix, default=1.0;
              ‘t’: number of random walk iteration times, default=1.

          ’network\_enhancing’:
          :   ’kNN’: number of nearest neighbors, default=20;
              ‘iterations’: number of iterations, default=1.

          ’logarithm’:
          :   ’epsilon’: numerator of error term, default=1;
              ‘log\_base’: denominator of error’s exponentiation, default=e.

          ’power’:
          :   ’pow’: exponent of exponentiation, default=0.5.

    Returns
    :   **mat** (*numpy.ndarray*) – Matrix after appling smoothing operator.

#### Module contents¶

PyHiC
\_\_author\_\_ == ‘Fan Feng’
=======

For conveniently dealing with single cell HiC data.

---

© Copyright 2020, Xinjun Li

Built with Sphinx using a theme provided by Read the Docs.
