## Supplementary material for "scHiCTools: a computational toolbox for analyzing single-cell Hi-C data": S4: search.html

  


Search — scHiCTools 0.0.3 documentation


scHiCTools

scHiCTools

- Docs »
- Search

---

Please activate JavaScript to enable the search
functionality.

---

© Copyright 2020, Xinjun Li

Built with Sphinx using a theme provided by Read the Docs.
