## Supplementary material for "scHiCTools: a computational toolbox for analyzing single-cell Hi-C data": S4: searchindex.js

```
Search.setIndex({docnames:["index","modules","scHiCTools","scHiCTools.analysis","scHiCTools.embedding","scHiCTools.load"],envversion:{"sphinx.domains.c":2,"sphinx.domains.changeset":1,"sphinx.domains.citation":1,"sphinx.domains.cpp":2,"sphinx.domains.index":1,"sphinx.domains.javascript":2,"sphinx.domains.math":2,"sphinx.domains.python":2,"sphinx.domains.rst":2,"sphinx.domains.std":1,sphinx:56},filenames:["index.rst","modules.rst","scHiCTools.rst","scHiCTools.analysis.rst","scHiCTools.embedding.rst","scHiCTools.load.rst"],objects:{"":{scHiCTools:[2,0,0,"-"]},"scHiCTools.analysis":{clustering:[3,0,0,"-"],visualization:[3,0,0,"-"]},"scHiCTools.analysis.clustering":{HAC:[3,1,1,""],kmeans:[3,1,1,""],spectral_clustering:[3,1,1,""]},"scHiCTools.analysis.visualization":{interactive_scatter:[3,1,1,""],scatter:[3,1,1,""]},"scHiCTools.embedding":{embedding:[4,0,0,"-"],reproducibility:[4,0,0,"-"]},"scHiCTools.embedding.embedding":{MDS:[4,1,1,""],PCA:[4,1,1,""],PHATE:[4,1,1,""],SpectralEmbedding:[4,1,1,""],VNE:[4,1,1,""],kl_divergence:[4,1,1,""],nMDS:[4,1,1,""],tSNE:[4,1,1,""]},"scHiCTools.embedding.reproducibility":{pairwise_distances:[4,1,1,""]},"scHiCTools.load":{ContactMaps:[5,0,0,"-"],load_hic_file:[5,0,0,"-"],processing_utils:[5,0,0,"-"]},"scHiCTools.load.ContactMaps":{scHiCs:[5,2,1,""]},"scHiCTools.load.ContactMaps.scHiCs":{__init__:[5,3,1,""],cal_strata:[5,3,1,""],clustering:[5,3,1,""],learn_embedding:[5,3,1,""],plot_contacts:[5,3,1,""],processing:[5,3,1,""],scHiCluster:[5,3,1,""],select_cells:[5,3,1,""]},"scHiCTools.load.load_hic_file":{file_line_generator:[5,1,1,""],get_chromosome_lengths:[5,1,1,""],load_HiC:[5,1,1,""]},"scHiCTools.load.processing_utils":{matrix_operation:[5,1,1,""]},scHiCTools:{analysis:[3,0,0,"-"],embedding:[4,0,0,"-"],load:[5,0,0,"-"]}},objnames:{"0":["py","module","Python module"],"1":["py","function","Python function"],"2":["py","class","Python class"],"3":["py","method","Python method"]},objtypes:{"0":"py:module","1":"py:function","2":"py:class","3":"py:method"},terms:{"1f77b4":5,"500kb":5,"class":5,"default":[3,4,5],"float":[3,4,5],"function":[0,3,4,5],"int":[3,4,5],"return":[3,4,5],"sch\u00fctze":3,"short":5,"true":[3,5],For:5,MDS:[4,5],The:[3,4,5],__author__:5,__init__:5,absolut:4,accord:5,addit:5,adjec:4,adjust:5,adjust_resolut:5,aes_label:3,affin:4,afflom:3,after:5,again:5,agglom:5,aggreg:5,algorithm:[3,4],all:5,all_strata:4,alpha:3,alreadi:5,alter:5,analysi:[1,2,4,5],analyz:0,appl:5,appli:5,argument:[3,5],arrai:[3,4,5],author:[3,4],avali:5,averag:3,axi:3,band:5,bao:5,base:[4,5],been:5,between:4,bin:5,binari:5,blend:3,bool:[3,4,5],cal_strata:5,calcul:[0,4,5],can:5,cell:[0,3,4,5],centroid:3,chang:5,chapter:3,check:5,chen:5,cheng:5,chr1:5,chr2:5,chr3:5,chr:5,christoph:3,chrom:5,chromesom:[4,5],chromosom:5,cluster:[1,2,5],clustering_method:5,color:5,column:3,complet:3,compon:[3,4,5],comput:0,condens:4,consid:5,constant:4,contact:[0,5],contact_read:5,contactmap:[1,2],contain:4,content:1,control:3,conveni:5,convert:5,convolut:[0,5],cool:5,coordiant:3,coordin:[3,4],correspond:5,custom:5,custom_format:5,customized_format:5,cutoff:5,data:[0,3,4,5],data_typ:3,dataset:0,deal:5,decend:4,defin:[3,4],degre:4,degrees_of_freedom:4,denomin:5,densiti:4,descent:4,descript:[],desir:4,dict:5,dictionari:5,differ:[4,5],diffus:4,dim:[4,5],dimens:[3,4,5],dimension:[4,5],dist_mat:4,distac:4,distanc:[3,4,5],distance_matrix:[3,5],distribut:4,diverg:4,docstr:5,don:5,each:[3,5],effect:5,effici:0,either:[3,5],embed:[0,1,2,3,5],embedding_method:5,enhanc:0,entropi:4,epsilon:5,error:[4,5],everi:[3,4,5],except:5,expon:5,exponenti:5,extract:5,fals:[4,5],fan:5,feng:5,file:[3,5],file_line_gener:5,filter:5,final_dist:5,find:[4,5],font:[],format:[3,5],freedom:4,from:5,full:5,funciton:5,gamma:4,gaussian:[4,5],gener:[3,5],genom:5,genome_length:5,get:5,get_chromosome_length:5,given:5,grad:4,gradient:4,graph:4,graph_matrix:4,gredient:4,grendient:4,group:3,gzip:5,hac:[3,5],happen:5,has:[3,5],have:[3,5],header:5,heat:4,hg19:5,hg38:5,hic:[0,5],hiclust:5,hicrep:[0,4,5],hierarch:[3,5],high:0,hinrich:3,hist:5,histogram:5,how:5,human:5,impliment:3,imput:5,includ:[0,5],index:0,inform:[3,4,5],init:4,initi:[3,4],inner_product:5,innerproduct:[0,4,5],input:[3,4,5],interact:3,interactive_scatt:3,introduct:3,iter:[3,4,5],joint:4,jul:5,just:5,keep:5,keep_n_strata:5,kept:5,kernel:[4,5],kernel_shap:5,kl_diverg:4,kmean:[3,5],knn:5,kr_norm:5,kullback:4,kwarg:[3,5],label:[3,5],learn_embed:5,legend:3,leibler:4,length:[3,4,5],like:5,line:5,linear:0,link:3,list:[3,5],list_of_fil:5,load:[0,1,2],load_hic:5,load_hic_fil:[1,2],local:4,log_bas:5,logarithm:5,low:5,lower:[4,5],luxburg:3,man:3,mani:5,map:[0,5],map_filt:5,mapping_filt:5,mapq1:5,mapq2:5,mat:[4,5],match:5,matplotlib:3,matric:5,matrix:[3,4,5],matrix_oper:5,max:4,max_dist:5,max_short_range_contact:5,maximum:[4,5],maximum_error_r:5,maxium:5,mcool:5,mean:[3,5],measur:[0,5],median:5,memori:5,method:[3,4,5],metric:4,min_n_contact:5,minimum:5,mitot:5,mm10:5,mm9:5,modul:[0,1],momentum:4,more:5,mous:5,much:5,multi:4,multidimension:4,must:[3,4,5],n_cluster:[3,5],n_compon:4,n_dim:4,n_iter:4,n_param:4,n_pc:5,n_sampl:4,n_strata:5,n_window:5,name:5,ndarrai:[3,4,5],nearest:5,need:5,neighbor:[4,5],neighborhood:4,network:0,network_enhanc:5,neumann:4,nmd:4,non:4,none:[3,5],norm:4,normal:3,now:[3,4,5],number:[3,4,5],numer:5,numpi:[3,4,5],object:[3,4,5],oe_norm:5,onli:5,oper:5,option:[3,4,5],order:5,other:[3,5],our:5,out_fil:3,output:3,p_ij:4,packag:1,page:0,pairwis:[4,5],pairwise_dist:[4,5],param:[4,5],paramet:[3,4,5],pass:[3,5],path:[3,5],pca:4,peng:5,perat:5,percent:5,perp:4,perplex:4,phate:[4,5],plot:[3,5],plot_contact:5,pna:5,point:[3,4,5],point_siz:3,pos1:5,pos2:5,posit:5,position_1:5,position_2:5,potenti:4,pow:5,power:5,prabhakar:3,pre:5,previou:5,princip:[4,5],print:5,print_tim:[4,5],probabl:4,process:5,processing_util:[1,2],project:[4,5],proper:5,proport:5,propot:5,provid:5,pyhic:5,pyplot:3,python:5,q_ij:4,qualifi:5,qualiti:5,raghavan:3,random:[0,5],random_walk:5,random_walk_ratio:5,rang:5,rate:[4,5],read:5,real:5,reduce_spars:[],ref:5,refer:[3,5],reference_genom:5,repres:3,reproduc:[0,1,2,5],requr:5,res:5,resolut:5,resolution_adjust:5,respect:4,retriev:3,return_dist:5,robust:5,row:3,run:4,same:3,sampl:4,scale:4,scatter:[3,5],schic:[3,5],schiclust:5,scipi:5,score:5,script:4,search:0,see:5,select:5,select_cel:5,selfish:[0,4,5],separ:5,sequenc:0,set:[3,5],shape:[4,5],should:3,sigma:[4,5],similar:[4,5],similarity_method:[4,5],simpli:5,singl:[0,3,4,5],situat:5,size:[3,4,5],skip:5,smooth:[0,5],sne:4,some:5,sometim:5,space:[4,5],spars:5,specif:[4,5],specifi:[3,5],specifii:3,spectral:[3,4],spectral_clust:[3,5],spectral_embed:5,spectralembed:4,stage:4,start:3,stochast:4,stop:[4,5],store:5,store_full_map:5,str:[3,4,5],strata:[4,5],student:4,sty:3,style:3,submodul:[1,2],subpackag:1,support:[3,4,5],t_max:4,term:5,than:5,thi:[3,4,5],threshold:[4,5],throughput:0,time:[4,5],titl:3,tol:4,too:5,toolbox:0,trajectori:4,tsne:[4,5],tutori:3,two:5,txt:5,type:[3,4,5],ulrik:3,unnorm:3,unravel:4,use:[3,5],used:[3,4,5],using:[4,5],valu:[3,5],vc_norm:5,vc_sqrt_norm:5,visual:[1,2],vne:4,von:[3,4],walk:[0,5],weight:3,when:5,whether:[3,4,5],which:[0,3,5],window:[4,5],window_s:4,within:5,xinjun:[3,4],yield:5,you:5,zhou:5,zip:5},titles:["Welcome to scHiCTools\u2019s documentation!","scHiCTools","scHiCTools package","scHiCTools.analysis package","scHiCTools.embedding package","scHiCTools.load package"],titleterms:{analysi:3,cluster:3,contactmap:5,content:[2,3,4,5],document:0,embed:4,indic:0,introduct:0,load:5,load_hic_fil:5,modul:[2,3,4,5],packag:[2,3,4,5],procedur:0,processing_util:5,reproduc:4,schictool:[0,1,2,3,4,5],setup:0,submodul:[3,4,5],subpackag:2,tabl:0,visual:3,welcom:0}})
```
