## Supplementary material for "scHiCTools: a computational toolbox for analyzing single-cell Hi-C data": S4: Makefile

```
# Minimal makefile for Sphinx documentation
#

# You can set these variables from the command line, and also
# from the environment for the first two.
SPHINXOPTS    ?=
SPHINXBUILD   ?= sphinx-build
SOURCEDIR     = source
BUILDDIR      = build

# Put it first so that "make" without argument is like "make help".
help:
	@$(SPHINXBUILD) -M help "$(SOURCEDIR)" "$(BUILDDIR)" $(SPHINXOPTS) $(O)

.PHONY: help Makefile

# Catch-all target: route all unknown targets to Sphinx using the new
# "make mode" option.  $(O) is meant as a shortcut for $(SPHINXOPTS).
%: Makefile
	@$(SPHINXBUILD) -M $@ "$(SOURCEDIR)" "$(BUILDDIR)" $(SPHINXOPTS) $(O)
```
