## Supplementary material for "scHiCTools: a computational toolbox for analyzing single-cell Hi-C data": S4: conf.py

```
# Configuration file for the Sphinx documentation builder.
#
# This file only contains a selection of the most common options. For a full
# list see the documentation:
# https://www.sphinx-doc.org/en/master/usage/configuration.html

# -- Path setup --------------------------------------------------------------

# If extensions (or modules to document with autodoc) are in another directory,
# add these directories to sys.path here. If the directory is relative to the
# documentation root, use os.path.abspath to make it absolute, like shown here.
#
import os
import sys
sys.path.insert(0, os.path.abspath('../../scHiCTools'))


# -- Project information -----------------------------------------------------

project = 'scHiCTools'
copyright = '2020, Xinjun Li'
author = 'Xinjun Li'

# The full version, including alpha/beta/rc tags
release = '0.0.3'


# -- General configuration ---------------------------------------------------

# Add any Sphinx extension module names here, as strings. They can be
# extensions coming with Sphinx (named 'sphinx.ext.*') or your custom
# ones.
extensions = [
    'rinoh.frontend.sphinx',
    'sphinx.ext.napoleon'
]
# Napoleon settings
napoleon_google_docstring = True
napoleon_numpy_docstring = True
napoleon_include_init_with_doc = True
napoleon_include_private_with_doc = False
napoleon_include_special_with_doc = True
napoleon_use_admonition_for_examples = False
napoleon_use_admonition_for_notes = False
napoleon_use_admonition_for_references = False
napoleon_use_ivar = False
napoleon_use_param = True
napoleon_use_rtype = False

# Add any paths that contain templates here, relative to this directory.
templates_path = ['_templates']

# List of patterns, relative to source directory, that match files and
# directories to ignore when looking for source files.
# This pattern also affects html_static_path and html_extra_path.
exclude_patterns = []


# -- Options for HTML output -------------------------------------------------

# The theme to use for HTML and HTML Help pages.  See the documentation for
# a list of builtin themes.
#
html_theme = 'sphinx_rtd_theme'

# Add any paths that contain custom static files (such as style sheets) here,
# relative to this directory. They are copied after the builtin static files,
# so a file named "default.css" will overwrite the builtin "default.css".
html_static_path = ['_static']
```
