## Supplementary material for "scHiCTools: a computational toolbox for analyzing single-cell Hi-C data": S4: example1.py

```
import sys
import os
import matplotlib.pyplot as plt

sys.path.insert(0, os.path.abspath(os.path.dirname(__file__)))

print(sys.path)
from scHiCTools import scHiCs,scatter

x = scHiCs(['../test/data/cell_01', '../test/data/cell_02',
            '../test/data/cell_03'],
           reference_genome='mm9',
           resolution=50000,
           max_distance=4000000,
           format='shortest_score',
           adjust_resolution=True,
           chromosomes='except Y',
           operations=['convolution'],
           kernel_shape=3,
           keep_n_strata=10,
           store_full_map=True
           )

x.plot_contacts()


emb, _ = x.learn_embedding(similarity_method='innerproduct',
                           embedding_method='MDS',
                           aggregation='median',
                           print_time=False,
                           return_distance=True)

print(emb)
print(_)


plt.figure()
scatter(emb,label=['01','02','03'],point_size=5)
plt.show()
```
