## Supplementary material for "scHiCTools: a computational toolbox for analyzing single-cell Hi-C data": S4: meta.yaml

```
package:
  name: schictools
  version: 0.0.3
  setup_py_install: true

python:
  version: 3.6
  install:
    - requirements: requirements.txt

about:
  summary: This is a computational tool for analyzing single cell Hi-C data.
```
