## Supplementary material for "scHiCTools: a computational toolbox for analyzing single-cell Hi-C data": S4: README.md

# scHiCTools

### Summary
A computational toolbox for analyzing single cell Hi-C (high-throughput sequencing for 3C) data which includes functions for:
1. Load single-cell HiC datasets
2. Smoothing the contact maps with linear convolution, random walk or network enhancing
3. Calculating embeddings for single cell HiC datasets efficiently with reproducibility measures include InnerProduct, HiCRep and Selfish

### Installation
  **Required Python Packages**
  - Python (version >= 3.6)
  - numpy (version >= 1.15.4)
  - scipy (version >= 1.0)
  - matplotlib (version >=3.1.1)
  - pandas (version >=0.19)
  - simplejson
  - six
  - h5py
  
  **`interactive_scatter` feature requirement**
  - plotly (version >= >=4.8.0)

  **Install from GitHub**

  You can install the package with following command:

  ```console
    $ git clone https://github.com/liu-bioinfo-lab/scHiCTools.git
    $ cd scHiCTools
    $ python setup.py install
  ```

  **Install from PyPI**

  ```console
    $ pip install scHiCTools
  ```

  **Install optional interactive dependencie**
  
  ```console
    $ pip install scHiCTools[interactive_scatter]
  ```
  or
  ```console
    $ pip install -e .[interactive_scatter]
  ```


### Usage
   **Supported Formats**

  - Pre-processed Matrices:
  If the data is already processed into matrices for intra-chromosomal contacts,
  the chromosome from the same cell must be stored in the same folder with
  chromosome names as file names (e.g., scHiC/cell_1/chr1.txt).
  You only need to provide the folder name for a cell (e.g., scHiC/cell_1).
    - npy: numpy.array / numpy.matrix
    - npz: scipy.sparse.coo_matrix
    - matrix: matrix stored as pure text
    - matrix_txt: matrix stored as .txt file
    - HiCRep: the format required by HiCRep package

  - Edge List <br />
   For all formats below:<br />
   &nbsp; str - strand (forward / reverse)<br />
   &nbsp; chr - chromosome<br />
   &nbsp; pos - position<br />
   &nbsp; score - contact reads<br />
   &nbsp; frag - fragments (will be ignored)<br />
   &nbsp; mapq - map quality<br />

    - Shortest
     ```
    <chr1> <pos1> <chr2> <pos2>
     ```
    - Shortest_Score
     ```
    <chr1> <pos1> <chr2> <pos2> <score>
     ```
    - Short
     ```
    <str1> <chr1> <pos1> <frag1> <str2> <chr2> <pos2> <frag2>
     ```
    - Short_Score
     ```
    <str1> <chr1> <pos1> <frag1> <str2> <chr2> <pos2> <frag2> <score>
     ```
    - Medium
     ```
    <readname> <str1> <chr1> <pos1> <frag1> <str2> <chr2> <pos2> <frag2> <mapq1> <mapq2>
     ```
    - Long
     ```
    <str1> <chr1> <pos1> <frag1> <str2> <chr2> <pos2> <frag2> <mapq1> <cigar1> <sequence1> <mapq2> <cigar2> <sequence2> <readname1> <readname2>
     ```
    - 4DN
     ```
    ## pairs format v1.0
    #columns: readID chr1 position1 chr2 position2 strand1 strand2
     ```
   - .hic format: we adapted "straw" from JuiceTools.

   - .mcool format: we adapted "dump" from cool.

   - Other formats: simply give the indices (start from 1) in the order of<br />
 "chromosome1 - position1 - chromosome2 - position2 - score" or<br />
 "chromosome1 - position1 - chromosome2 - position2" or<br />
 "chromosome1 - position1 - chromosome2 - position2 - mapq1 - mapq2".<br />
   For example, you can provide "2356" or [2, 3, 5, 6] if the file takes this format:
   ```
   <name> <chromosome1> <position1> <frag1> <chromosome2> <position2> <frag2> <strand1> <strand2>
   contact_1 chr1 3000000 1 chr1 3001000 1 + -
   ```

  **Import Package**
  ```console
  >>>import scHiCTools
  ```

  **Load scHiC data**

  The scHiC data is stored in a series of files, with each of the files corresponding to one cell.
  You need to specify the list of scHiC file paths.

  Only intra-chromosomal interactions are counted.
  ```console
  >>>from scHiCTools import scHiCs
  >>>files = ['./cell_1', './cell_2', './cell_3']
  >>>loaded_data = scHiCs(
  ... files, reference_genome='mm9',
  ... resolution=500000, keep_n_strata=10,
  ... format='customized', adjust_resolution=True,
  ... customized_format=12345, header=0, chromosomes='except Y',
  ... operations=['OE_norm', 'convolution']
  ... )
  ```
  - reference genome (dict or str): now supporting 'mm9', 'mm10', 'hg19', 'hg38'.
  If your reference genome is not in ['mm9', 'mm10', 'hg19', 'hg38'], you need to provide the lengths of chromosomes
  you are going to use with a Python dict. e.g. {'chr1': 150000000, 'chr2': 130000000, 'chr3': 200000000}
  - resolution (int): the resolution to separate genome into bins.
  If using .hic file format, the given resolution must match with the resolutions in .hic file.
  - keep_n_strata (None or int): only store contacts within n strata near the diagonal. Default: 10.
  If 'None', it will not store strata
  - store_full_map (bool): whether store full contact maps in numpy matrices or
  scipy sparse matrices，If False, it will save memory.
  - sparse (bool): whether to use sparse matrices
  - format (str): file format, supported formats: 'shortest', 'shortest_score', 'short',
  'short_score' , 'medium', 'long', '4DN', '.hic', '.mcool', 'npy', 'npz', 'matrix',
  'HiCRep', 'matrix_txt' and 'customized'. Default: 'customized'.
  - customized_format (int or str or list): the column indices in the order of
  "chromosome 1 - position 1 - chromosome 2 - position 2 - contact reads" or
  "chromosome 1 - position 1 - chromosome 2 - position 2" or
  "chromosome 1 - position 1 - chromosome 2 - position 2 - map quality 1 - map quality 2".
  e.g. if the line is "chr1 5000000 chr2 3500000 2", the format should be '12345' or [1, 2, 3, 4, 5]; if there is no column
  indicating number of reads, you can just provide 4 numbers like '1234', and contact read will be set as 1.
  Default: '12345'.
  - adjust_resolution: whether to adjust resolution for the input file.
  Sometimes the input file is already in the proper resolution (e.g. position 3000000 has already been changed to 6 with 500kb resolution).
  For this situation you can set adjust_resolution=False. Default: True.
  - map_filter (float): keep all contacts with mapq higher than this threshold. Default: 0.0
  - header (int): how many header lines does the file have. Default: 0.
  - chromosomes (list or str): chromosomes to use, eg. ['chr1', 'chr2'], or
  just 'except Y', 'except XY', 'all'.
  Default: 'all', which means chr 1-19 + XY for mouse and chr 1-22 + XY for human.
  - operations (list or None): the operations use for pre-processing or smoothing the maps given in a list.
  The operations will happen in the given order.
  Supported operations: 'logarithm', 'power', 'convolution', 'random_walk',
  'network_enhancing', 'OE_norm', 'VC_norm', 'VC_SQRT_norm', 'KR_norm'。
  Default: None.
  - For preprocessing and smoothing operations, sometimes you need additional arguments
  (introduced in next sub-section).

  You can also skip pre-processing and smoothing in loading step (operations=None),
  and do them in next lines.

  **Plot number of contacts and select cells**

  You can plot the number of contacts of your cells.
  ```console
  >>>loaded_data.plot_contacts(hist=True, percent=True)
  ```
  If hist is `True`, plot Histogram of the number of contacts. If percent is `True`, plot the scatter plot of cells with  of short-range contacts (< 2 Mb) versus contacts at the mitotic band (2-12 Mb).

  You can select cells based on number of contacts:
  ```console
  >>>loaded_data.select_cells(min_n_contacts=10000,max_short_range_contact=0.99)
  ```
  The above command select cells have number of contacts bigger than 10000 and percent of short range contacts small than .99.

  **Pre-processing and Smoothing Operations**
  Stand alone pre-processing and smoothing:
  ```console
  >>>loaded_data.processing(['random_walk', 'network_enhancing'])
  ```
  If you didn't store full map (i.e. store_full_map=False), processing is not
  doable in a separate step.

  - logarithm:
  new_W_ij = log_(base) (W_ij + epsilon). Additional arguments:
    - log_base: default: e
    - epsilon: default: 1
  - power: new_W_ij = (W_ij)^pow. Additional argument:
    - pow: default: 0.5 (i.e., sqrt(W_ij))

  - VC_norm: VC normalization - each value divided by the sum of
  corresponding row then divided by the sum of corresponding column
  - VC_SQRT_norm: VC_SQRT normalization - each value divided by the sqrt of the sum
  of corresponding row then divided by the sqrt of the sum of corresponding column
  - KR_norm: KR normalization - iterating until the sum of each row / column is one
  Argument:
    - maximum_error_rate (float): iteration ends when max error is smaller
    than (maximum_error_rate). Default: 1e-4
  - OE_norm: OE normalization -  each value divided by the average of its
  corresponding strata (diagonal line)

  - convolution: smoothing with a N by N convolution kernel, with each value equal to 1/N^2.
  Argument:
    - kernel_shape: an integer. e.g. kernel_shape=3 means a 3*3 matrix with each value = 1/9. Default: 3.
  - Random walk: multiply by a transition matrix (also calculated from contact map itself).
  Argument:
    - random_walk_ratio: a value between 0 and 1, e.g. if ratio=0.9, the result will be
    0.9 * matrix_after_random_walk + 0.1 * original_matrix. Default: 1.0.
  - Network enhancing: transition matrix only comes from k-nearest neighbors of each line.
  Arguments:
    - kNN: value 'k' in kNN. Default: 20.
    - iterations: number of iterations for network enhancing. Default: 1
    - alpha: similar with random_walk_ratio. Default: 0.9

  **Learn Embeddings**
  ```console
  >>>embs = loaded_data.learn_embedding(
  ... dim=2, similarity_method='inner_product', embedding_method='MDS',
  ... n_strata=None, aggregation='median', return_distance=False
  ... )
  ```
  This function will return the embeddings in the format of a numpy array with shape ( # of cells, # of dimensions).
  - dim (int): the dimension for the embedding
  - similarity_method (str): reproducibility measure, 'InnerProduct', 'HiCRep' or
  'Selfish'. Default: 'InnerProduct'
  - embedding_method (str): 'MDS', 'tSNE' or 'UMAP'
  - n_strata (int): only consider contacts within this genomic distance. Default: None.
  If it is None, it will use the all strata kept (the argument keep_n_strata from
  previous loading process). Thus n_strata and keep_n_strata (loading step) cannot be
  None at the same time.
  - aggregation (str): method to aggregate different chromosomes,
  'mean' or 'median'. Default: 'median'.
  - return_distance (bool): if True, return (embeddings, distance_matrix); if False, only return embeddings. Default: False.
  - Some additional argument for Selfish:
    - n_windows (int): split contact map into n windows, default: 10
    - sigma (float): sigma in the Gaussian-like kernel: default: 1.6


  **Clustering**

  There are two functions to cluster cells.

  ```console
  >>>label=loaded_data.clustering(
  ... n_clusters=4, clustering_method='kmeans', similarity_method='innerproduct',
  ... aggregation='median', n_strata=None
  ... )
  ```

  `clustering` function returns a numpy array of cell labels clustered.

  - n_clusters (int): Number of clusters.

  - clustering_method (str):
    Clustering method in 'kmeans', 'spectral_clustering' or 'HAC'(hierarchical agglomerative clustering).

  - similarity_method (str):
    Reproducibility measure.
    Value in ‘InnerProduct’, ‘HiCRep’ or ‘Selfish’.

  - aggregation (str):
    Method to aggregate different chromosomes.
    Value is either 'mean' or 'median'.
    Default: 'median'.

  - n_strata (int or None):
    Only consider contacts within this genomic distance.
    If it is None, it will use the all strata kept (the argument keep_n_strata) from previous loading process. Default: None.

  - print_time (bool):
    Whether to print the processing time. Default: False.


  ```console
  >>>hicluster=loaded_data.scHiCluster(dim=2,cutoff=0.8,n_PCs=10,k=4)
  ```

  `scHiCluster` function returns two componments.
  First componment is a numpy array of embedding of cells using HiCluster.
  Second componment is a numpy of cell labels clustered by HiCluster.

  - dim (int): Number of dimension of embedding. Default: 2.

  - cutoff (float): The cutoff proportion to convert the real contact matrix into binary matrix. Default: 0.8.

  - n_PCs (int): Number of principal components. Default: 10.

  - k (int): Number of clusters. Default: 4.


  **Visualization**

  ```console
  >>>scatter(data, dimension="2D", point_size=3, sty='default',
  ... label=None, title=None, alpha=None, aes_label=None
  ... )
  >>>plt.show()
  ```

  This function is to plot scatter plot of embedding points of single cell data.
    Scatter plot of either two-dimensions or three-dimensions will be generated.


  - data (numpy.array): A numpy array which has 2 or 3 columns, every row represent a point.

  - dimension (str): Specifiy the dimension of the plot, either "2D" or "3D". Default: "2D".

  - point_size (float): Set the size of the points in scatter plot. Default: 3.

  - sty (str): Styles of Matplotlib. Default: 'default'.

  - label (list or None): specifiy the label of each point. Default: None.

  - title (str): Title of the plot. Default: None.

  - alpha (float): The alpha blending value. Default: None.

  - aes_label (list): Set the label of every axis. Default: None.


"scHiCTools" also support interactive scatter plot which require the module 'plotly'

  ```console
  >>>interactive_scatter(loaded_data, data, out_file, dimension='2D', point_size=3,
  ... label=None, title=None, alpha=1, aes_label=None)
  ```

  This function is to generate an interactive scatter plot of embedded single cell data.
    The plot will be stored in a file.

  - schic (scHiCs): A `scHiCs` object.
  - data (numpy.array): A numpy array which has 2 or 3 columns, every row represent a point.
  - out_file (str): Output file path.
  - dimension (str): Specifiy the dimension of the plot, either "2D" or "3D". The default is "2D".
  - point_size (float): Set the size of the points in scatter plot. The default is 3.
  - label (list or None): Specifiy the label of each point. The default is None.
  - title (str): Title of the plot. The default is None.
  - alpha (float): The alpha blending value. The default is 1.
  - aes_label (list): Set the label of every axis. The default is None.


### Citation
Xinjun Li, Fan Feng, Wai Yan Leung and Jie Liu. "scHiCTools: a computational toolbox for analyzing single cell Hi-C data."
bioRxiv (2019): 769513.

### References
A. R. Ardakany, F. Ay, and S. Lonardi.  Selfish: Discovery of differential chromatininteractions via a self-similarity measure.bioRxiv, 2019.

N. C. Durand, J. T. Robinson, M. S. Shamim, I. Machol, J. P. Mesirov, E. S. Lander, and E. Lieberman Aiden. "Juicebox provides a visualization system for Hi-C contact maps with unlimited zoom." Cell Systems 3(1), 2016.

J. Liu, D. Lin, G. Yardimci, and W. S. Noble. Unsupervised embedding of single-cellHi-C data.Bioinformatics, 34:96–104, 2018.

T. Nagano, Y. Lubling, C. Várnai, C. Dudley, W. Leung, Y. Baran, N. M. Cohen,S.  Wingett,  P.  Fraser,  and  A.  Tanay.    Cell-cycle  dynamics  of  chromosomal organization at single-cell resolution.Nature, 547:61–67, 2017.

B. Wang, A. Pourshafeie, M. Zitnik, J. Zhu, C. D. Bustamante, S. Batzoglou, andJ.  Leskovec.   Network  enhancement  as  a  general  method  to  denoise  weighted biological networks.Nature Communications, 9(1):3108, 2018.

T. Yang, F. Zhang, G. G. Y. mcı, F. Song, R. C. Hardison, W. S. Noble, F. Yue, andQ. Li. HiCRep: assessing the reproducibility of Hi-C data using a stratum-adjusted correlation coefficient.Genome Research, 27(11):1939–1949, 2017.

G.  G.  Yardımcı,   H.  Ozadam,   M.  E.  Sauria,   O.  Ursu,   K. K.  Yan,   T.  Yang,A. Chakraborty, A. Kaul, B. R. Lajoie, F. Song, et al. Measuring the reproducibilityand quality of hi-c data.Genome Biology, 20(1):57, 2019.
