## Supplementary material for "scHiCTools: a computational toolbox for analyzing single-cell Hi-C data": S4: clustering.py

```
# -*- coding: utf-8 -*-
"""

 Clustering component of scHiCTools

 Author: Xinjun Li

 This module define functions to clustering single cell data.

"""

import numpy as np
import scipy.spatial.distance as dis
from scipy.sparse import csgraph


def kmeans(data,
           k=4,
           weights=None,
           iteration=1000,
           **kwargs):
    """
    k-means algorithm, with k-means++ to initialize the start points.
    

    Parameters
    ----------
    data : numpy.ndarray
        Coordiante of points.
        
    k : int, optional
        Number of clusters.
        The default is 4.
        
    weights : list, optional
        List of the weight of every point,
        should either be 'None' or have the length same as the number of rows in data. 
        The default is None.
        
    iteration : int, optional
        Number of iterations in k-means algorithm.
        The default is 1000.
    

    Returns
    -------
    label : numpy.ndarray
        A list of the label of every point.

    """
    
    # kmeans++
    data=np.array(data)
    n,p=data.shape
    cen=[None]*k
    p=np.ones(n)/n
    index=np.arange(n)
    for i in range(k):
        cen[i]=int(np.random.choice(index,size=1,p=p))
        p=np.array([np.linalg.norm(data-data[cen[j]],axis=1) for j in np.arange(i+1)]).min(axis=0)
        # print('p:',p,'\n','data:', data)
        p=p/sum(p)
    centroids=data[cen]
    
    
    label=np.zeros(len(data))

    # loop to find the proper cluster
    for i in range(iteration):

        # calculate the distance to the centroids respectively
        dist=np.zeros((len(data), k))
        for cent in range(k):
            dist[:,cent]=np.sum(np.square(data-centroids[cent]), axis=1)

        # find the lable under the centroids
        label0=np.argmin(dist,axis=1)
        if sum(label!=label0)==0:
            break
        else:
            label=label0

        # renew centroids
        if weights is not None:
            weights=np.array(weights)
            for j in range(k):
                group=data[label==j,:]
                centroids[j]=np.average(group, axis=0,weights=weights[label==j])
        else:
            for j in range(k):
                group=data[label==j,:]
                centroids[j]=np.average(group, axis=0)

    return label


# spectral_clustering can directly takes distance matrix.
def spectral_clustering(data,
                        data_type='points',
                        n_clusters=4,
                        normalize=True,
                        **kwargs):
    """
    This function is a impliment of unnormalized spectral clustering.
    Reference: "A Tutorial on Spectral Clustering" by Ulrike von Luxburg.
    

    Parameters
    ----------
    data : numpy.ndarray
        Distance matrix or coordinate of the points.
        
    data_type : str, optional
        Format of argument `data`, specify the type of data input. 
        Must be in either 'points' or 'distance_matrix'.
        The default is 'points'.
        
    n_clusters : int, optional
        Number of clusters.
        The default is 4.
        
    normalize : bool, optional
        Whether to use unnormalized or normalized spectral clustering.
        The default is True.
        
    **kwargs :
        Arguments pass to kmeans.


    Returns
    -------
    label : numpy.ndarray
        A list of the label of every point.

    """

    # compute graph similarity matrix
    if data_type=='distance_matrix' and len(data)==len(data[0]):
        if len(data)!=len(data[0]):
            raise ValueError('data is not a distance matrix!')
        elif np.sum(data.T!=data)>0:
            raise ValueError('data is not a distance matrix!')
        elif sum(np.diag(data)!=0)!=0:
            raise ValueError('data is not a distance matrix!')
        # using fully connected graph with Gaussian similarity function
        graph=np.exp(-np.square(data)/np.mean(data**2))
    elif data_type=='points':
        # pair-wise Euclidean distance
        graph = dis.pdist(data)
        graph = dis.squareform(graph)
        # graph = np.add(np.add(-2 * np.dot(data, data.T), graph).T, graph)
        # using fully connected graph with Gaussian similarity function
        graph = np.exp(-np.square(graph)/np.mean(graph**2))
    else:
        raise ValueError('data_type is not supported.')

    graph = graph-np.diag(graph.diagonal())


    if normalize: # normalized clustering

        # the normalized Laplacian L.
        L=csgraph.laplacian(graph, normed=True)

        # eigenvalues and eigenvectors.
        eig_vals, eig_vecs = np.linalg.eig(L)
        eig_vecs= eig_vecs[:, (eig_vals.argsort())]

        # first k=n_clusters eigenvectors.
        U=eig_vecs[:,:n_clusters]
        U=(U.T/np.linalg.norm(U, axis=1)).T

    else: # unnormalized clustering

        # the unnormalized Laplacian L.
        L=csgraph.laplacian(graph, normed=False)

        # eigenvalues and eigenvectors.
        eig_vals, eig_vecs = np.linalg.eig(L)
        eig_vecs= eig_vecs[:, (eig_vals.argsort())]

        # first k=n_clusters eigenvectors.
        U=eig_vecs[:,:n_clusters]

    # Clustering using k-means
    label=kmeans(U,k=n_clusters,**kwargs)

    return label


# Bottom-up hierarchical clustering algorithms:
#   hierarchical agglomerative clustering(HAC)
# HAC can directly take distance matrix.
def HAC(data,
        data_type='points',
        n_clusters=4,
        method='centroid'):
    """
    This function is a impliment of hierarchical afflomerative clustering.
    Reference: 
        Christopher D. Manning, Prabhakar Raghavan and Hinrich Schütze,
        "Introduction to Information Retrieval" chapter 17
    

    Parameters
    ----------
    data : numpy.ndarray
        Distance matrix or coordinate of the points.
    
    data_type : str, optional
        Format of argument `data`, specify the type of data input. 
        Must be in either 'points' or 'distance_matrix'.
        The default is 'points'.
        
    n_clusters : int, optional
        Number of clusters.
        The default is 4.
        
    method : str, optional
        Specify the hierarchical clustering method used.
        Now support 'single-link', 'complete-link', 'centroid','group-average'.
        The default is 'centroid'.


    Returns
    -------
    label : numpy.ndarray
        A list of the label of every point.

    """

    # compute graph similarity matrix
    if data_type=='distance_matrix' and len(data)==len(data[0]):
        if len(data)!=len(data[0]):
            raise ValueError('data is not a distance matrix!')
        elif np.sum(data.T!=data)>0:
            raise ValueError('data is not a distance matrix!')
        elif sum(np.diag(data)!=0)!=0:
            raise ValueError('data is not a distance matrix!')
        # using fully connected graph with Gaussian similarity function
        graph=np.exp(-np.square(data)/np.mean(data**2)) # sigma?
    elif data_type=='points':
        # pair-wise Euclidean distance
        # graph = np.sum(np.square(data), 1)
        # graph = np.add(np.add(-2 * np.dot(data, data.T), graph).T, graph)
        graph = dis.pdist(data)
        graph = dis.squareform(graph)
        # using fully connected graph with Gaussian similarity function
        graph = np.exp(-np.square(graph)/np.mean(graph**2)) # sigma?
    else:
        raise ValueError('data_type is not supported.')

    similarity=graph-np.diag(np.diag(graph))
    label=np.array(range(len(graph)))


    if method in ['single-link','complete-link']:

        while max(label)>=n_clusters:

            # find maximum similarity between groups
            index=np.argmax(similarity)
            index=[index//len(similarity), index%len(similarity)]

            # combine similarity groups
            label[label==index[1]]=index[0]
            label[label>index[1]]-=1

            # new similarity matrix
            if method=='single-link':
                new_simi=np.amax(similarity[:,index],axis=1)
            else:
                new_simi=np.amin(similarity[:,index],axis=1)
            similarity[:,index[0]]=new_simi
            similarity[index[0],:]=new_simi
            similarity=np.delete(
                    np.delete(similarity,index[1],axis=0),
                    index[1],
                    axis=1)
            similarity[index[0],index[0]]=0

    elif method=='centroid':
        while max(label)>=n_clusters:

            # find maximum similarity between groups
            index=np.argmax(similarity)
            index=[index//len(similarity), index%len(similarity)]

            # number of obs contains in the two group
            n0=sum(label==index[0])
            n1=sum(label==index[1])

            # combine similarity groups
            label[label==index[1]]=index[0]
            label[label>index[1]]-=1

            # new similarity matrix
            new_simi=(similarity[:,index[0]]*n0 + similarity[:,index[1]]*n1) / (n0+n1)
            similarity[:,index[0]]=new_simi
            similarity[index[0],:]=new_simi
            similarity=np.delete(
                    np.delete(similarity,index[1],axis=0),
                    index[1],axis=1)
            similarity[index[0],index[0]]=0

    elif method=='group-average':
        # A copy of similarity matrix
        simi0=similarity

        while max(label)>=n_clusters:
            # find maximum similarity between groups
            index=np.argmax(similarity)
            index=[index//len(similarity), index%len(similarity)]

            # combine similarity groups
            label[label==index[1]]=index[0]
            label[label>index[1]]-=1

            # new similarity matrix
            for i in range(max(label)):
                new_simi=np.mean(simi0[label==i][:,label==index[0]])
                similarity[i,index[0]]=new_simi
                similarity[index[0],i]=new_simi
            similarity=np.delete(
                    np.delete(similarity,index[1],axis=0),
                    index[1],axis=1)
            similarity[index[0],index[0]]=0

    else:
        raise ValueError('Method is not supported.')


    return label
```
