## Supplementary material for "scHiCTools: a computational toolbox for analyzing single-cell Hi-C data": S4: visualization.py

```
# -*- coding: utf-8 -*-
"""

 Visualization component of scHiCTools

 Author: Xinjun Li

 This module define a function to plot scatter plot of embedding points of single cell data.

"""


import numpy as np
import pandas as pd
import matplotlib as mpl
import matplotlib.pyplot as plt
from mpl_toolkits.mplot3d import Axes3D
try:
    import plotly.express as px
except:
    px=None


def scatter(data, dimension="2D", point_size=3, sty='default',
            label=None, title=None, alpha=None, aes_label=None,
            **kwargs):
    """
    This function is to plot scatter plot of embedding points
        of single cell data.
    
    Parameters
    ----------
    data : numpy.array
        A numpy array which has 2 or 3 columns, every row represent a point.
    dimension : str, optional
        Specifiy the dimension of the plot, either "2D" or "3D".
        The default is "2D".
    point_size : float, optional
        Set the size of the points in scatter plot.
        The default is 3.
    sty : str, optional
        Styles of Matplotlib. The default is 'default'.
    label : list or None, optional
        Specifiy the label of each point. The default is None.
    title : str, optional
        Title of the plot. The default is None.
    alpha : float, optional
        The alpha blending value. The default is None.
    aes_label : list, optional
        Set the label of every axis. The default is None.
    **kwargs :
        Other arguments passed to the `matplotlib.pyplot.legend`,
        controlling the plot's legend.

    Returns
    -------
        Scatter plot of either 2D or 3D.

    """

    # Error messages.
    if dimension not in ["2D", "3D"]:
        raise ValueError('Dimension must be "2D" or "3D".')
    if (dimension=="2D" and len(data[0])!=2) or (dimension=="3D" and len(data[0])!=3):
        raise ValueError('Data shape must match dimension!')
    if (label is not None) and len(data)!=len(label):
        raise ValueError('Number of rows in data must equal to length of label!')


    mpl.style.use(sty)

    # 2D scatter plot
    if dimension=="2D":

        # Plot with label
        if label is not None:
            label=np.array(label)
            lab=list(set(label))
            for index, l in enumerate(lab):
                plt.scatter(data[label==l,0], data[label==l,1],
                                c='C{!r}'.format(index),
                                s=point_size, label=l,
                                alpha=alpha)
            plt.legend(**kwargs)
        # Plot without label
        else:
            plt.scatter(data[:,0], data[:,1], s=point_size, alpha=alpha)

        if aes_label is not None:
                plt.xlabel(aes_label[0])
                plt.ylabel(aes_label[1])


    # 3D scatter plot
    if dimension=="3D":
        splot = plt.subplot(111, projection='3d')

        # Plot with label
        if label is not None:
            label=np.array(label)
            lab=list(set(label))

            for index, l in enumerate(lab):
                splot.scatter(data[label==l,0], data[label==l,1], data[label==l,2],
                              s=point_size,
                              color='C{!r}'.format(index),
                              label=l)
            plt.legend(**kwargs)
        # Plot without label
        else:
            splot.scatter(data[:,0], data[:,1], data[:,2],s=point_size)

        if aes_label is not None:
            splot.set_xlabel(aes_label[0])
            splot.set_ylabel(aes_label[1])
            splot.set_zlabel(aes_label[2])

    plt.title(title)


def interactive_scatter(schic, data, out_file, dimension="2D", point_size=3,
                        label=None, title=None, alpha=1, aes_label=None):
    """
    This function is to generate an interactive scatter plot of embedded single cell data.
    
    Parameters
    ----------
    schic : scHiCs
        A `scHiCs` object.
    data : numpy.array
        A numpy array which has 2 or 3 columns, every row represent a point.
    out_file : str
        Output file path.
    dimension : str, optional
        Specifiy the dimension of the plot, either "2D" or "3D".
        The default is "2D".
    point_size : float, optional
        Set the size of the points in scatter plot.
        The default is 3.
    label : list or None, optional
        Specifiy the label of each point. The default is None.
    title : str, optional
        Title of the plot. The default is None.
    alpha : float, optional
        The alpha blending value. The default is 1.
    aes_label : list, optional
        Set the label of every axis. The default is None.

    """
    
    # Error messages.
    if px is None:
        raise ImportError('Need `plotly` installed to use function `interactive_scatter`.')
    if dimension not in ["2D", "3D"]:
        raise ValueError('Dimension must be "2D" or "3D".')
    if (dimension=="2D" and len(data[0])!=2) or (dimension=="3D" and len(data[0])!=3):
        raise ValueError('Data shape must match dimension!')
    if (label is not None) and len(data)!=len(label):
        raise ValueError('Number of rows in data must equal to length of label!')
        

    # 2D scatter plot
    if dimension=="2D":
        if aes_label is None:
            aes_label=['x1','x2']

        # Plot with label
        if label is not None:
            df = pd.DataFrame({'cell':schic.files,
                               aes_label[0]:data[:,0],
                               aes_label[1]:data[:,1],
                               'label':label})
            df=df.astype({'label': 'category'})
            fig = px.scatter(df, x=aes_label[0], y=aes_label[1],
                             color="label", hover_data=['cell'],
                             opacity=alpha)
        # Plot without label
        else:
            df = pd.DataFrame({'cell':schic.files,
                               aes_label[0]:data[:,0],
                               aes_label[1]:data[:,1]})
            fig = px.scatter(df, x=aes_label[0], y=aes_label[1],
                             hover_data=['cell'], opacity=alpha)


    # 3D scatter plot
    if dimension=="3D":
        if aes_label is None:
            aes_label=['x1','x2','x3']

        # Plot with label
        if label is not None:
            df = pd.DataFrame({'cell':schic.files,
                               aes_label[0]:data[:,0],
                               aes_label[1]:data[:,1],
                               aes_label[2]:data[:,2],
                               'label':label})
            df=df.astype({'label': 'category'})
            fig = px.scatter_3d(df, x=aes_label[0], y=aes_label[1],
                                z=aes_label[2], color="label",
                                hover_data=['cell'], opacity=alpha)
            
        # Plot without label
        else:
            df = pd.DataFrame({'cell':schic.files,
                               aes_label[0]:data[:,0],
                               aes_label[1]:data[:,1],
                               aes_label[2]:data[:,2]})
            fig = px.scatter_3d(df, x=aes_label[0], y=aes_label[1],
                                z=aes_label[2], hover_data=['cell'],
                                opacity=alpha)

    fig.update_traces(marker=dict(size=point_size),
                      selector=dict(mode='markers'))
    fig.update_layout(title=title)

    fig.write_html(out_file)
```
