## Supplementary material for "scHiCTools: a computational toolbox for analyzing single-cell Hi-C data": S4: embedding.py

```
# -*- coding: utf-8 -*-
"""

 Embedding component of scHiCTools

 Author: Xinjun Li

 This script define functions to embedding single cell data to a lower-dimensional space.

"""


import numpy as np
import scipy.optimize as opt
from scipy.sparse import csgraph
import scipy.spatial.distance as dis
from scipy.optimize import curve_fit
from scipy.spatial.distance import squareform


# PCA :---------------------------------------------------

def PCA(X, dim=2):
    """
    
    Principal components analysis，PCA.
    
    
    Parameters
    ----------
    X : numpy.ndarray
        Coordinates of input data points.
        
    dim : int, optional
        The dimension of the projected points. 
        The default is 2.


    Returns
    -------
    Y : numpy.ndarray
        Coordinates of the projected points.

    """
    
    X=X-np.mean(X,axis=0)
    U, S, V = np.linalg.svd(X, full_matrices=False, compute_uv=True)
    Y = np.dot(V[:dim,:],X.T).T
    return(Y)
    


# MDS :---------------------------------------------------

def MDS(mat, n=2):
    """
    Multidimensional scaling, MDS.
    

    Parameters
    ----------
    mat : numpy.ndarray
        Distance matrix of the data points.
        
    n : int, optional
        The dimension of the projected points. 
        The default is 2.


    Returns
    -------
    co : numpy.ndarray
        Coordinates of the projected points.

    """
    
    
    # mat = np.sqrt(2 - 2 * mat)
    h = np.eye(len(mat)) - np.ones(mat.shape) / len(mat)
    k = -0.5 * h.dot(mat * mat).dot(h)
    if np.any(np.isnan(k)):
        k[np.isnan(k)] = 0
    w, v = np.linalg.eig(k)
    max_ = np.argsort(w)[:-n - 1:-1]
    co = np.real(v[:, max_].dot(np.sqrt(np.diag(w[max_]))))
    # co = np.real(v[:, :2].dot(np.sqrt(np.diag(w[:2]))))
    return co


# tSNE :--------------------------------------------------

MACHINE_EPSILON = np.finfo(np.double).eps

def kl_divergence(params, P, degrees_of_freedom, n_samples, n_components):
    """
    
    t-SNE objective function: gradient of the KL divergence
    of p_ijs and q_ijs and the absolute error.


    Parameters
    ----------
    params : numpy.ndarray, shape (n_params,)
        Unraveled embedding.

    P : numpy.ndarray, shape (n_samples * (n_samples-1) / 2,)
        Condensed joint probability matrix.

    degrees_of_freedom : int
        Degrees of freedom of the Student's-t distribution.

    n_samples : int
        Number of samples.

    n_components : int
        Dimension of the embedded space.
    
    
    Returns
    -------
    kl_divergence : float
        Kullback-Leibler divergence of p_ij and q_ij.

    grad : numpy.ndarray, shape (n_params,)
        Unraveled gradient of the Kullback-Leibler divergence with respect to the embedding.
        
    """
    X_embedded = params.reshape(n_samples, n_components)

    # Q is a heavy-tailed distribution: Student's t-distribution
    dist = dis.pdist(X_embedded, "sqeuclidean")
    dist /= degrees_of_freedom
    dist += 1.
    dist **= (degrees_of_freedom + 1.0) / -2.0
    Q = np.maximum(dist / (2.0 * np.sum(dist)), MACHINE_EPSILON)

    # Optimization trick below: np.dot(x, y) is faster than
    # np.sum(x * y) because it calls BLAS

    # Objective: C (Kullback-Leibler divergence of P and Q)
    kl_divergence = 2.0 * np.dot(
            P, np.log(np.maximum(P, MACHINE_EPSILON) / Q))
    

    # Gradient: dC/dY
    # pdist always returns double precision distances. Thus we need to take
    grad = np.ndarray((n_samples, n_components), dtype=params.dtype)
    PQd = squareform((P - Q) * dist)
    for i in range(n_samples):
        grad[i] = np.dot(np.ravel(PQd[i], order='K'),
                         X_embedded[i] - X_embedded)
    grad = grad.ravel()
    c = 2.0 * (degrees_of_freedom + 1.0) / degrees_of_freedom
    grad *= c

    return kl_divergence, grad


def tSNE(mat,
         n_dim=2,
         perp=30.0,
         n_iter=1000,
         momentum = 0.5,
         rate = 200.0,
         tol=1e-5):
    """
    t-Distributed Stochastic Neighbor Embedding, t-SNE.
    
    
    Parameters
    ----------
    mat : numpy.ndarray
        Distance matrix of the data points.
        
    n_dim : int, optional
        The dimension of the projected points.
        The default is 2.
        
    perp : float, optional
        Perplexity.
        The default is 30.0.
        
    n_iter : int, optional
        Max number of iteration.
        The default is 1000.
        
    momentum : float, optional
        Momentum of gredient decendent.
        The default is 0.5.
        
    rate : float, optional
         Gredient decendent rate.
         The default is 200.
         
    tol : float, optional
         The threshold of gradient norm to stop the iteration of grendient decendent.
         The default is 1e-5.

    Returns
    -------
    Y : numpy.ndarray
        Coordinates of the projected points.

    """

    # Error messagers
    if len(mat)!=len(mat[0]):
        raise ValueError('tSNE input mat is not a distance matrix!')
    elif np.sum(mat.T!=mat)>0:
        raise ValueError('tSNE input mat is not a distance matrix!')
    elif sum(np.diag(mat)!=0)!=0:
        raise ValueError('tSNE input mat is not a distance matrix!')

    
    def Hbeta(D, beta=1.0):
        """
            Compute the perplexity and the P-row for a specific value of the
            precision of a Gaussian distribution.
        """
        # Compute P-row and corresponding perplexity
        P = np.exp(-D.copy() * beta)
        sumP = sum(P)+10**-10
        H = np.log(sumP) + beta * np.sum(D * P) / sumP
        P = P / sumP
        return H, P
    
    
    def x2p(D, tol=1e-5, perplexity=30.0):
        """
            Performs a binary search to get P-values in such a way that each
            conditional Gaussian has the same perplexity.
        """
    
        # Initialize some variables
        n=len(D)
        P = np.zeros((n, n))
        beta = np.ones((n, 1))
        logU = np.log(perplexity)
    
        # Loop over all datapoints
        for i in range(n):
    
            # Compute the Gaussian kernel and entropy for the current precision
            betamin = -np.inf
            betamax = np.inf
            Di = D[i, np.concatenate((np.r_[0:i], np.r_[i+1:n]))]
            (H, thisP) = Hbeta(Di, beta[i])
    
            # Evaluate whether the perplexity is within tolerance
            Hdiff = H - logU
            tries = 0
            while np.abs(Hdiff) > tol and tries < 50:
    
                # If not, increase or decrease precision
                if Hdiff > 0:
                    betamin = beta[i].copy()
                    if betamax == np.inf or betamax == -np.inf:
                        beta[i] = beta[i] * 2.
                    else:
                        beta[i] = (beta[i] + betamax) / 2.
                else:
                    betamax = beta[i].copy()
                    if betamin == np.inf or betamin == -np.inf:
                        beta[i] = beta[i] / 2.
                    else:
                        beta[i] = (beta[i] + betamin) / 2.
    
                # Recompute the values
                (H, thisP) = Hbeta(Di, beta[i])
                Hdiff = H - logU
                tries += 1
    
            # Set the final row of P
            P[i, np.concatenate((np.r_[0:i], np.r_[i+1:n]))] = thisP
    
        return P
    
    
    
    distances = mat.astype(np.float32, copy=False)
    conditional_P = x2p(distances, tol, perp)
    P = conditional_P + conditional_P.T
    sum_P = np.maximum(np.sum(P), MACHINE_EPSILON)
    P = np.maximum(squareform(P) / sum_P, MACHINE_EPSILON)
    # P=_joint_probabilities(mat, perp)
    degrees_of_freedom = max(n_dim - 1, 1)
    n_samples=len(mat)
    X_embedded = np.random.normal(size=(n_samples, n_dim))
    
    # (it, n_iter, n_iter_check=1, momentum=0.8)
    
    params = X_embedded.ravel().copy()
    update = np.zeros_like(params)
    gains = np.ones_like(params)
    error = np.finfo(np.float).max
    
    
    for i in range(n_iter):

        error, grad = kl_divergence(params, P,
                                    degrees_of_freedom,
                                    n_samples, n_dim)
        grad_norm = np.linalg.norm(grad)

        inc = update * grad < 0.0
        dec = np.invert(inc)
        gains[inc] += 0.2
        gains[dec] *= 0.8
        np.clip(gains, 0.01, np.inf, out=gains)
        grad *= gains
        update = momentum * update - rate * grad
        params += update
        
        if (i+1)%100==0:
            print('t-SNE iteration {0}: KL-divergence is {1:.2e}.'.format(i+1,error))
            
        if grad_norm <= tol:
            break
    
    Y=params.reshape(n_samples, n_dim)
    return Y
    
    
    


# SpectralEmbedding :----------------------------------
# Takes in a graph adjecent matrix(instead of a distance matrix)

def SpectralEmbedding(graph_matrix, dim):
    """
    Spectral embedding.
    
    
    Parameters
    ----------
    graph_matrix : numpy.ndarray
        Adjecent matrix of the graph of points.
        
    dim : int
        Dimension of embedding space.


    Returns
    -------
    Y : numpy.ndarray
        Embedding points.

    """

    L=csgraph.laplacian(graph_matrix, normed=True)
    eig_vals, eig_vecs = np.linalg.eig(L)
    eig_vecs= eig_vecs[:, (eig_vals.argsort())]

    Y=eig_vecs[:,:dim]
    return Y


'''
# UMAP algorithm :-----------------------------------------

def LocalFuzzySimplicialSet(dist,x, n):

    knn_dist=np.sort(dist)[1:n+1]
    index=np.argsort(dist)[1:n+1]
    rho=knn_dist[0]

    # Function to solve
    def f(s):
        return(sum(np.exp(-(knn_dist-rho)/s))-np.log2(n))

    #Binary search for sigma such that sum of exp(-(knn_dists-rho)/sigma) = log2(n)
    if f(pow(.1,100))<=0:
        sigma=opt.bisect(f,pow(.1,100),3*knn_dist[n-1])
    else:
        sigma=1

    fs_set=[]

    for i in index:
        d=max(0,dist[i]-rho)/sigma
        fs_set.append([x,i,np.exp(-d)])

    return(fs_set)


def OptimizeEmbedding(fs_set,
                      Y,
                      min_dist,
                      n_epochs,
                      alpha=.1,
                      n_neg_samples=10):
    """

    """

    initial_alpha=alpha

    # Find a and b
    def curve(x, a, b):
        return 1 / (1 + a * x ** (2 * b))
    xv = np.linspace(0, 3, 300)
    yv = np.zeros(xv.shape)
    yv[xv < min_dist] = 1.0
    yv[xv >= min_dist] = np.exp(-(xv[xv >= min_dist] - min_dist))
    params, covar = curve_fit(curve, xv, yv)
    a=params[0]
    b=params[1]


    for n in range(n_epochs):
        alpha = initial_alpha * (1.0 - (n / n_epochs))
        for elem in fs_set:
            if np.random.rand()<elem[2]:
                dist=dis.pdist([Y[elem[0]],Y[elem[1]]])
                step = alpha*(
                    1+a*pow(dist**2,b))*(
                    -1)*pow((1+a*pow(dist**2,b)),-2
                    )*a*b*pow(dist**2,b-1)*(
                    Y[elem[0]]-Y[elem[1]])

                if np.log(curve(dist,a,b)) < np.log(curve(dis.pdist([Y[elem[0]]+step,Y[elem[1]]]),a,b)):
                    Y[elem[0]]+=step

                for i in range(n_neg_samples):
                    c = np.random.randint(len(Y)-1)
                    if c>=elem[0]:
                        c+=1
                    dist=dis.pdist([Y[elem[0]],Y[c]])
                    step = alpha*1/(
                        1-1/(1+a*pow(dist**2,b))
                        )*pow((1+a*pow(dist**2,b)),-2
                        )*a*b*pow(dist**2,b-1)*(
                        Y[elem[0]]-Y[c])/1000

                    if np.log(1-curve(dist,a,b)) < np.log(1-curve(dis.pdist([Y[elem[0]]+step,Y[elem[1]]]),a,b)):
                        Y[elem[0]]+=step

    return Y


def UMAP(mat,
         dim=2,
         n=5,
         min_dist=1,
         n_epochs=10,
         alpha=1.0,
         n_neg_samples=0):
    """

    Parameters
    ----------
    mat : numpy.ndarray
        distance matrix.
    dim : int, optional
        Dimension of embedding space. The default is 2.
    n : int, optional
        neighborhood size. The default is 5.
    min_dist : float, optional
        DESCRIPTION. The default is 1.
    n_epochs : int, optional
        DESCRIPTION. The default is 10.
    alpha : float, optional
        DESCRIPTION. The default is 1.0.
    n_neg_samples : int, optional
        DESCRIPTION. The default is 0.

    Returns
    -------
    Y :  numpy.ndarray
        embedding points.

    """

    fs_set=[]
    for i in range(len(mat)):
        fs_set=fs_set+LocalFuzzySimplicialSet(mat[i], i, n)

    G=np.zeros(shape=(len(mat),len(mat))) # graph matrix
    for x in fs_set:
        G[x[0],x[1]]=x[2] # weighted adjacency matrix
    G=(G+G.T)/2

    Y=SpectralEmbedding(G, dim)
    Y=OptimizeEmbedding(fs_set,
                        Y,
                        min_dist,
                        n_epochs,
                        alpha=alpha,
                        n_neg_samples=n_neg_samples)
    return(Y)

'''


# PHATE algorithm :------------------------------

def nMDS(dist_mat,
         init,
         momentum=.1, iteration=1000):
    """
    Non-metric multi-dimensional scaling, nMDS.
    
    
    Parameters
    ----------
    dist_mat : array
        Distance matrix of the points.
        
    init : array
        Initial embedding.
        
    momentum : float
        Dimension of the embedding space. 
        The default is 2.
    
    iteration : int
        Number of iteration of gradient descent.
        The default is 1000.

    Returns
    -------
    Y : array
        Embedding points.

    """
    # R is Dt
    R=dist_mat
    Y=init

    def cost_fun(R, Y):
        return(np.sum((R - dis.squareform(dis.pdist(Y)))**2)*.5)

    def cost_grad(R,Y):
        D=dis.squareform(dis.pdist(Y))
        K=(R-D)/(D+1e-10)
        G=np.zeros(Y.shape)
        for i in range(len(Y)):
            dyij=-(Y-Y[i])
            G[i]=np.sum(dyij.T*K[:,i],axis=1)
        return(-2*G)


    for i in range(iteration):
        step=-momentum*cost_grad(R,Y)
        if cost_fun(R,Y)>cost_fun(R,Y+step):
            Y=Y+step
        momentum-=momentum/iteration

    return(Y)


def VNE(P, t):
    """
    
    Von Neumann Entropy.
    

    Parameters
    ----------
    P : numpy.ndarray
        The density matrix.
        
    t : int
        Number of time stages to calculate VNE.


    Returns
    -------
    Ht : float
        Von Neumann Entropy.

    """

    eta=np.linalg.eigvals(P)
    eta=pow(eta,t)
    eta=eta/sum(eta)
    Ht=-sum(eta*np.log(eta))
    return Ht


def PHATE(mat, dim=2, k=5, a=1, gamma=1, t_max=100, momentum=.1, iteration=1000):
    """
    Potential of Heat-diffusion for Affinity-based Trajectory Embedding,
    PHATE.
    

    Parameters
    ----------
    mat : numpy.ndarray
        Distance matrix.
        
    dim : int, optional
        Desired embedding dimension.
        The default is 2.
        
    k : int, optional
        Neighborhood size.
        The default is 5.
        
    a : float, optional
        Locality scale.
        The default is 1.
        
    gamma : float, optional, must in [-1, 1]
        Informational distance constant between -1 and 1.
        The default is 1.
        
    t_max : int, optional
        Maximum time scale for diffusion.
        The default is 100.
    
    momentum : float, optional
        Momentum of gradient descent algorithm.
        The default is .1.
        
    iteration : TYPE, optional
        Number of iteration in gradient descent.
        The default is 1000.

    Returns
    -------
    Y : numpy.ndarray
        The PHATE embedding matrix.

    """

    epsilon=np.sort(mat, axis=0)[k]

    # Local affinity matrix from distance matrix and epsilon
    K=mat
    for i in range(len(mat)):
        for j in range(len(mat[0])):
            K[i][j]=np.exp(-pow(K[i][j]/epsilon[i],a))/2 +np.exp(-pow(K[i][j]/epsilon[j],a))/2

    # normalize K to form a Markov transition matrix
    # (diffusion operator P)
    P=K/np.sum(K,axis=0)
    P=P.transpose()

    # Find t(time scale via Von Neumann Entropy)
    t=1
    c=(VNE(P,1)-VNE(P,t_max))/t_max
    while (VNE(P,t)-VNE(P,t+1))>c:
        t+=1


    # Diffuse P for t time steps to obtain Pt
    Pt = np.linalg.matrix_power(P, t)

    # Compute potential representations Ut
    if gamma == 1:
        # handling small values
        Pt = Pt + 1e-7
        Ut = -1 * np.log(Pt)
    elif gamma == -1:
        Ut = Pt
    else:
        c = (1 - gamma) / 2
        Ut = (Pt ** c) / c


    # compute potential distance matrix from Ut
    Dt=mat
    for i in range(len(Ut)):
        for j in range(len(Ut[0])):
            Dt[i][j]=np.linalg.norm(Ut[j]-Ut[i])

    # apply classical MDS to Dt
    Y=MDS(Dt,dim)
    
    
    
    # apply non-metric MDS to Dt with Y as an initialization
    Y=nMDS(Dt,Y,momentum=.1, iteration=1000)

    return Y
```
