## Supplementary material for "scHiCTools: a computational toolbox for analyzing single-cell Hi-C data": S4: reproducibility.py

```
import numpy as np
from time import time
from scipy.stats import zscore
import scipy.spatial.distance as dis


def pairwise_distances(all_strata, similarity_method,
                       print_time=False, sigma=.5, window_size=10):
    """
    
    Find the pairwise distace using different similarity method, 
    and return a distance matrix.
    

    Parameters
    ----------
    all_strata : numpy.ndarray
        An array contained the strata of a specific chromesome for cells.
        
    similarity_method : str
        The similarity method used to calculate distance matrix.
        Now support 'innerproduct', 'hicrep', 'selfish'.
        
    print_time : bool, optional
        Whether to return the run time of similarity.
        The default is False.
        
    sigma : float, optional
        Parameter for method 'Selfish'.
        Sigma for the Gaussian kernel used in Selfish.
        The default is .5.
        
    window_size : int, optional
        Parameter for method 'Selfish'.
        Length of every window used in Selfish.
        The default is 10.
        

    Returns
    -------
    numpy.ndarray
        Distance matrix.

    """
    
    method = similarity_method.lower()
    t0 = time()

    if method in ['inner_product', 'innerproduct']:
        print(' Calculating z-scores...')
        zscores = []
        for stratum in all_strata:
            z = zscore(stratum, axis=1)
            # print(np.sum(np.isnan(z)))
            z[np.isnan(z)] = 0
            zscores.append(z)
        zscores = np.concatenate(zscores, axis=1)
        t1 = time()
        print(' Calculating inner product...')
        inner = zscores.dot(zscores.T) / zscores.shape[1]
        # print(np.max(inner), np.min(inner))
        inner[inner > 1] = 1
        inner[inner < -1] = -1
        distance_mat = np.sqrt(2 - 2 * inner)
        t2 = time()

    elif method == 'hicrep':
        print(' Calculating means and stds...')
        n_cells, n_bins = all_strata[0].shape
        n_strata = len(all_strata)
        weighted_std = np.zeros((n_cells, n_strata))
        for i, stratum in enumerate(all_strata):
            mean, std = np.mean(stratum, axis=1), np.std(stratum, axis=1)
            weighted_std[:, i] = np.sqrt(n_bins - i) * std
            all_strata[i] = all_strata[i] - mean[:, None]  # subtract a value for each row
        scores = np.concatenate(all_strata, axis=1)
        t1 = time()

        print(' Calculating fastHiCRep score...')
        inner = scores.dot(scores.T) / (weighted_std.dot(weighted_std.T) + 1e-8)  # avoid 0 / 0
        inner[inner > 1] = 1
        inner[inner < -1] = -1
        distance_mat = np.sqrt(2 - 2 * inner)
        t2 = time()

    elif method == 'old_hicrep':
        n_cells, n_bins = all_strata[0].shape
        similarity = np.ones((n_cells, n_cells))
        for i in range(n_cells):
            for j in range(i + 1, n_cells):
                corrs, weights = [], []
                for stratum in all_strata:
                    s1, s2 = stratum[i, :], stratum[j, :]
                    if np.var(s1)==0 or np.var(s2)==0:
                        weights.append(0)
                        corrs.append(0)
                    else:
                        # zero_pos = [k for k in range(len(s1)) if s1[k] == 0 and s2[k] == 0]
                        # s1, s2 = np.delete(s1, zero_pos), np.delete(s2, zero_pos)
                        weights.append(len(s1) * np.std(s1) * np.std(s2))
                        corrs.append(np.corrcoef(s1, s2)[0, 1])
                corrs=np.nan_to_num(corrs)
                s = np.inner(corrs, weights) / (np.sum(weights))
                similarity[i, j] = s
                similarity[j, i] = s
        t1 = time()
        distance_mat = np.sqrt(2 - 2 * similarity)
        t2 = time()


    elif method == 'selfish':
        n_cells, n_bins = all_strata[0].shape
        n_strata = len(all_strata), 
        # window_size = n_bins // (n_windows + 1) * 2
        # window_size=kwargs.pop('window_size', 10)
        n_windows=n_bins//window_size
        
        # if window_size > n_strata:
        #     print('Warning: first {0} strata cannot cover the full region for calculating map similarity.'.format(n_strata),
        #           'Required: {0} strata'.format(window_size),
        #           'Use zeros to fill the missing values.')
        print(' Calculating summation of sliding windows...')
        all_windows = np.zeros((n_cells, n_windows))
        for i, stratum in enumerate(all_strata):
            for j in range(n_windows):
                all_windows[:, j] += np.sum(stratum[:, j * window_size: (j + 1) * window_size - i],axis=1)
        t1 = time()

        print(' Pairwisely compare the windows...')
        fingerprints = np.zeros((n_cells, n_windows * (n_windows-1)//2))
        k=0
        for i in range(n_windows):
            for j in range(n_windows-i-1):
                fingerprints[:,k]=all_windows[:,i]>all_windows[:,j]
                k+=1
        
        # for idx in range(n_cells):
        #     for i, x in enumerate(all_windows[idx]):
        #         for j, y in enumerate(all_windows[idx]):
        #             if x > y:
        #                 fingerprints[idx, i * n_windows + j] = 1
        # print(fingerprints)
        # print(np.sum(fingerprints, axis=1))
        distance = dis.pdist(fingerprints, 'euclidean')
        distance = dis.squareform(distance)
        similarity = np.exp(- sigma * distance)
        distance_mat = np.sqrt(2 - 2 * similarity)
        t2 = time()
        
    else:
        raise ValueError('Method {0} not supported. Only "inner_product", "HiCRep", "old_hicrep" and "Selfish".'.format(method))
    
    if print_time:
        print('Time 1:', t1 - t0)
        print('Time 2:', t2 - t1)
        return distance_mat, t1 - t0, t2 - t1
    else:
        return distance_mat
```
