## Supplementary material for "scHiCTools: a computational toolbox for analyzing single-cell Hi-C data": S4: ContactMaps.py

```
import numpy as np
import pandas as pd
from copy import deepcopy
from scipy.sparse import coo_matrix
from .load_hic_file import get_chromosome_lengths, load_HiC
from ..embedding import pairwise_distances, MDS, tSNE, PHATE, SpectralEmbedding, PCA
from .processing_utils import matrix_operation
# from ..analysis import scatter
from ..analysis import kmeans, spectral_clustering, HAC
import matplotlib.pyplot as plt


class scHiCs:
    def __init__(self, list_of_files, reference_genome, resolution,
                 adjust_resolution=True, sparse=False, chromosomes='all',
                 format='customized', keep_n_strata=10, store_full_map=False,
                 operations=None, header=0, customized_format=None,
                 map_filter=0., gzip=False, **kwargs):
        """

        Parameters
        ----------
        list_of_files : list
            List of HiC file paths.
            
        reference_genome : str or dict
            Now supporting 'mm9', 'mm10', 'hg19', 'hg38',
            if using other references,you can simply provide the chromosome name 
            and corresponding size (bp) with a dictionary in Python.
            e.g. {'chr1': 150000000, 'chr2': 130000000, 'chr3': 200000000}
            
        resolution : int
            The resolution to separate genome into bins. If using .hic file format,
            the given resolution must match with the resolutions in .hic file.
            
        adjust_resolution : bool, optional
            Whether to adjust resolution for input file. 
            Sometimes the input file is already in the proper resolution
            (e.g. position 3000000 has already been changed to 6 in 500kb resolution),
            then you can set `adjust_resolution=False`.  The default is True.
            
        sparse : bool, optional
            Whether to use sparse matrix to store (only effective when max_distance=None). The default is False.
            
        chromosomes : list or str, optional
            Chromosomes to use, 
            eg. ['chr1', 'chr2'], or just 'except Y', 'except XY','all',
            which means chr 1-19 + XY for mouse and chr 1-22 + XY for human.
            The default is 'all'.
            
        format : str, optional
            HiC files' format.
            e.g., '.hic', 'customized', '.cool'. The default is 'customized'.
            
        keep_n_strata : int, optional
            Only consider contacts within this genomic distance. 
            If `None`, it will store full matrices in numpy matrix or 
            scipy sparse format, which will use too much memory sometimes.
            The default is 10.
            
        store_full_map : bool, optional
            Whether store contact maps. The default is False.
            
        operations : list, optional
            The methods use for pre-processing or smoothing the maps given in a list.
            The operations will happen in the given order. 
            Operations: 'convolution', 'random_walk', 'network_enhancing'.
            For pre-processing and smoothing operations, sometimes you need additional arguments.
            You can check docstrings for pre-processing and smoothing for more information. 
            The default is None.
        
        header : int, optional
            The number of header line(s). 
            If `header=0`, HiC files do not have header.
            The default is 0.
            
        customized_format : int or list, optional
            Format for each line. The default is None.
            
        map_filter : float, optional
            The threshold to filter some reads by map quality. 
            The default is 0..
            
        gzip : bool, optional
            If the HiC files are zip files. 
            If `True`, the HiC files are zip files. 
            The default is False.
            
        **kwargs : 
            Other arguments specify smoothing methods passed to function.
            See `scHiCTools.load.processing_utils.matrix_operation` function.


        Returns
        -------
        None.
        

        """

        self.resolution = resolution
        self.chromosomes, self.chromosome_lengths = get_chromosome_lengths(reference_genome, chromosomes, resolution)
        self.num_of_cells = len(list_of_files)
        self.sparse = sparse
        self.keep_n_strata = keep_n_strata
        self.contacts=np.array([0]*len(list_of_files))
        self.short_range=np.array([0]*len(list_of_files))
        self.mitotic=np.array([0]*len(list_of_files))
        self.files=list_of_files
        self.strata = {
            ch: [np.zeros((self.num_of_cells, self.chromosome_lengths[ch] - i)) for i in range(keep_n_strata)]
            for ch in self.chromosomes} if keep_n_strata else None
        self.full_maps = None
        self.similarity_method=None
        self.distance=None

        assert keep_n_strata is not None or store_full_map is True

        if not store_full_map:
            self.full_maps = None
        elif sparse:
            self.full_maps = {ch: [None] * self.num_of_cells for ch in self.chromosomes}
        else:
            self.full_maps = {
                ch: np.zeros((self.num_of_cells, self.chromosome_lengths[ch], self.chromosome_lengths[ch]))
                for ch in self.chromosomes}

        for idx, file in enumerate(list_of_files):
            print('Processing {0} out of {1} files: {2}'.format(idx+1,len(list_of_files),file))

            for ch in self.chromosomes:
                if ('ch' in ch) and ('chr' not in ch):
                    ch=ch.replace("ch", "chr")
                mat, strata = load_HiC(
                    file, genome_length=self.chromosome_lengths,
                    format=format, custom_format=customized_format,
                    header=header, chromosome=ch, resolution=resolution,
                    resolution_adjust=adjust_resolution,
                    map_filter=map_filter, sparse=sparse, gzip=gzip,
                    keep_n_strata=keep_n_strata, operations=operations,
                    **kwargs)
                
                self.contacts[idx]+=np.sum(mat)/2+ np.trace(mat)/2

                self.short_range[idx]+=sum([np.sum(mat[i,i:i+int(2000000/self.resolution)]) for i in range(len(mat))])
                self.mitotic[idx]+=sum([np.sum(mat[i,i+int(2000000/self.resolution):i+int(12000000/self.resolution)]) for i in range(len(mat))])


                if store_full_map:
                    self.full_maps[ch][idx] = mat

                if keep_n_strata:
                    # self.strata[ch][idx] = strata
                    for strata_idx, stratum in enumerate(strata):
                        self.strata[ch][strata_idx][idx, :] = stratum

    def cal_strata(self, n_strata):
        """
        
        Alter the number of strata kept in a `scHiCs` object.
        
        
        Parameters
        ----------
        n_strata : int
            Number of strata to keep.

        Returns
        -------
        dict
            Strata of cells.

        """
        
        if self.full_maps is None:
            if self.keep_n_strata <= n_strata:
                print(' Only {0} strata are kept!'.format(self.keep_n_strata))
                return deepcopy(self.strata)
            else:
                return deepcopy({ch: self.strata[ch][:n_strata] for ch in self.chromosomes})
        else:
            if self.keep_n_strata is None:
                new_strata = {
                    ch: [np.zeros((self.num_of_cells, self.chromosome_lengths[ch] - i))
                         for i in range(n_strata)] for ch in self.chromosomes}
                for ch in self.chromosomes:
                    for idx in range(self.num_of_cells):
                        fmap = self.full_maps[ch][idx].toarray() if self.sparse else self.full_maps[ch][idx]
                        for i in range(n_strata):
                            new_strata[ch][i][idx, :] = np.diag(fmap[i:, :-i])
                return new_strata
            elif self.keep_n_strata >= n_strata:
                return deepcopy({ch: self.strata[ch][:n_strata] for ch in self.chromosomes})
            else:
                for ch in self.chromosomes:
                    self.strata[ch] += [(np.zeros(self.num_of_cells, self.chromosome_lengths[ch] - i))
                                        for i in range(self.keep_n_strata, n_strata)]
                    for idx in range(self.num_of_cells):
                        fmap = self.full_maps[ch][idx].toarray() if self.sparse else self.full_maps[ch][idx]
                        for i in range(self.keep_n_strata, n_strata):
                            self.strata[ch][i][idx, :] = np.diag(fmap[i:, :-i])
                return deepcopy(self.strata)
            
            

    def processing(self, operations, **kwargs):
        """
        
        Apply a smoothing method to contact maps.
        Requre the `scHiCs` object to store the full map of contacts maps.
        

        Parameters
        ----------
        operations : str
            The methods use for smoothing the maps.
            Avaliable operations: 'convolution', 'random_walk', 'network_enhancing'.
            
        **kwargs : 
            Other arguments specify smoothing methods passed to function.
            See function `scHiCTools.load.processing_utils.matrix_operation`.
            

        Returns
        -------
        None.

        """
        
        if self.full_maps is None:
            raise ValueError('No full maps stored. Processing is not doable.')
        if self.sparse:
            for ch in self.chromosomes:
                for i, mat in enumerate(self.full_maps[ch]):
                    self.full_maps[ch][i] = coo_matrix(matrix_operation(mat.toarray(), operations, **kwargs))
        else:
            for ch in self.chromosomes:
                for i, mat in enumerate(self.full_maps[ch]):
                    self.full_maps[ch][i, :, :] = matrix_operation(mat, operations, **kwargs)
        # Update the strata
        if self.keep_n_strata is not None:
            for ch in self.chromosomes:
                for i, mat in enumerate(self.full_maps[ch]):
                    for j in range(self.keep_n_strata):
                        self.strata[ch][j][i, :] = np.diag(mat[j:, :len(mat) - j])


    def plot_contacts(self, hist=True, percent=True, 
                      size=1.0, bins=10, color='#1f77b4'):
        """
        
        Generate two plots:
        Histogram of contacts and 
        scatter plot of short-range contacts v.s. contacts at the mitotic band.
        

        Parameters
        ----------
        hist : bool, optional
            Whether to plot Histogram of contacts. 
            If `True`, plot Histogram of contacts. 
            The default is True.
            
        percent : int, optional
            Whether to plot scatter plot of short-range contacts v.s. contacts at the mitotic band.
            If `True`, plot scatter plot of short-range contacts v.s. contacts at the mitotic band.
            The default is True.
            
        size : float, optional
            The point size of scatter plot.
            The default is 1.0.
            
        bins : int, optional
            Number of bins in histogram.
            The default is 10.
            
        color : str, optional
            The color of the plot.
            The default is '#1f77b4'.

        Returns
        -------
        None.

        """

        if hist:
            if percent:
                plt.subplot(1,2,1)

            plt.hist(self.contacts,bins=bins,color=color)
            plt.xlabel("Number of contacts")
            plt.ylabel('Frequency')
            plt.title('Histogram of contacts')

        if percent:
            if hist:
                plt.subplot(1,2,2)

            plt.scatter(self.mitotic*100/self.contacts,self.short_range*100/self.contacts, s=size, c=color)
            plt.xlabel("% Mitotic contacts")
            plt.ylabel("% Short-range contacts")
            plt.title('Short-range contacts v.s. contacts at the mitotic band')


    def select_cells(self, min_n_contacts=0,max_short_range_contact=1):
        """
        
        Select qualify cells based on minimum number of contacts and
        maxium percent of short range contact.
        

        Parameters
        ----------
        min_n_contacts : int, optional
            The threshold of minimum number of contacts in each cell.
            The default is 0.
            
        max_short_range_contact : float, optional
            The threshold of maximum proportion of short range contact in every cell.
            The default is 1.

        Returns
        -------
        list
            Selected files.

        """
        
        files=np.array(self.files)
        selected=np.logical_and(self.short_range/self.contacts<=max_short_range_contact,self.contacts>=min_n_contacts)
        self.num_of_cells=sum(selected)
        self.files=[self.files[i] for i in  range(len(files)) if selected[i]]
        self.contacts=self.contacts[selected]
        self.short_range=self.short_range[selected]
        self.mitotic=self.mitotic[selected]
        if self.strata is not None:
            for ch in self.chromosomes:
                self.strata[ch]=[self.strata[ch][i] for i in np.arange(len(selected))[selected]]
        if self.full_maps is not None:
            for ch in self.chromosomes:
                self.full_maps[ch]=self.full_maps[ch][selected]
        if self.distance is not None:
            self.distance=self.distance[:,selected,:][:,:,selected]
            
        return files[selected]


    def scHiCluster(self,dim=2,n_clusters=4,cutoff=0.8,n_PCs=10,**kwargs):

        """
        
        Embedding and clustering single cells using HiCluster.
        Reference: 
            Zhou J, Ma J, Chen Y, Cheng C, Bao B, Peng J, et al.
            Robust single-cell Hi-C clustering by convolution- and random-walk–based imputation.
            PNAS. 2019 Jul 9;116(28):14011–8. 
        
        
        Parameters
        ----------
        dim : int, optional
            Number of dimension of embedding. The default is 2.
            
        n_clusters : int, optional
            Number of clusters. The default is 4.
            
        cutoff : float, optional
            The cutoff proportion to convert the real contact
            matrix into binary matrix. The default is 0.8.
            
        n_PCs : int, optional
            Number of principal components. The default is 10.
            
        **kwargs :
            Other arguments passed to kmeans.
            See `scHiCTools.analysis.clustering.kmeans` function.

        Returns
        -------
        embeddings : numpy.ndarray
            The embedding of cells using HiCluster.
            
        label : numpy.ndarray
            An array of cell labels clustered by HiCluster.
            
        """

        if self.full_maps is None:
            raise ValueError('No full maps stored. scHiCluster is not doable.')

        X=None
        for ch in self.chromosomes:
            print('HiCluster processing chromosomes {}'.format(ch))
            A=self.full_maps[ch].copy()
            if len(A.shape)==3:
                n=A.shape[1]*A.shape[2]
                A.shape=(A.shape[0],n)
            A=np.quantile(A,cutoff,axis=1)<np.transpose(A)
            A = PCA(A.T,n_PCs)
            if X is None:
                X=A
            else:
                X=np.append(X, A, axis=1)

        X=PCA(X,n_PCs)
        label=kmeans(X,n_clusters,kwargs.pop('weights',None),kwargs.pop('iteration',1000))

        return X[:,:dim], label


    def learn_embedding(self, similarity_method, embedding_method,
                        dim=2, aggregation='median', n_strata=None, return_distance=False, print_time=False,
                        **kwargs):
        """
        
        Function to find a low-dimensional embedding for cells.
        

        Parameters
        ----------
        similarity_method : str
            The method used to calculate similarity matrix.
            Now support 'inner_product', 'HiCRep' and 'Selfish'.
            
        embedding_method : str
            The method used to project cells into lower-dimensional space.
            Now support 'MDS', 'tSNE', 'phate', 'spectral_embedding'.
            
        dim : int, optional
            Dimension of the embedding space.
            The default is 2.
            
        aggregation : str, optional
            Method to find the distance matrix based on distance matrices of chromesomes.
            Must be 'mean' or 'median'.
            The default is 'median'.
            
        n_strata : int, optional
            Number of strata used in calculation.
            The default is None.
            
        return_distance : bool, optional
            Whether to return the distance matrix of cells.
            If True, return (embeddings, distance_matrix);
            if False, only return embeddings. 
            The default is False.
            
        print_time : bool, optional
            Whether to print process time. The default is False.
            
        **kwargs :
            Including two arguments for Selfish
            (see funciton `pairwise_distances`):\
            `n_windows`: number of Selfish windows\
            `sigma`: sigma in the Gaussian-like kernel\
            and some arguments specify different embedding method 
            (see functions in `scHiCTools.embedding.embedding`).
            

        Returns
        -------
        embeddings: numpy.ndarray
            The embedding of cells in lower-dimensional space.
        
        final_distance: numpy.ndarray, optional
            The pairwise distance calculated.
        
        """
        
        if self.distance is None or self.similarity_method!=similarity_method:
            self.similarity_method=similarity_method
            distance_matrices = []
            assert embedding_method.lower() in ['mds', 'tsne', 'umap', 'phate', 'spectral_embedding']
            assert n_strata is not None or self.keep_n_strata is not None
            n_strata = n_strata if n_strata is not None else self.keep_n_strata
            new_strata = self.cal_strata(n_strata)
            if print_time:
                time1=0
                time2=0
                for ch in self.chromosomes:
                    print(ch)
                    distance_mat,t1,t2 = pairwise_distances(new_strata[ch], similarity_method, print_time, kwargs.get('sigma',.5), kwargs.get('window_size',10))
                    time1=time1+t1
                    time2=time2+t2
                    distance_matrices.append(distance_mat)
                print('Sum of time 1:', time1)
                print('Sum of time 2:', time2)
            else:
                for ch in self.chromosomes:
                    print(ch)
                    distance_mat = pairwise_distances(new_strata[ch],
                                   similarity_method,
                                   print_time,
                                   kwargs.get('sigma',.5),
                                   kwargs.get('window_size',10))
                    distance_matrices.append(distance_mat)
            self.distance = np.array(distance_matrices)

        if aggregation == 'mean':
            final_distance = np.mean(self.distance, axis=0)
        elif aggregation == 'median':
            final_distance = np.median(self.distance, axis=0)
        else:
            raise ValueError('Aggregation method {0} not supported. Only "mean" or "median".'.format(aggregation))

        np.fill_diagonal(final_distance, 0)

        embedding_method = embedding_method.lower()
        if embedding_method == 'mds':
            embeddings = MDS(final_distance, dim)
        elif embedding_method == 'tsne':
            embeddings = tSNE(final_distance, dim, 
                              kwargs.pop('perp',30),
                              kwargs.pop('iteration',1000),
                              kwargs.pop('momentum', 0.5),
                              kwargs.pop('rate', 200),
                              kwargs.pop('tol',1e-5))
        # elif embedding_method == 'umap':
        #     embeddings = UMAP(final_distance, dim, 
        #                       kwargs.pop('n',5),
        #                       kwargs.pop('min_dist',1),
        #                       kwargs.pop('n_epochs',10),
        #                       kwargs.pop('alpha',1),
        #                       kwargs.pop('n_neg_samples',0))
        elif embedding_method == 'phate':
            embeddings = PHATE(final_distance, dim,
                               kwargs.pop('k',5), 
                               kwargs.pop('a',1),
                               kwargs.pop('gamma',1),
                               kwargs.pop('t_max',100),
                               kwargs.pop('momentum',.1),
                               kwargs.pop('iteration',1000))
        elif embedding_method == 'spectral_embedding':
            graph=np.exp(-np.square(final_distance)/np.mean(final_distance**2))
            graph = graph-np.diag(graph.diagonal())
            embeddings = SpectralEmbedding(graph, dim)
        else:
            raise ValueError('Embedding method {0} not supported. '.format(embedding_method))
            
        if return_distance:
            return embeddings, final_distance
        else:
            return embeddings


    def clustering(self,
                   n_clusters,
                   clustering_method,
                   similarity_method,
                   aggregation='median',
                   n_strata=None,
                   print_time=False,
                   **kwargs):
        """
        
        Parameters
        ----------
        n_clusters : int
            Number of clusters.
            
        clustering_method : str
            Clustering method in 'kmeans', 'spectral_clustering' or 'HAC'(hierarchical agglomerative clustering).
            
        similarity_method : str
            Reproducibility measure.
            Value in ‘InnerProduct’, ‘HiCRep’ or ‘Selfish’.
            
        aggregation : str, optional
             Method to aggregate different chromosomes.
             Value is either 'mean' or 'median'.
             The default is 'median'.
             
        n_strata : int or None, optional
            Only consider contacts within this genomic distance.
            If it is None, it will use the all strata kept from previous loading process.
            The default is None.
            
        print_time : bool, optional
            Whether to print the processing time. The default is False.
            
        **kwargs :
            Other arguments pass to function `scHiCTools.embedding.reproducibility.pairwise_distances `,
            and the clustering function in `scHiCTools.analysis.clustering`.


        Returns
        -------
        label : numpy.ndarray
            An array of cell labels clustered.
            
        """
        if self.distance is None or self.similarity_method!=similarity_method:
            self.similarity_method=similarity_method
            distance_matrices = []
            assert n_strata is not None or self.keep_n_strata is not None
            n_strata = n_strata if n_strata is not None else self.keep_n_strata
            new_strata = self.cal_strata(n_strata)

            for ch in self.chromosomes:
                print(ch)
                distance_mat = pairwise_distances(new_strata[ch],
                                              similarity_method,
                                              print_time,
                                              kwargs.get('sigma',.5),
                                              kwargs.get('window_size',10))
                distance_matrices.append(distance_mat)
            self.distance = np.array(distance_matrices)

        if aggregation == 'mean':
            final_distance = np.mean(self.distance, axis=0)
        elif aggregation == 'median':
            final_distance = np.median(self.distance, axis=0)
        else:
            raise ValueError('Aggregation method {0} not supported. Only "mean" or "median".'.format(aggregation))

        np.fill_diagonal(final_distance, 0)
        
        clustering_method=clustering_method.lower()
        if clustering_method=='kmeans':
            embeddings = MDS(final_distance, n_clusters)
            label=kmeans(embeddings,
                         k=n_clusters,
                         **kwargs)
        elif clustering_method=='spectral_clustering':
            label=spectral_clustering(final_distance,
                                      data_type='distance_matrix',
                                      n_clusters=n_clusters,
                                      **kwargs)
        elif clustering_method=='hac':
            label=HAC(final_distance,
                      'distance_matrix',
                      n_clusters,
                      kwargs.pop('method','centroid'))
        else:
            raise ValueError('Embedding method {0} not supported. '.format(clustering_method))

        return label
```
