## Supplementary material for "scHiCTools: a computational toolbox for analyzing single-cell Hi-C data": S4: cool.py

```
# -*- coding: utf-8 -*-
# Adapted from https://github.com/mirnylab/cooler
from .cooler_api import Cooler, parse_region, region_to_extent, annotate
import numpy as np
import pandas as pd
import sys


def _comes_before(a0, a1, b0, b1, strict=False):
    if a0 < b0:
        return a1 <= b0 if strict else a1 <= b1
    return False


def _contains(a0, a1, b0, b1, strict=False):
    if a0 > b0 or a1 < b1:
        return False
    if strict and (a0 == b0 or a1 == b1):
        return False
    return a0 <= b0 and a1 >= b1


def _prune_partition(edges, step):
    edges = np.asarray(edges)
    cumlen = np.r_[0, np.cumsum(np.diff(edges))]
    cuts = [step * i for i in range(0, int(np.ceil(cumlen[-1] / step)))]
    cuts.append(cumlen[-1])
    return np.unique(np.searchsorted(cumlen, cuts))


class CSRReader(object):
    def __init__(self, h5, field, chunksize):
        self.h5 = h5
        self.field = field
        self.chunksize = chunksize
        self.bin1_selector = h5["pixels"]["bin1_id"]
        self.bin2_selector = h5["pixels"]["bin2_id"]
        self.data_selector = h5["pixels"][field]
        self.offset_selector = h5["indexes"]["bin1_offset"]

    def __call__(self, i0, i1, j0, j1, transpose=False):
        isempty = True

        bin1_selector = self.bin1_selector
        bin2_selector = self.bin2_selector
        data_selector = self.data_selector
        chunksize = self.chunksize

        if (i1 - i0 > 0) or (j1 - j0 > 0):

            # coarsegrain the offsets to extract a big chunk of rows at a time
            offsets = self.offset_selector[i0 : i1 + 1]
            which_offsets = _prune_partition(offsets, chunksize)

            for o0, o1 in zip(which_offsets[:-1], which_offsets[1:]):

                # extract a chunk of rows
                slc = slice(offsets[o0], offsets[o1])
                bin2_extracted = bin2_selector[slc]
                data_extracted = data_selector[slc]

                i, j, v = [], [], []

                # filter each row
                delta = offsets[o0]
                for o in range(o0, o1):
                    # correct the offsets
                    lo = offsets[o] - delta
                    hi = offsets[o + 1] - delta

                    # this row
                    bin2 = bin2_extracted[lo:hi]

                    # filter for the range of j values we want
                    mask = (bin2 >= j0) & (bin2 < j1)
                    cols = bin2[mask]

                    # apply same mask for data
                    data = data_extracted[lo:hi][mask]

                    # shortcut for row data
                    rows = np.full(len(cols), i0 + o, dtype=bin1_selector.dtype)

                    i.append(rows)
                    j.append(cols)
                    v.append(data)

                if len(i):
                    isempty = False
                    i = np.concatenate(i, axis=0)
                    j = np.concatenate(j, axis=0)
                    v = np.concatenate(v, axis=0)
                    if transpose:
                        i, j = j, i
                    yield i, j, v

        if isempty:
            i = np.array([], dtype=bin1_selector.dtype)
            j = np.array([], dtype=bin2_selector.dtype)
            v = np.array([], dtype=data_selector.dtype)
            if transpose:
                i, j = j, i
            yield i, j, v


def query2d(triu_reader, i0, i1, j0, j1, duplex):
    # symmetric query
    if (i0, i1) == (j0, j1):
        for i, j, v in triu_reader(i0, i1, i0, i1):
            if duplex:
                nodiag = i != j
                i, j, v = np.r_[i, j[nodiag]], np.r_[j, i[nodiag]], np.r_[v, v[nodiag]]
            yield i, j, v

    # asymmetric query
    else:
        transpose = False
        if j0 < i0 or (i0 == j0 and i1 < j1):
            i0, i1, j0, j1 = j0, j1, i0, i1
            transpose = True

        # non-overlapping
        if _comes_before(i0, i1, j0, j1, strict=True):
            for i, j, v in triu_reader(i0, i1, j0, j1, transpose):
                yield i, j, v

        # partially overlapping
        elif _comes_before(i0, i1, j0, j1):
            for i, j, v in triu_reader(i0, j0, j0, i1, transpose):
                yield i, j, v
            for i, j, v in triu_reader(j0, i1, j0, i1, transpose):
                if duplex:
                    nodiag = i != j
                    i, j, v = (
                        np.r_[i, j[nodiag]],
                        np.r_[j, i[nodiag]],
                        np.r_[v, v[nodiag]],
                    )
                yield i, j, v
            for i, j, v in triu_reader(i0, i1, i1, j1, transpose):
                yield i, j, v

        # nested
        elif _contains(i0, i1, j0, j1):
            for i, j, v in triu_reader(i0, j0, j0, j1, transpose):
                yield i, j, v
            for j, i, v in triu_reader(j0, j1, j0, j1, transpose):
                if duplex:
                    nodiag = i != j
                    i, j, v = (
                        np.r_[i, j[nodiag]],
                        np.r_[j, i[nodiag]],
                        np.r_[v, v[nodiag]],
                    )
                yield i, j, v
            for j, i, v in triu_reader(j0, j1, j1, i1, transpose):
                yield i, j, v

        else:
            raise IndexError("This shouldn't happen")


def make_annotator(bins, balanced, join, annotate, one_based_ids, one_based_starts):
    # print(bins.keys())
    def annotator(chunk):
        if annotate is not None:
            extra_fields = list(annotate)
            try:
                extra_cols = bins[extra_fields]
            except KeyError as e:
                print("Column not found:\n {}".format(e))
                sys.exit(1)
            extra = annotate(
                chunk[["bin1_id", "bin2_id"]], extra_cols, replace=True
            )

        if balanced:
            df = annotate(chunk, bins[["weight"]])
            chunk["balanced"] = df["weight1"] * df["weight2"] * chunk["count"]

        if join:
            chunk = annotate(chunk, bins[["chrom", "start", "end"]], replace=True)

        if annotate is not None:
            chunk = pd.concat([chunk, extra], axis=1)

        if one_based_ids:
            for col in ["bin1_id", "bin2_id"]:
                if col in chunk.columns:
                    chunk[col] += 1

        if one_based_starts:
            for col in ["start1", "start2"]:
                if col in chunk.columns:
                    chunk[col] += 1

        return chunk

    return annotator


def dump(cool_uri, resolution, table='pixels', columns=None, header=False, na_rep='', float_format='g',
         range=None, range2=None,
         matrix=True, balanced=False, join=False, annotate=None, one_based_ids=False,
         one_based_starts=False, chunksize=None):
    """
    Args:
        cool_uri (str): .cool file path
        table (str): "chroms", "bins" or "pixels"


    Dump a cooler's data to a text stream.
    COOL_PATH : Path to COOL file or cooler URI.
    """
    cool_uri = cool_uri + '::resolutions/' + str(resolution)
    c = Cooler(cool_uri)

    # output stream
    # if out is None or out == "-":
    #     f = sys.stdout
    # elif out.endswith(".gz"):
    #     f = gzip.open(out, "wt")
    # else:
    #     f = open(out, "wt")

    # choose the source
    if table == "chroms":
        selector = c.chroms()
        if columns is not None:
            selector = selector[list(columns)]
        chunks = (selector[:],)
    elif table == "bins":
        selector = c.bins()
        if columns is not None:
            selector = selector[list(columns)]
        chunks = (selector[:],)
    else:
        # load all the bins
        bins = c.bins()[:]
        if chunksize is None:
            chunksize = len(bins)

        if balanced and "weight" not in bins.columns:
            print("Balancing weights not found", file=sys.stderr)
            sys.exit(1)

        h5 = c.open("r")
        if range:
            i0, i1 = region_to_extent(
                h5, c._chromids, parse_region(range[3:], c.chromsizes), binsize=c.binsize
            ) #??
            if range2 is not None:
                j0, j1 = region_to_extent(
                    h5,
                    c._chromids,
                    parse_region(range2[3:], c.chromsizes), #??
                    binsize=c.binsize,
                )
            else:
                j0, j1 = i0, i1

            triu_reader = CSRReader(h5, "count", chunksize)
            if matrix and c.storage_mode == "symmetric-upper":
                selector = query2d(triu_reader, i0, i1, j0, j1, duplex=True)
            else:
                selector = triu_reader(i0, i1, j0, j1, transpose=False)

            chunks = (
                pd.DataFrame(
                    {"bin1_id": i, "bin2_id": j, "count": v},
                    columns=["bin1_id", "bin2_id", "count"],
                )
                for i, j, v in selector
            )
        else:
            selector = c.pixels()
            if columns is not None:
                selector = selector[list(columns)]
            n = len(selector)
            edges = np.arange(0, n + chunksize, chunksize)
            edges[-1] = n

            if matrix and c.storage_mode == "symmetric-upper":

                def _select(lo, hi):
                    df = selector[lo:hi]
                    dfT = df.copy()
                    dfT["bin1_id"], dfT["bin2_id"] = df["bin2_id"], df["bin1_id"]
                    return pd.concat([df, dfT])

                chunks = (_select(lo, hi) for lo, hi in zip(edges[:-1], edges[1:]))
            else:
                chunks = (selector[lo:hi] for lo, hi in zip(edges[:-1], edges[1:]))

        if balanced or join or annotate:
            annotator = make_annotator(
                bins, balanced, join, annotate, one_based_ids, one_based_starts
            )
            chunks = map(annotator, chunks)

    first = True
    if float_format is not None:
        float_format = "%" + float_format

    for chunk in chunks:
        
        for idx, row in chunk.iterrows():
            if row.loc['bin1_id']<=row.loc['bin2_id']:
                yield row.loc['bin1_id'], row.loc['bin2_id'], row.loc['count']
            
            # break
    #     if first:
    #         if header:
    #             chunk[0:0].to_csv(
    #                 f, sep="\t", index=False, header=True, float_format=float_format
    #             )
    #         first = False
    #
    #     chunk.to_csv(
    #         f,
    #         sep="\t",
    #         index=False,
    #         header=False,
    #         float_format=float_format,
    #         na_rep=na_rep,
    #     )
    #
    # else:
    #     f.flush()


# dump('test.mcool',25000, 'pixels', range='chr1', range2='chr1')
```
