## Supplementary material for "scHiCTools: a computational toolbox for analyzing single-cell Hi-C data": S4: cooler_api.py

```
# Adapted from https://github.com/mirnylab/cooler
import simplejson as json
import six
import os
import re
from contextlib import contextmanager
from pandas.api.types import is_integer_dtype
from scipy.sparse import coo_matrix
import numpy as np
import pandas as pd
import h5py


# The 4DN data portal and hic2cool store these weight vectors in divisive form
_4DN_DIVISIVE_WEIGHTS = {"KR", "VC", "VC_SQRT"}


@contextmanager
def open_hdf5(fp, mode="r", *args, **kwargs):
    """
    Context manager like ``h5py.File`` but accepts already open HDF5 file
    handles which do not get closed on teardown.
    Parameters
    ----------
    fp : str or ``h5py.File`` object
        If an open file object is provided, it passes through unchanged,
        provided that the requested mode is compatible.
        If a filepath is passed, the context manager will close the file on
        tear down.
    mode : str
        * r        Readonly, file must exist
        * r+       Read/write, file must exist
        * a        Read/write if exists, create otherwise
        * w        Truncate if exists, create otherwise
        * w- or x  Fail if exists, create otherwise
    """
    if isinstance(fp, six.string_types):
        own_fh = True
        fh = h5py.File(fp, mode, *args, **kwargs)
    else:
        own_fh = False
        if mode == "r" and fp.file.mode == "r+":
            # warnings.warn("File object provided is writeable but intent is read-only")
            pass
        elif mode in ("r+", "a") and fp.file.mode == "r":
            raise ValueError("File object provided is not writeable")
        elif mode == "w":
            raise ValueError("Cannot truncate open file")
        elif mode in ("w-", "x"):
            raise ValueError("File exists")
        fh = fp
    try:
        yield fh
    finally:
        if own_fh:
            fh.close()


class closing_hdf5(h5py.Group):
    def __init__(self, grp):
        super(closing_hdf5, self).__init__(grp.id)

    def __enter__(self):
        return self

    def __exit__(self, *exc_info):
        return self.file.close()

    def close(self):
        self.file.close()


class TreeNode(object):
    def __init__(self, obj, depth=0, level=None):
        self.obj = obj
        self.depth = depth
        self.level = level

    def get_type(self):
        return type(self.obj).__name__

    def get_children(self):
        if hasattr(self.obj, "values"):
            if self.level is None or self.depth < self.level:
                depth = self.depth + 1
                children = self.obj.values()
                return [
                    self.__class__(o, depth=depth, level=self.level) for o in children
                ]
        return []

    def get_text(self):
        name = self.obj.name.split("/")[-1] or "/"
        if hasattr(self.obj, "shape"):
            name += " {} {}".format(self.obj.shape, self.obj.dtype)
        return name


MAGIC = u"HDF5::Cooler"
URL = u"https://github.com/mirnylab/cooler"
def _is_cooler(grp):
    fmt = grp.attrs.get("format", None)
    url = grp.attrs.get("format-url", None)
    if fmt == MAGIC or url == URL:
        keys = ("chroms", "bins", "pixels", "indexes")
        if not all(name in grp.keys() for name in keys):
            print("Cooler path {} appears to be corrupt".format(grp.name))
        return True
    return False


def visititems(group, func, level=None):
    """Like :py:method:`h5py.Group.visititems`, but much faster somehow.
    """

    def _visititems(node, func, result=None):
        children = node.get_children()
        if children:
            for child in children:
                result[child.obj.name] = func(child.obj.name, child.obj)
                _visititems(child, func, result)
        return result

    root = TreeNode(group, level=level)
    return _visititems(root, func, {})


def natsort_key(s, _NS_REGEX=re.compile(r"(\d+)", re.U)):
    return tuple([int(x) if x.isdigit() else x for x in _NS_REGEX.split(s) if x])


def natsorted(iterable):
    return sorted(iterable, key=natsort_key)


def list_coolers(filepath):
    """
    List group paths to all cooler data collections in a file.
    Parameters
    ----------
    filepath : str
    Returns
    -------
    list
        Cooler group paths in the file.
    """
    if not h5py.is_hdf5(filepath):
        raise OSError("'{}' is not an HDF5 file.".format(filepath))

    listing = []

    def _check_cooler(pth, grp):
        if _is_cooler(grp):
            listing.append("/" + pth if not pth.startswith("/") else pth)

    with h5py.File(filepath, "r") as f:
        _check_cooler("/", f)
        visititems(f, _check_cooler)

    return natsorted(listing)


def parse_cooler_uri(s):
    """
    Parse a Cooler URI string
    e.g. /path/to/mycoolers.cool::/path/to/cooler
    """
    parts = s.split("::")
    if len(parts) == 1:
        file_path, group_path = parts[0], "/"
    elif len(parts) == 2:
        file_path, group_path = parts
        if not group_path.startswith("/"):
            group_path = "/" + group_path
    else:
        raise ValueError("Invalid Cooler URI string")
    return file_path, group_path


def parse_humanized(s):
    _NUMERIC_RE = re.compile("([0-9,.]+)")
    _, value, unit = _NUMERIC_RE.split(s.replace(",", ""))
    if not len(unit):
        return int(value)

    value = float(value)
    unit = unit.upper().strip()
    if unit in ("K", "KB"):
        value *= 1000
    elif unit in ("M", "MB"):
        value *= 1000000
    elif unit in ("G", "GB"):
        value *= 1000000000
    else:
        raise ValueError("Unknown unit '{}'".format(unit))
    return int(value)


def parse_region_string(s):
    """
    Parse a UCSC-style genomic region string into a triple.
    Parameters
    ----------
    s : str
        UCSC-style string, e.g. "chr5:10,100,000-30,000,000". Ensembl and FASTA
        style sequence names are allowed. End coordinate must be greater than
        or equal to start.
    Returns
    -------
    (str, int or None, int or None)
    """

    def _tokenize(s):
        token_spec = [
            ("HYPHEN", r"-"),
            ("COORD", r"[0-9,]+(\.[0-9]*)?(?:[a-z]+)?"),
            ("OTHER", r".+"),
        ]
        tok_regex = r"\s*" + r"|\s*".join(r"(?P<%s>%s)" % pair for pair in token_spec)
        tok_regex = re.compile(tok_regex, re.IGNORECASE)
        for match in tok_regex.finditer(s):
            typ = match.lastgroup
            yield typ, match.group(typ)

    def _check_token(typ, token, expected):
        if typ is None:
            raise ValueError("Expected {} token missing".format(" or ".join(expected)))
        else:
            if typ not in expected:
                raise ValueError('Unexpected token "{}"'.format(token))

    def _expect(tokens):
        typ, token = next(tokens, (None, None))
        _check_token(typ, token, ["COORD"])
        start = parse_humanized(token)

        typ, token = next(tokens, (None, None))
        _check_token(typ, token, ["HYPHEN"])

        typ, token = next(tokens, (None, None))
        if typ is None:
            return start, None

        _check_token(typ, token, ["COORD"])
        end = parse_humanized(token)
        if end < start:
            raise ValueError("End coordinate less than start")

        return start, end

    parts = s.split(":")
    chrom = parts[0].strip()
    if not len(chrom):
        raise ValueError("Chromosome name cannot be empty")
    if len(parts) < 2:
        return (chrom, None, None)
    start, end = _expect(_tokenize(parts[1]))
    return (chrom, start, end)


def parse_region(reg, chromsizes=None):
    """
    Genomic regions are represented as half-open intervals (0-based starts,
    1-based ends) along the length coordinate of a contig/scaffold/chromosome.
    Parameters
    ----------
    reg : str or tuple
        UCSC-style genomic region string, or
        Triple (chrom, start, end), where ``start`` or ``end`` may be ``None``.
    chromsizes : mapping, optional
        Lookup table of scaffold lengths to check against ``chrom`` and the
        ``end`` coordinate. Required if ``end`` is not supplied.
    Returns
    -------
    A well-formed genomic region triple (str, int, int)
    """
    if isinstance(reg, six.string_types):
        chrom, start, end = parse_region_string(reg)
    else:
        chrom, start, end = reg
        start = int(start) if start is not None else start
        end = int(end) if end is not None else end
    try:
        clen = chromsizes[chrom] if chromsizes is not None else None
    except KeyError:
        raise ValueError("Unknown sequence label: {}".format(chrom))

    start = 0 if start is None else start
    if end is None:
        if clen is None:  # TODO --- remove?
            raise ValueError("Cannot determine end coordinate.")
        end = clen

    if end < start:
        raise ValueError("End cannot be less than start")

    if start < 0 or (clen is not None and end > clen):
        raise ValueError("Genomic region out of bounds: [{}, {})".format(start, end))

    return chrom, start, end


class Cooler(object):
    """
    A convenient interface to a cooler data collection.
    Parameters
    ----------
    store : str, :py:class:`h5py.File` or :py:class:`h5py.Group`
        Path to a cooler file, URI string, or open handle to the root HDF5
        group of a cooler data collection.
    root : str, optional [deprecated]
        HDF5 Group path to root of cooler group if ``store`` is a file.
        This option is deprecated. Instead, use a URI string of the form
        :file:`<file_path>::<group_path>`.
    kwargs : optional
        Options to be passed to :py:class:`h5py.File()` upon every access.
        By default, the file is opened with the default driver and mode='r'.
    Notes
    -----
    If ``store`` is a file path, the file will be opened temporarily in
    when performing operations. This allows :py:class:`Cooler` objects to be
    serialized for multiprocess and distributed computations.
    Metadata is accessible as a dictionary through the :py:attr:`info`
    property.
    Table selectors, created using :py:meth:`chroms`, :py:meth:`bins`, and
    :py:meth:`pixels`, perform range queries over table rows,
    returning :py:class:`pd.DataFrame` and :py:class:`pd.Series`.
    A matrix selector, created using :py:meth:`matrix`, performs 2D matrix
    range queries, returning :py:class:`numpy.ndarray` or
    :py:class:`scipy.sparse.coo_matrix`.
    """

    def __init__(self, store, root=None, **kwargs):
        if isinstance(store, six.string_types):
            if root is None:
                self.filename, self.root = parse_cooler_uri(store)
            elif h5py.is_hdf5(store):
                with open_hdf5(store, **kwargs) as h5:
                    self.filename = h5.file.filename
                    self.root = root
            else:
                raise ValueError("Not a valid path to a Cooler file")
            self.uri = self.filename + "::" + self.root
            self.store = self.filename
            self.open_kws = kwargs
        else:
            # Assume an open HDF5 handle, ignore open_kws
            self.filename = store.file.filename
            self.root = store.name
            self.uri = self.filename + "::" + self.root
            self.store = store.file
            self.open_kws = {}
        self._refresh()

    def _refresh(self):
        try:
            with open_hdf5(self.store, **self.open_kws) as h5:
                grp = h5[self.root]
                _ct = chroms(grp)
                _ct["name"] = _ct["name"].astype(object)
                self._chromsizes = _ct.set_index("name")["length"]
                self._chromids = dict(zip(_ct["name"], range(len(_ct))))
                self._info = info(grp)
                mode = self._info.get("storage-mode", u"symmetric-upper")
                self._is_symm_upper = mode == u"symmetric-upper"
        except KeyError:
            err_msg = "No cooler found at: {}.".format(self.store)
            listing = list_coolers(self.store)
            if len(listing):
                err_msg += (
                    " Coolers found in {}. ".format(listing)
                    + "Use '::' to specify a group path"
                )
            raise KeyError(err_msg)

    def _load_dset(self, path):
        with open_hdf5(self.store, **self.open_kws) as h5:
            grp = h5[self.root]
            return grp[path][:]

    def _load_attrs(self, path):
        with open_hdf5(self.store, **self.open_kws) as h5:
            grp = h5[self.root]
            return dict(grp[path].attrs)

    def open(self, mode="r", **kwargs):
        """ Open the HDF5 group containing the Cooler with :py:mod:`h5py`
        Functions as a context manager. Any ``open_kws`` passed during
        construction are ignored.
        Parameters
        ----------
        mode : str, optional [default: 'r']
            * ``'r'`` (readonly)
            * ``'r+'`` or ``'a'`` (read/write)
        Notes
        -----
            For other parameters, see :py:class:`h5py.File`.
        """
        grp = h5py.File(self.filename, mode, **kwargs)[self.root]
        return closing_hdf5(grp)

    @property
    def storage_mode(self):
        """Indicates whether ordinary sparse matrix encoding is used
        (``"square"``) or whether a symmetric matrix is encoded by storing only
        the upper triangular elements (``"symmetric-upper"``).
        """
        return self._info.get("storage-mode", u"symmetric-upper")

    @property
    def binsize(self):
        """ Resolution in base pairs if uniform else None """
        return self._info["bin-size"]

    @property
    def chromsizes(self):
        """ Ordered mapping of reference sequences to their lengths in bp """
        return self._chromsizes

    @property
    def chromnames(self):
        """ List of reference sequence names """
        return list(self._chromsizes.index)

    def offset(self, region):
        """ Bin ID containing the left end of a genomic region
        Parameters
        ----------
        region : str or tuple
            Genomic range
        Returns
        -------
        int
        Examples
        --------
        # >>> c.offset('chr3')  # doctest: +SKIP
        1311
        """
        with open_hdf5(self.store, **self.open_kws) as h5:
            grp = h5[self.root]
            return region_to_offset(
                grp, self._chromids, parse_region(region, self._chromsizes)
            )

    def extent(self, region):
        """ Bin IDs containing the left and right ends of a genomic region
        Parameters
        ----------
        region : str or tuple
            Genomic range
        Returns
        -------
        2-tuple of ints
        Examples
        --------
        # >>> c.extent('chr3')  # doctest: +SKIP
        (1311, 2131)
        """
        with open_hdf5(self.store, **self.open_kws) as h5:
            grp = h5[self.root]
            return region_to_extent(
                grp, self._chromids, parse_region(region, self._chromsizes)
            )

    @property
    def info(self):
        """ File information and metadata
        Returns
        -------
        dict
        """
        with open_hdf5(self.store, **self.open_kws) as h5:
            grp = h5[self.root]
            return info(grp)

    @property
    def shape(self):
        return (self._info["nbins"],) * 2

    def chroms(self, **kwargs):
        """ Chromosome table selector
        Returns
        -------
        Table selector
        """

        def _slice(fields, lo, hi):
            with open_hdf5(self.store, **self.open_kws) as h5:
                grp = h5[self.root]
                return chroms(grp, lo, hi, fields, **kwargs)

        return RangeSelector1D(None, _slice, None, self._info["nchroms"])

    def bins(self, **kwargs):
        """ Bin table selector
        Returns
        -------
        Table selector
        """

        def _slice(fields, lo, hi):
            with open_hdf5(self.store, **self.open_kws) as h5:
                grp = h5[self.root]
                return bins(grp, lo, hi, fields, **kwargs)

        def _fetch(region):
            with open_hdf5(self.store, **self.open_kws) as h5:
                grp = h5[self.root]
                return region_to_extent(
                    grp, self._chromids, parse_region(region, self._chromsizes)
                )

        return RangeSelector1D(None, _slice, _fetch, self._info["nbins"])

    def pixels(self, join=False, **kwargs):
        """ Pixel table selector
        Parameters
        ----------
        join : bool, optional
            Whether to expand bin ID columns into chrom, start, and end
            columns. Default is ``False``.
        Returns
        -------
        Table selector
        """

        def _slice(fields, lo, hi):
            with open_hdf5(self.store, **self.open_kws) as h5:
                grp = h5[self.root]
                return pixels(grp, lo, hi, fields, join, **kwargs)

        def _fetch(region):
            with open_hdf5(self.store, **self.open_kws) as h5:
                grp = h5[self.root]
                i0, i1 = region_to_extent(
                    grp, self._chromids, parse_region(region, self._chromsizes)
                )
                lo = grp["indexes"]["bin1_offset"][i0]
                hi = grp["indexes"]["bin1_offset"][i1]
                return lo, hi

        return RangeSelector1D(None, _slice, _fetch, self._info["nnz"])

    def matrix(
        self,
        field=None,
        balance=True,
        sparse=False,
        as_pixels=False,
        join=False,
        ignore_index=True,
        divisive_weights=None,
        max_chunk=500000000,
    ):
        """ Contact matrix selector
        Parameters
        ----------
        field : str, optional
            Which column of the pixel table to fill the matrix with. By
            default, the 'count' column is used.
        balance : bool, optional
            Whether to apply pre-calculated matrix balancing weights to the
            selection. Default is True and uses a column named 'weight'.
            Alternatively, pass the name of the bin table column containing
            the desired balancing weights. Set to False to return untransformed
            counts.
        sparse: bool, optional
            Return a scipy.sparse.coo_matrix instead of a dense 2D numpy array.
        as_pixels: bool, optional
            Return a DataFrame of the corresponding rows from the pixel table
            instead of a rectangular sparse matrix. False by default.
        join : bool, optional
            If requesting pixels, specifies whether to expand the bin ID
            columns into (chrom, start, end). Has no effect when requesting a
            rectangular matrix. Default is True.
        ignore_index : bool, optional
            If requesting pixels, don't populate the index column with the
            pixel IDs to improve performance. Default is True.
        divisive_weights : bool, optional
            Force balancing weights to be interpreted as divisive (True) or
            multiplicative (False). Weights are always assumed to be
            multiplicative by default unless named KR, VC or SQRT_VC, in which
            case they are assumed to be divisive by default.
        Returns
        -------
        Matrix selector
        Notes
        -----
        If ``as_pixels=True``, only data explicitly stored in the pixel table
        will be returned: if the cooler's storage mode is symmetric-upper,
        lower triangular elements will not be generated. If
        ``as_pixels=False``, those missing non-zero elements will
        automatically be filled in.
        """
        if balance in _4DN_DIVISIVE_WEIGHTS and divisive_weights is None:
            divisive_weights = True

        def _slice(field, i0, i1, j0, j1):
            with open_hdf5(self.store, **self.open_kws) as h5:
                grp = h5[self.root]
                return matrix(
                    grp,
                    i0,
                    i1,
                    j0,
                    j1,
                    field,
                    balance,
                    sparse,
                    as_pixels,
                    join,
                    ignore_index,
                    divisive_weights,
                    max_chunk,
                    self._is_symm_upper,
                )

        def _fetch(region, region2=None):
            with open_hdf5(self.store, **self.open_kws) as h5:
                grp = h5[self.root]
                if region2 is None:
                    region2 = region
                region1 = parse_region(region, self._chromsizes)
                region2 = parse_region(region2, self._chromsizes)
                i0, i1 = region_to_extent(grp, self._chromids, region1)
                j0, j1 = region_to_extent(grp, self._chromids, region2)
                return i0, i1, j0, j1

        return RangeSelector2D(field, _slice, _fetch, (self._info["nbins"],) * 2)

    def __repr__(self):
        if isinstance(self.store, six.string_types):
            filename = os.path.basename(self.store)
            container = "{}::{}".format(filename, self.root)
        else:
            container = repr(self.store)
        return '<Cooler "{}">'.format(container)


def _region_to_extent(h5, chrom_ids, region, binsize):
    chrom, start, end = region
    cid = chrom_ids[chrom]
    if binsize is not None:
        chrom_offset = h5["indexes"]["chrom_offset"][cid]
        yield chrom_offset + int(np.floor(start / binsize))
        yield chrom_offset + int(np.ceil(end / binsize))
    else:
        chrom_lo = h5["indexes"]["chrom_offset"][cid]
        chrom_hi = h5["indexes"]["chrom_offset"][cid + 1]
        chrom_bins = h5["bins"]["start"][chrom_lo:chrom_hi]
        yield chrom_lo + np.searchsorted(chrom_bins, start, "right") - 1
        yield chrom_lo + np.searchsorted(chrom_bins, end, "left")


def region_to_offset(h5, chrom_ids, region, binsize=None):
    return next(_region_to_extent(h5, chrom_ids, region, binsize))


def region_to_extent(h5, chrom_ids, region, binsize=None):
    return tuple(_region_to_extent(h5, chrom_ids, region, binsize))


def get(grp, lo=0, hi=None, fields=None, convert_enum=True, as_dict=False):
    """
    Query a range of rows from a table as a dataframe.
    A table is an HDF5 group containing equal-length 1D datasets serving as
    columns.
    Parameters
    ----------
    grp : ``h5py.Group`` or any dict-like of array-likes
        Handle to an HDF5 group containing only 1D datasets or any similar
        collection of 1D datasets or arrays
    lo, hi : int, optional
        Range of rows to select from the table.
    fields : str or sequence of str, optional
        Column or list of columns to query. Defaults to all available columns.
        A single string returns a Series instead of a DataFrame.
    convert_enum : bool, optional
        Whether to convert HDF5 enum datasets into ``pandas.Categorical``
        columns instead of plain integer columns. Default is True.
    kwargs : optional
        Options to pass to ``pandas.DataFrame`` or ``pandas.Series``.
    Returns
    -------
    DataFrame or Series
    Notes
    -----
    HDF5 ASCII datasets are converted to Unicode.
    """
    series = False
    if fields is None:
        fields = list(grp.keys())
    elif isinstance(fields, six.string_types):
        fields = [fields]
        series = True

    data = {}
    for field in fields:
        dset = grp[field]

        if convert_enum:
            dt = h5py.check_dtype(enum=dset.dtype)
        else:
            dt = None

        if dt is not None:
            data[field] = pd.Categorical.from_codes(
                dset[lo:hi], sorted(dt, key=dt.__getitem__), ordered=True
            )
        elif dset.dtype.type == np.string_:
            data[field] = dset[lo:hi].astype("U")
        else:
            data[field] = dset[lo:hi]

    if as_dict:
        return data

    if data and lo is not None:
        index = np.arange(lo, lo + len(next(iter(data.values()))))
    else:
        index = None

    if series:
        return pd.Series(data[fields[0]], index=index, name=field)
    else:
        return pd.DataFrame(data, columns=fields, index=index)


def info(h5):
    """
    File and user metadata dict.
    Parameters
    ----------
    h5 : :py:class:`h5py.File` or :py:class:`h5py.Group`
        Open handle to cooler file.
    Returns
    -------
    dict
    """
    d = {}
    for k, v in h5.attrs.items():
        if isinstance(v, six.string_types):
            try:
                v = json.loads(v)
            except ValueError:
                pass
        d[k] = v
    return d


def chroms(h5, lo=0, hi=None, fields=None, **kwargs):
    """
    Table describing the chromosomes/scaffolds/contigs used.
    They appear in the same order they occur in the heatmap.
    Parameters
    ----------
    h5 : :py:class:`h5py.File` or :py:class:`h5py.Group`
        Open handle to cooler file.
    lo, hi : int, optional
        Range of rows to select from the table.
    fields : sequence of str, optional
        Subset of columns to select from table.
    Returns
    -------
    :py:class:`DataFrame`
    """
    if fields is None:
        fields = (
            pd.Index(["name", "length"])
            .append(pd.Index(h5["chroms"].keys()))
            .drop_duplicates()
        )
    return get(h5["chroms"], lo, hi, fields, **kwargs)


def bins(h5, lo=0, hi=None, fields=None, **kwargs):
    """
    Table describing the genomic bins that make up the axes of the heatmap.
    Parameters
    ----------
    h5 : :py:class:`h5py.File` or :py:class:`h5py.Group`
        Open handle to cooler file.
    lo, hi : int, optional
        Range of rows to select from the table.
    fields : sequence of str, optional
        Subset of columns to select from table.
    Returns
    -------
    :py:class:`DataFrame`
    """
    if fields is None:
        fields = (
            pd.Index(["chrom", "start", "end"])
            .append(pd.Index(h5["bins"].keys()))
            .drop_duplicates()
        )

    # If convert_enum is not explicitly set to False, chrom IDs will get
    # converted to categorical chromosome names, provided the ENUM header
    # exists in bins/chrom. Otherwise, they will return as integers.
    out = get(h5["bins"], lo, hi, fields, **kwargs)

    # Handle the case where the ENUM header doesn't exist but we want to
    # convert integer chrom IDs to categorical chromosome names.
    if "chrom" in fields:
        convert_enum = kwargs.get("convert_enum", True)
        if isinstance(fields, six.string_types):
            chrom_col = out
        else:
            chrom_col = out["chrom"]

        if is_integer_dtype(chrom_col.dtype) and convert_enum:
            chromnames = chroms(h5, fields="name")
            chrom_col = pd.Categorical.from_codes(chrom_col, chromnames, ordered=True)
            if isinstance(fields, six.string_types):
                out = pd.Series(chrom_col, out.index)
            else:
                out["chrom"] = chrom_col

    return out


def pixels(h5, lo=0, hi=None, fields=None, join=True, **kwargs):
    """
    Table describing the nonzero upper triangular pixels of the Hi-C contact
    heatmap.
    Parameters
    ----------
    h5 : :py:class:`h5py.File` or :py:class:`h5py.Group`
        Open handle to cooler file.
    lo, hi : int, optional
        Range of rows to select from the table.
    fields : sequence of str, optional
        Subset of columns to select from table.
    join : bool, optional
        Whether or not to expand bin ID columns to their full bin description
        (chrom, start, end). Default is True.
    Returns
    -------
    :py:class:`DataFrame`
    """
    if fields is None:
        fields = (
            pd.Index(["bin1_id", "bin2_id"])
            .append(pd.Index(h5["pixels"].keys()))
            .drop_duplicates()
        )

    df = get(h5["pixels"], lo, hi, fields, **kwargs)

    if join:
        bins = get(h5["bins"], 0, None, ["chrom", "start", "end"], **kwargs)
        df = annotate(df, bins, replace=True)

    return df


def annotate(pixels, bins, replace=False):
    """
    Add bin annotations to a data frame of pixels.
    This is done by performing a relational "join" against the bin IDs of a
    table that describes properties of the genomic bins. New columns will be
    appended on the left of the output data frame.
    .. versionchanged:: 0.8.0
       The default value of ``replace`` changed to False.
    Parameters
    ----------
    pixels : :py:class:`DataFrame`
        A data frame containing columns named ``bin1_id`` and/or ``bin2_id``.
        If columns ``bin1_id`` and ``bin2_id`` are both present in ``pixels``,
        the adjoined columns will be suffixed with '1' and '2' accordingly.
    bins : :py:class:`DataFrame` or DataFrame selector
        Data structure that contains a full description of the genomic bins of
        the contact matrix, where the index corresponds to bin IDs.
    replace : bool, optional
        Remove the original ``bin1_id`` and ``bin2_id`` columns from the
        output. Default is False.
    Returns
    -------
    :py:class:`DataFrame`
    """
    columns = pixels.columns
    ncols = len(columns)

    if "bin1_id" in columns:
        if len(bins) > len(pixels):
            bin1 = pixels["bin1_id"]
            lo = bin1.min()
            hi = bin1.max() + 1
            lo = 0 if np.isnan(lo) else lo
            hi = 0 if np.isnan(hi) else hi
            right = bins[lo:hi]
        else:
            right = bins[:]

        pixels = pixels.merge(right, how="left", left_on="bin1_id", right_index=True)

    if "bin2_id" in columns:
        if len(bins) > len(pixels):
            bin2 = pixels["bin2_id"]
            lo = bin2.min()
            hi = bin2.max() + 1
            lo = 0 if np.isnan(lo) else lo
            hi = 0 if np.isnan(hi) else hi
            right = bins[lo:hi]
        else:
            right = bins[:]

        pixels = pixels.merge(
            right, how="left", left_on="bin2_id", right_index=True, suffixes=("1", "2")
        )

    # rearrange columns
    pixels = pixels[list(pixels.columns[ncols:]) + list(pixels.columns[:ncols])]

    # drop bin IDs
    if replace:
        cols_to_drop = [col for col in ("bin1_id", "bin2_id") if col in columns]
        pixels = pixels.drop(cols_to_drop, axis=1)

    return pixels


def matrix(
    h5,
    i0,
    i1,
    j0,
    j1,
    field=None,
    balance=True,
    sparse=False,
    as_pixels=False,
    join=True,
    ignore_index=True,
    divisive_weights=False,
    max_chunk=500000000,
    is_upper=True,
):
    """
    Two-dimensional range query on the Hi-C contact heatmap.
    Depending on the options, returns either a 2D NumPy array, a rectangular
    sparse ``coo_matrix``, or a data frame of pixels.
    Parameters
    ----------
    h5 : :py:class:`h5py.File` or :py:class:`h5py.Group`
        Open handle to cooler file.
    i0, i1 : int, optional
        Bin range along the 0th (row) axis of the heatap.
    j0, j1 : int, optional
        Bin range along the 1st (col) axis of the heatap.
    field : str, optional
        Which column of the pixel table to fill the matrix with. By default,
        the 'count' column is used.
    balance : bool, optional
        Whether to apply pre-calculated matrix balancing weights to the
        selection. Default is True and uses a column named 'weight'.
        Alternatively, pass the name of the bin table column containing the
        desired balancing weights. Set to False to return untransformed counts.
    sparse: bool, optional
        Return a scipy.sparse.coo_matrix instead of a dense 2D numpy array.
    as_pixels: bool, optional
        Return a DataFrame of the corresponding rows from the pixel table
        instead of a rectangular sparse matrix. False by default.
    join : bool, optional
        If requesting pixels, specifies whether to expand the bin ID columns
        into (chrom, start, end). Has no effect when requesting a rectangular
        matrix. Default is True.
    ignore_index : bool, optional
        If requesting pixels, don't populate the index column with the pixel
        IDs to improve performance. Default is True.
    Returns
    -------
    ndarray, coo_matrix or DataFrame
    Notes
    -----
    If ``as_pixels=True``, only data explicitly stored in the pixel table
    will be returned: if the cooler's storage mode is symmetric-upper,
    lower triangular elements will not be generated. If ``as_pixels=False``,
    those missing non-zero elements will automatically be filled in.
    """
    if field is None:
        field = "count"

    if isinstance(balance, str):
        name = balance
    elif balance:
        name = "weight"

    if balance and name not in h5["bins"]:
        raise ValueError(
            "No column 'bins/{}'".format(name)
            + "found. Use ``cooler.balance_cooler`` to "
            + "calculate balancing weights or set balance=False."
        )

    if as_pixels:
        reader = CSRReader(h5, field, max_chunk)
        index = None if ignore_index else reader.index_col(i0, i1, j0, j1)
        i, j, v = reader.query(i0, i1, j0, j1)

        cols = ["bin1_id", "bin2_id", field]
        df = pd.DataFrame(dict(zip(cols, [i, j, v])), columns=cols, index=index)

        if balance:
            weights = Cooler(h5).bins()[[name]]
            df2 = annotate(df, weights, replace=False)
            if divisive_weights:
                df2[name + "1"] = 1 / df2[name + "1"]
                df2[name + "2"] = 1 / df2[name + "2"]
            df["balanced"] = df2[name + "1"] * df2[name + "2"] * df2[field]

        if join:
            bins = Cooler(h5).bins()[["chrom", "start", "end"]]
            df = annotate(df, bins, replace=True)

        return df

    elif sparse:
        reader = CSRReader(h5, field, max_chunk)
        if is_upper:
            i, j, v = query_rect(reader.query, i0, i1, j0, j1, duplex=True)
        else:
            i, j, v = reader.query(i0, i1, j0, j1)
        mat = coo_matrix((v, (i - i0, j - j0)), (i1 - i0, j1 - j0))

        if balance:
            weights = h5["bins"][name]
            bias1 = weights[i0:i1]
            bias2 = bias1 if (i0, i1) == (j0, j1) else weights[j0:j1]
            if divisive_weights:
                bias1 = 1 / bias1
                bias2 = 1 / bias2
            mat.data = bias1[mat.row] * bias2[mat.col] * mat.data

        return mat

    else:
        reader = CSRReader(h5, field, max_chunk)
        if is_upper:
            i, j, v = query_rect(reader.query, i0, i1, j0, j1, duplex=True)
        else:
            i, j, v = reader.query(i0, i1, j0, j1)
        arr = coo_matrix((v, (i - i0, j - j0)), (i1 - i0, j1 - j0)).toarray()

        if balance:
            weights = h5["bins"][name]
            bias1 = weights[i0:i1]
            bias2 = bias1 if (i0, i1) == (j0, j1) else weights[j0:j1]
            if divisive_weights:
                bias1 = 1 / bias1
                bias2 = 1 / bias2
            arr = arr * np.outer(bias1, bias2)

        return arr


class _IndexingMixin(object):
    def _unpack_index(self, key):
        if isinstance(key, tuple):
            if len(key) == 2:
                row, col = key
            elif len(key) == 1:
                row, col = key[0], slice(None)
            else:
                raise IndexError("invalid number of indices")
        else:
            row, col = key, slice(None)
        return row, col

    def _isintlike(self, num):
        try:
            int(num)
        except (TypeError, ValueError):
            return False
        return True

    def _process_slice(self, s, nmax):
        if isinstance(s, slice):
            if s.step not in (1, None):
                raise ValueError("slicing with step != 1 not supported")
            i0, i1 = s.start, s.stop
            if i0 is None:
                i0 = 0
            elif i0 < 0:
                i0 = nmax + i0
            if i1 is None:
                i1 = nmax
            elif i1 < 0:
                i1 = nmax + i1
            return i0, i1
        elif self._isintlike(s):
            if s < 0:
                s += nmax
            if s >= nmax:
                raise IndexError("index is out of bounds")
            return int(s), int(s + 1)
        else:
            raise TypeError("expected slice or scalar")


class RangeSelector1D(_IndexingMixin):
    """
    Selector for out-of-core tabular data. Provides DataFrame-like selection of
    columns and list-like access to rows.
    Examples
    --------
    Passing a column name or list of column names as subscript returns a new
    selector.
    # >>> sel[ ['A', 'B'] ]  # doctest: +SKIP
    # >>> sel['C']
    # Passing a scalar or slice as subscript invokes the slicer.
    # >>> sel[0]  # doctest: +SKIP
    # >>> sel['A'][50:100]
    # Calling the fetch method invokes the fetcher to parse the input into an
    # integer range and then invokes the slicer.
    # >>> sel.fetch('chr3:10,000,000-12,000,000') # doctest: +SKIP
    # >>> sel.fetch(('chr3', 10000000, 12000000))
    """

    def __init__(self, fields, slicer, fetcher, nmax):
        self.fields = fields
        self._slice = slicer
        self._fetch = fetcher
        self._shape = (nmax,)

    @property
    def shape(self):
        return self._shape

    @property
    def columns(self):
        return self._slice(self.fields, 0, 0).columns

    @property
    def dtypes(self):
        return self._slice(self.fields, 0, 0).dtypes

    def keys(self):
        return list(self.columns)

    def __len__(self):
        return self._shape[0]

    def __contains__(self, key):
        return key in self.columns

    def __getitem__(self, key):
        # requesting a subset of columns
        if isinstance(key, (list, str)):
            return self.__class__(key, self._slice, self._fetch, self._shape[0])

        # requesting an interval of rows
        if isinstance(key, tuple):
            if len(key) == 1:
                key = key[0]
            else:
                raise IndexError("too many indices for table")
        lo, hi = self._process_slice(key, self._shape[0])
        return self._slice(self.fields, lo, hi)

    def fetch(self, *args, **kwargs):
        if self._fetch is not None:
            lo, hi = self._fetch(*args, **kwargs)
            return self._slice(self.fields, lo, hi)
        else:
            raise NotImplementedError


class RangeSelector2D(_IndexingMixin):
    """
    Selector for out-of-core sparse matrix data. Supports 2D scalar and slice
    subscript indexing.
    """

    def __init__(self, field, slicer, fetcher, shape):
        self.field = field
        self._slice = slicer
        self._fetch = fetcher
        self._shape = shape

    @property
    def shape(self):
        return self._shape

    def __len__(self):
        return self._shape[0]

    def __getitem__(self, key):
        s1, s2 = self._unpack_index(key)
        i0, i1 = self._process_slice(s1, self._shape[0])
        j0, j1 = self._process_slice(s2, self._shape[1])
        return self._slice(self.field, i0, i1, j0, j1)

    def fetch(self, *args, **kwargs):
        if self._fetch is not None:
            i0, i1, j0, j1 = self._fetch(*args, **kwargs)
            return self._slice(self.field, i0, i1, j0, j1)
        else:
            raise NotImplementedError


class CSRReader(object):
    """
    Retrieves data from a 2D range query on the pixel table of a Cooler.
    Parameters
    ----------
    h5 : ``h5py.File`` or ``h5py.Group``
        Root node of a cooler tree.
    field : str
        Column of the pixel table to query.
    max_chunk : int
        Size of largest chunk to read into memory in a single disk fetch.
        Increase this to increase performance for large queries at the cost of
        memory usage.
    """

    def __init__(self, h5, field, max_chunk):
        self.h5 = h5
        self.field = field
        self.max_chunk = max_chunk

    def index_col(self, i0, i1, j0, j1):
        """Retrieve pixel table row IDs corresponding to query rectangle."""
        edges = self.h5["indexes"]["bin1_offset"][i0 : i1 + 1]
        index = []
        for lo1, hi1 in zip(edges[:-1], edges[1:]):
            if hi1 - lo1 > 0:
                bin2 = self.h5["pixels"]["bin2_id"][lo1:hi1]
                mask = (bin2 >= j0) & (bin2 < j1)
                index.append(lo1 + np.flatnonzero(mask))
        if not index:
            return np.array([], dtype=int)
        else:
            return np.concatenate(index, axis=0)

    def query(self, i0, i1, j0, j1):
        """Retrieve sparse matrix data inside a query rectangle."""
        h5 = self.h5
        field = self.field

        i, j, v = [], [], []
        if (i1 - i0 > 0) or (j1 - j0 > 0):
            edges = h5["indexes"]["bin1_offset"][i0 : i1 + 1]
            data = h5["pixels"][field]
            p0, p1 = edges[0], edges[-1]

            if (p1 - p0) < self.max_chunk:
                all_bin2 = h5["pixels"]["bin2_id"][p0:p1]
                all_data = data[p0:p1]
                dtype = all_bin2.dtype
                for row_id, lo, hi in zip(
                    range(i0, i1), edges[:-1] - p0, edges[1:] - p0
                ):
                    bin2 = all_bin2[lo:hi]
                    mask = (bin2 >= j0) & (bin2 < j1)
                    cols = bin2[mask]

                    i.append(np.full(len(cols), row_id, dtype=dtype))
                    j.append(cols)
                    v.append(all_data[lo:hi][mask])
            else:
                for row_id, lo, hi in zip(range(i0, i1), edges[:-1], edges[1:]):
                    bin2 = h5["pixels"]["bin2_id"][lo:hi]
                    mask = (bin2 >= j0) & (bin2 < j1)
                    cols = bin2[mask]
                    dtype = bin2.dtype

                    i.append(np.full(len(cols), row_id, dtype=dtype))
                    j.append(cols)
                    v.append(data[lo:hi][mask])

        if not i:
            i = np.array([], dtype=int)
            j = np.array([], dtype=int)
            v = np.array([])
        else:
            i = np.concatenate(i, axis=0)
            j = np.concatenate(j, axis=0)
            v = np.concatenate(v, axis=0)

        return i, j, v


def _comes_before(a0, a1, b0, b1, strict=False):
    if a0 < b0:
        return a1 <= b0 if strict else a1 <= b1
    return False


def _contains(a0, a1, b0, b1, strict=False):
    if a0 > b0 or a1 < b1:
        return False
    if strict and (a0 == b0 or a1 == b1):
        return False
    return a0 <= b0 and a1 >= b1


def query_rect(triu_reader, i0, i1, j0, j1, duplex=True):
    """
    Process a 2D range query on a symmetric matrix using a reader that
    retrieves only upper triangle pixels from the matrix.
    This function is responsible for filling in the missing data in the query
    rectangle and for ensuring that diagonal elements are not duplicated.
    Removing duplicates is important because various sparse matrix constructors
    sum duplicates instead of ignoring them.
    Parameters
    ----------
    triu_reader : callable
        Callable that takes a query rectangle but only returns elements from
        the upper triangle of the parent matrix.
    i0, i1, j0, j1 : int
        Bounding matrix coordinates of the query rectangle. Assumed to be
        within the bounds of the parent matrix.
    Returns
    -------
    i, j, v : 1D arrays
    Details
    -------
    Query cases to consider based on the axes ranges (i0, i1) and (j0, j1):
    1. they are identical
    2. different and non-overlapping
    3. different but partially overlapping
    4. different but one is nested inside the other
    - (1) requires filling in the lower triangle.
    - (3) and (4) require splitting the selection into instances of
      (1) and (2).
    In some cases, the input axes ranges are swapped to retrieve the data,
    then the final result is transposed.
    """

    # symmetric query
    if (i0, i1) == (j0, j1):
        i, j, v = triu_reader(i0, i1, i0, i1)
        if duplex:
            nodiag = i != j
            i, j, v = np.r_[i, j[nodiag]], np.r_[j, i[nodiag]], np.r_[v, v[nodiag]]

    # asymmetric query
    else:
        transpose = False
        if j0 < i0 or (i0 == j0 and i1 < j1):
            i0, i1, j0, j1 = j0, j1, i0, i1
            transpose = True

        # non-overlapping
        if _comes_before(i0, i1, j0, j1, strict=True):
            i, j, v = triu_reader(i0, i1, j0, j1)

        # partially overlapping
        elif _comes_before(i0, i1, j0, j1):
            ix, jx, vx = triu_reader(i0, j0, j0, i1)
            iy, jy, vy = triu_reader(j0, i1, j0, i1)
            iz, jz, vz = triu_reader(i0, i1, i1, j1)
            if duplex:
                nodiag = iy != jy
                iy, jy, vy = (
                    np.r_[iy, jy[nodiag]],
                    np.r_[jy, iy[nodiag]],
                    np.r_[vy, vy[nodiag]],
                )
            i, j, v = np.r_[ix, iy, iz], np.r_[jx, jy, jz], np.r_[vx, vy, vz]

        # nested
        elif _contains(i0, i1, j0, j1):
            ix, jx, vx = triu_reader(i0, j0, j0, j1)
            jy, iy, vy = triu_reader(j0, j1, j0, j1)
            jz, iz, vz = triu_reader(j0, j1, j1, i1)
            if duplex:
                nodiag = iy != jy
                iy, jy, vy = (
                    np.r_[iy, jy[nodiag]],
                    np.r_[jy, iy[nodiag]],
                    np.r_[vy, vy[nodiag]],
                )
            i, j, v = np.r_[ix, iy, iz], np.r_[jx, jy, jz], np.r_[vx, vy, vz]

        else:
            raise IndexError("This shouldn't happen")

        if transpose:
            i, j = j, i

    return i, j, v
```
