## Supplementary material for "scHiCTools: a computational toolbox for analyzing single-cell Hi-C data": S4: hic_straw.py

```
"""Straw module

Straw enables programmatic access to .hic files.
.hic files store the contact matrices from Hi-C experiments and the
normalization and expected vectors, along with meta-data in the header.

The main function, straw, takes in the normalization, the filename or URL,
chromosome1 (and optional range), chromosome2 (and optional range),
whether the bins desired are fragment or base pair delimited, and bin size.

It then reads the header, follows the various pointers to the desired matrix
and normalization vector, and stores as [x, y, count]

Usage: straw <NONE/VC/VC_SQRT/KR> <hicFile(s)> <chr1>[:x1:x2] <chr2>[:y1:y2] <\
BP/FRAG> <binsize>

See https://github.com/theaidenlab/straw/wiki/Python for more documentation
"""

from __future__ import absolute_import, division, print_function
# unicode_literals

__author__ = "Yue Wu and Neva Durand"
__license__ = "MIT"

import sys
import struct
import zlib
import requests
import io

blockMap = dict()
# global version
version=0


def __readcstr(f):
    """ Helper function for reading in C-style string from file
    """
    buf = b""
    while True:
        b = f.read(1)
        if b is None or b == b"\0":
            return buf.decode("utf-8")
        elif b == "":
            raise EOFError("Buffer unexpectedly empty while trying to read null-terminated string")
        else:
            buf += b


def readHeader(req, chr1, chr2, posilist):
    """ Reads the header

    Args:
       req (file): File to read from
       chr1 (str): Chromosome 1
       chr2 (str): Chromosome 2
       c1pos1 (int, optional): Starting range of chromosome1 output
       c1pos2 (int, optional): Stopping range of chromosome1 output
       c2pos1 (int, optional): Starting range of chromosome2 output
       c2pos2 (int, optional): Stopping range of chromosome2 output

    Returns:
       list: master index, chromosome1 index, chromosome2 index
    """
    magic_string = struct.unpack('<3s', req.read(3))[0]
    req.read(1)
    if (magic_string != b"HIC"):
        print('This does not appear to be a HiC file magic string is incorrect')
        return -1
    global version
    version = struct.unpack('<i', req.read(4))[0]
    if (version < 6):
        print("Version {0} no longer supported".format(str(version)))
        return -1
    # print('HiC version:', version)

    master = struct.unpack('<q', req.read(8))[0]
    # print('Master: ', master)

    genome = b""
    c = req.read(1)
    while (c != b'\0'):
        genome += c
        c = req.read(1)
    # print('Genome: ', genome.decode('utf-8'))

    # read and throw away attribute dictionary (stats+graphs)
    nattributes = struct.unpack('<i', req.read(4))[0]
    for x in range(nattributes):
        key = __readcstr(req)
        # print('Attributes:', key, ': ', end='')
        value = __readcstr(req)
        # print(value)
    nChrs = struct.unpack('<i', req.read(4))[0]
    found1 = False
    found2 = False
    for i in range(0, nChrs):
      name = __readcstr(req)
      if not name.startswith('chr'):
          name = 'chr' + name
      length = struct.unpack('<i', req.read(4))[0]

      if name == chr1:
          # print(name, ': ', length)
          found1 = True
          chr1ind = i
          if (posilist[0] == -100):
              posilist[0] = 0
              posilist[1] = length
      if name == chr2:
          # print(name, ': ', length)
          found2 = True
          chr2ind = i
          if (posilist[2] == -100):
              posilist[2] = 0
              posilist[3] = length
    if (not found1) or (not found2):
      print("One of the chromosomes wasn't found in the file. Check that the chromosome name matches the genome.\n")
      return -1
    return [master, chr1ind, chr2ind, posilist[0], posilist[1], posilist[2], posilist[3]]


def readFooter(req, c1, c2, norm, unit, resolution):
    """Reads the footer, which contains all the expected and normalization
    vectors. Presumes file pointer is in correct position
    Args:
       req (file): File to read from; presumes file pointer is in correct position
       chr1 (str): Chromosome 1
       chr2 (str): Chromosome 2
       norm (str): Normalization type, one of NONE, VC, KR, VC_SQRT
       unit (str): One of BP or FRAG
       resolution (int): Bin size

    Returns:
       list: File position of matrix, position+size chr1 normalization vector,
             position+size chr2 normalization vector
    """
    c1NormEntry = dict()
    c2NormEntry = dict()
    nBytes = struct.unpack('<i', req.read(4))[0]
    # print('Footer nBytes: ', nBytes)

    key = str(c1) + "_" + str(c2)
    nEntries = struct.unpack('<i', req.read(4))[0]
    # print('Footer nEntries: ', nEntries)

    found = False
    for i in range(nEntries):
        stri = __readcstr(req)
        # print('Footer: ', stri)
        fpos = struct.unpack('<q', req.read(8))[0]
        # print('Footer fpos: ', fpos)
        sizeinbytes = struct.unpack('<i', req.read(4))[0]
        # print('Footer size in bytes: ', sizeinbytes)
        if stri == key:
            myFilePos = fpos
            found = True
            # print('Footer: ', stri, key, fpos)
    if not found:
        print("File doesn't have the given chr_chr map\n")
    if norm == "NONE":
        return [myFilePos, 0, 0]

    nExpectedValues = struct.unpack('<i', req.read(4))[0]
    for i in range(nExpectedValues):
        str_ = __readcstr(req)
        binSize = struct.unpack('<i', req.read(4))[0]
        nValues = struct.unpack('<i', req.read(4))[0]
        for j in range(nValues):
            v = struct.unpack('<d',req.read(8))[0]
        nNormalizationFactors = struct.unpack('<i',req.read(4))[0]
        for j in range(nNormalizationFactors):
            chrIdx = struct.unpack('<i',req.read(4))[0]
            v = struct.unpack('<d',req.read(8))[0]
    nExpectedValues = struct.unpack('<i',req.read(4))[0]
    for i in range(nExpectedValues):
        str_ = __readcstr(req)
        str_ = __readcstr(req)
        binSize = struct.unpack('<i',req.read(4))[0]
        nValues = struct.unpack('<i',req.read(4))[0]
        for j in range(nValues):
            v = struct.unpack('<d',req.read(8))[0]
        nNormalizationFactors = struct.unpack('<i',req.read(4))[0]
        for j in range(nNormalizationFactors):
            chrIdx = struct.unpack('<i',req.read(4))[0]
            v = struct.unpack('<d',req.read(8))[0]
    nEntries = struct.unpack('<i',req.read(4))[0]
    found1 = False
    found2 = False
    for i in range(nEntries):
        normtype = __readcstr(req)
        chrIdx = struct.unpack('<i',req.read(4))[0]
        unit1 = __readcstr(req)
        resolution1 = struct.unpack('<i',req.read(4))[0]
        filePosition = struct.unpack('<q',req.read(8))[0]
        sizeInBytes = struct.unpack('<i',req.read(4))[0]
        if (chrIdx==c1 and normtype==norm and unit1==unit and resolution1==resolution):
            c1NormEntry['position']=filePosition
            c1NormEntry['size']=sizeInBytes
            found1=True
        if (chrIdx==c2 and normtype==norm and unit1==unit and resolution1==resolution):
            c2NormEntry['position']=filePosition
            c2NormEntry['size']=sizeInBytes
            found2=True
    if ((not found1) or (not found2)):
        print("File did not contain {0} normalization vectors for one or both chromosomes at {1} {2}\n".format(norm, resolution, unit))
        return -1
    return [myFilePos, c1NormEntry, c2NormEntry]


def readMatrixZoomData(req, myunit, mybinsize):
    """ Reads the Matrix Zoom Data, which gives pointer list for blocks for
    the data. Presumes file pointer is in correct position

    Args:
       req (file): File to read from; presumes file pointer is in correct position
       myunit (str): Unit (BP or FRAG) we're searching for
       mybinsize (int): Resolution we're searching for

    Returns:
       list containing boolean indicating if we found appropriate matrix,
       and if so, the counts for the bins and columns
    """
    unit = __readcstr(req)
    # print(unit)
    temp = struct.unpack('<i', req.read(4))[0]
    # print(temp)
    temp2 = struct.unpack('<f', req.read(4))[0]
    # print(temp2)
    temp2 = struct.unpack('<f', req.read(4))[0]
    # print(temp2)
    temp2 = struct.unpack('<f', req.read(4))[0]
    # print(temp2)
    temp2 = struct.unpack('<f', req.read(4))[0]
    # print(temp2)

    binSize = struct.unpack('<i', req.read(4))[0]
    #print('MatrixZoom: ', binSize)
    blockBinCount = struct.unpack('<i', req.read(4))[0]
    #print('MatrixZoom: ', blockBinCount)
    blockColumnCount = struct.unpack('<i', req.read(4))[0]
    #print('MatrixZoom: ', blockColumnCount)
    storeBlockData = False
    # print(unit, binSize)
    
    #for the initial
    myBlockBinCount = -1
    myBlockColumnCount = -1
    if myunit == unit and mybinsize == binSize:
        myBlockBinCount = blockBinCount
        myBlockColumnCount = blockColumnCount
        storeBlockData = True

    nBlocks = struct.unpack('<i', req.read(4))[0]
    # print(nBlocks)
    for b in range(nBlocks):
        blockNumber = struct.unpack('<i', req.read(4))[0]
        filePosition = struct.unpack('<q', req.read(8))[0]
        blockSizeInBytes = struct.unpack('<i', req.read(4))[0]
        entry = dict()
        entry['size'] = blockSizeInBytes
        entry['position'] = filePosition
        # print(b, blockNumber, entry)

        if storeBlockData:
            blockMap[blockNumber] = entry
    return [storeBlockData, myBlockBinCount, myBlockColumnCount, binSize]
    #  , binSize ?


def readMatrix(req, unit, binsize):
    """ Reads the matrix - that is, finds the appropriate pointers to block
    data and stores them. Needs to read through headers of zoom data to find
    appropriate matrix. Presumes file pointer is in correct position.

    Args:
       req (file): File to read from; presumes file pointer is in correct position
       unit (str): Unit to search for (BP or FRAG)
       binsize (int): Resolution to search for

    Returns:
       list containing block bin count and block column count of matrix
    """
    c1 = struct.unpack('<i', req.read(4))[0]
    #print('read matrix c1:', c1)
    c2 = struct.unpack('<i', req.read(4))[0]
    #print('read matrix c2:', c2)

    nRes = struct.unpack('<i', req.read(4))[0]
    #print('nRes: ', nRes)

    i = 0
    found = False
    blockBinCount = -1
    blockColumnCount = -1

    res = []
    while i < nRes and (not found):
        list1 = readMatrixZoomData(req, unit, binsize)
        found = list1[0]
        if not found:
            res.append(list1[3])
        if list1[1] != -1 and list1[2] != -1:
            blockBinCount = list1[1]
            blockColumnCount = list1[2]
        i = i+1
    if not found:
        raise ValueError('Error finding block data. Resolution should be in: {}'.format(res))
    return [blockBinCount, blockColumnCount]

def getBlockNumbersForRegionFromBinPosition(regionIndices, blockBinCount, blockColumnCount, intra):
    """ Gets the block numbers we will need for a specific region; used when
    the range to extract is sent in as a parameter

    Args:
       regionIndices (array): Array of ints giving range
       blockBinCount (int): The block bin count of the matrix
       blockColumnCount (int): The block column count of the matrix
       intra: Flag indicating if this is an intrachromosomal matrix

    Returns:
       blockSet (set): A set of blocks to print
    """
    col1 = int(regionIndices[0] / blockBinCount)
    col2 = int((regionIndices[1] + 1) / blockBinCount)
    row1 = int(regionIndices[2] / blockBinCount)
    row2 = int((regionIndices[3] + 1) / blockBinCount)
    blocksSet = set()
    # print(str(col1)+"\t"+str(col2)+"\t"+str(row1)+"\t"+str(row2))
    for r in range(row1, row2+1):
        for c in range(col1, col2+1):
            blockNumber = r * blockColumnCount + c
            blocksSet.add(blockNumber)
    if intra:
        for r in range(col1, col2+1):
            for c in range(row1, row2+1):
                blockNumber = r * blockColumnCount + c
                blocksSet.add(blockNumber)
    # print(str(blocksSet))
    return blocksSet


def readBlock(req, size):
    """ Reads the block - reads the compressed bytes, decompresses, and stores
    results in array. Presumes file pointer is in correct position.

    Args:
       req (file): File to read from. Presumes file pointer is in correct
       position
       size (int): How many bytes to read

    Returns:
       array containing row, column, count data for this block
    """
    compressedBytes = req.read(size)
    uncompressedBytes = zlib.decompress(compressedBytes)
    nRecords = struct.unpack('<i', uncompressedBytes[0:4])[0]
    v = []
    global version
    if version < 7:
        for i in range(nRecords):
            binX = struct.unpack('<i',uncompressedBytes[(12*i+4):(12*i+8)])[0]
            binY = struct.unpack('<i',uncompressedBytes[(12*i+8):(12*i+12)])[0]
            counts = struct.unpack('<f',uncompressedBytes[(12*i+12):(12*i+16)])[0]
            record = dict()
            record['binX'] = binX
            record['binY'] = binY
            record['counts'] = counts
            v.append(record)
    else:
        binXOffset = struct.unpack('<i', uncompressedBytes[4:8])[0]
        binYOffset = struct.unpack('<i', uncompressedBytes[8:12])[0]
        useShort = struct.unpack('<b', uncompressedBytes[12:13])[0]
        type_ = struct.unpack('<b', uncompressedBytes[13:14])[0]
        # print(binXOffset, binYOffset, useShort, type_)
        # print('Type: ', type_)
        index = 0
        if type_ == 1:
            rowCount = struct.unpack('<h', uncompressedBytes[14:16])[0]
            #print('RowCount: ', rowCount)
            temp = 16
            for i in range(rowCount):
                y = struct.unpack('<h', uncompressedBytes[temp:(temp+2)])[0]
                temp = temp + 2
                binY = y + binYOffset
                colCount = struct.unpack('<h', uncompressedBytes[temp:(temp+2)])[0]
                temp = temp + 2
                for j in range(colCount):
                    x = struct.unpack('<h', uncompressedBytes[temp:(temp+2)])[0]
                    temp = temp + 2
                    binX = binXOffset + x

                    if useShort == 0:
                        c = struct.unpack('<h', uncompressedBytes[temp:(temp+2)])[0]
                        temp = temp + 2
                        counts = c
                    else:
                        counts = struct.unpack('<f', uncompressedBytes[temp:(temp+4)])[0]
                        temp = temp + 4

                    # print(binX, binY, counts)

                    record = dict()
                    record['binX'] = binX
                    record['binY'] = binY
                    record['counts'] = counts
                    v.append(record)
                    index = index + 1
        elif (type_== 2):
            temp=14
            nPts = struct.unpack('<i',uncompressedBytes[temp:(temp+4)])[0]
            temp=temp+4
            w = struct.unpack('<h',uncompressedBytes[temp:(temp+2)])[0]
            temp=temp+2
            for i in range(nPts):
                row=int(i/w)
                col=i-row*w
                bin1=int(binXOffset+col)
                bin2=int(binYOffset+row)
                if (useShort==0):
                    c = struct.unpack('<h',uncompressedBytes[temp:(temp+2)])[0]
                    temp=temp+2
                    if (c != -32768):
                        record = dict()
                        record['binX'] = bin1
                        record['binY'] = bin2
                        record['counts'] = c
                        v.append(record)
                        index = index + 1
                else:
                    counts = struct.unpack('<f',uncompressedBytes[temp:(temp+4)])[0]
                    temp=temp+4
                    if (counts != 0x7fc00000):
                        record = dict()
                        record['binX'] = bin1
                        record['binY'] = bin2
                        record['counts'] = counts
                        v.append(record)
                        index = index + 1
    return v


def readNormalizationVector(req):
    """ Reads the normalization vector from the file; presumes file pointer is
    in correct position

    Args:
       req (file): File to read from; presumes file pointer is in correct
       position

    Returns:
      Array of normalization values

    """
    value = []
    nValues = struct.unpack('<i',req.read(4))[0]
    for i in range(nValues):
        d = struct.unpack('<d',req.read(8))[0]
        value.append(d)
    return value


def straw(norm, infile, chr1loc, chr2loc, unit, binsize, is_synapse=False):
    """ This is the main workhorse method of the module. Reads a .hic file and
    extracts the given contact matrix. Stores in an array in sparse upper
    triangular format: row, column, (normalized) count

    Args:
       norm(str): Normalization type, one of VC, KR, VC_SQRT, or NONE
       infile(str): File name or URL of .hic file
       chr1loc(str): Chromosome name and (optionally) range, i.e. "1" or "1:10000:25000"
       chr2loc(str): Chromosome name and (optionally) range, i.e. "1" or "1:10000:25000"
       unit(str): One of BP or FRAG
       binsize(int): Resolution, i.e. 25000 for 25K
    """
    # clear the global variable blockMap so that it won't keep the data from previous calls
    for blockNum in list(blockMap.keys()):
        blockMap.pop(blockNum)

    magic_string = ""
    if infile.startswith("http"):
        # try URL first. 100K should be sufficient for header
        headers = {'range': 'bytes=0-100000', 'x-amz-meta-requester': 'straw'}
        if is_synapse:
            headers = {'range': 'bytes=0-100000'}
        s = requests.Session()
        r = s.get(infile, headers=headers)
        if r.status_code >= 400:
            print("Error accessing " + infile)
            print("HTTP status code " + str(r.status_code))
            return -1
        req = io.BytesIO(r.content)
        myrange = r.headers['content-range'].split('/')
        totalbytes = myrange[1]
    else:
        req = open(infile, 'rb')

    if not (norm=="NONE" or norm=="VC" or norm=="VC_SQRT" or norm=="KR"):
        print("Norm specified incorrectly, must be one of <NONE/VC/VC_SQRT/KR>\nUsage: straw <NONE/VC/VC_SQRT/KR> <hicFile(s)> <chr1>[:x1:x2] <chr2>[:y1:y2] <BP/FRAG> <binsize>\n")
        return -1
    if not (unit=="BP" or unit=="FRAG"):
        print("Unit specified incorrectly, must be one of <BP/FRAG>\nUsage: straw <NONE/VC/VC_SQRT/KR> <hicFile(s)> <chr1>[:x1:x2] <chr2>[:y1:y2] <BP/FRAG> <binsize>\n")
        return -1

    c1pos1 = -100
    c1pos2 = -100
    c2pos1 = -100
    c2pos2 = -100
    chr1_arra = chr1loc.split(":")
    chr2_arra = chr2loc.split(":")
    chr1 = chr1_arra[0]
    chr2 = chr2_arra[0]
    if len(chr1_arra) == 3:
        c1pos1 = chr1_arra[1]
        c1pos2 = chr1_arra[2]
    if len(chr2_arra) == 3:
        c2pos1 = chr2_arra[1]
        c2pos2 = chr2_arra[2]

    list1 = readHeader(req, chr1, chr2, [c1pos1, c1pos2, c2pos1, c2pos2])
    # print(chr1loc, chr2loc)
    # print(req, chr1, chr2, [c1pos1, c1pos2, c2pos1, c2pos2])
    # print(list1)

    master = list1[0]
    chr1ind = list1[1]
    chr2ind = list1[2]
    c1pos1 = int(list1[3])
    c1pos2 = int(list1[4])
    c2pos1 = int(list1[5])
    c2pos2 = int(list1[6])
    c1 = min(chr1ind, chr2ind)
    c2 = max(chr1ind, chr2ind)

    #####################
    # eg. [7712922943 (length), 21 (chr X), 1 (chr 1), 0, 10000000, 0, 10000000]
    #####################

    origRegionIndices = []
    regionIndices = []

    if chr1ind > chr2ind:
        origRegionIndices.append(c2pos1)
        origRegionIndices.append(c2pos2)
        origRegionIndices.append(c1pos1)
        origRegionIndices.append(c1pos2)
        regionIndices.append(int(c2pos1/binsize))
        regionIndices.append(int(c2pos2/binsize))
        regionIndices.append(int(c1pos1/binsize))
        regionIndices.append(int(c1pos2/binsize))
    else:
        origRegionIndices.append(c1pos1)
        origRegionIndices.append(c1pos2)
        origRegionIndices.append(c2pos1)
        origRegionIndices.append(c2pos2)
        regionIndices.append(int(c1pos1/binsize))
        regionIndices.append(int(c1pos2/binsize))
        regionIndices.append(int(c2pos1/binsize))
        regionIndices.append(int(c2pos2/binsize))

    # Get footer: from master to end of file
    if (infile.startswith("http")):
        headers = {'range': 'bytes={0}-{1}'.format(master, totalbytes), 'x-amz-meta-requester': 'straw'}
        if is_synapse:
            headers = {'range' : 'bytes={0}-{1}'.format(master, totalbytes)}
        r = s.get(infile, headers=headers)
        req = io.BytesIO(r.content)
    else:
        req.seek(master)

    list1 = readFooter(req, c1, c2, norm, unit, binsize)
    # print('Footer: ', list1)
    ###########################
    # eg. [7542224070, 0, 0] if no normalization
    ###########################

    myFilePos = list1[0]
    c1NormEntry = list1[1]
    c2NormEntry = list1[2]

    if norm != "NONE":
        if infile.startswith("http"):
            endrange = 'bytes={0}-{1}'.format(c1NormEntry['position'], c1NormEntry['position']+c1NormEntry['size'])
            headers = {'range': endrange, 'x-amz-meta-requester': 'straw'}
            if is_synapse:
                headers={'range': endrange}
            r = s.get(infile, headers=headers)
            req = io.BytesIO(r.content)
            c1Norm = readNormalizationVector(req)

            endrange = 'bytes={0}-{1}'.format(c2NormEntry['position'], c2NormEntry['position']+c2NormEntry['size'])
            headers = {'range': endrange, 'x-amz-meta-requester': 'straw'}
            if is_synapse:
                headers = {'range': endrange}
            r = s.get(infile, headers=headers)
            req = io.BytesIO(r.content)
            c2Norm = readNormalizationVector(req)
        else:
            req.seek(c1NormEntry['position'])
            c1Norm = readNormalizationVector(req)
            req.seek(c2NormEntry['position'])
            c2Norm = readNormalizationVector(req)

    if infile.startswith("http"):
        headers = {'range': 'bytes={0}-'.format(myFilePos), 'x-amz-meta-requester': 'straw'}
        if is_synapse:
            headers = {'range' : 'bytes={0}-'.format(myFilePos)}
        r = s.get(infile, headers=headers, stream=True)
        list1 = readMatrix(r.raw, unit, binsize)
    else:
        req.seek(myFilePos)
        list1 = readMatrix(req, unit, binsize)
        # print(list1)
    blockBinCount = list1[0]
    blockColumnCount = list1[1]
    blockNumbers = getBlockNumbersForRegionFromBinPosition(regionIndices, blockBinCount, blockColumnCount, c1 == c2)

    yActual = []
    xActual = []
    counts = []

    # print(blockMap)
    # print('All blocks: ', blockNumbers)
    for i_set in blockNumbers:
        idx = dict()
        if i_set in blockMap:
            idx = blockMap[i_set]
        else:
            idx['size'] = 0
            idx['position'] = 0
        # print(i_set, idx)
        if idx['size'] == 0:
            records = []
        else:
            if (infile.startswith("http")):
                endrange='bytes={0}-{1}'.format(idx['position'], idx['position']+idx['size'])
                headers={'range' : endrange, 'x-amz-meta-requester' : 'straw'}
                if is_synapse:
                    headers={'range' : endrange}
                r = s.get(infile, headers=headers)
                req = io.BytesIO(r.content)
            else:
                req.seek(idx['position'])
            records = readBlock(req, idx['size'])

        for j in range(len(records)):
            rec = records[j]
            # x = rec['binX'] * binsize
            x = rec['binX']
            # y = rec['binY'] * binsize
            y = rec['binY']
            c = rec['counts']
            if (norm != "NONE"):
                a = c1Norm[rec['binX']]*c2Norm[rec['binY']]
                if (a!=0.0):
                    c=(c/(c1Norm[rec['binX']]*c2Norm[rec['binY']]))
                else:
                    c="inf"
            if ((x>=origRegionIndices[0] and x<=origRegionIndices[1]
                 and y>=origRegionIndices[2] and y<=origRegionIndices[3])
                    or ((c1==c2) and y>=origRegionIndices[0] and y<=origRegionIndices[1]
                        and x>= origRegionIndices[2] and x<=origRegionIndices[3])):
                yield x, y, c
```
