## Supplementary material for "scHiCTools: a computational toolbox for analyzing single-cell Hi-C data": S4: load_hic_file.py

```
# -*- coding: utf-8 -*-
"""
This file is for loading a single HiC contact map from different type of file.
"""

from scipy.sparse import coo_matrix, load_npz
import numpy as np
from gzip import open as gopen
import os.path
from .hic_straw import straw
from .cool import dump
from .processing_utils import matrix_operation

my_path = os.path.abspath(os.path.dirname(__file__))


def get_chromosome_lengths(ref, chromosomes, res=1):
    """
    Get lengths for all chromosomes in a given resolution according to the reference genome.

    Args:
        ref (str or dict): name of reference genome, eg.'mm10', 'hg38'
        res (int): resolution
        chromosomes (str): 'All', 'except X', 'except Y' or a list of chromosomes like ['chr1', 'chr2']

    Returns:
        chromosomes (set): 
        lengths (dict): eg. {'chr1': 395, 'chr2': 390, ...}
    """
    def _res_change(length, res):
        if length % res == 0:
            return length // res
        else:
            return length // res + 1

    if isinstance(ref, str):
        try:
            path = os.path.join(my_path, 'reference_genome/' + ref)
            rg = open(path)
        except FileNotFoundError:
            try:
                path = os.path.join(my_path, 'reference_genome\\' + ref)
                rg = open(path)
            except FileNotFoundError:
                raise FileNotFoundError('Reference genome {0} not supported!'.format(ref))
        rg = [line.strip().split() for line in rg]
        lengths = {lst[0]: _res_change(int(lst[1]), res) for lst in rg}
    elif isinstance(ref, dict):
        lengths = {elm: _res_change(ref[elm], res) for elm in ref}
    else:
        raise ValueError('Unsupported reference genome!')

    if chromosomes in ['all', 'All']:
        pass
    elif chromosomes == 'expect X':
        del lengths['chrX']
    elif chromosomes == 'except Y':
        del lengths['chrY']
    elif chromosomes == 'except XY':
        del lengths['chrX']
        del lengths['chrY']
    elif type(chromosomes) == list:
        lengths = {elm: lengths[elm] for elm in chromosomes}

    chromosomes = set(lengths.keys())
    return chromosomes, lengths


def file_line_generator(file, format=None, chrom=None, header=0, resolution=1,
                        resolution_adjust=True, mapping_filter=0., gzip=False):
    """
    For formats other than .hic and .mcool

    Args:
        file (str): File path.
        
        format (int or list): Format 
            1) [chr1 pos1 chr2 pos2 mapq1 mapq2];
            2) [chr1 pos1 chr2 pos2 score];
            3) [chr1 pos1 chr2 pos2].
    
        chrom (str): Chromosome to extract.
        
        header (None or int): Whether to skip header (1 line).
    
        mapping_filter (float): The threshold to filter some reads by map quality.
        

    Returns:
        No return value.
        Yield a line each time in the format of (position_1, position_2, contact_reads).
    """

    f = gopen(file) if gzip else open(file)
    if header:
        for _ in range(header):
            next(f)
    for line in f:
        lst = line.strip().split()
        if len(format) not in [4, 5, 6]:
            raise ValueError('Wrong custom format!')

        if format[0] != 0 and format[2] != 0:
            # chr1 chr2
            c1, c2 = lst[format[0]-1], lst[format[2]-1]
            if (c1 != chrom and 'chr' + c1 != chrom) or (c2 != chrom and 'chr' + c2 != chrom):
                continue

        if len(format) == 6:  # [chr1 pos1 chr2 pos2 mapq1 mapq2]
            # mapq1 mapq2
            q1, q2 = float(lst[format[4]-1]), float(lst[format[4]-1])
            if q1 < mapping_filter or q2 < mapping_filter:
                continue

        # pos1 pos2
        p1, p2 = int(lst[format[1]-1]), int(lst[format[3]-1])
        if resolution_adjust:
            p1 = p1 // resolution  # * resolution
            p2 = p2 // resolution  # * resolution

        if len(format) == 4 or len(format) == 6:  # [chr1 pos1 chr2 pos2]
            v = 1.0
        elif len(format) == 5:
            v = float(lst[format[4]-1])
        else:
            raise ValueError('Wrong custom format!')

        yield p1, p2, v
    f.close()


def load_HiC(file, genome_length, format=None, custom_format=None, 
             header=0, chromosome=None, resolution=10000,
             resolution_adjust=True, map_filter=0., sparse=False, gzip=False,
             keep_n_strata=False, operations=None, **kwargs):
    """
    Load HiC contact map into a matrix
    Args:
        file (str): File path.
        genome_length (dict): The length of each genome.
        format (str): Now support .txt, .hic, and .mcool file.
        custom_format (int or list): If the format is not in our provided list.
        header (int): How many header lines to skip.
        chromosome (str): Specify the chromosome.
        resolution (int): Resolution.
        resolution_adjust (bool): In some situations, the input file is already pre-processed, and we don't need to adjust resolution again.
        map_filter (float): The threshold to filter some reads by map quality
        sparse (bool): Whether store in sparse matrices
        gzip (bool): Whether the file is zipped.
        keep_n_strata (int or None): Number of strata to keep.

    Return:
        Numpy.array: loaded contact map
    """
    size = genome_length[chromosome]

    format = format.lower()
    # If the inputs are pre-processed matrices
    if format in ['npy', 'npz', 'hicrep', 'matrix_txt', 'matrix']:
        if format == 'npy':
            mat = np.load('{0}/{1}.npy'.format(file, chromosome))
        elif format == 'npz':
            mat = load_npz('{0}/{1}.npz'.format(file, chromosome))
            mat = mat.toarray()
        elif format == 'matrix_txt':
            mat = np.loadtxt('{0}/{1}.txt'.format(file, chromosome))
        elif format == 'matrix':
            mat = np.loadtxt('{0}/{1}'.format(file, chromosome))
        else:  # HiCRep
            mat = np.loadtxt('{0}/{1}'.format(file, chromosome), dtype=str)
            mat = mat[1:, 5:].astype(float)

    # If the inputs are other files, pick a generator to generate a line of the file each time
    else:
        # .hic and .mcool files
        if format in ['hic', '.hic']:
            gen = straw('NONE', file, chromosome, chromosome, 'BP', resolution)
        elif format in ['mcool', 'cool', '.cool', '.mcool']:
            gen = dump(file, resolution=resolution, range=chromosome, range2=chromosome, header=header > 0)

        # text files
        elif format == 'shortest':
            gen = file_line_generator(
                file, format=[1, 2, 3, 4], chrom=chromosome, header=0, gzip=gzip,
                resolution=resolution, resolution_adjust=resolution_adjust, mapping_filter=0
            )
        elif format == 'shortest_score':
            gen = file_line_generator(
                file, format=[1, 2, 3, 4, 5], chrom=chromosome, header=0, gzip=gzip,
                resolution=resolution, resolution_adjust=resolution_adjust, mapping_filter=0
            )
        elif format == 'short':
            gen = file_line_generator(
                file, format=[2, 3, 6, 7], chrom=chromosome, header=0, gzip=gzip,
                resolution=resolution, resolution_adjust=resolution_adjust, mapping_filter=0
            )
        elif format == 'short_score':
            gen = file_line_generator(
                file, format=[2, 3, 6, 7, 9], chrom=chromosome, header=0, gzip=gzip,
                resolution=resolution, resolution_adjust=resolution_adjust, mapping_filter=0
            )
        elif format == 'medium':
            gen = file_line_generator(
                file, format=[3, 4, 7, 8, 10, 11], chrom=chromosome, 
                header=0, gzip=gzip, resolution=resolution,
                resolution_adjust=resolution_adjust,
                mapping_filter=map_filter
            )
        elif format == 'long':
            gen = file_line_generator(
                file, format=[2, 3, 6, 7, 9, 12], chrom=chromosome, header=0,
                gzip=gzip, resolution=resolution,
                resolution_adjust=resolution_adjust,
                mapping_filter=map_filter
            )
        elif format == '4dn':
            gen = file_line_generator(
                file, format=[2, 3, 4, 5], chrom=chromosome, header=2,
                gzip=gzip, resolution=resolution,
                resolution_adjust=resolution_adjust, mapping_filter=0
            )
        elif format is None or format == 'customized':
            if custom_format is None:
                raise ValueError('Please provide file format!')
            else:
                if isinstance(custom_format, int):
                    custom_format = [int(elm) for elm in str(custom_format)]
                gen = file_line_generator(
                    file, format=custom_format, header=header,
                    chrom=chromosome, gzip=gzip, resolution=resolution,
                    resolution_adjust=resolution_adjust,
                    mapping_filter=map_filter
                )
        else:
            raise ValueError('Unrecognized format: ' + format)

        mat = np.zeros((size, size))
        count = 0
        for p1, p2, val in gen:
            if count==0: #?
                p1_m=p1#?
                p2_m=p2#?
            count += 1
            p1-=p1_m#?
            p2-=p2_m#?
            if count % 100000 == 0:
                print('Line: ', count)
            # print(chromosome, p1, p2, val)
            mat[p1, p2] += val
            if p1 != p2:
                mat[p2, p1] += val
        
    if operations is not None:
        mat = matrix_operation(mat, operations, **kwargs)

    if keep_n_strata:
        strata = [np.diag(mat[i:, :len(mat)-i]) for i in range(keep_n_strata)]
    else:
        strata = None

    if sparse:
        mat = coo_matrix(mat)

    return mat, strata
```
