## Supplementary material for "scHiCTools: a computational toolbox for analyzing single-cell Hi-C data": S4: processing_utils.py

```
import numpy as np
from scipy.signal import convolve2d


def matrix_operation(mat, operations, **kwargs):
    """
    

    Parameters
    ----------
    mat : numpy.ndarray
        Matrix to apply smoothing operators.
        
    operations : list[str]
        A list of smoothing operators to apply.
        Now support perators: 'oe_norm', 'vc_norm', 'vc_sqrt_norm',
        'kr_norm', 'convolution', 'random_walk', 'network_enhancing',
        'logarithm', 'power'.
        
    **kwargs :
        Arguments for specific smoothing operators.
        
        'kr_norm':
            'maximum_error_rate': error rate to stop iteration, default=1e-4.
        'convolution':
            'kernel_shape': shape of kernel ('kernel_shape' ,'kernel_shape'), default=3.
        'random_walk':
            'random_walk_ratio': propotion of random walk smooth matrix, default=1.0;
            't': number of random walk iteration times, default=1.
        'network_enhancing':
            'kNN': number of nearest neighbors, default=20;
            'iterations': number of iterations, default=1.
        'logarithm':
            'epsilon': numerator of error term, default=1;
            'log_base': denominator of error's exponentiation, default=e.
        'power':
            'pow': exponent of exponentiation, default=0.5.
        

    Returns
    -------
    mat : numpy.ndarray
        Matrix after appling smoothing operator.

    """
    
    for op in operations:
        op = op.lower()
        
        if op == 'oe_norm':
            new_mat = np.zeros(mat.shape)
            averages = np.array([np.mean(mat[i:, :len(mat) - i]) for i in range(len(mat))])
            averages = np.where(averages == 0, 1, averages)
            for i in range(len(mat)):
                for j in range(len(mat)):
                    d = abs(i - j)
                    new_mat[i, j] = mat[i, j] / averages[d]
            mat = new_mat
            
        elif op == 'vc_norm':
            sm = np.sum(mat, axis=0)
            sm = np.where(sm == 0, 1, sm)
            sm_v = np.tile(sm, (len(sm), 1))
            sm_c = sm_v.T
            mat = mat / sm_c / sm_v
        
        elif op == 'vc_sqrt_norm':
            sm = np.sum(mat, axis=0)
            sm = np.where(sm == 0, 1, sm)
            sm = np.sqrt(sm)
            sm_v = np.tile(sm, (len(sm), 1))
            sm_c = sm_v.T
            mat = mat / sm_c / sm_v
            
        elif op == 'kr_norm':
            mat = KR_norm(mat, kwargs.pop('maximum_error_rate', 1e-4))
            
        elif op == 'convolution':
            mat = convolution(mat, kwargs.pop('kernel_shape', 3))
            
        elif op == 'random_walk':
            mat = random_walk(mat, kwargs.pop('random_walk_ratio', 1.0),kwargs.pop('t', 1))
            
        elif op == 'network_enhancing':
            mat = network_enhancing(mat, kwargs.pop('kNN', 20),
                                    kwargs.pop('iterations', 1), kwargs.pop('alpha', 0.9))
        
        elif op == 'logarithm':
            mat = np.log(mat + kwargs.pop('epsilon', 1)) / np.log(kwargs.pop('log_base', np.e))
        
        elif op == 'power':
            mat = np.power(mat, kwargs.pop('pow', 0.5))
        
    return mat


'''
def OE_norm(mat):
    new_mat = np.zeros(mat.shape)
    averages = np.array([np.mean(mat[i:, :len(mat) - i]) for i in range(len(mat))])
    averages = np.where(averages == 0, 1, averages)
    for i in range(len(mat)):
        for j in range(len(mat)):
            d = abs(i - j)
            new_mat[i, j] = mat[i, j] / averages[d]
    return new_mat


def VC_norm(mat):
    sm = np.sum(mat, axis=0)
    sm = np.where(sm == 0, 1, sm)
    sm_v = np.tile(sm, (len(sm), 1))
    sm_c = sm_v.T
    new_mat = mat / sm_c / sm_v
    return new_mat


def VC_SQRT_norm(mat):
    sm = np.sum(mat, axis=0)
    sm = np.where(sm == 0, 1, sm)
    sm = np.sqrt(sm)
    sm_v = np.tile(sm, (len(sm), 1))
    sm_c = sm_v.T
    new_mat = mat / sm_c / sm_v
    return new_mat
'''


def KR_norm(mat, maximum_error_rate=1e-4):
    bias = np.mean(mat) * maximum_error_rate
    # Remove all-zero rows and columns
    sm = np.sum(mat, axis=0)
    zeros = []
    for i in range(len(sm)):
        if sm[i] == 0:
            zeros.append(i)
    new_mat = np.delete(mat, zeros, axis=0)
    new_mat = np.delete(new_mat, zeros, axis=1)

    # Iteration
    x = np.random.random(size=len(new_mat))
    k = 0
    while True:
        # I forgot where I found this iteration formula
        # But it does work...
        # I'll check later...
        k += 1
        aa = np.diag(x).dot(new_mat) + np.diag(new_mat.dot(x))
        aa = np.linalg.inv(aa)
        bb = np.diag(x).dot(new_mat).dot(x) - np.ones(x.shape)
        delta = aa.dot(bb)
        new_x = x - aa.dot(bb)

        max_error = np.max(np.abs(delta))
        # print(f'Iteration: {k}, Max Error: {max_error}')
        if max_error < bias:
            break
        else:
            x = new_x

    # Normalization
    dg = np.diag(new_x)
    new_mat = dg.dot(new_mat).dot(dg)

    # Put all-zero rows and columns back
    for zero in zeros:
        new_mat = np.insert(new_mat, zero, 0, axis=0)
        new_mat = np.insert(new_mat, zero, 0, axis=1)
    return new_mat


def convolution(mat, kernel_shape=3):
    conv = np.ones((kernel_shape, kernel_shape)) / (kernel_shape ** 2)
    mat = convolve2d(mat, conv, 'same')
    return mat


def random_walk(mat, random_walk_ratio=1.0,t=1):
    sm = np.sum(mat, axis=1)
    sm = np.where(sm == 0, 1, sm)
    sm = np.tile(sm, (len(mat), 1)).T
    walk = mat / sm
    for i in range(t):
        mat = random_walk_ratio * mat.dot(walk) + (1 - random_walk_ratio) * mat
    return mat


def reduce_sparsity(mat, sparsity_method='log', power=0.5):
    if sparsity_method == 'log':
        return np.log(mat + 1)
    elif sparsity_method == 'power':
        return np.power(mat, power)
    else:
        raise ValueError('Method {0} not supported while reducing sparsity.'.format(sparsity_method))


def network_enhancing(mat, kNN=20, iteration=1, alpha=0.9):
    argsort = np.argsort(-mat, axis=1)
    new_mat = np.zeros(mat.shape)
    for i in range(len(mat)):
        for j in range(kNN):
            pos = argsort[i, j]
            new_mat[i, pos] = mat[i, pos]

    sm = np.sum(new_mat, axis=1)
    sm = np.where(sm == 0, 1, sm)
    sm = np.tile(sm, (len(mat), 1)).T
    walk = new_mat / sm

    for k in range(iteration):
        if k == 0:
            new_mat = alpha * walk.T.dot(mat).dot(walk) + (1 - alpha) * mat
        else:
            new_mat = alpha * walk.T.dot(new_mat).dot(walk) + (1 - alpha) * new_mat
    return mat
```
