## Supplementary material for "scHiCTools: a computational toolbox for analyzing single-cell Hi-C data": S4: setup.py

```
import setuptools
from os import path

this_directory = path.abspath(path.dirname(__file__))
with open(path.join(this_directory, 'README.md'), encoding='utf-8') as f:
    long_description = f.read()


def get_requirements(path):
    with open(path, "r") as fh:
        content = fh.read()
    return [
        req
        for req in content.split("\n")
        if req != '' and not req.startswith('#')
    ]

install_requires = get_requirements('requirements.txt')


setuptools.setup(
    name="scHiCTools",
    version="0.0.3",
    author="Fan Feng",
    author_email="",
    description="A user-friendly package for processing single cell HiC data",
    long_description=long_description,
    long_description_content_type="text/markdown",
    url="https://github.com/liu-bioinfo-lab/scHiCTool",
    packages=setuptools.find_packages(),
    install_requires=install_requires,
    include_package_data = True,
    classifiers=[
        "Programming Language :: Python :: 3",
        "License :: OSI Approved :: MIT License",
        "Operating System :: OS Independent",
    ],
    extras_require={
        "interactive_scatter": ["plotly>=4.8.0"],
    }
)
```
