## Supplementary material for "scHiCTools: a computational toolbox for analyzing single-cell Hi-C data": S4: test_all.py

```
import pytest
import matplotlib.pyplot as plt
import numpy as np

import os
import sys
sys.path.insert(0, os.path.abspath(
    os.path.join(
        os.path.dirname(__file__), '..')))


from scHiCTools import scHiCs
from scHiCTools import scatter

sys.path.insert(0, os.path.abspath(
    os.path.join(os.path.dirname(__file__), '.')))

def test():

    y = scHiCs(['data/cell_03','data/cell_01','data/cell_02'],
                reference_genome='mm9', resolution=100000,
                max_distance=4000000, format='shortest_score',
                adjust_resolution=False, chromosomes='except Y',
                operations=['convolution'], kernel_shape=3, keep_n_strata=10,
                store_full_map=True)

    
    emb1 = y.learn_embedding(similarity_method='innerproduct',
                         return_distance=True,
                         embedding_method='mds',
                         aggregation='median')
    
    emb2=y.learn_embedding(similarity_method='HiCRep',
                       return_distance=True,
                       embedding_method='mds',
                       aggregation='median')
    
    emb3 = y.learn_embedding(similarity_method='Selfish',
                         return_distance=True,
                         embedding_method='mds',
                         aggregation='median')
    
    emb4 = y.learn_embedding(similarity_method='innerproduct',
                         return_distance=True,
                         embedding_method='mds',
                         aggregation='mean')
    
    emb5 = y.learn_embedding(similarity_method='innerproduct',
                         return_distance=True,
                         embedding_method='tSNE',
                         aggregation='median')
    
    # emb6 = y.learn_embedding(similarity_method='innerproduct',
    #                      return_distance=True,
    #                      embedding_method='UMAP',
    #                      aggregation='median',
    #                      print_time=False)
    
    emb7 = y.learn_embedding(similarity_method='innerproduct',
                         return_distance=True,
                         embedding_method= 'phate',
                         aggregation='median',
                         k=2)
    
    emb8 = y.learn_embedding(similarity_method='innerproduct',
                         return_distance=True,
                         embedding_method= 'spectral_embedding',
                         aggregation='median',
                         print_time=False)
    
    label1=y.clustering(n_clusters=2,
                    clustering_method='kmeans',
                    similarity_method='innerproduct',
                    aggregation='median',
                    n_strata=None)
    
    label2=y.clustering(n_clusters=2,
                    clustering_method='spectral_clustering',
                    similarity_method='innerproduct',
                    aggregation='median',
                    n_strata=None)
    
    hicluster=y.scHiCluster(dim=2,cutoff=0.8,n_PCs=10,n_clusters=2)
    
    assert len(set(label1))==2
    assert len(set(label2))==2
    assert len(set(hicluster[1]))==2
    assert emb1[0].shape==(3,2)
    assert emb1[1].shape==(3,3)
    assert emb2[0].shape==(3,2)
    assert emb2[1].shape==(3,3)
    assert emb3[0].shape==(3,2)
    assert emb3[1].shape==(3,3)
    assert emb4[0].shape==(3,2)
    assert emb4[1].shape==(3,3)
    assert emb5[0].shape==(3,2)
    assert emb5[1].shape==(3,3)
    # assert emb6[0].shape==(3,2)
    # assert emb6[1].shape==(3,3)
    assert emb7[0].shape==(3,2)
    assert emb7[1].shape==(3,3)
    assert emb8[0].shape==(3,2)
    assert emb8[1].shape==(3,3)
    
    plt.figure()
    plt.subplot(1,2,1)
    scatter(emb1[0]*100, label=label1)
    plt.subplot(1,2,2)
    scatter(emb2[0])
    
    
    plt.figure()
    scatter(hicluster[0],label=hicluster[1])
    
```
