## Supplementary material for "scHiCTools: a computational toolbox for analyzing single-cell Hi-C data": S4: test_clustering.py

```
# -*- coding: utf-8 -*-

import pytest
import numpy as np
import sys
import os
sys.path.insert(0, os.path.abspath(
    os.path.join(
        os.path.dirname(__file__), '..')))


from scHiCTools import kmeans, spectral_clustering, HAC


center=np.array([[0,0],[100,100],[-100,100]])
rand_data = np.random.normal(size=(90,2)) + np.repeat(center, 30, axis=0)

def test_kmeans():
    label=kmeans(rand_data, k=3)
    l1 = np.equal(label[:30], label[0]).all()
    l2 = np.equal(label[30:-30], label[30]).all()
    l3 = np.equal(label[-30:], label[-30]).all()
    assert l1
    assert l2
    assert l3


def test_SC():
    label=spectral_clustering(rand_data, n_clusters=3)
    l1 = np.equal(label[:30], label[0]).all()
    l2 = np.equal(label[30:-30], label[30]).all()
    l3 = np.equal(label[-30:], label[-30]).all()
    assert l1
    assert l2 
    assert l3
    
def test_HAC():
    label=HAC(rand_data, n_clusters=3)
    l1 = np.equal(label[:30], label[0]).all()
    l2 = np.equal(label[30:-30], label[30]).all()
    l3 = np.equal(label[-30:], label[-30]).all()
    assert l1
    assert l2
    assert l3
    
```
