## Supplementary material for "scHiCTools: a computational toolbox for analyzing single-cell Hi-C data": S4: test_embeding.py

```
# -*- coding: utf-8 -*-


import pytest
import numpy as np
from scipy.spatial import distance
import sys
import os
sys.path.insert(0, os.path.abspath(
    os.path.join(
        os.path.dirname(__file__), '../scHiCTools/embedding')))

from embedding import PCA, MDS, tSNE, SpectralEmbedding, PHATE # , UMAP

rand_data = np.random.normal(size=(60,5))
dist_mat = distance.squareform(distance.pdist(rand_data))
dim=np.random.randint(1,5)

def test_PCA():
    emb = PCA(rand_data, dim=dim)
    assert len(emb[0])==dim
    assert len(emb)==len(rand_data)
    assert not np.any(np.isnan(emb))
    assert not np.any(np.isinf(emb))
    
    
def test_MDS():
    emb = MDS(dist_mat, n=dim)
    assert len(emb[0])==dim
    assert len(emb)==len(dist_mat)
    assert not np.any(np.isnan(emb))
    assert not np.any(np.isinf(emb))
    
    
def test_tSNE():
    emb = tSNE(dist_mat, n_dim=dim)
    assert len(emb[0])==dim
    assert len(emb)==len(dist_mat)
    assert not np.any(np.isnan(emb))
    assert not np.any(np.isinf(emb))
    
    
def test_SpectralEmbedding():
    emb = SpectralEmbedding(dist_mat, dim=dim)
    assert len(emb[0])==dim
    assert len(emb)==len(dist_mat)
    assert not np.any(np.isnan(emb))
    assert not np.any(np.isinf(emb))
    
    
# def test_UMAP():
#     emb = UMAP(dist_mat, dim=dim)
#     assert len(emb[0])==dim
#     assert len(emb)==len(dist_mat)
#     assert not np.any(np.isnan(emb))
#     assert not np.any(np.isinf(emb))
    
    
def test_PHATE():
    emb = PHATE(dist_mat, dim=dim)
    assert len(emb[0])==dim
    assert len(emb)==len(dist_mat)
    assert not np.any(np.isnan(emb))
    assert not np.any(np.isinf(emb))
    
```
