## Supplementary material for "scHiCTools: a computational toolbox for analyzing single-cell Hi-C data": S4: test_load_data.py

```
# -*- coding: utf-8 -*-

import pandas as pd
import numpy as np
import joblib
import matplotlib.pyplot as plt


import sys
import os
sys.path.insert(0, os.path.abspath(
    os.path.join(
        os.path.dirname(__file__), '..')))

from scHiCTools import scHiCs

sys.path.insert(0, os.path.abspath(
    os.path.join(os.path.dirname(__file__), '.')))

print(sys.path[:2])


def test_readhic():
    files = ['data/test.hic']
    x = scHiCs(files, reference_genome='hg19', resolution=2500000,
               max_distance=4000000, format='.hic', adjust_resolution=True,
               chromosomes='all', operations=['convolution'], kernel_shape=3,
               keep_n_strata=10, store_full_map=True)
    assert x.num_of_cells==len(files)


def test_readtxt():
    files = ['data/cell_03','data/cell_01','data/cell_02']
    x = scHiCs(files, reference_genome='mm9', resolution=100000,
               max_distance=4000000, format='shortest_score',
               adjust_resolution=False, chromosomes='except Y',
               operations=['convolution', 'random_walk', 'network_enhancing'], kernel_shape=3, keep_n_strata=10,
               store_full_map=True)
    assert x.num_of_cells==len(files)


def test_readmcool():
    files=['data/test.mcool']
    y = scHiCs(files, reference_genome='hg19', resolution=2500000,
                max_distance=4000000, format='mcool', adjust_resolution=True,
                chromosomes='all', operations=['convolution'],
                kernel_shape=3, keep_n_strata=10, store_full_map=True)
    assert y.num_of_cells==len(files)

def test_select_cells():
    files = ['data/cell_03','data/cell_01','data/cell_02']
    x = scHiCs(files, reference_genome='mm9', resolution=50000,
               max_distance=4000000, format='shortest_score',
               adjust_resolution=True, chromosomes='except Y',
               operations=['convolution'], kernel_shape=3, keep_n_strata=10,
               store_full_map=False)
    x.plot_contacts()
    x.select_cells(min_n_contacts=240000,max_short_range_contact=0.583)
    assert (x.contacts>240000).all()
    assert (x.short_range/x.contacts<0.583).all()
```
