## Supplementary material for "scHiCTools: a computational toolbox for analyzing single-cell Hi-C data": S4: test_smoothing.py

```
import pandas as pd
import numpy as np
import joblib
import matplotlib.pyplot as plt


import sys
import os
sys.path.insert(0, os.path.abspath(
    os.path.join(
        os.path.dirname(__file__), '..')))

from scHiCTools import scHiCs

sys.path.insert(0, os.path.abspath(
    os.path.join(os.path.dirname(__file__), '.')))

print(sys.path[:2])

def test_smoothing():
    files = ['data/cell_03','data/cell_01','data/cell_02']
    x = scHiCs(files, reference_genome='mm9', resolution=100000,
               max_distance=4000000, format='shortest_score',
               adjust_resolution=False, chromosomes='except Y',
               operations=['convolution', 'random_walk', 'network_enhancing'], kernel_shape=3, keep_n_strata=10,
               store_full_map=True)
    x.processing('convolution')
    x.processing('random_walk')
    x.processing('network_enhancing')
    assert x.num_of_cells==len(files)
```
